# Overexpression of *Thalassiosira pseudonana* violaxanthin de-epoxidase-like 2 (VDL2) increases fucoxanthin while stoichiometrically reducing diadinoxanthin cycle pigment abundance

**DOI:** 10.1101/2020.01.06.896316

**Authors:** Olga Gaidarenko, Dylan W. Mills, Maria Vernet, Mark Hildebrand

## Abstract

Despite the ubiquity and ecological importance of diatoms, much remains to be understood about their physiology and metabolism, including their carotenoid biosynthesis pathway. Early carotenoid biosynthesis steps are well-conserved, while the identity of the enzymes that catalyze the later steps and their order remain unclear. Those steps lead to the biosynthesis of the final pathway products: the main accessory light-harvesting pigment fucoxanthin (Fx) and the main photoprotective pigment pool comprised of diadinoxanthin (Ddx) and its reversibly de-epoxidized form diatoxanthin (Dtx). We used sequence comparison to known carotenoid biosynthesis enzymes to identify novel candidates in the diatom *Thalassiosira pseudonana*. Microarray and RNA-seq data was used to select candidates with transcriptomic responses similar to known carotenoid biosynthesis genes and to create full-length gene models, and we focused on those that encode proteins predicted to be chloroplast-localized. We identified a violaxanthin de-epoxidase-like gene (Thaps3_11707, VDL2) that when overexpressed results in increased Fx abundance while stoichiometrically reducing Ddx+Dtx. Based on transcriptomics, we hypothesize that Thaps3_10233 may also contribute to Fx biosynthesis, in addition to VDL2. Separately using antisense RNA to target VDL2, VDL1, and both LUT1-like copies (hypothesized to catalyze an earlier step in the pathway) simultaneously, reduced the overall cellular photosynthetic pigment content, including chlorophylls, suggesting destabilization of light-harvesting complexes by Fx deficiency. Based on transcriptomic and physiological data, we hypothesize that the two predicted *T. pseudonana* zeaxanthin epoxidases have distinct functions and that different copies of phytoene synthase and phytoene desaturase may serve to initiate carotenoid biosynthesis in response to different cellular needs. Finally, nine carotene cis/trans isomerase (CRTISO) candidates identified based on sequence identity to known CRTISO proteins were narrowed to two most likely to be part of the *T. pseudonana* carotenoid biosynthesis pathway based on transcriptomic responses and predicted chloroplast targeting.

## INTRODUCTION

Diatoms are incredibly diverse, environmentally flexible, and productive photosynthetic microalgae that belong to the Stramenopile or Heterokont class. There are over 100,000 estimated diatom species that occupy a wide variety of habitats. Diatoms are of great ecological importance, as they are responsible for approximately 40% of marine primary productivity and play a major role in the global cycling of carbon, nitrogen, phosphorus, and silicon [Hildebrand et al. 2012, Wilhelm et al. 2006]. They have a complex evolutionary history that includes a secondary endosymbiotic event, wherein an ancient red alga was engulfed by a heterotrophic eukaryote, at least 800Ma ago. Because of this event, heterokonts, including diatoms, possess physiological and metabolic features that differ from other algal groups, such as the more extensively studied chlorophytes [Hildebrand et al. 2012, Wilhelm et al. 2006].

Photosynthetic pigments are among the distinguishing features of diatoms. Carotenoids are utilized by all known photosynthetic organisms, and the initial part of the carotenoid biosynthesis pathway from phytoene to β-carotene (β-car) (Fig. 1) is well-conserved [Bertrand 2010, Coesel et al. 2008]. β-car is synthesized from lycopene, as is α-carotene (α-car) in many organisms. Both β- and α-car have a variety of derivatives that can differ substantially depending on the taxonomic class, sometimes with further inter-species differences. Diatoms do not possess the α-car branch of carotenoids, and their main photoprotective and accessory light-harvesting pigments are β-car derivatives. Knowledge about the sequence of the post-β-car biosynthetic steps in diatoms, as well as the enzymes that catalyze them, is sparse [Bertrand 2010, Coesel et al. 2008]. One of the final products of the pathway is fucoxanthin (Fx), the main accessory light-harvesting pigment in diatoms that is responsible for their characteristic golden-brown color. It is bound to photoantenna proteins along with chlorophylls a and c (Chl a, Chl c). Most diatoms have Chl c1 and Chl c2, although Chl c3 has been found in some species as well. Unlike chlorophytes, diatoms do not make chlorophyll b [Wilhelm et al. 2006]. Diadinoxanthin (Ddx) and diatoxanthin (Dtx) are the other final products of the diatom carotenoid biosynthesis pathway. They form a xanthophyll cycle by reversibly interconverting via epoxidation and de-epoxidation (Fig. 1), comprising the major photoprotective mechanism employed by diatoms [Goss et al. 2006, Lohr and Wilhelm 1999, Wilhelm et al. 2006]. Some Ddx molecules are bound by photoantenna proteins, while others are dissolved in the lipid shield that surrounds the photoantennae [Lepetit et al. 2010]. The other xanthophyll cycle found in diatoms is the violaxanthin (Vx) cycle, in which zeaxanthin (Zx) is reversibly converted to (Vx) via antheraxanthin (Ax) by two epoxidation steps. Both cycles serve to allow the switch between harvesting (using epoxidized forms of the pigments) and dissipation (using de-epoxidized forms of the pigments) of light energy as a rapid adjustment to irradiance changes. The Vx cycle is conserved and plays a major photoprotective role in chlorophytes. While functional in diatoms, its pigments are present in small amounts compared to other photosynthetic pigments, and mainly serve as precursors to the Ddx cycle pigments and Fx [Coesel et al. 2008, Lohr and Wilhelm 1999].

**Fig. 1.**
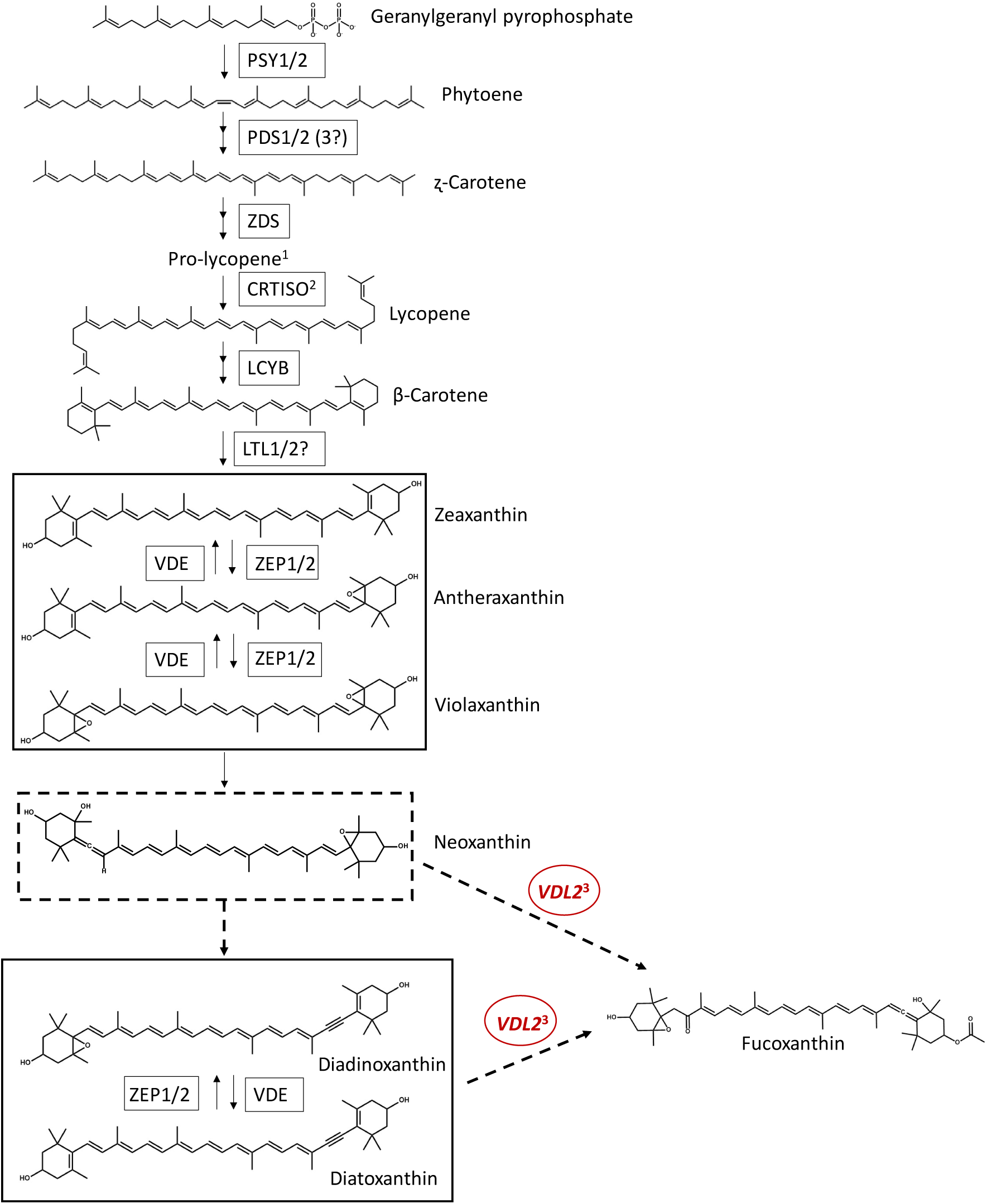
Putative carotenoid biosynthesis pathway in *T. pseudonana.* VDL2 is putatively placed based on results of this study. ^1^Pro-lycopene is a stereoisomer of lycopene. ^2^Thaps3_21900 is hypothesized to catalyze this step. Thaps3_10233 might also be involved. ^3^Thaps3_10233 might also be involved. PSY = phytoene synthase, PDS = phytoene desaturase, ZDS = ʐ-carotene desaturase, CRTISO = carotene cis/trans isomerase, prolycopene isomerase, LCYB = lycopene cyclase b, LTL = LUT-like, ZEP = zeaxanthin epoxidase, VDE = violaxanthin de-epoxidase, VDL = VDE-like. Dashed arrows and box represent steps yet to be experimentally confirmed.

There are two major hypotheses regarding biosynthesis of the Ddx+Dtx pool and Fx from Vx (Fig. 1). Ddx+Dtx and Fx may have a common precursor, or Fx may be synthesized from Ddx+Dtx. Pigment flux-based evidence supports the latter hypothesis [Goericke and Welschmeyer 1992, Lohr and Wilhelm 1999], but it is yet to be directly tested by genetic manipulation, and enzymes involved in the biosynthetic steps have not been identified. Furthermore, Phaeophyceae and Chrysophyceae contain Fx but not Ddx cycle pigments, and therefore Ddx+Dtx is not an obligate precursor of Fx [Lohr and Wilhelm 1999]. Neoxanthin has been hypothesized as an intermediate between Vx and (Ddx+Dtx) + Fx based on its structure. Although not typically observed in diatom pigment extracts, it was detected in an enriched fraction from the diatom *Phaeodactylum tricornutum* by Dambeck et al. [2012].

The sequencing of diatom genomes and development of genetic manipulation tools have facilitated the study of diatom carotenoid biosynthesis. Putative carotenoid biosynthesis gene candidates have been identified in the first two available diatom genomes, *Thalassiosira pseudonana* [Armbrust et al. 2004] and *P. tricornutum* [Bowler et al. 2008], by Coesel et al. [2008] and Dambeck et al. [2012], based on sequence identity of their protein products to those of known carotenoid biosynthesis genes in other organisms. Functional complementation confirming enzyme functions for the pre-β-car part of the pathway was performed by Dambeck et al. [2012]. Lavaud et al. [2012] silenced the *P. tricornutum* gene that encodes Vx de-epoxidase (VDE), which performs de-epoxidation in both xanthophyll cycles, resulting in a Dtx de-epoxidation deficiency (Vx de-epoxidation was not examined). Phytoene synthase (PSY), which catalyzes the first committed step of carotenoid biosynthesis, has been silenced [Kaur and Spillane 2014] and overexpressed [Kadono et al. 2015] in *P. tricornutum*, resulting in reduced and increased cellular carotenoid content, respectively. Nevertheless, much remains to be understood about the post-β-car part of the carotenoid biosynthesis pathway in diatoms. In this study, we confirm and expand the list of putative carotenoid biosynthesis gene candidates using sequence identity-based reciprocal probing of the *T. pseudonana* and *P. tricornutum* genomes. We then focus on *T. pseudonana* and use available transcriptomic and physiological data to narrow the carotenoid biosynthesis gene candidate list and form hypotheses about the functions of their protein products. Furthermore, we directly test the function of several enzymes hypothesized to be involved in diatom carotenoid biosynthesis through genetic manipulation.

## RESULTS

### Identification of Carotenoid Biosynthesis Gene Candidates Based on Sequence Identity

The genomes of *T. pseudonana* and *P. tricornutum*, accessed on the Department of Energy Joint Genome Institute (DOE JGI) website [Grigoriev et al. 2012, Nordberg et al. 2014], were searched for genes that encode proteins with sequence similarity to those known to participate in carotenoid biosynthesis in both organisms, in order to confirm what has been previously reported [Coesel et al. 2008, Dambeck et al. 2012] and identify new candidates. Some Basic Local Alignment Search Tool (BLAST) results indicated previously unreported gene models that overlap with previously published ones. The latter likely represent outdated models that have since been replaced. Large chromosomal pieces, sometimes encompassing genes identified as part of the carotenoid biosynthesis pathway or those found using the latter as BLAST queries, also frequently showed up in BLAST results. They may contain multiple partial sequence matches, possibly including those of domains that are present in, but not necessarily limited to, enzymes of the carotenoid biosynthesis pathway. Some open reading frames without available gene models were also found. The findings are detailed in **Appendix A**, and the most current model IDs of known carotenoid biosynthesis genes and corresponding BLAST hits for which gene models are available are summarized in Table 1. The unnamed enzymes are not listed in a specific order and no relationship between those listed in the same row but different columns is implied. The putative diatom carotenoid biosynthesis pathway is depicted in Fig. 1.

**Table 1.** Model IDs of known carotenoid biosynthesis genes/enzymes and corresponding BLAST results.

| Sequence Identity BLAST Results For: | Enzyme Name | *T. pseudonana* | *P. tricornutum* |
| --- | --- | --- | --- |
| Phytoene Synthase (PSY) | PSY1 | Thaps3_268908 | Phatr2_56481 |
|  | PSY2 | Thaps3_263269 | ----- |
| Phytoene Desaturase (PDS) | PDS1 | Thaps3_23291 | Phatr2_45735 |
|  | PDS2 | Thaps3_1383 | Phatr2_55102 |
|  | ----- | Thaps3_bd_1474 | ----- |
| ζ-Carotene Desaturase (ZDS) | ZDS | Thaps3_24832 | Phatr2_53974 |
| Carotene Cis-Trans Isomerase<br>(Prolycopene Isomerase) (CRTISO) | ----- | Thaps3_7094 | Phatr2_45243 |
|  | ----- | Thaps3_21900 | Phatr2_9210 |
|  | ----- | Thaps3_21847 | Phatr2_54842 |
|  | ----- | Thaps3_5221 | Phatr2_51868 |
|  | ----- | Thaps3_10233 | Phatr2_54826 |
|  | ----- | Thaps3_11636 | Phatr2_54800 |
|  | ----- | Thaps3_5859 | Phatr2_42980 |
|  | ----- | Thaps3_25361 | ----- |
|  | ----- | Thaps3_10254 | ----- |
| Lycopene β-cyclase (LCYB) | LCYB | Thaps3_270357 | Phatr2_56484 |
| β-Carotene Hydroxylase (BCH) | BCH | Thaps3_263437 | ----- |
| LUT-Like (Lutein Deficient-Like) (LTL) | LTL1 | Thaps3_9541 | Phatr2_50101 |
|  | LTL2 | Thaps3_270336 | Phatr2_26422 |
|  | ----- | Thaps3_33926 | Phatr2_34027 |
|  | ----- | Thaps3_32491 | Phatr2_33568 |
|  | ----- | Thaps3_1549 | Phatr2_6940 |
|  | ----- | Thaps3_264647 | Phatr2_46438 |
|  | ----- | Thaps3_25944 | Phatr2_31339 |
|  | ----- | Thaps3_14875 | Phatr2_47234 |
|  | ----- | Thaps3_4027 | Phatr2_37006 |
|  | ----- | Thaps3_4026 | Phatr2_43466 |
|  | ----- | Thaps3_bd_518 | Phatr2_43467 |
|  | ----- | Thaps3_269400 | Phatr2_43562 |
|  | ----- | Thaps3_264325 | Phatr2_50619 |
|  | ----- | Thaps3_263399 | Phatr2_43469 |
|  | ----- | ----- | Phatr2_32833 |
|  | ----- | ----- | Phatr2_43537 |
| Zeaxanthin Epoxidase (ZEP) | ZEP1 | Thaps3_270370 | Phatr2_45845 |
|  | ZEP2 | Thaps3_261390 | Phatr2_56488 |
|  | ZEP3 | ----- | Phatr2_56492 |
|  | ----- | Thaps3_1961 | Phatr2_43425 |
|  | ----- | Thaps3_6395 | Phatr2_47925 |
|  | ----- | Thaps3_20663 | Phatr2_45936 |
|  | ----- | Thaps3_22671 | ----- |
| Violaxanthin De- Epoxidase (VDE), | VDE | Thaps3_7677 | Phatr2_44635 |
| VDE-Like (VDL), VDE-Related (VDR) | VDL1 | Thaps3_22076 | Phatr2_46155 |
|  | VDL2 | Thaps3_11707 | Phatr2_45846 |
|  | VDR | Thaps3_270211 | Phatr2_56450 |
|  | ----- | ----- | Phatr2_bd_1281 |

### Phylogenetic Analyses of Candidate Carotenoid Biosynthesis Enzymes Based on Sequence Identity

Carotene cis/trans isomerase (CRTISO), LUT1-like (LTL), and zeaxanthin epoxidase (ZEP) candidate searches yielded numerous new candidates. Because many of the publicly available gene models for the LTL group are clearly incomplete (i.e., missing start and/or stop codons), sequence identity analysis was not performed at this stage. However, it should be mentioned that the only BLAST hits for Phatr2_34027 in the *T. pseudonana* genome were Thaps3 9541 (LTL1) and Thaps3 2703365 (LTL2). Most enzymes in this group were functionally annotated as cytochrome p450, with some exceptions. Phatr2_47234, Phatr2_37006, Thaps3_4027, and Thaps3_4026 had no functional annotation available on the DOE JGI website. Phatr2_43563 was functionally annotated as an iron-binding oxidoreductase, Phatr2_50619 chitinase, Phatr2_43537 chitinase with a diacylglycerol binding domain, Phatr2_43469 diacylglycerol acyltransferase, Phatr2_32833 N-acetyltransferase, and Thaps3_14875 glycosyltransferase (UDP-N-acetylglucosamine dolichyl-phosphate N-acetylglucosamine-phosphotransferase).

Sequence identity analysis for the CRTISO group (Fig. 2A**, Appendix B1)** revealed that Thaps3_5221, Thaps3_21847, Thaps3_10233, Thaps3_21900, and Thaps3_7094 are more similar to the *P. tricornutum* enzymes identified by Dambeck et al. [2012] as CRTISO candidates, while Thaps3_25361, Thaps3_11636, Thaps3_5859, and Thaps_10254 grouped closer to each other and were more dissimilar from the former group. However, both groups had members that either had Gene Ontology (GO) terms including carotenoid biosynthetic process, oxidoreductase activity, and flavin adenine dinucleotide (FAD) binding, or did not have GO terms available. All of the proteins had the EuKaryotic Orthologous Groups Identity (KOG ID) “phytoene desaturase.”

**Fig. 2.**
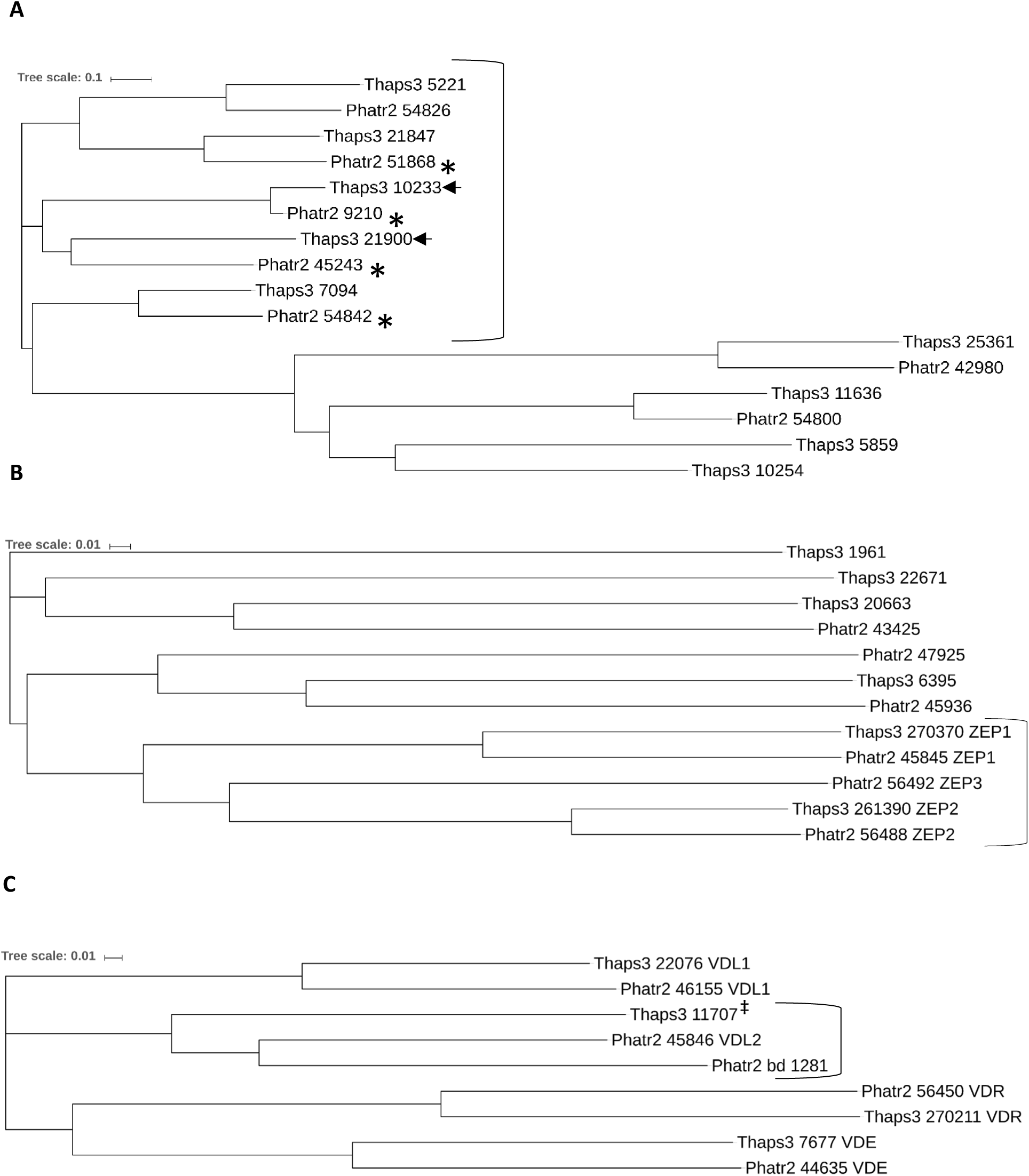
Sequence identity-based phylogenetic trees. Predicted protein sequences were obtained on the DOE JGI website. **A.** CRTISO candidates and related proteins. The bracketed group includes those previously identified as CRTISO candidates by Dambeck et al. [2012], indicated with an asterisk (*). Candidates hypothesized to be involved in carotenoid biosynthesis based on transcriptomic analysis and predicted chloroplast targeting are indicated with an arrow (←). **B.** ZEPs (bracketed) and related proteins. **C.** Known VDEs, VDLs, VDRs, and related proteins. Group including the *P. tricornutum* VDL2 is bracketed. The novel *T. pseudonana* protein Thaps3_11707, hereafter designated VDL2, is indicated with a double dagger (‡).

ZEP candidate sequence identity analysis (Fig. 2B**, Appendix B2)** showed that previously identified ZEPs [Coesel et al. 2008] form a group without additional members. Within the group, the ZEP1s and ZEP2s clustered with their counterparts, and ZEP3 was separate. No terms, predictions, or annotations were available for Phatr2_48545 (ZEP1). All of the other proteins that had a KOG ID were designated as kynurenine 3-monooxygenase and related flavoprotein monooxygenases, and the two that did not were indicated to have a FAD-binding domain (Phatr2_56492, ZEP3) or to be a FAD-dependent oxidoreductase (Phatr2_56488, ZEP2) by Pfam. All of the proteins besides Phatr2_48545 (ZEP1) had GO terms that included monooxygenase activity and cellular aromatic compound metabolic process. Most also included oxidoreductase activity, except for Thaps3_22671 and Phatr2_43425. GO terms for Phatr2_45936 also included small nuclear ribonucleoprotein complex, and for Thaps3_261390 (ZEP2), xanthophyll biosynthetic process.

Sequence identity analysis was also performed on the VDE/VDL/VDR group (Fig. 2C**, Appendix B3)**. Four clusters resulted, with the known VDEs, VDL1s, and VDRs clustering with their counterparts, and the *P. tricornutum* VDL2 in a cluster with the two newly identified proteins, Phatr2_bd_1281 and Thaps3_11707. Based on the sequence similarity, the latter is hereafter designated as the *T. pseudonana* VDL2, which has been previously asserted to be absent from the *T. pseudonana* genome [Coesel et al. 2008].

### Gene Expression Patterns of Carotenoid Biosynthesis Candidates

For all *T. pseudonana* genes listed in Table 1, expression patterns were obtained from the microarray dataset described in Smith et al. [2016]. Briefly, Smith et al. performed a 24-hour silicon starvation time course experiment, with cells placed in silica-free media at 0 h, sampling for a variety of physiological variables as well as the transcriptome throughout. At 4 h, a major portion of cells underwent chloroplast division. There was also evidence of sustained light-induced stress throughout the duration of the experiment. Genes with similar functions appeared to be highly co-regulated, as evidenced by their sorting into clusters with highly similar expression patterns.

Most genes known to participate in *T. pseudonana* carotenoid biosynthesis followed a distinct expression pattern that spiked at 4 h, during chloroplast division, then decreased (Fig. 3A). Some increased expression by 4 h and remained elevated throughout, and ZEP2 was unique in having lower transcript levels throughout the experiment compared to 0 h (Fig. 3B). One half (15 out of 30) of the candidate genes identified based on sequence identity (Table 1) had expression patterns similar to those known to be in the carotenoid biosynthesis pathway (Fig. 3C, D), the other half did not (Fig. 3E, F).

**Fig. 3.**
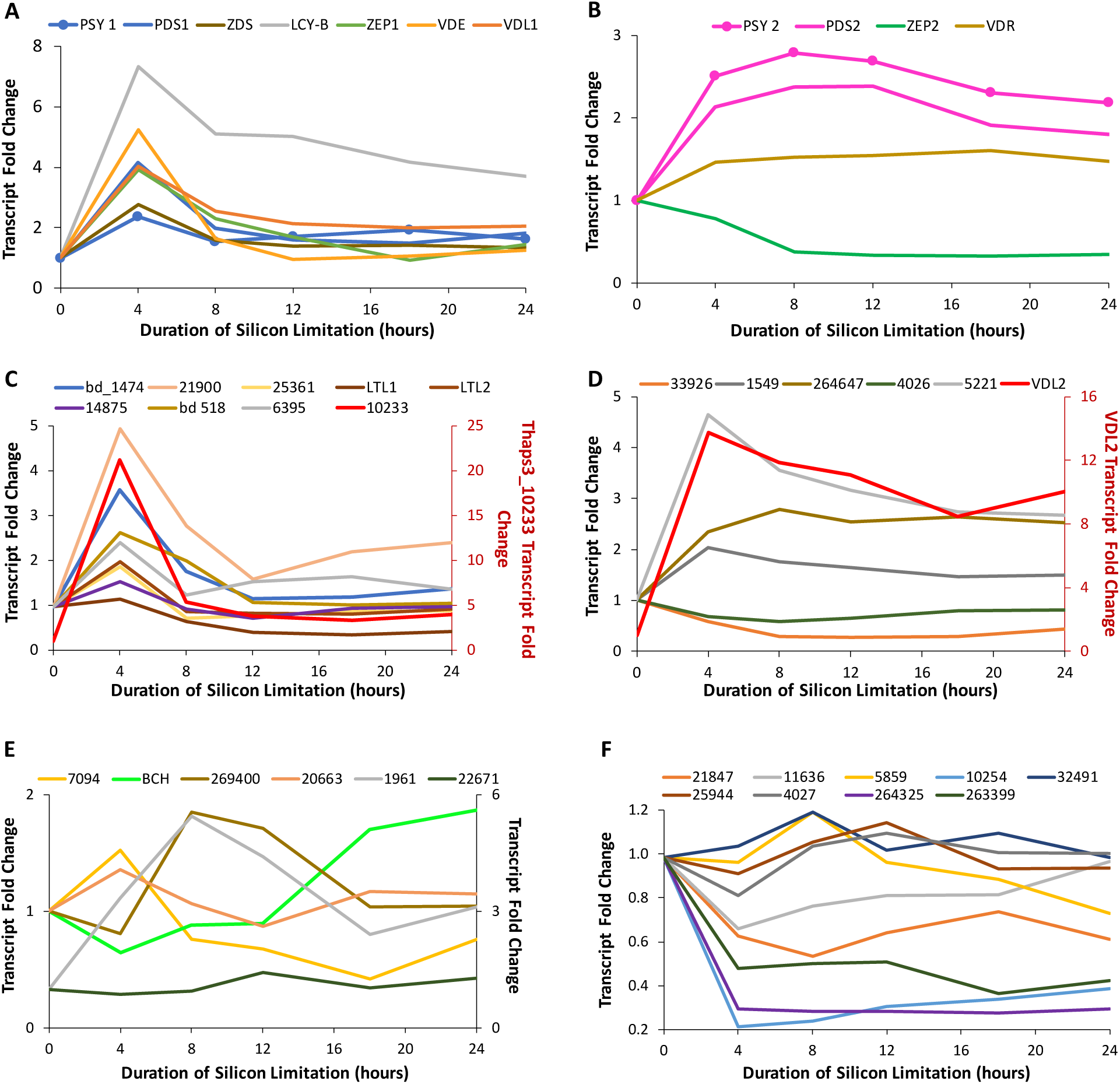
Candidate gene silicon starvation microarray expression patterns [Smith et. al. 2016]. **A, B.** Genes known to be involved in carotenoid biosynthesis. **C, D.** Genes with expression patters similar to those in A and B, chosen for further study. **E, F.** Genes with expression patterns different from those in A and B, not chosen for further study. BCH refers to the BCH-like protein, Thaps3_263437.

Thaps3_17707, the *T. pseudonana* VDL2 identified in this study, and Thaps3_10233, found in the sequence identity-based search for CRTISO candidates, were upregulated to a substantially higher extent than the rest of the carotenoid biosynthesis genes (Fig. 3A-D) and were previously found by Smith et al. [2016] to be in a co-regulated gene cluster with photoantenna protein-encoding genes. We examined the expression patterns of those photoantenna protein genes as well as carotenoid biosynthesis genes using an additional set of transcriptomic data [Abbriano 2017]. This RNA-seq data was obtained during a 9-hour time course experiment after re-addition of silica to silicon-starved *T. pseudonana* cultures, which typically results in a synchronous progression through the cell cycle and thus synchronous chloroplast division. VDL2 and Thaps3_10233 had a distinct co-expression pattern in the silica re-addition RNA-seq dataset as well and were upregulated later and to a higher extent than the rest of the carotenoid biosynthesis genes (Fig. 4A). All but one photoantenna protein-encoding gene that clustered with VDL2 and Thaps3_10233 in the microarray dataset [Smith et al. 2016] also clustered with them in the RNA-seq dataset [Abbriano 2017] (Fig. 4B). Additionally, VDL2 and Thaps3_10233 were found to be co-expressed (hierarchical cluster 871, bootstrapped hierarchical cluster 2194) across a variety of conditions in a work that combined transcriptomic data from multiple studies comprising 380 *T. pseudonana* samples [Ashworth and Ralph 2018].

**Fig. 4.**
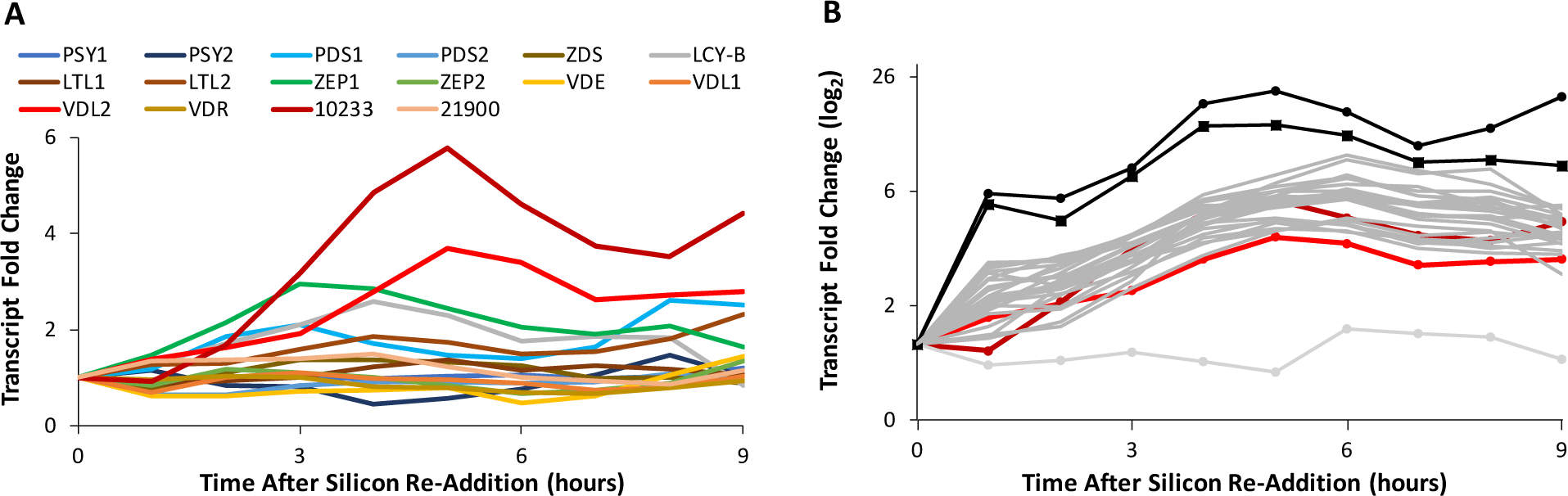
RNA-seq gene expression patterns [Abbriano 2017]. **A.** Carotenoid biosynthesis genes, VDL2 in bright red and Thaps3_10233 in dark red; **B.** VDL2 (bright red), Thaps3_10233 (dark red), and photoantenna protein genes: Thaps3_32723 (light grey, circles), Thaps3_38667 (black, circles), Thaps3_42962 (black, squares), Thaps3_2601, 2845, 3815, 5174, 6139, 7916, 10219, 29375, 30385, 31749, 31983, 33018, 33131, 33606, 34276, 36081, 38122, 39813, 40747, 262313, 262332, 268127, 270092 (dark grey).

### Full-length Gene Models and Predicted Protein Targeting

Because the publicly available gene models are sometimes incomplete/incorrect, full-length gene models **(Appendix C)** were constructed manually by using available RNA-seq data [Abbriano 2017, Smith et al. 2016] for all the genes known to be part of the carotenoid biosynthesis pathway, those that had similar microarray gene expression patterns [Smith et al. 2016], and Thaps3_263437 (β-carotene hydroxylase-like, (BCH-like)), which did not (Fig. 3). Full-length gene models were translated **(Appendix C)**, and the likelihood of chloroplast targeting for the resultant peptides was assessed **(Appendix C)**, summarized in Table 2.

**Table 2.**
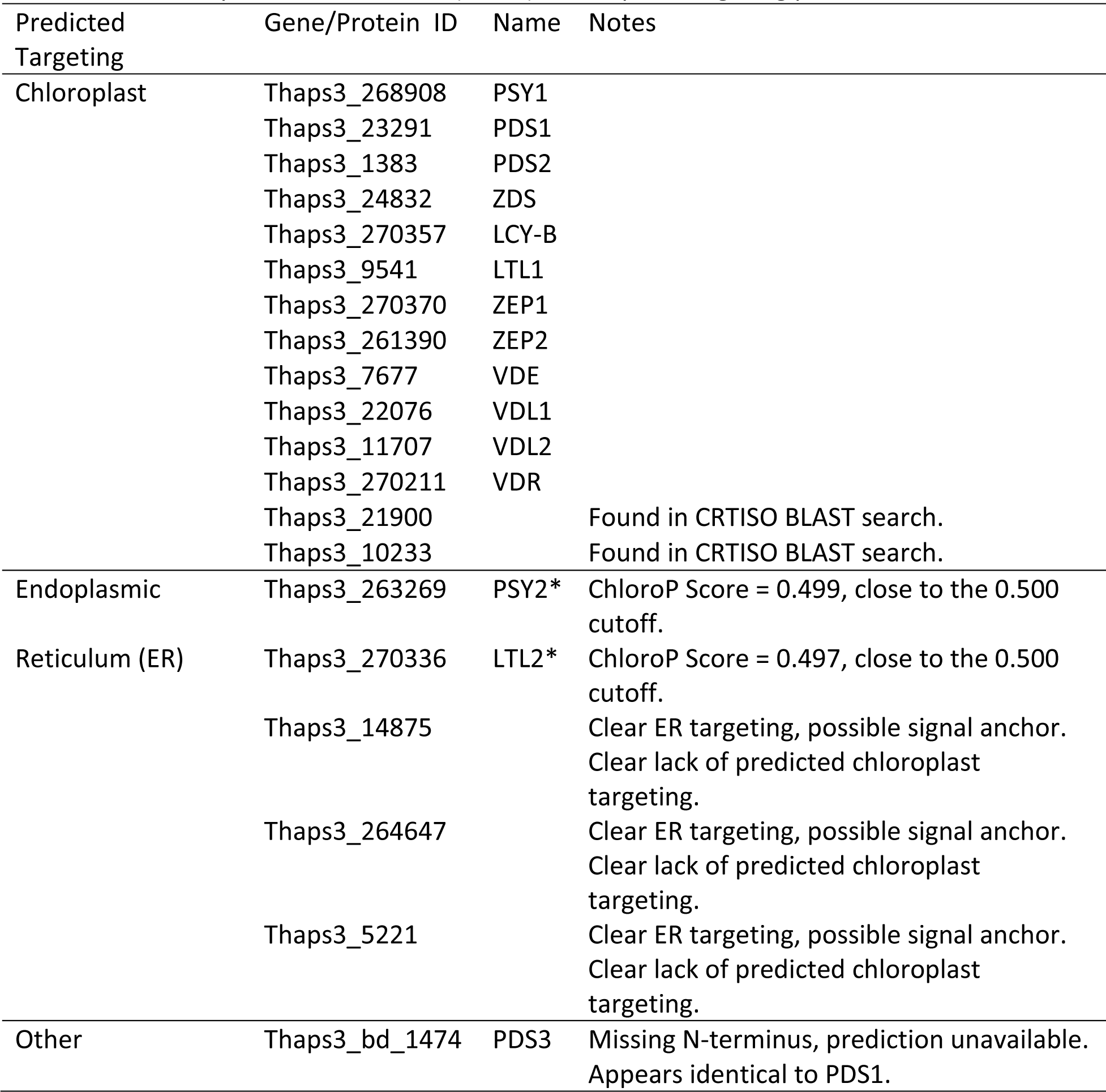
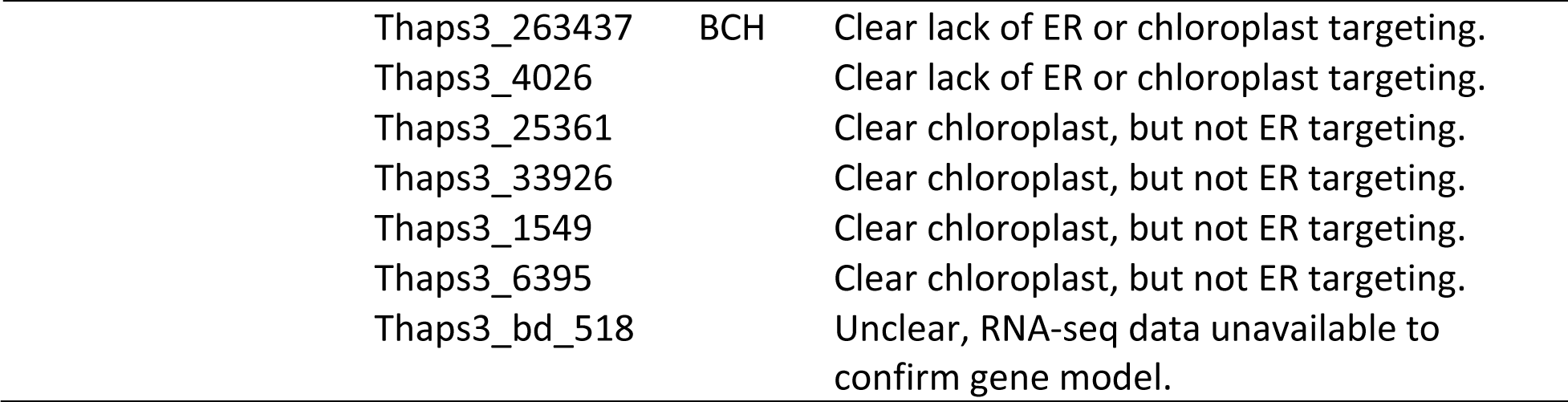
Targeting predictions for known carotenoid biosynthesis enzymes and candidates. *ChloroP score very close to the cutoff (0.500), chloroplast targeting possible.

| Predicted Targeting | Gene/Protein ID | Name | Notes |
| --- | --- | --- | --- |
| Chloroplast | Thaps3_268908 | PSY1 |  |
|  | Thaps3_23291 | PDS1 |  |
|  | Thaps3_1383 | PDS2 |  |
|  | Thaps3_24832 | ZDS |  |
|  | Thaps3_270357 | LCY-B |  |
|  | Thaps3_9541 | LTL1 |  |
|  | Thaps3_270370 | ZEP1 |  |
|  | Thaps3_261390 | ZEP2 |  |
|  | Thaps3_7677 | VDE |  |
|  | Thaps3_22076 | VDL1 |  |
|  | Thaps3_11707 | VDL2 |  |
|  | Thaps3_270211 | VDR |  |
|  | Thaps3_21900 |  | Found in CRTISO BLAST search. |
|  | Thaps3_10233 |  | Found in CRTISO BLAST search. |
| Endoplasmic | Thaps3_263269 | PSY2* | ChloroP Score = 0.499, close to the 0.500 cutoff. |
| Reticulum (ER) | Thaps3_270336 | LTL2* | ChloroP Score = 0.497, close to the 0.500 cutoff. |
|  | Thaps3_14875 |  | Clear ER targeting, possible signal anchor. Clear lack of predicted chloroplast targeting. |
|  | Thaps3_264647 |  | Clear ER targeting, possible signal anchor. Clear lack of predicted chloroplast targeting. |
|  | Thaps3_5221 |  | Clear ER targeting, possible signal anchor. Clear lack of predicted chloroplast targeting. |
| Other | Thaps3_bd_1474 | PDS3 | Missing N-terminus, prediction unavailable. Appears identical to PDS1. |

|  |  |  |
| --- | --- | --- |
| Thaps3_263437 | BCH | Clear lack of ER or chloroplast targeting. |
| Thaps3_4026 |  | Clear lack of ER or chloroplast targeting. |
| Thaps3_25361 |  | Clear chloroplast, but not ER targeting. |
| Thaps3_33926 |  | Clear chloroplast, but not ER targeting. |
| Thaps3_1549 |  | Clear chloroplast, but not ER targeting. |
| Thaps3_6395 |  | Clear chloroplast, but not ER targeting. |
| Thaps3_bd_518 |  | Unclear, RNA-seq data unavailable to confirm gene model. |

### Additional Sequence-Based Analyses

#### Phytoene Desaturase (PDS) and Thaps3_bd_1474

Thaps3_bd_1474, found in the PDS candidate search (Table 1**, Appendix A)**, is part of the unmapped sequence assembly of the *T. pseudonana* genome and is truncated on the 5’ end due to the way those sequences were assembled. The unmapped sequence assembly is also referred to as the “bottom drawer,” and “_bd_” is added to the identification numbers of genes that are part of it. RNA-seq data is not available for the “bottom drawer” sequences and could not be used to construct a full-length gene model. The DOE JGI predictions were used instead **(Appendix D)**. The DOE JGI-predicted peptides for PDS1 and Thaps3_bd_1474 are identical where they align. The latter is approximately 200 amino acids shorter on the N-terminal side and approximately 120 amino acids longer on the C-terminal side. The PDS1 gene contains no introns, and one intron is predicted by DOE JGI for Thaps3_bd_1474. If the predicted intron is disregarded and the sequence is translated as one open reading frame **(Appendix D)**, the resulting peptide is identical to PDS1, except for the N-terminal truncation due to assembly **(Appendix D)**. When genomic DNA sequences that include the DOE JGI models for PDS1 and Thaps3_1474 and 10 kb downstream are examined, they appear identical where the gene models align and for approximately 1.8 kb downstream, with occasional single base pair differences that may be attributed to naturally occurring polymorphisms, and then abruptly become substantially different thereafter **(Appendix D)**. The identical region downstream PDS1 and Thaps3_bd_1474 includes two more genes, one predicted to encode the ribosomal protein L33 (Thaps3_41211 and Thaps3_bd_1472), and the other predicted to encode a DNA methylase (Thaps3_6523 and Thaps3_bd_1109). Polymerase chain reaction (PCR) amplification followed by Sanger sequencing confirmed that both loci are present in the *T. pseudonana* genome. By contrast to the apparent complete sequence identity between PDS1 and the available portion of Thaps3_bd_1474, PDS1 and PDS2 have 62.9% sequence identity **(Appendix D)**.

### Thaps3_263437 (BCH-like)

Approximately one half of the *T. pseudonana* protein, on the N-terminal side, aligns with known non-heme di-iron BCH sequences from other organisms **(Appendix D)**. No known motifs or conserved domains based on sequence or predicted structure were found in the C-terminal half of Thaps3_263437. Portions of it did exhibit limited sequence identity to unknown proteins from diverse organisms (Table 3). Neither the N-terminal BCH-like half nor the unknown C-terminal half of Thaps3_263437 were found in the other currently available diatom genomes (*Phaeodactylum tricornutum, Cyclotella cryptica*, *Thalassiosira oceanica*, *Fragilariopsis cylindrus*, *Pseudo-nitzschia multiseries*).

**Table 3.** Partial sequence identity matches for the Thaps3_263437 C-terminal, non-BCH-like peptide.

| Portion of C-terminal part | BLAST Result Type | Sequence Identity | Organism |
| --- | --- | --- | --- |
| 81-88% | Unidentified or hypothetical proteins | 28-39% | *Chrysochromulina* sp. CCMP291 (haptophyte)<br>*Aurantiochytrium* sp. FCC1311 (labyrinthulomycete)<br>*Oikopleura dioica* (tunicate) |
| 26-37%, C-terminal | Translated nucleotide sequences | 41-43% | *Oikopleura dioica* (tunicate)<br>*Homo sapiens* (primate)<br>*Pan troglodytes* (primate) |
| 17%, middle | Translated nucleotide sequence | 38% | *Arachis hypogaea* (green plant) |

#### Functional Annotation Predictions for the T. pseudonana Carotenoid Biosynthesis Enzymes

Sequence-based functional annotation predictions were compiled for the enzymes of the *T. pseudonana* carotenoid biosynthesis pathway (Table 4). Additionally, predicted protein structures for VDL2, Thaps3_10233, and the C-terminal half of Thaps3_263437 were analyzed for functional predictions. The analysis yielded no additional information.

**Table 4.** Sequence-based functional annotation of *T. pseudonana* carotenoid biosynthesis enzymes.

| Name/ID | Gene Ontology (GO) | EuKaryotic Orthologous Group (KOG) | Protein Family (Pfam) | InterPro Family |
| --- | --- | --- | --- | --- |
| PSY1<br>Thaps3_268908 | GO:0009058<br>biosynthetic process | KOG1459 Squalene synthetase | PF00494<br>Squalene/phytoene synthase | IPR002060<br>Squalene/phytoene synthase |
|  | GO:0016740<br>transferase activity |  |  | IPR008949<br>Isoprenoid synthase domain superfamily |
| PSY2<br>Thaps3_263269 | GO:0003824<br>catalytic activity | KOG1459 Squalene synthetase | PF00494<br>Squalene/phytoene synthase | IPR002060<br>Squalene/phytoene synthase |
|  | GO:0008299<br>isoprenoid biosynthetic process<br>GO:0009058<br>biosynthetic process<br>GO:0009507 chloroplast<br>GO:0016117 carotenoid biosynthetic process<br>GO:0016740 transferase activity |  |  | IPR008949<br>Isoprenoid synthase domain superfamily |
| PDS1<br>Thaps3_23291 | GO:0004497<br>monooxygenase activity | KOG0029 Amine oxidase | PF01593 Flavin-containing amine oxidase | IPR014102<br>Phytoene desaturase |
|  | GO:0009507 chloroplast |  |  | IPR036188<br>FAD/NAD(P)-binding domain superfamily |
|  | GO:0016117 carotenoid biosynthetic process<br>GO:0016491<br>oxidoreductase activity |  |  | IPR002937 Amine oxidase |
| Thaps3_bd_1474 | GO:0016491<br>oxidoreductase activity | KOG0029 Amine oxidase | PF01593 Flavin-containing amine oxidase | IPR036188<br>FAD/NAD(P)-binding domain superfamily<br>IPR002937 Amine oxidase |
| PDS2<br>Thaps3_1383 | GO:0009507 chloroplast | KOG0029 Amine oxidase | PF01593 Flavin-containing amine oxidase | IPR014102<br>Phytoene desaturase |
|  | GO:0016117 carotenoid biosynthetic process |  |  | IPR036188 FAD/NAD(P)-binding domain superfamily |
|  | GO:0016120 carotene biosynthetic process |  |  |  |
|  | GO:0016166 phytoene dehydrogenase activity<br>GO:0016491 oxidoreductase activity |  |  | IPR002937 Amine oxidase |
| ZDS<br>Thaps3_24832 | GO:0016117 carotenoid biosynthetic process | KOG0029 Amine oxidase | PF01593 Flavin-containing amine oxidase | IPR014103<br>Zeta-carotene desaturase |
|  | GO:0016491 oxidoreductase activity |  |  | IPR036188<br>FAD/NAD(P)-binding domain superfamily |
|  | GO:0016719 carotene 7,8-desaturase activity |  |  | IPR002937 Amine oxidase |
| CRTISO<br>Thaps3_21900 | GO:0016117 carotenoid biosynthetic process | KOG4254 Phytoene desaturase | PF13450 NAD(P)-binding Rossmann-like domain | IPR036188<br>FAD/NAD(P)-binding domain superfamily |
|  | GO:0016705 oxidoreductase activity, acting on paired donors, with incorporation or reduction of molecular oxygen<br>GO:0050660 flavin adenine dinucleotide binding |  |  |  |
| Thaps3_10233 | GO:0015036 disulfide oxidoreductase activity | KOG4254 Phytoene desaturase | PF13450 NAD(P)-binding Rossmann-like domain | IPR036188<br>FAD/NAD(P)-binding domain superfamily |
|  | GO:0016117 carotenoid biosynthetic process | PF01593 Flavin-containing amine oxidase |  | IPR002937 Amine oxidase |
|  | GO:0016491 oxidoreductase activity<br>GO:0016705 oxidoreductase activity, acting on paired donors, with incorporation or reduction of molecular oxygen<br>GO:0050660 flavin adenine dinucleotide binding |  |  |  |
| LCY-B<br>Thaps3_270357 | GO:0008152 metabolic process<br>GO:0009507 chloroplast | N/A | PF05834 Lycopene cyclase | IPR010108 Lycopene cyclase, beta/epsilon<br>IPR036188 FAD/NAD(P)-binding domain superfamily |
|  | GO:0016117 carotenoid biosynthetic process<br>GO:0016491 oxidoreductase activity<br>GO:0016705 oxidoreductase activity, acting on paired donors, with incorporation or reduction of molecular oxygen |  |  |  |
| LTL1<br>Thaps3_9541 | GO:0004497 monooxygenase activity | N/A | PF00067 Cytochrome P450 | IPR001128 Cytochrome P450<br>IPR002401 Cytochrome P450, E-class, group I |
| LTL2<br>Thaps3_270336 | GO:0004497 monooxygenase activity<br>GO:0005506 iron ion binding<br>GO:0020037 heme binding | N/A | PF00067 Cytochrome P450 | IPR001128 Cytochrome P450<br>IPR002401 Cytochrome P450, E-class, group I<br>IPR036396 Cytochrome P450 superfamily |
| ZEP1<br>Thaps3_270370 | GO:0004497 monooxygenase activity | KOG2614 Kynurenine 3-monooxygenase | PF01494 FAD binding domain | IPR036188 FAD/NAD(P)-binding domain superfamily |
|  | GO:0006118 obsolete<br>electron transport<br>GO:0006725 cellular<br>aromatic compound<br>metabolic process<br>GO:0008152 metabolic<br>process<br>GO:0009507 chloroplast<br>GO:0016491 oxidoreductase<br>activity | and related<br>flavoprotein<br>monooxygenases |  | IPR002938 FAD-binding domain |
| ZEP2<br>Thaps3_261390 | GO:0004497 monooxygenase<br>activity<br>GO:0006725 cellular<br>aromatic compound<br>metabolic process<br>GO:0008152 metabolic<br>process<br>GO:0016123 xanthophyll<br>biosynthetic process<br>GO:0016491 oxidoreductase<br>activity | KOG2614 Kynurenine<br>3-monooxygenase<br>and related<br>flavoprotein<br>monooxygenases | PF01494 FAD<br>binding domain | IPR036188<br>FAD/NAD(P)-binding domain superfamily<br>IPR002938 FAD-binding domain<br>IPR003953<br>FAD-dependent oxidoreductase 2,<br>FAD binding domain |
| VDE<br>Thaps3_7677 | GO:0055114 oxidation-reduction process<br>GO:0046422 violaxanthin de-epoxidase activity<br>GO:0009507 chloroplast | N/A | PF07137 VDE<br>lipocalin domain | IPR010788 VDE lipocalin domain<br>IPR012674 Calycin |
| VDL1<br>Thaps3_22076 | GO:0055114 oxidation-reduction process<br>GO:0046422 violaxanthin de-epoxidase activity<br>GO:0009507 chloroplast | N/A | PF07137 VDE lipocalin domain | IPR010788 VDE lipocalin domain<br>IPR012674 Calycin |
| VDL2<br>Thaps3_11707 | GO:0055114 oxidation-reduction process<br>GO:0046422 violaxanthin de-epoxidase activity<br>GO:0009507 chloroplast | KOG4157 β-1,6-N-acetylglucosaminyl-transferase, contains WSC domain | PF07137 VDE lipocalin domain | IPR010788 VDE lipocalin domain<br>IPR012674 Calycin |
| VDR<br>Thaps3_270211 | N/A | N/A | N/A | IPR012674 Calycin |

#### Identification of Carotenoid Biosynthesis Genes in Additional Diatom Species

Orthologs of the carotenoid biosynthesis genes identified in *T. pseudonana* and *P. tricornutum* were found in the four other currently available diatom genomes (Table 5). Where gene models were not available, genomic coordinates were given. Three of the *T. oceanica* genes appeared to be split into two adjacent gene models. Those were listed together, separated by a plus sign (+). The number of copies of each gene varied between species, with the most for several genes found in *C. cryptica* (Table 5).

**Table 5.**
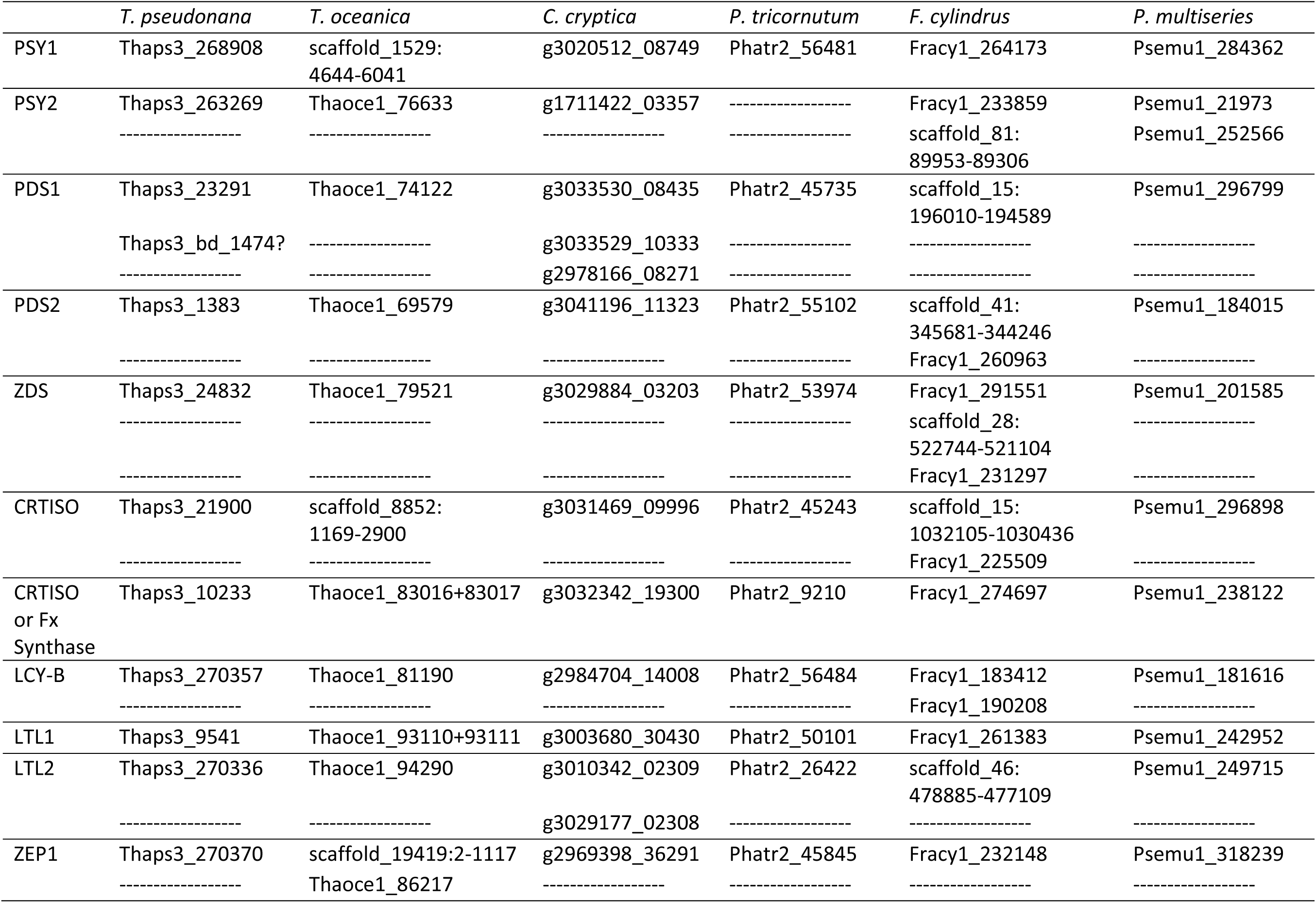
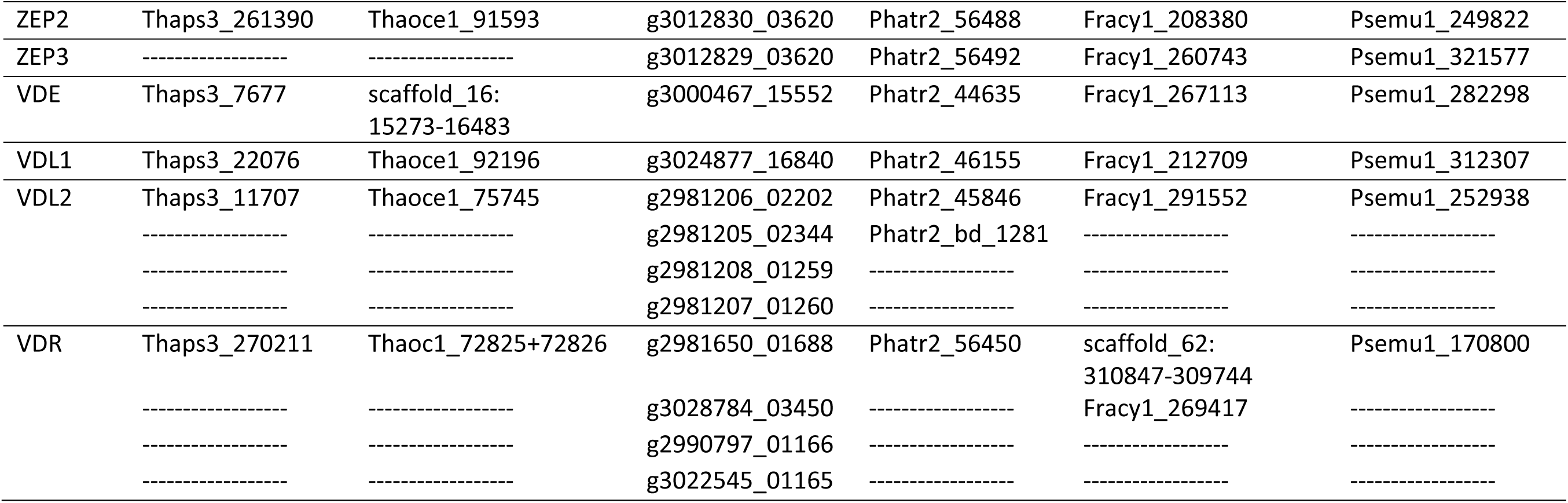
Carotenoid biosynthesis genes in currently available diatom genomes.

### Genetic Manipulation and Resultant Photosynthetic Pigment Phenotypes

The following transgenic *T. pseudonana* lines were created to investigate the roles of several enzymes hypothesized to participate in carotenoid biosynthesis: overexpression (OE) of VDL2, knockdown (KD) of VDL2, KD of VDL1, and a simultaneous KD of LUT1-like 1 (LTL1) and LTL2, which share 48% sequence identity **(Appendix D)**. Four independent clones per line were then allowed to adapt to cultivation conditions, and their photosynthetic pigment content was assessed using high-performance liquid chromatography (HPLC).

For VDL2 OE, total cellular photosynthetic pigment content (Tot) varied between the four clones and two WT cultures at both low (30 µmol photons m^-2^ sec^-1^) and high light (300 µmol photons m^-2^ sec^-1^), without an observable trend (Fig. 5A). Nevertheless, some pigment ratios exhibited a consistent difference between WT and VDL2 OE. Fx/Tot was increased (low light p-value = 0.02, high light p-value = 0.0008) and (Ddx+Dtx)/Tot was reduced (low light p-value = 0.0006, high light p-value = 0.0008) in VDL2 OE, but (Ddx+Dtx+Fx)/Tot was not significantly different from WT (low light p-value = 0.9, high light p-value = 0.8) (Fig. 5B). No differences were observed in β-car/Tot, Chl a/Tot, and Chl c/Tot (Fig. 5C, D).

**Fig. 5.**
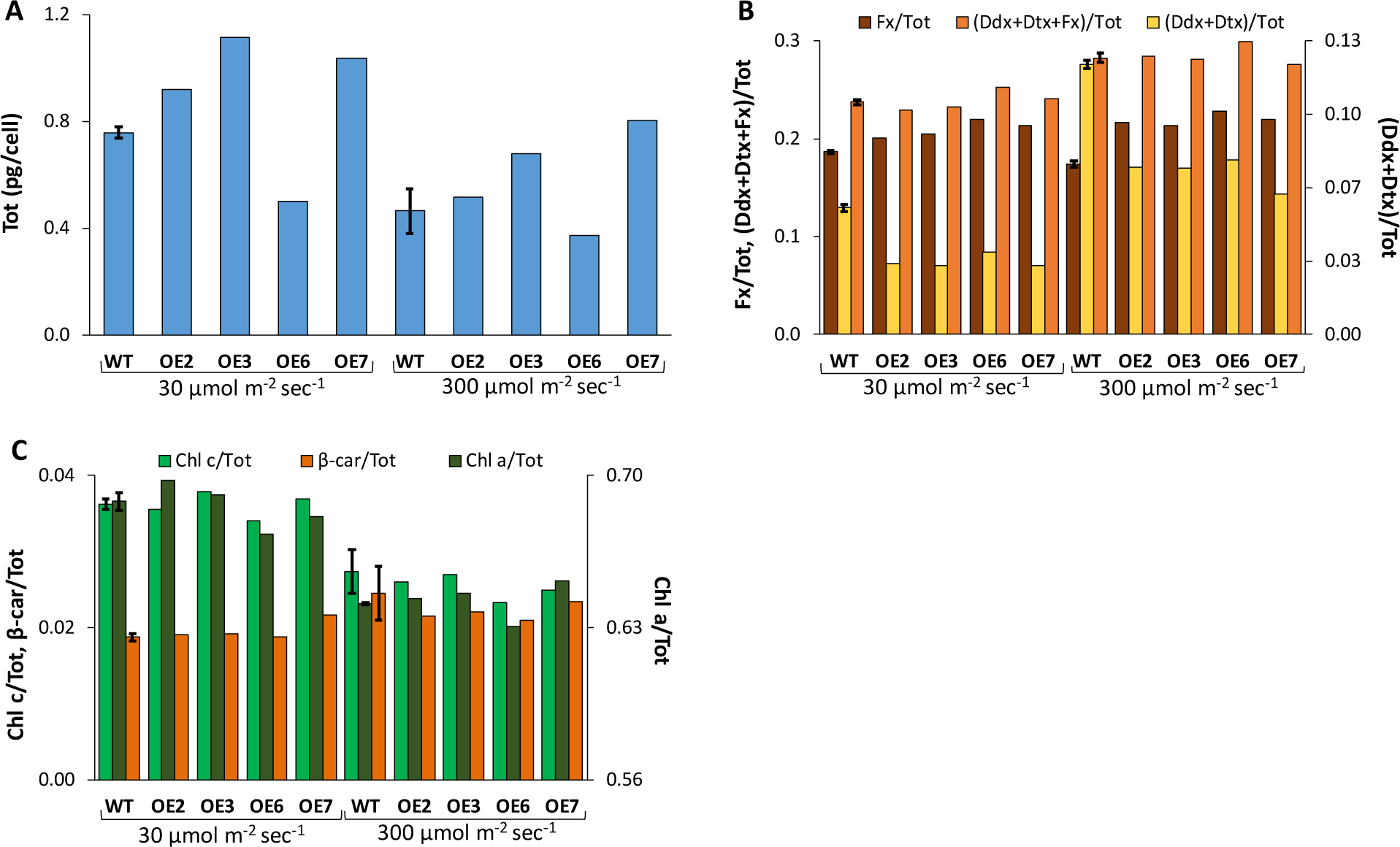
HPLC-based pigment analysis of VDL2 overexpression (OE) lines cultured at 30 µmol m^-2^ sec^-1^ and 300 µmol m^-2^ sec^-1^. Wild-type (WT) data is an average of two independent cultures. **A**. Total cellular photosynthetic pigments (Tot); **B.** Fx/Tot, (Ddx+Dtx)/Tot, (Ddx+Dtx+Fx)/Tot; **C.** Chl a/Tot, Chl c/Tot, β-car/Tot.

For LTL KD lines, initial PCR-based screening for the presence of the full KD construct (promoter to terminator) revealed a selection against it, as most of the screened clones had integrated the portion allowing them to grow on selection but lost all or parts of the promoter and/or the antisense portion. Approximately 3 weeks after plating, substantial reduction in pigmentation became apparent in some of the clones (Fig. 6A). Four LTL KD clones were selected based on lighter pigmentation, and genomic integration of the entire KD construct was subsequently confirmed by PCR. There was a reduction in Tot in LTL KDs compared to WT, with significant differences only in high light (Fig. 6B). In low light, LTL KD lines had 75-93% Tot of average WT (p-value = 0.2), and 60-76% of average WT Tot in high light (p-value = 0.009). No statistically significant differences in the ratios of individual photosynthetic pigments to Tot were found, although a trend of reduction in β-car is observable (low light p-value = 0.09, high light p-value = 0.08) (Fig. 6B, C).

**Fig. 6.**
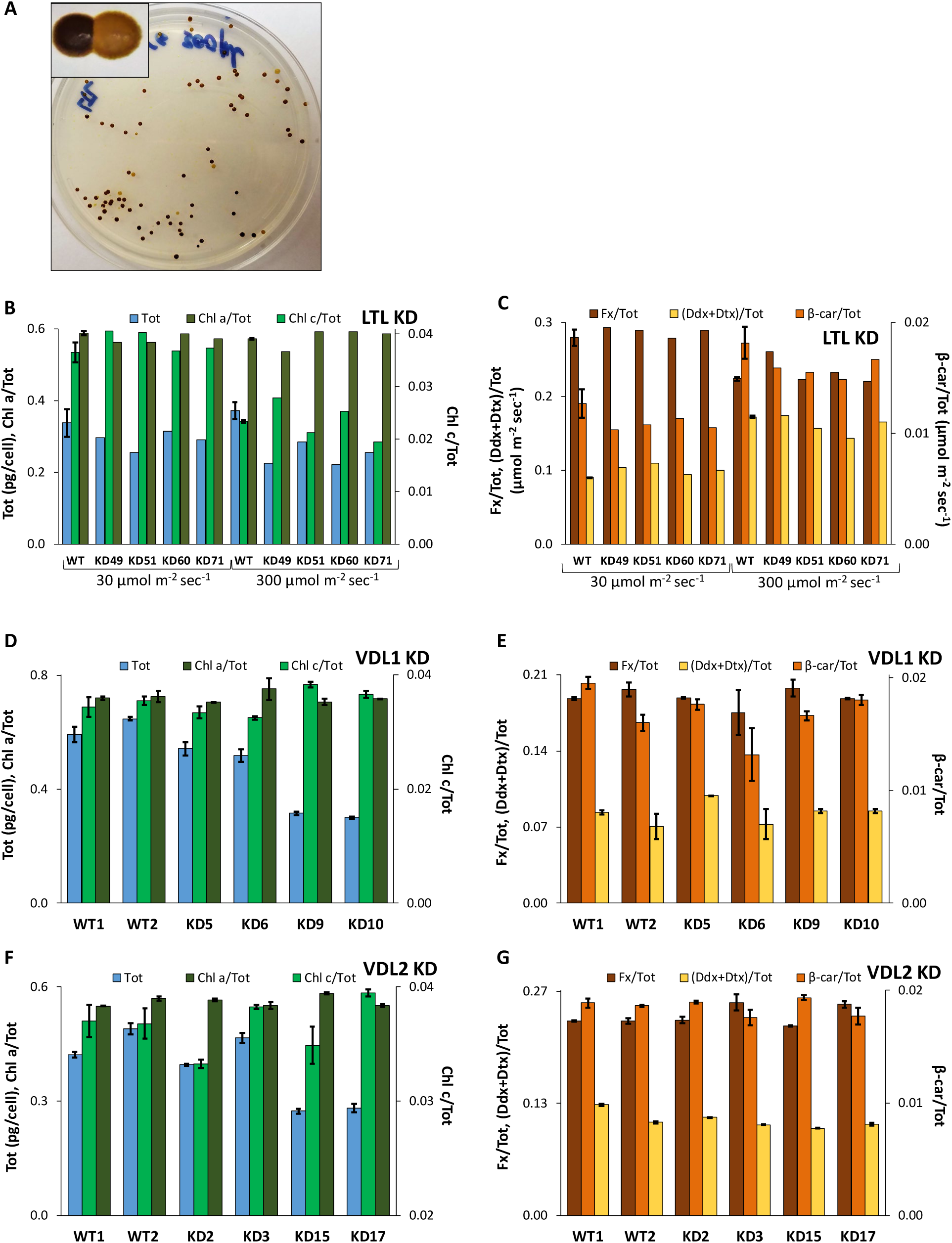
Knockdown (KD) line data. **A.** Lighter-pigmented LTL KD colonies among those with wild-type (WT)-equivalent pigmentation; HPLC-based pigment analysis of LTL KD lines cultured at 30 µmol m^-2^ sec^-1^ and 300 µmol m^-2^ sec^-1^. WT data is an average of two independent cultures. **B.** Tot, Chl a/Tot, Chl c/Tot; **C.** Fx/Tot, (Ddx+Dtx)/Tot, β-car/Tot. HPLC-based pigment analysis of VDL1 and VDL2 knockdown (KD) lines, cultured at 300 µmol m^-2^ sec^-1^, respectively. Each data point is an average of two samples taken from the same culture. **D, F.** Tot, Chl a/Tot, Chl c/Tot; **E, G.** Fx/Tot, (Ddx+Dtx)/Tot, β-car/Tot.

Four clones each of VDL1 KD and VDL2 KD were selected based on the ability to grow on selection and PCR-confirmed genomic integration of full-length KD constructs. Photosynthetic pigment content was assessed after acclimation to high light cultivation conditions and Tot was found to be reduced in both lines, with the exception of the clone VLD2 KD3 (Figs. 6D, F). VDL1 KD lines contained 48-91% of the average WT Tot (p-value = 0.01), and VDL2 KD lines were at 59-87% of average WT Tot (p-value = 0.004), excluding KD3. KD3 did not significantly differ from WT in Tot. No substantial differences in the ratios of individual pigments to Tot were observed between WT and the VDL1 or VDL2 KD lines (Fig. 6D-G).

## DISCUSSION

### Identification of Carotenoid Biosynthesis Gene/Enzyme Candidates

Building on earlier analyses [Coesel et al. 2008, Dambeck et al. 2012], we sought to identify novel gene candidates based on sequence identity of their protein products to those of previously identified carotenoid biosynthesis genes. The genomes of *T. pseudonana* [Armbrust et al. 2004] and *P. tricornutum* [Bowler et al. 2008] were chosen for reciprocal BLAST searches as the first two sequenced and best annotated of the currently available diatom genomes. A notable finding was that of an additional, previously unreported VDL protein in *T. pseudonana* (Thaps3_11707), designated VDL2 in this study. It should be noted that the *T. pseudonana* VDR (Thaps3_270211) is mis-annotated as VDL2 on the DOE JGI website.

The list of candidates for the *T. pseudonana* carotenoid biosynthesis pathway obtained by BLAST was narrowed to those whose gene expression patterns were similar to those of genes known to participate in carotenoid biosynthesis, using available transcriptomic (microarray) data [Smith et al. 2016] (Fig. 3). Smith et al. [2016] report substantial co-regulation of genes with related functions, as evidenced by numerous gene clusters with highly similar expression patterns. Nevertheless, it is possible that some gene candidates whose expression patterns differed from those of known carotenoid biosynthesis genes and were therefore not chosen for further study (Fig. 3) may be relevant to carotenoid biosynthesis but regulated differently.

The microarray data was obtained from a 24-hour silicon starvation time course during which there was a major chloroplast division event at 4 h, and light-induced stress was sustained throughout. The latter is at least partially attributable to the culture being resuspended in silica-free media at half the density that it had grown to in replete media prior to being harvested, resulting in reduced shading [Smith et al. 2016]. Both chloroplast division and exposure to higher light are expected to result in carotenoid biosynthesis [Gaidarenko et al. 2019, Ragni and d’Alcala 2007], and Smith et al. [2016] did measure an overall steady increase in photosynthetic pigment content throughout the time course, as well as a preferential accumulation of the photoprotective de-epoxide form (Dtx) of the interconvertible Ddx/Dtx cycle pigments. Thus, carotenoid biosynthesis activities associated with both chloroplast division and light-induced stress occurred during the experiment.

Smith et al. [2016] also collected RNA-seq data alongside the microarray data, with a notable difference of the culture used for the former being resuspended in silica-free media at the same density that it had attained in replete media prior to harvesting. Thus, unlike the culture used for the microarray, the culture used for RNA-seq would not have been exposed to higher light due to reduced shading at the beginning of the silicon starvation experiment. Even though expression patterns for many genes replicated well between the microarray and RNA-seq silicon starvation datasets, those of genes known to be involved in the light-induced stress response [Smith et al. 2016], as well as those of genes known to participate in carotenoid biosynthesis (Figs. 3**, S1)** did not, suggesting that the responses observed in the microarray dataset could not be attributed solely to silicon starvation but were affected by increased light exposure of the microarray culture as well. No physiological data was collected from the silicon starvation RNA-seq culture, and thus the status of light stress, chloroplast division, and pigment abundance cannot be directly referenced for that dataset [Smith et al. 2016].

Candidates selected based on their expression in the microarray dataset [Smith et al. 2016] were further narrowed to those predicted to be targeted to the chloroplast by using full-length gene models constructed using available RNA-seq data [Abbriano 2017, Smith et al. 2016] **(Appendix C,** Table 2). Due to diatoms having evolved via secondary endosymbiosis, their chloroplasts are surrounded by the endoplasmic reticulum (ER) [Smith et al. 2012]. Thus, diatom proteins targeted to the chloroplast must also be targeted to the ER. PSY2 and LTL2 had clear ER targeting, but their predictive scores for chloroplast targeting were just below the cutoff value **(Appendix C,** Table 2). It is likely that they are indeed targeted to the chloroplast, as chloroplast targeting predictions are not always completely accurate [Emanuelsson et al. 1999].

The majority of candidates identified by BLAST, including the numerous LTL, CRTISO, and ZEP candidates, did not meet the criteria of both having microarray gene expression patterns similar to genes known to participate in carotenoid biosynthesis and predicted chloroplast targeting. Barring potentially having excluded novel carotenoid biosynthesis genes that had different gene expression patterns or incorrectly predicted protein product targeting, this indicates that most of these candidates were found due to containing domains that are utilized by, but not limited to, carotenoid biosynthesis enzymes. Indeed, each of the CRTISO, LTL, and ZEP candidate groups had shared functional annotation predictions.

### Stepwise Analysis of the T. pseudonana Carotenoid Biosynthesis Pathway

Major known and hypothesized diatom carotenoid biosynthesis steps are depicted in Fig. 1. However, there may be additional/alternative biosynthesis steps and pathway intermediates.

The condensation of two geranylgeranyl pyrophosphate molecules to synthesize phytoene by PSY is the first committed step of carotenoid biosynthesis and is considered rate-limiting [Bertrand 2010, Eilers et al. 2016]. It is followed by several desaturation steps catalyzed by PDS [Melendez-Martinez et al. 2015]. There are two previously reported copies of each in *T. pseudonana* [Coesel et al. 2008], and a possible additional *T. pseudonana* PDS identified in this study. The novel PDS candidate Thaps3_bd_1474 appears to have arisen as a result of a duplication event of the PDS1 locus, encompassing it, as well as two downstream genes. Because of the way “bottom drawer” sequences were processed, the 5’ end as well as the genomic sequence upstream of Thaps3_bd_1474 are not available, and the size of the duplicated region is unknown.

In vascular plants that possess more than one copy of PSY, the different genes are differentially regulated in response to developmental and environmental signals. Differential regulation based on different cellular needs may be hypothesized for microalgae that possess more than one PSY gene, but this has yet to be experimentally investigated [Melendez-Martinez et al. 2015, Tran et al. 2009]. Because PDS is responsible for the next immediate steps in carotenoid biosynthesis and has also been previously suggested to be rate-limiting [Chamovitz et al. 1993], different PDS copies might be differentially co-regulated with different copies of PSY. In the present study, both PSY and PDS genes were found to exhibit two distinct gene expression patterns in the microarray dataset [Smith et al. 2016]. PSY1, PDS1, and Thaps3_bd_1474 had a spike in expression at 4 h, during the major chloroplast division event (Fig. 3A, C). By contrast, the transcript levels of PSY2 and PDS2 increased by 4 h and remained elevated throughout the time course (Fig. 3B). In the RNA-seq dataset [Smith et al. 2016], although the magnitude of transcript level changes for carotenoid biosynthesis genes was modest, PSY1 expression varied more similarly to PDS1, while PSY2 expression varied more similarly to PDS2 (Fig. S1A). In an additional set of transcriptomic microarray data from a study by Ashworth et al. [2013], PSY1/PDS1 had similar and consistent patterns of diel changes in their expression levels during exponential and stationary growth phases in ambient and elevated CO^2^, whereas PSY2/PDS2 varied less predictably, but similarly to each other (Fig. S1C, D). The cellular accumulation of diatom photosynthetic pigments is influenced by cell cycle progression and therefore chloroplast division, with the exception of the Ddx cycle pigments, which appear to primarily respond to light intensity [Gaidarenko et al. 2019, Ragni and d’Alcala 2007]. The differential accumulation of the Ddx cycle pigments and Fx, the other end product of carotenoid biosynthesis, as well as β-car, their common precursor, suggests a differential regulatory mechanism. We hypothesize that in *T. pseudonana*, one set of PSY/PDS genes may serve to initiate carotenoid biosynthesis during chloroplast division and the other set may serve as an independent way to initiate carotenoid biosynthesis in response to increased irradiance, which leads to an increase in Ddx cycle pigments [Lavaud et al. 2004].

Following PDS, more desaturation is carried out by ʐ-carotene desaturase (ZDS), resulting in pro-lycopene, a lycopene stereoisomer. There is only one known copy of ZDS in *T. pseudonana*, and we present no new hypotheses or observations for this step. Pro-lycopene is isomerized to lycopene by CRTISO, and the identity of the enzyme(s) responsible for this step is not yet confirmed. Many potential CRTISO copies have been previously found in diatom genomes, including *T. pseudonana* [Bertrand 2010]. Our analysis has narrowed the list to two candidates, Thaps3_21900 and Thaps3_10233 (Fig. 2A, Table 2). It is possible, however, that other candidates identified by BLAST (Table 1**, Appendix A)** might participate in this step but were excluded based on having a gene expression pattern that differed from genes known to participate in carotenoid biosynthesis (Fig. 3), or due to a false negative prediction for chloroplast targeting (Table 2).

Lycopene is converted into β-car by lycopene cyclase B (LCYB), for which no new candidates or hypotheses were generated by this study. The subsequent conversion of β-car via hydroxylation to Zx is intriguing. That reaction is typically catalyzed by one of the two known types of BCH [Martin et al. 2008], neither of which are found in any of the currently available diatom genomes, except a partial 238 amino acid BCH-like sequence in *T. pseudonana*. The latter, however, is not predicted to be targeted to the chloroplast, and has a 342 amino acid C-terminal addition which is entirely unknown except for limited sequence identity to unidentified peptides from several diverse organisms (Table 3). Its gene expression pattern in the microarray dataset [Smith et al. 2016] is markedly different from any known carotenoid biosynthesis genes (Fig. 3E). Like the BCH-like sequence, the C-terminal peptide is not found in any other currently available diatom genomes (*C. cryptica*, *T. oceanica*, *F. cylindrus*, *P. multiseries*, *P. tricornutum*), which makes the *T. pseudonana* protein an evolutionary curiosity. It is unlikely to participate in carotenoid biosynthesis and has possibly evolved to perform a different function by losing chloroplast targeting and gaining the C-terminal sequence.

Diatom genomes contain genes encoding LUT1-like (LTL) enzymes, which are similar to the *Arabidopsis thaliana* enzyme (LUT1) that converts α-car to lutein (respective isomers of β-car and Zx) and has been demonstrated to have weak β-car hydroxylation activity. Since diatoms do not make α-car, it has been hypothesized that diatom LTLs function in place of BCH, but this has not been previously experimentally assessed [Bertrand 2010]. To do so, we simultaneously targeted both *T. pseudonana* LTL copies for silencing with antisense RNA, which resulted in a reduction of all photosynthetic pigments (Fig. 6A-C). Although this does not conclusively place the LTLs in a specific carotenoid biosynthesis step, it does implicate them as being involved. Both LTLs are cytochrome P450 monooxygenases (Table 4), and there are no oxygen atoms in β-car or the pigments that precede it (Fig. 1). Thus, it is unlikely that the LTLs catalyze anything upstream of β-car hydroxylation.

Zx, whether synthesized from β-car by the LTLs or another enzyme or set of enzymes yet to be discovered, is part of the Vx xanthophyll cycle. The de-epoxidation reactions of both the Vx and the Ddx xanthophyll cycles are catalyzed by ZEPs [Bertrand 2010, Wilhelm et al. 2006]. Chlorophytes lack the Ddx cycle and typically possess one ZEP copy [Frommolt et al. 2008], while multiple ZEPs are found in all diatom genomes available to date (Table 5). It may be hypothesized that the multiple diatom ZEPs evolved to have different functions. In support of that idea, we found that the two *T. pseudonana* ZEP copies had distinct gene expression patterns in the microarray data [Smith et al. 2006] (Fig. 3A, B). Additionally, distinct transcriptomic responses for the three *P. tricornutum* ZEP genes during a time course of exposure to blue, red, or white light have been documented [Coesel et al. 2008].

De-epoxidation of the Vx cycle pigments in chlorophytes is catalyzed by VDE. Only one VDE copy is found in the currently available diatom genomes (Table 5), and it has been shown to participate in both diatom xanthophyll cycles [Jakob et al. 2000, Lavaud et al. 2012]. In addition to the VDE, diatom genomes encode VDE-like (VDL) and VDE-related (VDR) proteins (Fig. 2C), the function of which has not been determined. They have been stipulated to have xanthophyll cycle activity in addition to the VDE, perhaps differing in localization [Bertrand 2010, Lavaud et al. 2012]. Because VDR appears to be generally present in chlorophytes [Coesel et al. 2008], it is unlikely to catalyze reactions that are unique to diatoms. In the present study, we identified a previously unreported VDL2 in *T. pseudonana* (Thaps3_11707) based on sequence identity to the VDL2 in *P. tricornutum*. Overexpressing it resulted in increased cellular Fx abundance, together with a stoichiometric decrease in the cellular abundance of Ddx+Dtx. The ratio of Ddx+Dtx+Fx to total cellular pigments remained unchanged (Fig. 5). Since the abundance of Chl a did not increase along with Fx, we suggest that the excess Fx in the VDL2 OE lines was either localized to the lipid shield around the photoantennae, along with non-protein bound Ddx cycle pigments, or that it may have replaced some of the photoantenna-bound Ddx [Lepetit et al. 2010]. Eilers et al. [2015] also found that increased Fx abundance in their *P. tricornutum* PSY OE lines was not accompanied by a concomitant increase in Chl a content. While our results do not clarify whether Ddx+Dtx and Fx share a common precursor or Fx is derived from Ddx+Dtx, they do implicate VDL2 in Fx biosynthesis. Our findings do not support the hypothesis that VDLs participate in xanthophyll cycling. Based on our data, the function of VDL1 remains unclear. Multiple chemical reactions must take place during Fx biosynthesis from either Ddx or the other hypothetical precursor, neoxanthin. However, based on our findings, if other enzymes besides VDL2 are involved in this step, they are not rate-limiting. VDL2 analysis based on sequence as well as predicted structure revealed only a VDE-like lipocalin domain, a fold that is shared by VDEs, VDLs, and ZEPs, and allows them to bind to their pigment substrates [Coesel et al. 2008]. The chemistry necessary to synthesize Fx includes additions of one acetyl and one keto group and oxidoreductase activity. VDL2 is predicted to be capable of oxidoreductase activity (Table 4), which could catalyze everything except for the acetyl group addition. It is also designated as part of the N-acetylglucosaminyltransferase containing a carbohydrate-binding WSC domain KOG group on the DOE JGI website (Table 4), which could suggest an ability to transfer acetyl groups. Almost half of the VDL2 protein on the N-terminal side shows no similarity to other known proteins based on either sequence or predicted structure and may perform previously undescribed chemistry.

Because Thaps3_10233, an oxidoreductase (Table 4) initially identified by BLAST during the CRTISO candidate search, exhibited transcript responses that were similar to VDL2 and photoantenna protein genes and distinct from other carotenoid biosynthesis genes, it may be hypothesized to be involved in Fx biosynthesis as well. The co-expression of the genes encoding proteins involved in Fx biosynthesis with photoantenna protein genes may represent another layer of carotenoid biosynthesis regulation, in addition to the hypothesized separate means of inducing the pathway in response to irradiance increase and chloroplast division. Synthesizing Fx only when photoantenna proteins are also being made may be a way to ensure that pigment precursors are funneled into Fx biosynthesis only when it is needed to populate newly synthesized photoantenna proteins, and otherwise are used to make Ddx cycle pigments. While some Ddx cycle pigments are bound to photoantenna proteins, the majority of those synthesized in response to an increase in illumination are dissolved in the lipid shield around photoantennae [Lepetit et al. 2010]. Therefore, their accumulation would not need to be coordinated with photoantenna protein production.

### Total Photosynthetic Pigment Reduction in Knockdown Lines

An overall reduction in cellular photosynthetic pigment content was observed in LTL, VDL1, and VDL2 KD lines, without significant differences in the ratios of individual pigments (Fig. 6). Thus, β-car, presumably upstream of the biosynthetic steps targeted by the KDs, as well as Chl a and Chl c, products of a separate biosynthetic pathway, were reduced proportionately with Fx and Ddx cycle pigments in the KD lines. Carotenoids are known to play a crucial role in the assembly and stabilization of light-harvesting complexes in photosynthetic bacteria, algae, and green plants [Moskalenko and Karapetyan 1996, Santabarbara et al. 2013]. Unlike chlorophytes that adjust the size of the photoantenna complexes associated with photosynthetic reaction centers in response to changes in light intensity, diatoms mainly co-regulate the number of reaction centers and photoantenna units. Thus, the ratio of Chl a to Fx and β-car does not vary substantially, while Chl c abundance appears to be more variable [Brunet et al. 2014, Lepetit et al. 2012]. In the KD lines, Chl a, Chl c, and β-car may have failed to accumulate to WT levels due to being prevented from stable binding to light-harvesting complex proteins because of light-harvesting complex destabilization caused by a Fx deficit, which may be especially important since the photosynthetic pigment ratios appear to be generally maintained in diatoms. Lohr and Wilhelm [1999] monitored photosynthetic pigment accumulation in *P. tricornutum* upon reducing illumination intensity, with and without the addition of the *de novo* carotenoid biosynthesis inhibitor norflurazon, which acts upon PDS. In the norflurazon-treated sample, the abundance of Fx increased as accumulated precursor pigments were depleted. In the untreated samples, following the initial conversion of precursors to Fx, Fx abundance continued to increase, presumably via *de novo* biosynthesis. Despite the differences in final Fx content between the treated and untreated samples, final Chl a/Fx and β-car/Fx ratios did not differ substantially, demonstrating that Chl a and β-car accumulation was proportional to that of Fx. Chl c abundance was not reported [Lohr and Wilhelm 1999]. This observation provides further support for the notion that reducing the abundance of Fx will also reduce that of the other photosynthetic pigments, as was observed in our KD lines (Fig. 6).

### Broader Implications of the Findings

The *T. pseudonana* genes identified and discussed in this study are also found in all other currently available diatom genomes (Table 5). Thus, our findings are relevant to diatom carotenoid biosynthesis in general, and not limited to our model species. There may exist, however, interspecies differences in certain aspects of the pathway. For example, *P. tricornutum* has only one copy of PSY, unlike *T. pseudonana* and other diatoms with sequenced genomes available that have at least two (Table 5). Therefore, the step that PSY catalyzes could not be used by *P. tricornutum* to differentially initiate carotenoid biosynthesis in response to chloroplast division and illumination increase. However, *P. tricornutum* does have two copies of PDS, which we hypothesize may be differentially used by *T. pseudonana* along with the two PSY copies. Consequently, *P. tricornutum* might use PDS, but not PSY, to differentiate between the two carotenogenic needs of the cell. Another observation is that some carotenoid biosynthesis genes in *P. tricornutum* are adjacent and appear to be divergently transcribed [Coesel et al. 2008], whereas no such relationships exist between the carotenoid biosynthesis genes in *T. pseudonana*. Interestingly, two such pairs in *P. tricornutum* are VDE with ZEP3 and VDL2 with ZEP1. A more in-depth discussion on the syntenic arrangement (or lack thereof) of ZEPs with VDE and VDL genes in diatoms and beyond can be found in the manuscript by Frommolt et al. [2008]. Diatoms are incredibly diverse and adapted to a wide variety of environmental conditions over the course of evolution. Strategies employed by different species may vary [Hildebrand et al. 2012], including those related to photosynthesis and photoprotection [Lavaud and Lepetit 2013]. However, increasing what is known about a given process in one diatom species ultimately facilitates future efforts for studying it in others, and improves our overall understanding of these environmentally important and commercially promising organisms.

Our results have important implications for studying diatom carotenoid biosynthesis. If knocking down (or knocking out) most steps in the pathway will result in the same phenotype of overall photosynthetic pigment reduction, as was the case with all knockdowns performed in our study, it will not reveal enzyme function. However, if it is possible to perform chemical rescue experiments by feeding various pigments in the pathway to knockdown lines, that may be a viable approach to further pathway elucidation. There are some exceptions for which the knockdown approach alone may be helpful. VDE, for example, does not participate in forward *de novo* carotenoid biosynthesis. It has been knocked down in *P. tricornutum* without reducing pigment content, resulting only in impaired Ddx de-epoxidation (and likely that of Vx as well, which was not assessed) [Lavaud et al. 2012]. Our hypothesis about there being two ways to independently induce carotenoid biosynthesis in response to either chloroplast division or an increase in illumination may be tested by knocking down PSY1 and PSY2 separately as well. If one of them indeed serves to initiate carotenoid biosynthesis during chloroplast division in order to populate newly divided chloroplasts with pigments, knocking it down should result in an overall photosynthetic pigment reduction. If the other copy initiates carotenoid biosynthesis in order to make more Ddx cycle pigments in response to increased illumination, knocking it down should not affect cellular pigmentation at low light intensities, but should result in impaired Ddx cycle pigment pool accumulation upon being transferred to higher light. Overexpression, as a different approach to assessing enzyme function, may be limited by substrate availability and not usable for all the pathway steps. Heterologous complementation, such as performed by Dambeck et al. [2012] for the earlier carotenoid biosynthesis steps in *P. tricornutum*, may help in the study of the function of enzymes involved in the later steps as well.

Our findings must be taken into consideration if altering diatom pigment content by genetically manipulating carotenoid biosynthesis is of interest. At least in *T. pseudonana*, it does not appear possible to use this approach to reduce the amount of the main accessory light-harvesting pigment Fx without affecting the abundance of other photosynthetic pigments, in order to hyperaccumulate β-car for commercial purposes or obtain a photoantenna reduction phenotype that has been achieved in chlorophytes [e.g., Kirst et al. 2012], for example. On the other hand, it is possible to increase cellular Fx content, which may be of interest due to its various health-promoting activities [Peng et al. 2011].

## METHODS

### Carotenoid Biosynthesis Gene Candidate Search Based on Sequence Identity

*T. pseudonana* and *P. tricornutum* genome sequence data was accessed on the DOE JGI website [Grigoriev et al. 2012, Nordberg et al. 2014] at https://genome.jgi.doe.gov/Thaps3/Thaps3.home.html and https://genome.jgi.doe.gov/Phatr2/Phatr2.home.html, respectively. Database searches were performed using the available tBLASTn function, comparing protein queries to translated nucleotide sequences. Sequence identity analyses were performed on protein sequences predicted by the DOE JGI website. Multiple sequence alignment, percent identity matrix generation, and phylogenetic tree construction were completed using Clustal Omega (https://www.ebi.ac.uk/ Tools/msa/clustalo/) [Sievers et al. 2011]. Trees were visualized using Interactive Tree of Life (iTOL) (https://itol.embl.de/) [Letunic and Bork 2016]. Gene Ontology (GO) terms, EuKaryotic Orthologous Groups Identity (KOG ID), and Protein Family (Pfam) information were provided by the DOE JGI website.

### Full-length Gene Model Construction

RNA-seq data [Abbriano 2017, Smith et al. 2016] was visualized using the Integrated Genomics Viewer (IGV) [Robinson et al. 2011, Thorvaldsdottir et al. 2013]. Exon boundary coordinates were used to obtain genomic sequences on the DOE JGI website.

### Predicted Protein Targeting Analysis

Open reading frame translations for the full-length gene models were obtained using the online ExPASy Translate tool (https://web.expasy.org/translate/). Because chloroplast-targeted diatom proteins must cross the ER membrane first, online programs SignalP 3.0 [Bendtsen et al. 2004] and SignalP 4.1 [Petersen et al. 2011] were used to predict ER targeting, and ChloroP 1.1 [Emanuelsson et al. 1999] was used to predict the presence of chloroplast transit domains, as previously discussed in Smith et al. [2012].

### Additional Sequence-Based Analyses

Alignments and sequence identity analyses were performed with Clustal Omega. BCH sequences were obtained from the National Center for Biotechnology Information (NCBI) website (https://www.ncbi.nlm.nih.gov/) using previously published GenBank accession numbers [Tian and DellaPenna 2004]. BLASTp and tBLASTn programs on the NCBI website were used to search for proteins and translated nucleotide sequences with sequence identity to the C-terminal portion of the BCH-like Thaps3_263437. Conserved protein domain and motif analyses were performed using the NCBI Conserved Domains Search (https://www.ncbi.nlm.nih.gov/Structure/cdd/wrpsb.cgi) [Marchler-Bauer et al. 2017], the ScanProsite tool (https://prosite.expasy.org/scanprosite/) [De Castro et al. 2006], and InterPro (https://www.ebi.ac.uk/interpro/) [Finn et al. 2017]. Functional annotation predictions for the *T. pseudonana* carotenoid biosynthesis enzymes were obtained from the DOE JGI website as well as by protein sequence analysis using InterPro and Pfam (https://pfam.xfam.org/search/sequence). Structure-based functional annotation predictions were performed with Phyre2 [Kelley et al. 2015] (http://www.sbg.bio.ic.ac.uk/phyre2/html/page.cgi?id=index).

Additional diatom genomes were accessed at: http://genomes3.mcdb.ucla.edu/cgibin/hgGateway?hgsid=23003&clade=plant&org=Ahi+simulation+02&db=0 for *Cyclotella cryptica* (2011 Assembly), https://genome.jgi.doe.gov/Fracy1/Fracy1.home.html for *Fragilariopsis cylindrus*, https://genome.jgi.doe.gov/Psemu1/Psemu1.home.html for *Pseudo-nitzschia multiseries*, and https://genome.jgi.doe.gov/Thaoce1/Thaoce1.home.html for *Thalassiosira oceanica*.

Carotenoid biosynthesis genes in those genomes were found by using BLAT for *C. cryptica* and tBLASTn for the others, with predicted *T. pseudonana* protein sequences and Phatr2_56492 (*P. tricornutum* ZEP3) as queries.

### Genetic Manipulation

Cloning was carried out using MultiSite Gateway technology (Thermo Fisher Scientific, Waltham, MA, USA) as previously described [Shrestha and Hildebrand 2015]. VDL2 OE was driven by the *T. pseudonana* nitrate reductase (NR) promoter, LTL KD by the *T. pseudonana* ribosomal protein L41 (rpL41) promoter, and VDL1 and VDL2 KDs by the *T. pseudonana* acetyl CoA carboxylase (ACCase) promoter, with the corresponding terminators used respectively. The rpL41 promoter is the strongest out of the three, and NR is the weakest [Shrestha et al. 2013]. The KD constructs contained a nourseothricin resistance gene encoding nourseothricin N-acetyltransferase (NAT1) upstream 500-600 bp of antisense per gene of interest on the same transcript. Plasmid and primer details are available in **Appendix E**. The constructs were transformed into WT *T. pseudonana* using tungsten microparticle bombardment with Bio-Rad PDS-1000/He, following procedures described by Davis et al. [2017], with the modification of incubating cells in dim light overnight following the bombardment. VDL2 OE was co-transformed with another plasmid carrying the NAT1 gene under the *T. pseudonana* ACCase promoter (received from N. Kroger, Germany). Genomic integration of the constructs was confirmed by PCR, as previously described [Shrestha and Hildebrand 2015]. For VDL2 OE, RNA extraction, reverse transcription, qRT-PCR primer design, and qRT-PCR with normalization to the TATA box binding protein Thaps3_264095 were performed as in Shrestha and Hildebrand [2015]. Four clones were chosen for further analysis (Fig. S2).

### Cultivation Conditions and Photosynthetic Pigment Analysis

*T. pseudonana* WT and transgenic cultures were cultivated at either 30 or 300 µmol photons m^-2^ sec^-1^ (natural white LED lighting, superbrightleds.com, NFLS-NW300X3-WHT-LC2), using a 12:12 h light:dark regime, at 18°C. 50 mL cultures in Erlenmeyer flasks were maintained in Artificial Sea Water (ASW) medium [Darley and Volcani 1969] with rapid stirring. Each experimental set included 2 WT and 4 transgenic cultures. After inoculation, the cultures were grown to 1-3×10^6^ cells/mL, then allowed to adapt to the cultivation conditions by daily dilutions that maintained exponential growth with culture density under 2.5×10^6^ cells/mL for a minimum of 2 weeks prior to sampling for pigments. Cultures were rotated between stir plates daily to minimize any position-specific differences. Sampling was performed within the first two hours of the photoperiod. Immediately prior to sampling, cultures were allowed to adapt to the light environment of the biosafety cabinet for 45-90 min. 10-20 mL of 1×10^6^-2.5×10^6^ cells/mL cultures were harvested, processed, and analyzed by HPLC as previously described [Kozlowski et al. 2011]. HPLC analysis employed an internal control of including the same sample in two different quantities to ensure precision in pigment quantification, which scaled accordingly. An online one-way ANOVA calculator accessed at https://www.socscistatistics.com/tests/anova/default2.aspx was used for statistical analysis of pigmentation differences between WT and transgenic lines.

## Supporting information

Appendix A - BLAST Results

Appendix B - Phylogeny

Appendix C - Gene Models and Targeting Predictions

Appendix D - Additional Sequence-Based Analyses

Appendix E - Plasmid Maps and Primers

## AUTHOR CONTRIBUTIONS

Conception and design (OG and MH), experimental work (OG and DWM), provision of HPLC equipment and supplies (MV), data analysis and interpretation (OG), drafting of the manuscript (OG), critical revision for important intellectual content and final approval (OG, MV, DWM). MH is deceased.

---

**Fig. S1.**
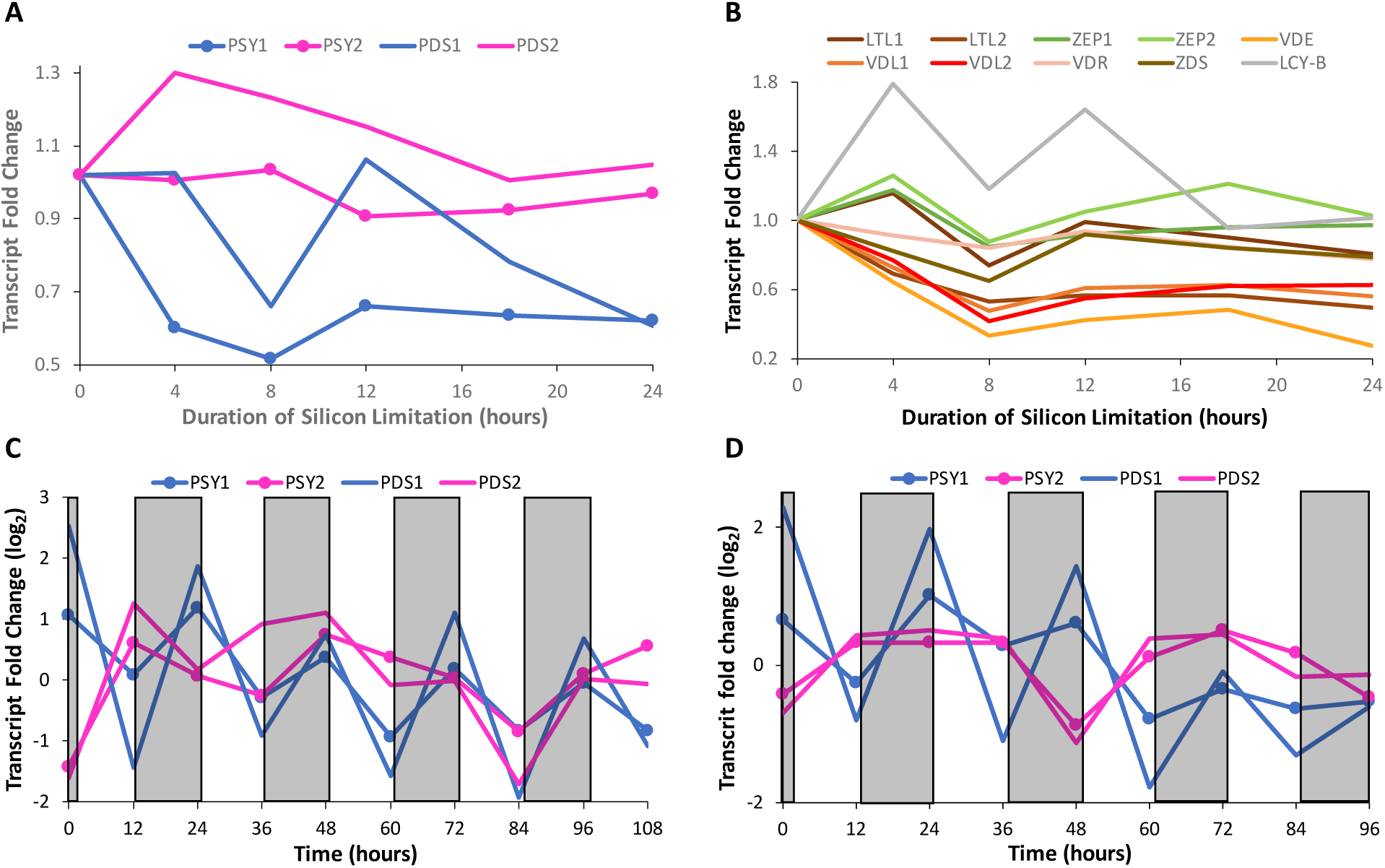
RNA-seq carotenoid biosynthesis gene expression patterns during silicon starvation [Smith et. al. 2016]. **A.** PSY1, PSY2, PDS1, PDS2; **B.** The other genes in the carotenoid biosynthesis pathway; PSY1, PSY2, PDS1, and PDS2 gene expression patterns in the diel transcriptomic microarray data [Ashworth et al. 2013]. Stationary phase begins after 48 hours. **C.** 400 ppm CO_2_. **D.** 800 ppm CO_2_.

**Fig. S2.**
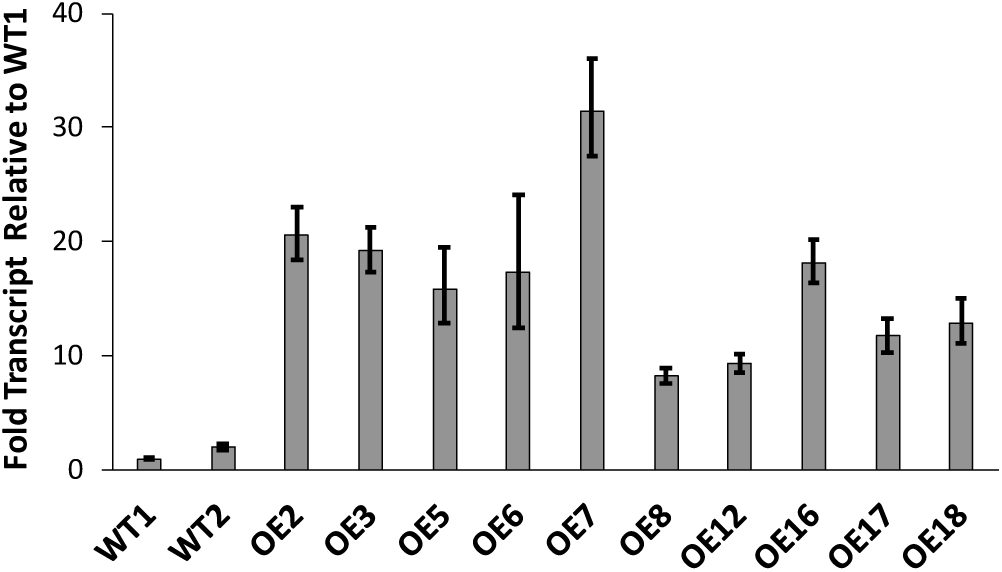
qRT-PCR screen for VDL2 overexpression (OE) compared to wild-type (WT) transcript levels. Clones 2, 3, 6, 7 were chosen for analysis. Each data point represents an average of two wells.

