## Appendix A - BLAST Results for "Overexpression of *Thalassiosira pseudonana* violaxanthin de-epoxidase-like 2 (VDL2) increases fucoxanthin while stoichiometrically reducing diadinoxanthin cycle pigment abundance"

1. Phytoene Synthase (PSY)

Previously reported single-copy PSY in *P. tricornutum* (Phatr2_56481) [Coesel et. al. 2008], was confirmed to have no other BLAST hits in the *P. tricornutum* genome. A BLAST search against the *T. pseudonana* genome yielded two hits, one previously published as PSY1 (Thaps3_268908) [Coesel et. al. 2008], and previously unpublished Thaps3_263269, which overlaps with previously published PSY2 (Thaps3_258309) [Coesel et. al. 2008].

2. Phytoene Desaturase (PDS)

BLAST queries of the previously published *P. tricornutum* PDS1 (Phatr2_45735) and PDS2 (Phatr2_55102) [Coesel et. al. 2008] against the *P. tricornutum* genome yielded each other, as well as a large chromosomal region (chr_1:926023-1979107) that includes Phatr2_53974, the ζ-carotene desaturase (ZDS) (3). In *T. pseudonana*, BLAST searches with the aforementioned *P. tricornutum* gene products yielded Thaps3_23291, which overlaps with the previously published PDS1 (Thaps3_6524) [Coesel et. al. 2008], Thaps3_1383, previously published as PDS2 [Coesel et. al. 2008], Thaps3_bd_1474 (not previously published), as well as the *T. pseudonana* ZDS (3), Thaps3_28432.

3. ζ-Carotene Desaturase (ZDS)

No candidates besides Phatr2_53974 and Thaps3_24832 (2) were found. The former had been reported by Coesel et. al. [2008], and the latter overlaps with the previously published gene model, Thaps3_37288 [Coesel et. al. 2008].

4. Carotene Cis-Trans Isomerase (Prolycopene Isomerase) (CRTISO)

Exhaustive reciprocal BLAST searches between the *P. tricornutum* and *T. pseudonana* genomes starting with the products of four genes identified by Dambeck et. al. [2012] as *P. tricornutum* CRTISO candidates (Phatr2_45243, Phatr2_9210, Phatr2_54842, Phatr2_51868) as queries yielded numerous hits in both organisms.

For *P. tricornutum*, the findings (in addition to the aforementioned genes) were Phatr2_54826, Phatr2_54800, Phatr2_42980. Three large chromosomal regions were repeatedly found as well: chr_1:1011072-1894441 (containing Phatr2_42890, Phatr2_9210, and Phatr2_53974, the ZDS), chr_6:586147-620230 (containing Phatr2_45243), and chr_15:52824-647530 (containing Phatr2_54826 and Phatr2_54800).

The findings for *T. pseudonana* were Thaps3_7094, Thaps3_21900, Thaps3_21847, Thaps3_5221, Thaps3_10233, Thaps3_11636, Thaps3_5859, Thaps3_25361, and Thaps3_10254. Two large chromosomal pieces were also found: chr_6:68923-1558412 (containing Thaps3_5859) and chr_15:676997-739441 (containing Thaps3_10233 and Thaps3_10254).

5. Lycopene β-cyclase (LCYB)

Phatr2_56484 [Coesel et. al. 2008] generated no additional BLAST hits in the *P. tricornutum* genome, and only Thaps3_270357 in the *T. pseudonana* genome. The latter generated no additional BLAST hits in either of the genomes, and overlapped with the previously reported gene model, Thaps3_261407 [Coesel et. al. 2008].

6. β-Carotene Hydroxylase (BCH)

As reported in Coesel et. al. [2008], no BCH was found in the *P. tricornutum* genome, and only Thaps3_263437, previously reported as a partial sequence [Coesel et. al. 2008], was found in the *T. pseudonana* genome.

7. LUT-Like (Lutein Deficient-Like) (LTL)

Exhaustive reciprocal BLAST searches between the *P. tricornutum* and *T. pseudonana* genomes, starting with Phatr2_50101 and Phatr2_26422 reported as LTL1 and LTL2, respectively, by Coesel et. al. [2008], yielded many hits in both genomes.

For *P. tricornutum*, the findings were Phatr2_34027, Phatr2_33568, Phatr2_6940, Phatr2_46438, Phatr2_31339, Phatr2_47234, Phatr2_37006, Phatr2_43466, Phatr2_43467, Phatr2_43562, Phatr2_50619, Phatr2_43469, Phatr2_32833, Phatr2_43537. Several regions without available gene models also appeared in the BLAST results: chr_2:488287-489217, chr_4:1314546-1315166, chr_8:134705-134894, chr_8:983359-983442, chr_15:149534-149668, and chr_15:494943-495059. Where open reading frames were readily apparent, the hypothetical protein products were included as BLAST search queries. Additionally, there were several larger regions: chr_2:487571-543110, chr_2: 540661-973489, chr_4:48892-1315175 (contains Phatr2_34027), chr_8:134708-983593, and chr_11:518522-869181 (includes Phatr2_37006).

For *T. pseudonana*, Thaps3_9541 was confirmed to be LTL1, and a more complete model for LTL2 (Thaps3_270336) compared to the previously reported one that was missing the N-terminus (Thaps3_36235), was found [Coesel et. al. 2008]. Additional results were Thaps3_33926, Thaps3_32491, Thaps3_1549, Thaps3_264647, Thaps3_25944, Thaps3_14875, Thaps3_4027, Thaps3_4026, Thaps3_bd_518, Thaps3_269400, Thaps3_264325, Thaps3_263399. Two large chromosomal regions were also found: chr_9:88966-685832 (includes Thaps3_270336) and chr_3:2376984-2382496 (includes Thaps3_14875, Thaps3_4026, Thaps3_4027, and Thaps3_25944, which appear immediately adjacent to each other, in the order listed).

8. Zeaxanthin Epoxidase (ZEP)

For *P. tricornutum*, previously published [Coesel et. al. 2008] ZEP1 (Phatr2_45845), ZEP2 (Phatr2_56488), and ZEP3 (Phatr2_56492) were confirmed. Additionally, Phatr2_43425, Phatr2_47925, Phatr2_45936, and chr_21:64775 – 64879 (no gene model) were found.

For *T. pseudonana*, Thaps3_270370 was identified as ZEP1, overlapping with the previously reported Thaps3 269147, and Thaps3_261390 was confirmed as ZEP2 [Coesel et. al. 2008]. Additional findings were Thaps3_1961, Thaps3_6395, Thaps3_20663, Thaps3_22671, and chr_9:932102-933624 (no gene model).

9. Violaxanthin De-Epoxidase (VDE), VDE-Like (VDL), VDE-Related (VDR)

In *P. tricornutum*, previously published VDE (Phatr2_44635), VDL1 (Phatr2_46155), and VDL2 (Phatr2_45846) were confirmed [Coesel et. al. 2008], and Phatr2_bd_1281 was found. Additional found sequences without available gene models were bd_29x34:989-1110, chr_2:9199-9410, and chr_1:974981-975118.

In *T. pseudonana*, previously reported VDE (Thaps3_7677) and VDL1 (Thaps3_22076) were confirmed [Coesel et. al 2008], and Thaps3_11707 as well as chr8: 84698 – 842033 were found.

Only previously reported Phatr2_56450 and Thaps3_270211 [Coesel et. al. 2008] were found in the VDR search.
