## Appendix B - Phylogeny for "Overexpression of *Thalassiosira pseudonana* violaxanthin de-epoxidase-like 2 (VDL2) increases fucoxanthin while stoichiometrically reducing diadinoxanthin cycle pigment abundance"

1)

### CRTISO Percent Identity Matrix

| | | | | | | | | | | | | | | | | |
|---|---|---|---|---|---|---|---|---|---|---|---|---|---|---|---|---|
| 1: Thaps3_25361 | 100.00 | 47.05 | 16.73 | 16.24 | 16.79 | 18.23 | 19.17 | 18.06 | 16.46 | 17.20 | 15.98 | 19.00 | 18.10 | 18.94 | 17.98 | 19.76 |
| 2: Phatr2_42980 | 47.05 | 100.00 | 16.61 | 16.85 | 16.60 | 16.04 | 16.96 | 18.22 | 16.96 | 17.09 | 19.25 | 18.32 | 19.62 | 19.64 | 17.65 | 19.75 |
| 3: Thaps3_5221 | 16.73 | 16.61 | 100.00 | 57.64 | 35.21 | 36.94 | 32.19 | 32.73 | 30.20 | 31.70 | 32.04 | 32.56 | 18.32 | 18.30 | 17.49 | 19.42 |
| 4: Phatr2_54826 | 16.24 | 16.85 | 57.64 | 100.00 | 36.55 | 35.94 | 34.01 | 32.73 | 31.65 | 33.06 | 32.34 | 34.94 | 17.92 | 20.03 | 18.49 | 19.77 |
| 5: Thaps3_21847 | 16.79 | 16.60 | 35.21 | 36.55 | 100.00 | 58.92 | 36.35 | 37.59 | 32.29 | 33.98 | 32.16 | 35.21 | 17.42 | 18.71 | 18.18 | 22.05 |
| 6: Phatr2_51868 | 18.23 | 16.04 | 36.94 | 35.94 | 58.92 | 100.00 | 35.51 | 34.78 | 31.49 | 33.59 | 31.83 | 34.95 | 20.00 | 19.75 | 18.37 | 20.99 |
| 7: Thaps3_7094 | 19.17 | 16.96 | 32.19 | 34.01 | 36.35 | 35.51 | 100.00 | 59.07 | 34.94 | 38.37 | 37.07 | 38.36 | 22.20 | 22.60 | 20.13 | 22.95 |
| 8: Phatr2_54842 | 18.06 | 18.22 | 32.73 | 32.73 | 37.59 | 34.78 | 59.07 | 100.00 | 33.74 | 37.45 | 35.55 | 38.13 | 21.05 | 20.56 | 20.48 | 21.98 |
| 9: Thaps3_10233 | 16.46 | 16.96 | 30.20 | 31.65 | 32.29 | 31.49 | 34.94 | 33.74 | 100.00 | 85.09 | 36.67 | 36.57 | 18.40 | 20.49 | 17.93 | 20.19 |
| 10: Phatr2_9210 | 17.20 | 17.09 | 31.70 | 33.06 | 33.98 | 33.59 | 38.37 | 37.45 | 85.09 | 100.00 | 37.65 | 38.39 | 19.05 | 21.46 | 20.10 | 22.53 |
| 11: Thaps3_21900 | 15.98 | 19.25 | 32.04 | 32.34 | 32.16 | 31.83 | 37.07 | 35.55 | 36.67 | 37.65 | 100.00 | 43.01 | 18.59 | 19.59 | 19.51 | 19.76 |
| 12: Phatr2_45243 | 19.00 | 18.32 | 32.56 | 34.94 | 35.21 | 34.95 | 38.36 | 38.13 | 36.57 | 38.39 | 43.01 | 100.00 | 20.35 | 22.35 | 20.78 | 21.30 |
| 13: Thaps3_11636 | 18.10 | 19.62 | 18.32 | 17.92 | 17.42 | 20.00 | 22.20 | 21.05 | 18.40 | 19.05 | 18.59 | 20.35 | 100.00 | 59.69 | 22.22 | 27.49 |
| 14: Phatr2_54800 | 18.94 | 19.64 | 18.30 | 20.03 | 18.71 | 19.75 | 22.60 | 20.56 | 20.49 | 21.46 | 19.59 | 22.35 | 59.69 | 100.00 | 21.68 | 24.68 |
| 15: Thaps3_5859 | 17.98 | 17.65 | 17.49 | 18.49 | 18.18 | 18.37 | 20.13 | 20.48 | 17.93 | 20.10 | 19.51 | 20.78 | 22.22 | 21.68 | 100.00 | 28.43 |
| 16: Thaps3_10254 | 19.76 | 19.75 | 19.42 | 19.77 | 22.05 | 20.99 | 22.95 | 21.98 | 20.19 | 22.53 | 19.76 | 21.30 | 27.49 | 24.68 | 28.43 | 100.00 |

### CRTISO Alignment

```
Thaps3_25361      MRMGRPNKKLRSTSKQTTNPNPPKYSSPTLVVGQVSSNVIHSIYGPALTKLAVESV----        56
Phatr2_42980      MRIGKPLRKSRAWKKV-GVNGTPKYVSPESVVGRINSDIISSVLAPKIAALASASL----        55
Thaps3_5221       ------------------------------------------------------------        0
Phatr2_54826      ------------------------------------------------------------        0
Thaps3_21847      ------------------------------------------------------------        0
Phatr2_51868      ------------------------------------------------------------        0
Thaps3_7094       ------------------------------------------------------------        0
Phatr2_54842      ------------------------------------------------------------        0
Thaps3_10233      ------------------------------------------------------------        0
Phatr2_9210       ------------------------------------------------------------        0
Thaps3_21900      ------------------------------------------------------------        0
Phatr2_45243      ------------------------------------------------------------        0
Thaps3_11636      ------------------------------------------------------------        0
Phatr2_54800      --------------------MTPVTQSPPVELSTD-PPVALSLALPPLSPTADGTLQHHH        39
Thaps3_5859       ------------------------------------------------------------        0
Thaps3_10254      ------------------------------------------------------------        0


Thaps3_25361      ----------------------EEYADAV-LRWEASLPEVLVK-----------------        76
Phatr2_42980      ----------------------ERYAGEL-LVYEDIMKKVNSN-----------------        75
Thaps3_5221       ------------------------------------------------------------        0
Phatr2_54826      ------------------------------------------------------------        0
Thaps3_21847      ------------------------------------------------------------        0
Phatr2_51868      ------------------------------------------------------------        0
Thaps3_7094       ------------------------------------------------------------        0
Phatr2_54842      ------------------------------------------------------------        0
Thaps3_10233      ------------------------------------------------------------        0
Phatr2_9210       ------------------------------------------------------------        0
Thaps3_21900      ------------------------------------------------------------        0
Phatr2_45243      ------------------------------------------------------------        0
Thaps3_11636      ------------------------------------------------------------        0
Phatr2_54800      LTTTDDESSSPWPVIRSVFRGQNNFSDPLNRGWNPWRPGISSRQDKCGVEYVKMHGQYFP        99
Thaps3_5859       ------------------------------------------------------------        0
Thaps3_10254      ------------------------------------------------------------        0


Thaps3_25361      ---------P------SQLDDAEDIDVDADGTFEKGEEVEVDLDGSILPSHDNDDDKTSS        121
Phatr2_42980      ---------E------S-----------IDGLLESDS--------SIIIQ----DG---I        94
Thaps3_5221       ------------------------------------------------------------        0
Phatr2_54826      ------------------------------------------------------------        0
Thaps3_21847      ------------------------------------------------------------        0
```

```
Phatr2_51868      ------------------------------------------------------------      0
Thaps3_7094       ------------------------------------------------------------      0
Phatr2_54842      ------------------------------------------------------------      0
Thaps3_10233      ------------------------------------------------------------      0
Phatr2_9210       ------------------------------------------------------------      0
Thaps3_21900      ------------------------------------------------------------      0
Phatr2_45243      ------------------------------------------------------------      0
Thaps3_11636      ------------------------------------------------------------      0
Phatr2_54800      TSGSGFSTGPVGQHDGHELEQRCDTERQGGGHVNKGNHVEANVATSVLCATE-----CCL      154
Thaps3_5859       ------------------------------------------------------------      0
Thaps3_10254      -----------------------------MGFIVRGTRARSVVS------SR-----LAV      20

Thaps3_25361      PTMPTSNRL--FTTQSSIDNLTALLTDTSQH-FSTTNAWKIHANAAKFERLLDEKYGRFR      178
Phatr2_42980      PQQ-PQRPV--FHSQFSVDEAASIFSETSEHFFAKAGKWKAHANAAKFERILDEKYGILR      151
Thaps3_5221       ------------------------------------------------------------      0
Phatr2_54826      -----------MLV-------------------ESKKSRDG----SRTSS----RS----      18
Thaps3_21847      ------------------------------------------------------------      0
Phatr2_51868      ------------------------------------------------------------      0
Thaps3_7094       ------------------------------------------------------------      0
Phatr2_54842      ------------------------------------------------------------      0
Thaps3_10233      ------------------------------------------------------------      0
Phatr2_9210       ------------------------------------------------------------      0
Thaps3_21900      ------------------------------------------------------------      0
Phatr2_45243      ------------------------------------------------------------      0
Thaps3_11636      ------------------------------------------------------------      0
Phatr2_54800      PHREPEPKTVCVVVLRSLQRDPTFQQDEVSAFLGAFRGWIGNEVDRLFEW----AA----      206
Thaps3_5859       ------------------------------------------------------------      0
Thaps3_10254      KGQSPNPL-----------------------------------ESQLTNW----RE----      37

Thaps3_25361      PFIESHPELEVFIKKVQRKYAMGQFSPLRKGEGPMSTTSSIMLLFMMHRNGVRKELVALV      238
Phatr2_42980      PFITNHPEIEHFIRGVQRKYAMGYFSPFRQGDPPIPRSTAVIILFMMQRGQMRWEIMLLT      211
Thaps3_5221       ------------------------------------------------------------      0
Phatr2_54826      --------------LDTKTHCVCSTS------------KQ--------------------      32
Thaps3_21847      ------------------------------------------------------------      0
Phatr2_51868      ------------------------------------------------------------      0
Thaps3_7094       ------------------------------------------------------------      0
Phatr2_54842      ------------------------------------------------------------      0
Thaps3_10233      ------------------------------------------------------------      0
Phatr2_9210       ------------------------------------------------------------      0
Thaps3_21900      ------------------------------------------------------------      0
Phatr2_45243      ------------------------------------------------------------      0
Thaps3_11636      ------------------------------------------------------------      0
Phatr2_54800      --------------TAAKALCMANWQ------------GKSRLLK--K---FGATVRRLD      235
Thaps3_5859       -----------------------------------------------------------M      1
Thaps3_10254      --------------PPRRPFSKVQTH------------ASNQL------------QSEVM      59

Thaps3_25361      ALFTLVGLEPWALVGLVCVGKYSVDQRRRKRIGGMP--------------KKVKV----V      280
Phatr2_42980      TLFFLIGLQPWALVAVVGVLQGLLMRRKAKPLGKMK--------------RFIPA----V      253
Thaps3_5221       -----------------------------------MKLLF-VASTLIGVLSFTPP--QVL      22
Phatr2_54826      ----------NSVRPANTLLRSR-------ARHTLIWLVFYVWEWNTTTTAFAPSPSRIA      75
Thaps3_21847      ----------------------MK-------VSTTAT-FALLQIGTAAVSAFTSP-----      25
Phatr2_51868      ------------------MANISKD-R--L-TGRFLA-FLLLVLANKETSSFCVQSGYRS      37
Thaps3_7094       ------------------MLPHT-------TVHGVVALATLLLNAFVLVDSFAPS-----      30
Phatr2_54842      ------------------MFAISSQLT--LTLVGHLILLHMM-ENSAICSAFVPASQRTT      39
Thaps3_10233      ------------------MIGRKY--S--LAASA----LAIIA-SLTSTTAFAPPSSSLL      33
Phatr2_9210       ------------------------------------------------------------      0
Thaps3_21900      -------------------MIR---------SISA----LALLAACCPSVFSFAPLSVF--      27
Phatr2_45243      -------------------MR---------FSER----SLIACAICSISSAFVPIIHTPQ      28
Thaps3_11636      -----MNT------ITDLLFQNPSTFIV-L--------LPLLFIASFIFYIT--------      32
Phatr2_54800      KACGPFGWTAWILFPCQTLMQRRDHYTR-YAIGRS-R-TLGDAESCFLVWLF--------      284
Thaps3_5859       ELFSSINYDPWTLVPAS--------FYY-PTIITA-C-IPLLFIATAYWLLIRRA-----      45
Thaps3_10254      DSLSKISS---SLLGDGKHT---KVTTV-ATVAGL-T-LGTLFIARRIYLS---------      101

Thaps3_25361      ESYYAHGVVGE----------EEEESEEVERSK----KYAILEKPVGT-------IFNPA      319
Phatr2_42980      ESYYTDAKTDT----------EK---------------HELLLHPVGE-------PL-PS      280
Thaps3_5221       T--D------------------------TRHR-----------P----PALCNSGDATN      40
Phatr2_54826      AFRA------------------------SRGR-----------KLTTSVSSLVSGDKRD      99
```

```
Thaps3_21847      ----SI-------------NSVIRSPSTHLRS-----------SPSATAST---------     48
Phatr2_51868      RHYFSA-------------NFLSVQPSDVARG-----------SSTAPAAAI---ADAPT     70
Thaps3_7094       -PRCSH-------------------------R------YH----I---SSAA---STTLH     48
Phatr2_54842      FSNCRR-------------------------S------KN----RVGRHGCF---LLASQ     61
Thaps3_10233      RSTRLHSTVEETTNGEAATNTNVEQIKDTSRDKVMTFSYDMSIEPKYEKPTY---PGTGN     90
Phatr2_9210       ------------------------------------------------------------     0
Thaps3_21900      ---------------------------------------------RANAP-------SSL     35
Phatr2_45243      -----H------------------QSPRTTRH---QFT------RIYAAV-------SSV     49
Thaps3_11636      -------------------RWPQARP-VQFRR-----------------AD--RFRPEKV     53
Phatr2_54800      -------------------HWPARRVKLHPRR-----------------AS--RFRPELV     306
Thaps3_5859       ---QLH-------------REKGKLPKYDAIP-----------------SSVLKHIASKQ     72
Thaps3_10254      --------------------WMKEFPSSDSLP-----------------STNPVKQ-GFS     123


Thaps3_25361      DLSLRDEEYDVILLGCGPEVLYTASLL-SRAGKKTLVLSPREDASGCLTLQNG-------     371
Phatr2_42980      KEEIDASLFDALILGSGPASLYIASLL-SRAGRKVLVLSSRNDASGCLSIKHAE------     333
Thaps3_5221       DGGDEVHEVDIAVVGAGIGGLCAGAILNTLYDKKVGVYESHYLAGGCAHSFSRSVK----     96
Phatr2_54826      CASPADDLVDVAIIGAGLGGLCAGAILNTLYGKKVGIYEAHYLAGGCAHAFDRRAA----     155
Thaps3_21847      ITDADEEEWDVVVVGSGVGGLSAAAMC-ARYGLKTICVEAHDAPGGVAHSFERRAS----     103
Phatr2_51868      GSIYREETVDVVVIGAGVGGLSCAALS-SKYGMDTLCLEAHDTAGGCAHSFERYSA----     125
Thaps3_7094       ATTTPHSEYDAIIVGSGIGGLSAAALL-SHYGYSVAVFEAHSTPGGAAHGYTVNA-----     102
Phatr2_54842      SATPGTTSPTPVVVGSGIGGLCAAAML-AKYGYTVAVLESHNVPGGAAHGFTARDP----     116
Thaps3_10233      GMSGDSGEYDIIVIGSGMGGLACSALS-AKYGSRVLCLESHIKVGGSAHTFSRMHN----     145
Phatr2_9210       ---------DVIVIGSGMGGLACGALS-AKYGDKVLVLESHIKCGGSAHTFSRMHN----     46
Thaps3_21900      ASTTYQDEVDCIVIGSGIGGLSCAALL-AATGRTVRVLEQHYEIGGCAHAFYMDMNGKTV     94
Phatr2_45243      PSNSIPDEADVVVIGSGLAGLSCAALL-AHCGKRVVVLESHDAPGGAAHGWE--------     100
Thaps3_11636      P-----SNIDTIVIGSGSGGSTVANLL-AQSGQRVLVLEQHSVTGGCTHSFR--------     99
Phatr2_54800      LENGKQRRFDTIVIGSGSGGCACANLL-AQSGQRVLILEQHTKTGGCTHSFR--------     357
Thaps3_5859       VLRDLSGKIDVAIVGSGIAALSNASAL-AHQGYKIAVFEQNEIVGGCTHTFE--------     123
Thaps3_10254      IKSVSSTNWDVIVIGSGAGGLTTAALL-SKEGKKVLVLEQHDIAGGNLHTFS--------     174
                              ::*.*      .    :  .      . .   .*

Thaps3_25361      ------------KTNVPFDIDGSNIAHLARQ---------QSLLAPA------LCTTTDT     404
Phatr2_42980      ------------YSNVPFDVEASNVAKISRQ---------QQILAPA------LCTETDT     366
Thaps3_5221       ------IGDDEQPTTFTFDSGPTIVLGCSK----EPYNPLQQVLRAVGVDDQIEWLPYDG     146
Phatr2_54826      ------D-----GVNFTFDSGPTILLGCSS----PPFNALQQVLDAVGQK----------     190
Thaps3_21847      ---------SSPNRPFVFDSGPSLLSGMSS----KGTNPLRQVLDAVGTADDIDWVTYDG     150
Phatr2_51868      ---------ASKTTPFRFDSGPSLVSGLSE----KGTNPLRQVLDAVGTAEEVQWKTYDG     172
Thaps3_7094       ----------KDVGPLTFDTGPSFFSGLNSNYPAKSSNPLRSILDIID--EKVECIPYTT     150
Phatr2_54842      ----------KIEGEFRFDTGPSFFSGINSDTPAKASNPLRTVLDAID--ERVECVPYTT     164
Thaps3_10233      ------------GGKYSFEVGPSIFEGLDR----PSLNPLRMIFDILE--ETMPVKTYKG     187
Phatr2_9210       ------------GEKYSFEVGPSIFEGLDR----PSLNPLRMIFDVLE--EEMPVKTYTG     88
Thaps3_21900      PSSALKDDPTKKGELFHFEAGPSLYSGLSEE---RTPNPLKHIYQMIE--EEPEWLTYDQ     149
Phatr2_45243      ------------RRGFHFESGPSLYSGFAME---RSPNPLKNIFQITG--EDCEWITYDR     143
Thaps3_11636      ------------EEGCEWDTGLHYVSKAMA----TPTKRAGAIMSFMS-RGKQSFTPFPT     142
Phatr2_54800      ------------DRGCEWDTGLHYTSAGMG----RSTCRPGAIMHFMT-QGLQKWTPL--     398
Thaps3_5859       ------------KQGFEFDVGVHYVGGFG--------TVVKHMYDELS-DGQLKWTKL--     160
Thaps3_10254      ------------EKGYEFDTGLHYVGGKVG----DKSSSVRKQLDYVM-DTDVEWEKM--     215
                                   ::

Thaps3_25361      QGGIRFA--------RIG-SEVDGYAHSILSVPGLGTDSISNECIPIVLT-----AEGEV     450
Phatr2_42980      QGGVRFA--------QIG-SNEDAHAFEILSIPGMGTDSYDEELPFILNA-----DGGTA     412
Thaps3_5221       WGMIEHPMQ----------PKEKRWKF--KV----G----PNHFEDGPLQVF---ASNLN     183
Phatr2_54826      -----NPGK----------DNELRWKV--IL----G----RDEFQRGPLTRF---GG-PK     221
Thaps3_21847      WMVHDTAFP----------MDDSRSSFRLTT----G---SDGTWEDAIEAKAG--VDSRR     191
Phatr2_51868      WLVHDTS---------------DDKVFKVTT----G---DSGAFEDALEKKAG--INAKR     208
Thaps3_7094       FGLMFPE---------------GVFVHSSNF----G----KEG--STVEA-----VSGSN     180
Phatr2_54842      FGLQFPE---------------GNFEHSCFF----G----AQG--GLLEQ-----LQGTT     194
Thaps3_10233      LGYWTPS---------------GYWRFPIGS---------REGFEQLLMEQCG--EDGEK     221
Phatr2_9210       LGYWTPT---------------GYWRFPIGS---------QSKFEDLLMEQA---EDGPK     121
Thaps3_21900      WGAFLPE---------------APEGYQMSI----G----AENFCKILET-----YGGEG     181
Phatr2_45243      WGTVMPD---------------GT-KFAAKI----G----PEEFQDVLES-----QGGPG     174
Thaps3_11636      STPYDEIVFPKDANVKDGAPNEFSHKF--YD----G---VNRTVSSVIGSIDPSDNELKH     193
Phatr2_54800      QDPYDEVIFPPDDFVKLGVPNESSYRF--VS----G---ADETIQSVLASIDPEHRELEK     449
Thaps3_5859       DRVYDVMYNGRTG---------ERYEI--TD----D---HDK-----------NRRVLTK     191
Thaps3_10254      DDIYDVAICDEEQ---------F--NF--CS----S---WKT-----------LKVELKK     244


Thaps3_25361      ALAEYCSTYLGDAFPGTDLDGNDNGNSTSLSYLKACGQINAGSGDFYL-------AKLFP     503
Phatr2_42980      GLIDDAAKYLNDGWPDAE---GGNGNSVTGAYAAACEAINSTANEFYI-------SKILS     462
Thaps3_5221       ALEEF------------------------NQLREITKPLVTGAATIPAMAMRPGQSALV-     218
```

```
Phatr2_54826    ALEEF------------------------EALREATKDLLAG-AKIPAMAMRPGPSALV-          255
Thaps3_21847    EFTKF------------------------KKKMMSSGGLSESSALLPPMALRGDFGALF-          226
Phatr2_51868    EFIEF------------------------KRKVLEEGGLAEASAYIPPFALRGGITALA-          243
Thaps3_7094     GVQEW------------------------ASLMKSMDPLAQAVDAMPTLALRADLGLLA-          215
Phatr2_54842    AQKEW------------------------QALMQSMGPLEKAVAALPTAALRGDIGLLL-          229
Thaps3_10233    AIGEW------------------------KALRERLRTLGGSTQAVALLNLRQDAGFLA-          256
Phatr2_9210     AVEEW------------------------NMLRKRLKTLGGSTTAVSLLNLRQDPGFLA-          156
Thaps3_21900    AVEDW------------------------EKLAEQLRPMAGGIKGIPHAAIRGDWGIFL-          216
Phatr2_45243    AREEF------------------------AALMERMKPLSDAAQALTSLALREDPAVVV-          209
Thaps3_11636    RVDTF------------------------MDI---CLDVHNG----F-VAL--GIYRLLP          219
Phatr2_54800    RARLY------------------------MDL---CTDINSG----F-TAL--GISRVLP          475
Thaps3_5859     DFGID------------------------EQSW-RKFDRKKAYAKFWAMVV--FSLKLFH          224
Thaps3_10254    KFPEE------------------------SDAIDKHFQLVQSTVKLFPVFM--GIKNLPT          278
                                                                         .

Thaps3_25361    KAAES--------FKSSDSNV-----YQQ-ASIRPASTFL-----N--KCLPLNTHVRAL          542
Phatr2_42980    EKVNS--------LRS--SPT-----YQD-SGIRYAQSFL-----N--KTFTINPHTRSL          499
Thaps3_5221     PLLRY---------LPSLISI---ISNGVEASTGPFAPYM-----NGPIFTVKDPWLRSW          261
Phatr2_54826    PLIRY---------FSTLVTL---LSQGSK-ATGTFASFI-----DGPNFTVTDPWLRSW          297
Thaps3_21847    T-MGS---------YVFKFLT---IGLQGTLLTGPFTECM-----N--LYGLNDRFNRQW          266
Phatr2_51868    S-LAN---------YMFKLLS---IGSKGALLTGPFSKVM-----D--LHGLKDPFVRKW          283
Thaps3_7094     STSQF---------LPNFAKL---NPLQNLKLTKPFSNII-----N--EAGVKDTFIRNW          256
Phatr2_54842    TAAPF---------LPNFTTL---NPLENLKLTQPFSAIV-----N--P-SVSNVFTRNW          269
Thaps3_10233    TTAGS---------LPFVVTH---PDVFG-DLSLTFDDLS-----KTVDEFVTVPFLRNF          298
Phatr2_9210     TTAGS---------LPFVATH---PDVFL-DLSLTFDSLH-----KTVDKIVTVPFLRNF          198
Thaps3_21900    TLILK---------YPLSFMN---VLKYAPAFTAPFD--L-----D--KLGVTNKFLRNY          255
Phatr2_45243    -TLLK---------YPRDLIA---TLAQGQALNEPFKNIM-----D--EMKIENKFVKNW          249
Thaps3_11636    SYLKFLMKDKVERLYKYGSMTVKDAQHAVLKLGYSKEELLK-NCPTAP-EMEDDPSIRRM          277
Phatr2_54800    SWMHFLVRSRIDRLMKFAAMTVRDVQYGMLNLGLTIEELLKDGCPPAPAGSEPDPSIRRL          535
Thaps3_5859     PMVLRLA-------WPFVC-----IPYRRCALRSTIDVLI-----------NDCGFSQEA          261
Thaps3_10254    PLFRLVM-------WLFDS-----K---LGVYRKTTKEVL-----------ESITSNRKL          312
                                                                         :

Thaps3_25361    MAAI--GMANENLSPDKTSMAAHVTNVCAMTSTEGYA--------YPVGGPRALCHALTS          592
Phatr2_42980    MAGI--GMKGENIRPGATSMAAHVTNISAALSGEGMH--------YPIGGPRALCRALAN          549
Thaps3_5221     LNALAFSLSG---LPADRTSAGAMAYVLFDMHREGAA------LDYPRGGLGEVVKALVN          312
Phatr2_54826    LDALAFSLSG---LPASRTAAAAMAFTLSDMHRPGAA------LDYPKGGMGAIAEALVR          348
Thaps3_21847    FDYLAFALSG---LDAAHTQAAPVAYTMIDLHKDGAV------LDYPKGGMDSMIQALVN          317
Phatr2_51868    FDYLAFALSG---VDASHTQAAAVAYMMMDLHKKDAV------LDYPMGGMDSLVQALVS          334
Thaps3_7094     LDVLCFCLSG---VPSDGTITAEMAMMMGEFYDEDAI------MDCPVGGASAIVDALVR          307
Phatr2_54842    LDLLCFCLSG---LPAKGTITAEMAMMMGEFYAPGAV------MDCPKGGAQSIVKALVR          320
Thaps3_10233    IDTMCI-FCG---FPAKGAMTAHLLYILERFFEETAA------FSVPIGGTCELGNTLQR          348
Phatr2_9210     IDTMCI-FCG---FPAKGAMTAHMLYILERFFEESAC------YSVPIGGTCEMGNTLVR          248
Thaps3_21900    LEMLAFLLQG---LPADQTLTVVMAYMVEDFFRENAV------MDFPKGGSGELMGALAR          306
Phatr2_45243    LDMLCFLLQG---LPASDTMNAVMAYMLADWYRPGVT------LDFPKGGSSSIVSALVR          300
Thaps3_11636    TAVLTHPIGDYAVQPRDATF--AAHGVTMAHYVNGSPNHNLVITKHTVGATQNISTRLTS          335
Phatr2_54800    KAVLTHPIGDYAVQPRDATM--AAHGVTMAHYQDGAC--------YCVGPTQQISVRSSS          585
Thaps3_5859     AGALTYHWGDHVVPPHRCPF--FMTALLDTHYKGGGY--------FPRGGSRSIAKCLVS          311
Thaps3_10254    QGVLSYHYGDYGEHPSRGAF--VMHSMICVHYRGGAY--------YPVGGPLSIAKSIAT          362
                   :     .                                      *    :

Thaps3_25361    VIEQ-----NGGRVVSGVLLQELLFEKLEKKEPKEETKDGES-KEPKPRCKGIRLENGL-          645
Phatr2_42980    VVLR-----SGGRVLTSVDVAELIFGEPREQASKGKQKEGDNDGPPPPRCVGVKLSDGR-          603
Thaps3_5221     GVEQK---SIGSKVHLSRHVESIDTNEE---------------G-DR--VIGLTVRKNGG          351
Phatr2_54826    GVQQG---SNGSQVHLRQPVEKIDFSED---------------G-TI--ATGLTLRNGR-          386
Thaps3_21847    GLEMKRDNVESGELRLKSRVERFVLNEV---------------K-NKATCTGVVLEN-G-          359
Phatr2_51868    GIKT-----NGGELRLNSRVERMILEDN---------------N-GRVECKGVVLTD-G-          371
Thaps3_7094     GIEK-----KGGKVFCNSRIDEICIENG---------------K-----AVGVRLAKNY-          341
Phatr2_54842    GIEK-----YGGEVVCNTHVQEIVVENE---------------K-----AVGVVIKQGK-          354
Thaps3_10233    GLEK-----YGGKLQLNAHVDEILVENG---------------R-----AVGVRLMN-G-          381
Phatr2_9210     GLEK-----FGGKIQLNAHVDEILVENG---------------R-----AVGVRLKN-G-          281
Thaps3_21900    GVTKR----EGCSVEVSTSVDEVIVENG---------------R-----AVGVKLAKSG-          341
Phatr2_45243    AVQK-----NGSSVCVNSHVDEILVENG---------------K-----TVGVRLTD-G-          333
Thaps3_11636    MVRS-----FGGEALIDATVRGIIIENG--------------------RAVGVKVSNTD-          369
Phatr2_54800    MVRE-----FGGEVLTDATVREIILEHG--------------------RAVGVRVSNTS-          619
Thaps3_5859     AITR-----RGGHVFALSPVDEILTKKN---------------MFGKFIATGVSVRGID-          350
Thaps3_10254    TIEK-----HGGKVLVRAPVSSVLVDEK---------------N----RAYGVVVKGKE-          397
                 :        .        :  .   .                        *: :

Thaps3_25361    --ELSVS-D-------------KGAVVSFMGMIPTFLQLVSPDVRTAEGV------PAGL          683
Phatr2_42980    --EIKFA-SDRFDE---KNGSCLPAVISMEGFIWTFINMLPDDIRMKYKV------PRGL          651
```

```
Thaps3_5221         KKVIVKA---------------KEGVVCNVPMWSLRKLIKNRNALSVLGGDKATSSSSGL     396
Phatr2_54826        ---RILA---------------REGVICNAPVWSLKSLRPTR------------------    410
Thaps3_21847        --TILKA---------------RRGVICNAPLWNMAKLLEDSITNPLDL-----------    391
Phatr2_51868        --TVVNA---------------RKGVVSNAPIWNMARILEDSVPGEVND-----------    403
Thaps3_7094         --SRIKA---------------TKGVISNLSVWDLMNSGIV---D---------------    366
Phatr2_54842        --QRVAA---------------SKAVISNLSVWDLFGSGIL---D---------------    379
Thaps3_10233        --NVVKA---------------RKAVVSNATPFDTVKLMPKAEGEPKG------------    412
Phatr2_9210         --NVVKA---------------NKAVVSNATPFDTVKMLGEKQALPEG------------    312
Thaps3_21900        --RIIKA---------------KEAVISNADLYNTYKFVPEGKHEGFD-KER--------    375
Phatr2_45243        --RKVHA---------------TQAVVSNADPYISNKLLLNARKSGQLNKAA--------    368
Thaps3_11636        --ELEECTSEEDLAKVPAV---------------EYNKFLPQDLPVV-------------    399
Phatr2_54800        --ALAECKSDAERAQVPVTELRAKAVVCATSVYNLYNNLLPQDLAQV-------------    664
Thaps3_5859         --IVVKKC-----------------VVSDAGFLNTFGIDSEGNKPALVDSNAAASQR---    388
Thaps3_10254        --VLAK-T-----------------IVSSIGAPATFGKLLPESHRHLV------------    425


Thaps3_25361        PALEERRPLMRVMISLKGNKDDLNLTGADWYRLPNATLPRDELDPMTGQVKFGTIGVDDD    743
Phatr2_42980        PALSSRRPVFKVLFALKGSADQLNVTGADYYRLPNAAVARDEFDQSSGQIKHGEIGWSDS    711
Thaps3_5221         KAKQSWMTSFDTD-PSTGRGSVLRPKPAEDTTIEKSLLEKCDSAEMTGSFLHLHLALNAT    455
Phatr2_54826        -----------------------------SGEADETLLGACDTAEKTGSFLHLHLALESS    441
Thaps3_21847        ----------------------------SVAAAVNDVRSQANEMEMTGSFMHLHLGIPND    423
Phatr2_51868        ----------------------------ARRSIVKAIQKQADDMSMTGSFMHLHLGIPKA    435
Thaps3_7094         ---------------------------TDLF--PEDFVKERKATPACPSFMHLHVGFQIT    397
Phatr2_54842        ---------------------------TTLL--PNSLVQQKLSTPLGKSFMHLHVGFRMS    410
Thaps3_10233        ---------------------------------LTKWREELGKLPRHGAISHLFLAIDAE    439
Phatr2_9210         ---------------------------------VAKWKEELGKLPRHGAIMHLFLAIDAK    339
Thaps3_21900        ---------------------------IEYL--GLTAKPKDGSVPFCKSFMHLHLAVKAE    406
Phatr2_45243        ---------------------------TDHLDALINTDKTEGGIADLKSFIHIHAGIDAA    401
Thaps3_11636        ---------------------------------KKFKDE--ATIRQSNGHVFLFCKLRGN    424
Phatr2_54800        ---------------------------------KEFQDPEKRTIQQSNGHIFLFCKIKGD    691
Thaps3_5859         -----------------------------ALLHNAKGFPTLDSVTPCISNLSLFIGLDRT    419
Thaps3_10254        ---------------------------------SKQLESMKDNMIASNLTLMSMFVGISDP    453


Thaps3_25361        NTGASEELILGEATDETEATT----SHTRGKRNKAATSKAPRSKFTSGVSWMKVSFPSAK    799
Phatr2_42980        DTGDNGDAYADGGKNLMDVINQDPGSISDEHIVNSSRKRARKTKFEAGSSWLHVSFPSAK    771
Thaps3_5221         GLDLQS---LE-PH-YTVMDRGL---EGDG-KV--IDG--VKDDSSGELNMIAVSNPCVL    502
Phatr2_54826        GLNLDN---LE-AH-YTVMDRSL---GGDG-SS--VNG--VLDGPCGILNMIAVSNPCKI    488
Thaps3_21847        GLPA----DLD-CH-HSVLNLEH--------------------DVTAAQNLVIVSIPTIF    457
Phatr2_51868        GLPE----HLE-CH-HSVLNMQD--------------------DVTAEQNMVIISIPTVF    469
Thaps3_7094         KEELSK---LQ-AH-YIFMNDWE---R----------------GVTAEENCALVSIPSVH    433
Phatr2_54842        KGELQT---LQ-AH-YMHMEDWG---R----------------GVQDEDNAVLVSIPSVH    446
Thaps3_10233        GLDLSHI--QD-PA-HLVVQDWD---R----------------SLQDSQNLCSFFIPSIL    476
Phatr2_9210         DLDLSHI--QD-PA-HLVVQDWG---R----------------SLQDSQNLCSFFIPSLL    376
Thaps3_21900        LIPE----DAP-PQ-WTVVQDWD---K----------------GIDATGNVVVVSVGSKL    441
Phatr2_45243        GLPDQPSADFP-AQ-WAVVRDWD---A--------------PEGVESPRNIVLCSMPSLI    442
Thaps3_11636        ADEIG----LP-DHNLWYFNGYD---LDDA-FDKYFAN-----PTEVRPPTVYIGFPCTK    470
Phatr2_54800        PTELK----LP-AHNLWYFNSYD---IDDA-FEAYFTD-----PVGQRPPTVYIGFPCTK    737
Thaps3_5859         DEELE----LP-AQNVWHVHDWD---HDAA-WKNMMNAISPYQSLADQTPFLFISNESAK    470
Thaps3_10254        ENSLA----LP-KRNYWIHDSWD---HDKN-IE-------AFKKNPTKPPVFFVSFSSAK    497


Thaps3_25361        ------DPSWQDRHGDVSTCVVTVEA-DDDFVQMFDTKPKIYSV--------------LK    838
Phatr2_42980        ------DPSFEERHGKTTTCVVTIEA-DDDFVTYFDTKPKIYVI--------------KN    810
Thaps3_5221         ------DNTLAPEGFIIMHAYG---AGNEPFEIWKPPTASKGNASPNTAGEGEIIGGERC    553
Phatr2_54826        ------DNSLAPDGTIVVHAYS---AGNEPYEIWEGLDR---------------------    518
Thaps3_21847        ------DPSLAPEGYHIIHAYT---AASEDFADWERMLIGELDGGK-------PEFTDYK    501
Phatr2_51868        ------DPSLAPEGYHVVHAYT---AACDGFDQWTPYLDSGKETG--------------K    506
Thaps3_7094         ------DNTLAPDNHAVLHIYT---PATELYERWENVKRNT-------------------    465
Phatr2_54842        ------DDTLAPEGYAVLHIYT---PATEDFTRWENVQSK--------------------    477
Thaps3_10233        ------DKTLCPEGKHVIHVYS---SGGEPYEPWEKLTPGS-------------------    508
Phatr2_9210         ------DKTLCPEGKHVIHVYS---SGGEPYEPWEKLKPGT-------------------    408
Thaps3_21900        ------DQSLAPPGYHVIHAYT---AGNESYEDWEQFEHLMDDAA------------VRD    480
Phatr2_45243        ------DPSLAPEGKHVLHAYV---PATEPYADWAGMDRKS-------------------    474
Thaps3_11636        FGLIRQDITWQKRFPNVSNCILISDGLYEWFEQWSDKPV-------------------RN    511
Phatr2_54800        ------DTSWKQRFPGVSNCILISDGLWEWFEKWQDKPV-------------------HN    772
Thaps3_5859         ------DPDFGTKHPGKATSEVFAVCKYDLFEKWADTAH-------------------NS    505
Thaps3_10254        ------DPTYSSRNPGKQVALVVGPGFFDHVAVFQNERV-------------------KH    532
                          *                     :

Thaps3_25361        A---GNGERERLRDRVLKDLLETFPQLQGQLE----TVQICGPVR---------------    876
```

```
Phatr2_42980      A-SATKGDLDRLLERVKKDVYHIFPQLRDKVD----HCEICGPFQ---------------      850
Thaps3_5221       SPSTYQALKDSRSKVLWRAVESVIPDARERTV----LALIGSPRT---HERFLRRPCG-S      605
Phatr2_54826      RSDGYMCLKEDRAEVLWRAVESIIPDARNRVV----ISEIGSPIT---HERFLNRPRG-T      570
Thaps3_21847      RTKAYKDLKQEKAEALWLALERIIPDVRERAKREGSVVEVGTPLT---HRRYNRRYRG-T      557
Phatr2_51868      VVDGYNELKDEKADVLWRAVERVIPDVRLRAKQKGSIILVGTPLT---HRRYNQRYRG-T      562
Thaps3_7094       --PEYNQLKEERSAFLWKVLEKIIPDIRQRAV----HSKVGTPLT---HQRFLNRYRG-S      515
Phatr2_54842      --EAYEKLKEERSQYLWKVLTRIVPDIRERAR----IVRVGTPLT---HQRFLRRYKG-S      527
Thaps3_10233      --EEYEAYKNERAEVLWRAVERCIPDVRDRVE----FSIVGSPLA---HEAFLRRDRG-T      558
Phatr2_9210       --QEYDDYKNERAKVLWEAVERCIPDVRDRLE----FSIVGSPLA---HEAFLRRDRG-T      458
Thaps3_21900      KDAAYQTFKDERAQPIWDAIQKRAPAVVKGA-C--VIEKVATPLT---HARFLNRHRG-N      533
Phatr2_45243      --EEYTKKKEQAADFLWSAIEEYIPNARDRAVP--GTVQIGTPLT---HERFLRRTRG-T      526
Thaps3_11636      RGEEYLEFKDKLTHHLLDILQEFVPQVKGRIE----YHHLGTPLS---EETFLASYRGGS      564
Phatr2_54800      RGSDYEEFKEKLSKHLLEILFEFVPEVKDKIE----FSFLGTPLS---EQTYLNSFCAGS      825
Thaps3_5859       RGDDYTELKEKIIESYLNVFYLHFPKTKGHEG----NLAKCTVLTMVSADSMA*------      554
Thaps3_10254      RGKEYTDMKKEWEIVYMEAFLKQFPELKDKVD----YVEFGTALS---NDFYLGTNRGAV      585
                            .         .    *

Thaps3_25361      SGLTHNGPRF-----------AIKGNRPETPYPGLYIGGADLTVGDSFSGAIVGGWLAAN      925
Phatr2_42980      KGLSHNPERF-----------AAKGIRADTPYPGLFVGGSDLTVGESFSGDIVGAWLAAN      899
Thaps3_5221       YGAAFED----------------CLKDGSTPISNLVLSGDGVF--PGIG-------IPAV      640
Phatr2_54826      YGSATED----------------YLADGSTSIGNLLLAGDGIF--PGIG-------LPAV      605
Thaps3_21847      YGPAPSNGND-----------VWELPGPKTPIEGLLACGDCCF--PGIG-------LPGV      597
Phatr2_51868      YGPAPGPGKD-----------VWELAGATTKIKGLLACGDSTF--PGIG-------LPGV      602
Thaps3_7094       YGPAIRAGDA-------------SFPFPNTPIQGLLLCGDSCF--PGIG-------VPAV      553
Phatr2_54842      YGPAIQAGVG-------------SFPFAGTPIRQLLTCGDSCF--PGIG-------VPAV      565
Thaps3_10233      YGMAWAAGSSAPQSGILGSVLPFPFPFPNLKTPVDGLLRCGDSCF--PGIG-------TPSA      609
Phatr2_9210       YGMAWAAGTSAPQAGLLQNILPFPFPFPNLKTPVDGLLRCGDSCF--PGIG-------TPSA      509
Thaps3_21900      YGLAIAPDNA----------EGWKFPDVKTPLEGYYRCGDSTT--SGIG-------VPAT      574
Phatr2_45243      YGPRVEV--G----------AGQTLPGHKTPLPGFYMVGDFTF--PGIG-------VPAT      565
Thaps3_11636      YGTQCVTEMF-------APINRNWTTTPFTEVPGLYLAGSDAFL-PSVTGAMYGGCLSAS      616
Phatr2_54800      YGTKCLPSMF-------AKSNRRWTTSPHTSIPGLYLAGSDAFL-PAVCGAMYGGCFGAI      877
Thaps3_5859       ------------------------------------------------------------      554
Thaps3_10254      YGLSHTPERF-------N---LQWL-KPKTPIQNFYLTGQDVCS-CGITGALVGGYLSAY      633

Thaps3_25361      AIMGYSFM---------DHMY--L-GKN-ITSDL-----------------QQFIEEPIL      955
Phatr2_42980      AVEQYGPL---------DHLF--L-QKN-ITTDI-----------------EQFLEEPGW      929
Thaps3_5221       ALNGASAANGF--VGIFDQWR-CM-DYLKAKGIIA*------------------------      671
Phatr2_54826      AISGASAANAM--VSVFKQWE-CL-DELGKSQKL*-------------------------      635
Thaps3_21847      AASGTIAANTL--VDSSVQLD-LM-SELKDSGALQ*------------------------      628
Phatr2_51868      AASGTIAANTM--TTIANQRN-LM-KELKAKGALQ*------------------------      633
Thaps3_7094       AGSGMIAANSVSLDSIGAQLE-VL-SKIKQQ*----------------------------      582
Phatr2_54842      AGSGLLAAHSVSWDSIGPQQD-LL-KTLQKRK*---------------------------      595
Thaps3_10233      AASGAIAANTM--THVDNHLK-ML-SEASKLDPMYKFLDAGIMGQVYKPLVQGFTPSPEL      665
Phatr2_9210       AASGAIAANTM--NPVGKHLD-LL------------------------------------      530
Thaps3_21900      ASSGAVCANAI--MSVWDQLS-LN-QKIKMP*----------------------------      601
Phatr2_45243      AASGAIAANTL--VSVFDHLA-ML-DKVRLPEKEQKS*----------------------      598
Thaps3_11636      AVLGTLGTMRLGH-AILTHLAMRLREENPKLSKIE----------AYMLAVKKFTE*---      661
Phatr2_54800      AVLGHLRALKLTL-AFIAHFAGCITDEDPKIGWIQ----------AYILAWKKFMND*--      923
Thaps3_5859       ------------------------------------------------------------      554
Thaps3_10254      AISPRCFLR--TA-SLLN*-----------------------------------------      648

Thaps3_25361      ATERNGVIVDDVAVPFKEVVVDMQKGITDADRSTAAESSKEE*      997
Phatr2_42980      VDEE------DVAIPYKSADAKKDKDV*---------------      950
Thaps3_5221       -------------------------------------------      671
Phatr2_54826      -------------------------------------------      635
Thaps3_21847      -------------------------------------------      628
Phatr2_51868      -------------------------------------------      633
Thaps3_7094       -------------------------------------------      582
Phatr2_54842      -------------------------------------------      595
Thaps3_10233      RTDQYVSGAGVAPVDYTATDPSVSERIDL*-------------      694
Phatr2_9210       -------------------------------------------      530
Thaps3_21900      -------------------------------------------      601
Phatr2_45243      -------------------------------------------      598
Thaps3_11636      -------------------------------------------      661
Phatr2_54800      -------------------------------------------      923
Thaps3_5859       -------------------------------------------      554
Thaps3_10254      -------------------------------------------      648
```

2)

### ZEP Percent Identity Matrix

| | | | | | | | | | | | | |
|---|---|---|---|---|---|---|---|---|---|---|---|---|
| 1: Phatr2_47925 | 100.00 | 31.88 | 32.79 | 20.94 | 21.18 | 19.89 | 21.58 | 21.35 | 23.27 | 20.05 | 22.40 | 19.67 |
| 2: Thaps3_6395 | 31.88 | 100.00 | 46.63 | 22.93 | 18.67 | 18.94 | 21.71 | 26.03 | 24.36 | 20.46 | 21.14 | 19.61 |
| 3: Phatr2_45936 | 32.79 | 46.63 | 100.00 | 21.19 | 22.89 | 19.45 | 19.89 | 25.15 | 22.67 | 19.10 | 19.40 | 19.32 |
| 4: Thaps3_1961 | 20.94 | 22.93 | 21.19 | 100.00 | 22.54 | 22.92 | 24.21 | 25.13 | 24.02 | 22.81 | 26.11 | 22.77 |
| 5: Thaps3_270370_ZEP1_ | 21.18 | 18.67 | 22.89 | 22.54 | 100.00 | 65.42 | 33.85 | 33.48 | 33.67 | 20.00 | 20.93 | 21.04 |
| 6: Phatr2_45845_ZEP1_ | 19.89 | 18.94 | 19.45 | 22.92 | 65.42 | 100.00 | 33.41 | 35.44 | 35.52 | 19.56 | 22.22 | 22.65 |
| 7: Phatr2_56492_ZEP3_ | 21.58 | 21.71 | 19.89 | 24.21 | 33.85 | 33.41 | 100.00 | 44.88 | 42.80 | 20.00 | 21.03 | 22.19 |
| 8: Thaps3_261390_ZEP2_ | 21.35 | 26.03 | 25.15 | 25.13 | 33.48 | 35.44 | 44.88 | 100.00 | 78.51 | 21.62 | 23.16 | 25.08 |
| 9: Phatr2_56488_ZEP2_ | 23.27 | 24.36 | 22.67 | 24.02 | 33.67 | 35.52 | 42.80 | 78.51 | 100.00 | 21.76 | 23.33 | 23.84 |
| 10: Thaps3_22671 | 20.05 | 20.46 | 19.10 | 22.81 | 20.00 | 19.56 | 20.00 | 21.62 | 21.76 | 100.00 | 25.82 | 24.67 |
| 11: Thaps3_20663 | 22.40 | 21.14 | 19.40 | 26.11 | 20.93 | 22.22 | 21.03 | 23.16 | 23.33 | 25.82 | 100.00 | 44.80 |
| 12: Phatr2_43425 | 19.67 | 19.61 | 19.32 | 22.77 | 21.04 | 22.65 | 22.19 | 25.08 | 23.84 | 24.67 | 44.80 | 100.00 |

### ZEP Alignment

```
Phatr2_47925              ------------------------------------------------------------ 0
Thaps3_6395               ------------------------------------------------------------ 0
Phatr2_45936              MADQSAARKTLPSLLRHFEGTELTVELKTGRLYRGTLSSADQAMNLTLEDASLLQRLIVN 60
Thaps3_1961               ------------------------------------------------------------ 0
Thaps3_270370(ZEP1)       ----------------------MTV---------RRIASLAIGISLSTLTCAFVTIS--- 26
Phatr2_45845(ZEP1)        -----------------------------------------MKFSTTVSSALFLIASV-- 17
Phatr2_56492(ZEP3)        ---------------------------------------------MK-RSCSIVTILY-- 12
Thaps3_261390(ZEP2)       ------------------------------------------------------------ 0
Phatr2_56488(ZEP2)        -----------------------------------------MGLSFL-SLCAVLTASS-- 16
Thaps3_22671              ------------------------------------------------------------ 0
Thaps3_20663              --------------------------------------MVSTILIFILVACLLLQST--- 19
Phatr2_43425              ------------------------------------------------------------ 0


Phatr2_47925              ------------------------------------------------------------ 0
Thaps3_6395               -------------------MSSSDRHSSQLNQPKRPRHEEPSHSI-----MAFDL--KN- 33
Phatr2_45936              QQHKGAFRRGSSSAAVPSTLSLVHIRGSTIRFIHFPDQLDLTLTIKQGIDRSWRMEHNSE 120
Thaps3_1961               ------------------------------------------------------------ 0
Thaps3_270370(ZEP1)       -----------SSRTTIKPLNVVGEQASSIGPATLLRNLKQNL-----PQIDWLAEGKGS 70
Phatr2_45845(ZEP1)        -----------STTTSFTPVQSFGVHR--------------------------------- 33
Phatr2_56492(ZEP3)        -----------VATT--------------------------------------------- 16
Thaps3_261390(ZEP2)       ------------------------------------------------------------ 0
Phatr2_56488(ZEP2)        -----------AMAF--------------------------------------------- 20
Thaps3_22671              ------------------------------------------------------------ 0
Thaps3_20663              -----------CDAFTF------------------------------------------- 25
Phatr2_43425              ------------------------------------------------------------ 0


Phatr2_47925              ------------------------------------------------------------ 0
Thaps3_6395               --RSPYEKKMERVITACPKCNGEGKVRAP---------LSKKARAQRKRMQQSQTGDTTN 82
Phatr2_45936              PGWDETTTIHEGPFTVCPKCHGDGHIVHQ---------ASKKQKLRHKRART--NGDYTD 169
Thaps3_1961               -----------------MPCSQLLSTAFNNDYNHLI--------------------TSQH 23
Thaps3_270370(ZEP1)       PSNK--IDIPDHVATVLAQPNAPKREAESEERTHKIRSRAKQASEDA--MALRGMLIG-D 125
Phatr2_45845(ZEP1)        --RT--LL----VTPRHATVEPPVREPETSDRVRQVRDRFRKASQDA--ANAKGCVAQDD 83
Phatr2_56492(ZEP3)        ------------VRAFAPAP--LVQ-----------SSCFFQRQPTT--TAR--FVSGTA 47
Thaps3_261390(ZEP2)       -------------------------------------------------MAD----DEAD 7
Phatr2_56488(ZEP2)        ------------VTTRSPACNDVTRSLH-RINTRHMTYPFYPASSLR--IST--RVASTA 63
Thaps3_22671              -------------------------------------------------MADSPTASSEE 11
Thaps3_20663              ------------------PSSGVLRDVRVSINGSVERRCSLPASEHVQHVAASSTSSSSS 67
Phatr2_43425              ------------------------------------------------------------ 0
```

```
Phatr2_47925                --------------------------MGKKRQSRPRLDPGAHIAIIGSGLAGLSTALSLE 34
Thaps3_6395                 APNLAILKKPCKECDGSGLIAINPLDTTERKQTPPQIQPNFSVAIVGGGIGGIALAAALQ 142
Phatr2_45936                TPAP-QRLETCRECDSSGLVQSDT--------DPPVDTTLPEIAVVGGGLAGLALAAACR 220
Thaps3_1961                 I----------------------NHI-MFIKQATLNHAKYASVAVVGGGIGGLTAANALL 60
Thaps3_270370(ZEP1)         DDAN-------------AWWR-EQRS-IPEGGRVVTTDDPLTVLVAGGGLAGLVVAAACH 170
Phatr2_45845(ZEP1)          GDES-------------SWWR-K--P-LPEDNDVISNQRPLRVVIAGGGVAGLVTAAACH 126
Phatr2_56492(ZEP3)          PPS-------------------SNVA-SEEKVDAISEAHRLKVLIAGAGVGGLSLAKVLT 87
Thaps3_261390(ZEP2)         ADF-------------------NSSD-YELLGRPARPGRPLKVAIAGGGVGGLTAALCML 47
Phatr2_56488(ZEP2)          VPP-------------------EDVA-FDKLSLPAREGRPLKIAIAGGGVGGLTTALCML 103
Thaps3_22671                VPS-------------------APPH-TTV--DDISNLEHHPLVIIGGGIGGLVLALCLD 49
Thaps3_20663                TRS-------------------RSST-TTLQAATAPHQPVQKVAIIGSGIAGLALAHAFL 107
Phatr2_43425                ------------------------------------------------------------ 0

Phatr2_47925                QG--------------------GFTNVHIYERDGSHDARKEGYGLTLTYNPTGVLHQLNV 74
Thaps3_6395                 HR--------------------NIP-CIVYERDLSFEERKQGYGLTMQQGARALR-SLGF 180
Phatr2_45936                HR--------------------GMK-YTVYERDLDFHQRSQGLD---------------- 243
Thaps3_1961                 NK----------------NPNLIER-LTVYEQAKEFTPT-AGAGFGFSPNGQICLSSIGI 102
Thaps3_270370(ZEP1)         S-----------------K---GMK-VALFEQASSYAPY-GGP-IQIQSNALRALQQINP 207
Phatr2_45845(ZEP1)          A-----------------K---GMQ-VAIFEQASQYAPY-GGP-IQIQSNALRALERINP 163
Phatr2_56492(ZEP3)          KM----------------P---TMD-VTVLEQTSEFKRF-GGP-IQLASNAMEILKHMDK 125
Thaps3_261390(ZEP2)         K-----------------K---GFD-VTVYEKTAAFARF-GGP-IQFASNALSVIKEIDE 84
Phatr2_56488(ZEP2)          K-----------------K---GFD-VTVYEKTAAFARF-GGP-IQFASNALSVLKEIDE 140
Thaps3_22671                QVYNHSITTDDANGEPITSSSVKFP-IHVYESTAEYSAN-AGGAIGLYPNGLRVLRNLSR 107
Thaps3_20663                S------------NNPSSSNNNKIQ-IDIFDSRTNLDEK-AGSGIQLT-GGLVALNEISN 152
Phatr2_43425                ------------------------------------------------------------ 0

Phatr2_47925                LE-----EIA------------------------QSDCPSRSHYM--FNA-NGEIQGYFG 102
Thaps3_6395                 FSFSDDGEDD----NNNCSGKKA-VDENTSNTKQKFGIHSTRHVV--HKP-DGTVVGEWG 232
Phatr2_45936                ----------------------------------SDGIMSTKHVV--HEP-DGAIVGEWG 266
Thaps3_1961                 YGYKKFILPFNSM--------KR-LNKEGNL----------------VNQSDV------- 130
Thaps3_270370(ZEP1)         EIFQELVTAGTCTADRVSG-LKIGYKKGNK--LA--------GL---YDAGDW------- 246
Phatr2_45845(ZEP1)          VICEEIRKAGTVTADRVSG-LKIGYKKGVFLGLG--------KQ---YEKGDW------- 204
Phatr2_56492(ZEP3)          PVFDKVMEKFTFTGDKENG-IKDGIRT------------------------EW------- 153
Thaps3_261390(ZEP2)         ELFERVMDKFTFTGTRACG-IKDGLRADGSFRMTNDSLDYLWNP---EAPADW------- 133
Phatr2_56488(ZEP2)          TLFERVMDKFTFTGTRTCG-IKDGLRADGSFRMTEDRLDYLWNP---DAPADW------- 189
Thaps3_22671                GSSPSYLDSEHVKFGANCPNLLQNVR--------TAGCDYIYRRWMRHDGLQVAVAREDE 159
Thaps3_20663                NLYNEVVESS----------LPL------------ERLVSKCRPWFGGNKDDAGVEQGWQ 190
Phatr2_43425                ------MDAG----------LLQ------------TGVRSRCKPWNPASPFDT------- 25
                                                                               :

Phatr2_47925                NA-----------F-------------------------------------------A-- 106
Thaps3_6395                 MK-----------V--------------------------------------WGGRFE-- 241
Phatr2_45936                LR-----------K--------------------------------------WGRSER-- 275
Thaps3_1961                 LR-----------EL--------------------------------------------- 134
Thaps3_270370(ZEP1)         LV-----------RFDTIGP---------------------------------------- 255
Phatr2_45845(ZEP1)          LV-----------RFDTLQP---------------------------------------- 213
Phatr2_56492(ZEP3)          YA-----------KFDLKTP---------------------------------------- 162
Thaps3_261390(ZEP2)         FV-----------KFPLRQC---------------------------------------- 142
Phatr2_56488(ZEP2)          FV-----------KFPLKQC---------------------------------------- 198
Thaps3_22671                LLPDIKVDESEMAKLEVLDESEKGTGSRSSAVSRADSTKSQDVEGERANRRPHGGSFANA 219
Thaps3_20663                LL-----------ELDIQNA---------------------------------------- 199
Phatr2_43425                LL-----------DLDLLKT---------------------------------------- 34

Phatr2_47925                ---------------------------------------R-----NRGWGQRGNLRVPRQ 122
Thaps3_6395                 ---------------------------------------KN----GRKHAKRQNAHISRQ 258
Phatr2_45936                ----------------------------------------------AKKPKRQNIHIARQ 289
Thaps3_1961                 ---------------------------------------------SNRHGFGIAGC-LRS 148
Thaps3_270370(ZEP1)         ---------------------------------------------ALEAGLPATVVVDRP 270
Phatr2_45845(ZEP1)          ---------------------------------------------ALDAGLYPTVVVDRP 228
Phatr2_56492(ZEP3)          ---------------------------------------------AENRNMPYTGVIERP 177
Thaps3_261390(ZEP2)         ---------------------------------------------ADLFGLPYTGVIDRP 157
Phatr2_56488(ZEP2)          ---------------------------------------------ADLFGLPYTGVIDRP 213
Thaps3_22671                MGALEAMKDMSQNLSQRLSRISFTGSDATTTSAAGGSDKSTPRASRVVDTELLSLGIRRW 279
Thaps3_20663                ---------IRENA--------AADASKQHGAEEGDSNKQY--SLVREDGEVVAYTILRG 240
Phatr2_43425                ---------VQNA----------------------GSDV-S--NALIREGKLVWTSIMRG 60
                                                                                  *
```

```
Phatr2_47925              RVRQILASRL---KITETHWDHKLVGVSGCENGE-------NICLAFQLEGA-AEEKLLV 171
Thaps3_6395               NLRQLLMEML---HPGTIQWGQKFVGYSGQSSDDDSSQDQPSLQVRFRRRSNDCDEEVAT 315
Phatr2_45936              SLRWQLYKAA-GGRTANIAWNHRLLQYKQRV-------DAPGWELKFQV----DDQIIAH 337
Thaps3_1961               DLVNLLVEQL-----DTQHGGKGALKYSEKLVGINPIH--DKVELEF------ESGRQD- 194
Thaps3_270370(ZEP1)       VIQQILVKYG--FPEGTVRIKSRIQSYEDL-----GKG--RGVSVTL------EDGTKA- 314
Phatr2_45845(ZEP1)        VIQQILLEHG--IPEKTVRIKSRIANYEEL-----GPG--KGVRILL------EDGTVA- 272
Phatr2_56492(ZEP3)        DLQQIFLDSLPK---GTVKNGDGVARYEKL-----PDG---GVKAVL------KSGKEV- 219
Thaps3_261390(ZEP2)       DLQEILLDECRKIKPDFIQNGNPVNGYVSK-----GKG--NGVTVNL------ADGTTA- 203
Phatr2_56488(ZEP2)        DLQEILIDECRKLKPDFLINGNPVVGYEDL-----GKG--QGVTINL------NDQTTA- 259
Thaps3_22671              KYQQVLYDQC-KEVGIQFHMGKRLQSVTSIPASGEDGD--AKSLLLF------KDGSRI- 329
Thaps3_20663              TLQRILREQLAQEHGVDVQFDKRLCGMAY---SNEENG----VKCQF------NDGTTTG 287
Phatr2_43425              ALQEALYGALPSNVRQNVQFGKVLVDLR----SVREGG----IECLF------SDGSVAG 106
                               :                                        :       .

Phatr2_47925              GADLVVAADGIRSAVLQHAYP-QAPPI--------------------------------- 197
Thaps3_6395               TASVLVGCDGIRSSVRSAKLGEDGTPL--------------------------------- 342
Phatr2_45936              KADLIVGADGLRSQVRRSLIGEDRTPL--------------------------------- 364
Thaps3_1961               LVDLVIGADGINSSVSKLLNIDDEIA---------------------------------P 221
Thaps3_270370(ZEP1)       YADVLVGADGIWSQVRKNLHGLDDGAGGFAASGAAGGALDDAEARKLARDTVAIAAKADR 374
Phatr2_45845(ZEP1)        YADVLIGSDGIWSSVRRIMHGLDQGADGFAASGAAGGALNEAEARRMAKDSVLMANNANR 332
Phatr2_56492(ZEP3)        YGDVLIGADGIWSAVRATMRDS-----------------------------PAKGDGSGA 250
Thaps3_261390(ZEP2)       EADVLVGSDGIWSAIRAQMYGEEI-KKS----------------------SNNALKRQGC 240
Phatr2_56488(ZEP2)        SADVLVGSDGIWSAVRDQMYKEGGVKST----------------------SANKKKRQGC 297
Thaps3_22671              TASLVIGADGINSKVRNYVTNPKPTAAT----------------------TKQQEEYVP 366
Thaps3_20663              PYDLVVGCDGIQSKVKQYVNTGSLQPNA---------------------------DSSSA 320
Phatr2_43425              PFDVVVGCDGIKSACKEYVENGRILPKD----------------------AKREGDSVA 143
                            .:::..**: *

Phatr2_47925              QSLGIRLILGISSSF----------------THVHLKER--GFYTLDSGKRLFVMPFART 239
Thaps3_6395               RYLDCIVILGIAPSP---------------TSALTDGET--VFQTADGITRLYVMPFAEA 385
Phatr2_45936              RFLDCIVILGICPIAGISL--------EGQQSDLLDGET--VFQTADGVTRIYIMPFTTT 414
Thaps3_1961               IYSGANIFYGKIPNPDGHE--------YLRGHPIFTEG------SVTNGPGTGEFI---- 263
Thaps3_270370(ZEP1)       RFSGFTCYAALAPHRASNI--------ENVSYQILLGEK-KYFVSTDGGGDRQQWF---- 421
Phatr2_45845(ZEP1)        RYSKFTCYAALTEHRASNI--------EEVSYQILLGKD-KYFVSTDGGGERQQWF---- 379
Phatr2_56492(ZEP3)        TYSGYTVFAGELAYDSFDN--------GQVGYKVYIGPG-QYFVITDIGNGNYQWY---- 297
Thaps3_261390(ZEP2)      TYSGYTVFAGETVLKTEDY--------YETGYKVYIGPQ-RYFVTSDVGDGRVQWY---- 287
Phatr2_56488(ZEP2)        DYSGYTVFAGETILKTPDY--------YATGYKVYIGPK-RYFVTSDVGDGRIQWY---- 344
Thaps3_22671              AYTGVTCLMGCASVPRIRGICFPSSAT-TKCHACYYPTRAPKEVDDEGNADDTVRPVSGD 425
Thaps3_20663              IYSGIRITFAIQEGDADDN---PVQAKKGAQFTQFFGNG-AYALTSSYGAGKGVPPAKG- 375
Phatr2_43425              VYSGLRIRYAVKDGNSEEK---QA---ETATLSQYFGEG-AYGLDGIYGAGPGQPHTKC- 195
                                   .

Phatr2_47925              AEWAVIDDNATSCSQPTGEQYMWQLSFASSDDKTY-----SSTEL----LQQALFHCRDW 290
Thaps3_6395               GD-----DSSGLSTDNTKGLSMWQLSFPMDETDATRLSQLGSSAL----KEEALKRCGAW 436
Phatr2_45936              -------------------AYMWQLSFPMAEEQALSLSSKGPSAL----KKEAIRRCQSW 451
Thaps3_1961               ---AF----HTGAEDN--KTFIWANTY--ASNSPPPK-------------------REDW 293
Thaps3_270370(ZEP1)       ---AL----IREPAGGVDPEPT--------PEDPHPKLTRLRKEFACNGS--GDADGNVW 464
Phatr2_45845(ZEP1)        ---AL----IREPAGGVDPEPT--------PENPTPKLTRLLQEFNHEEP--GDQNGDVW 422
Phatr2_56492(ZEP3)        ---AF----LARPADSASST------------DMPDGQSKYLQEI-----------FAGW 327
Thaps3_261390(ZEP2)       ---AF----FALPPGTKKAPSGWG---------------DYIKSL-----------HQGW 314
Phatr2_56488(ZEP2)        ---AF----FALPPGTKKAPSGWGGSTRDGQTDPEENLVDYVKGL-----------HEGW 386
Thaps3_22671              YEQVF----QIYFPSPIERPDTWRTLTP-------TEAKEECRELA------KKLREDGW 468
Thaps3_20663              ---AF----LIYTDDDYYGP-----FPK-------SVFKRDESEVAVKPSVEAAAENADW 416
Phatr2_43425              ---AF----LVYLDPDYVGP-----FKK-------KRAR-----LESKPS---VDENADW 228
                                                                                   *

Phatr2_47925              HEPVQE----------------------LMLS-TSQESIWGTLLYDRNP---EILHKH-- 322
Thaps3_6395               HDPILK----------------------LLRS-TPEDFITGYPCYDRALVERKELRDG-- 471
Phatr2_45936              HTPVPD----------------------ILYS-TPIELVSGYPVYDRALLTPELLQE--- 485
Thaps3_1961               SEGNFHELKDILLKYPTS-----HPIHKFAEL-TGESDLLHFGLYYR----------HHK 337
Thaps3_270370(ZEP1)       DPFALE----------------------LINA-ASEEDIKRRDLYDGAPLLTTLDPQRLL 501
Phatr2_45845(ZEP1)        DDFAYE----------------------LFKA-TPEEDIKRRDLYDGSPLL--------M 451
Phatr2_56492(ZEP3)        SEEVHH----------------------ILRA-TQEHEIEQRDLYDRPPSA--------M 356
Thaps3_261390(ZEP2)       SDEVMT----------------------VLDS-TPPDSVEQRDLYDRPPEL--------L 343
Phatr2_56488(ZEP2)        SDEVMM----------------------VLDS-TSPDSVEQRDLYDRAPEL--------F 415
Thaps3_22671              DEQFLAPLESETL--TGVL-RVGLRSRE------------ALDV-WHVGG---------S 503
Thaps3_20663              TQDNRVPREHVAE-CIKVLKTAAIPGNDVADIVSNSNRFFDLGVYFHNPFS--------W 467
Phatr2_43425              TQDVRKSIEVARETMLDQTKSLGVPDTDLSPTISAADRFFELGVYFHNPFS--------T 280

Phatr2_47925              ----LQINDTLPRRILIVGDACHAMSPFKGQGANQALQDGRVLVKHLTSARVE------- 371
```

```
Thaps3_6395              ----CDKSQSANAFVTLLGDACHPMSPFKGQGANQALLDAVLLSQKLFDISRIHNGKTNV 527
Phatr2_45936             -----------TASVTLVGDACHPMSPFKGQGANQALLDALALVRSIYKHCKTAGSY--- 531
Thaps3_1961              NTWS-------KDRVVLLGDACHATLPYVGQGANQAIEDAIYLAVCLNR----------- 379
Thaps3_270370(ZEP1)      SPWA-------KGPVALCGDAAHPMMPNLGQGGCQATEDGYRLVEELAKVQH-------- 546
Phatr2_45845(ZEP1)       QGWS-------KGQVAICGDAAHPMMPNLGQGGCQATEDGYRLAEELATVRT-------- 496
Phatr2_56492(ZEP3)       KPWT-------DGPVALLGDGVHAMMPNLGQGGCQAIEDAFVIGQELGSATK-------- 401
Thaps3_261390(ZEP2)      RSWA-------DGNVVLIGDAVHPMMPNLGQGGCQAIEDAFVLSETLEACES-------- 388
Phatr2_56488(ZEP2)       RSWA-------NGNVVLIGDAVHAMMPNLGQGGCQAIEDAYVLTETLANTRT-------- 460
Thaps3_22671             IGVASGEDNDDVGRAVLLGDAAHPPVPYIGQGAMMAMEDAGTLALLLARYCPLDTTNSPT 563
Thaps3_20663             NGWVREFDKSA-KYAVLAGDAAHAMPPFLGQGANQALQDAYLLAEKVFEYNDQVEQYSPV 526
Phatr2_43425             QGWSREMTDSKGGSVVLCGDAAHALPPFLGQGSNQAIQDAYCLAKQLYAYNAEIEQG--- 337
                                         : **. *   *  ***.  *  *.  :   :

Phatr2_47925             --------------IAVSNTQREIVQRTASVVAASRQASVYWHEPQLVMPKDGQDTQKFA 417
Thaps3_6395              NEQQPTISLNESTPQALAEFENDMLQRCEVKVKKSADAAKFLHSDVA------------- 574
Phatr2_45936             --------DSNSLERAVKEFEVEMLVRSAVKVEASAEAARFLHTEIA------------- 570
Thaps3_1961              ---------HDNYSDAFADYYDKRFPRTKRIVQFAGIMHKLYHT---------------- 414
Thaps3_270370(ZEP1)      ---------SRDVPGALGRYSRVRVIRTAIIQGFAQLGSDLLVD--------------FD 583
Phatr2_45845(ZEP1)       ---------TKDIEGALQEYYRKRIPRTTIIQALAQLGSDLLVD--------------FD 533
Phatr2_56492(ZEP3)       ---------RSQIVDKLREYQQRRLIRSAAVQGLSRFASDIIIR-------------GFD 439
Thaps3_261390(ZEP2)      ---------TQKLEDALQDFYKKRIVRVSIVQFLSRLASDLIIN-------------AFD 426
Phatr2_56488(ZEP2)       ---------TEKLQDALQEYYRKRIVRVSIVQFLSKLASDLIIN-------------AFD 498
Thaps3_22671             VD-------FSLFKKAMHAYESLRVSRTKTILGSSVELGKTQQKRAE----S-----KLY 607
Thaps3_20663             VRGGE-STAEPNLKALLNEYEKRRWLPTTSITAKAAGLGYLETG-S-----------GFF 573
Phatr2_43425             --------RDANLNAMLKDYENTRWPSTFGIFWKSTFLGYLETG-GE----D-----GLY 379
                                         .                 :

Phatr2_47925             GVCSQDIPALLHALQKKNIKANSANDLDKSVQCTI---DELQLIGPPEVRRKAESELKEA 474
Thaps3_6395              -------------IQEGNITRGAAA-LD--------------AKG*-------------- 591
Phatr2_45936             -------------IQKGNVTRGAAS-RS--------------ILIKSNEEDTATTE*--- 598
Thaps3_1961              DS--------------WLVHKA----LDVLIGSIINGGAALKQLEREIINECPVKDYQQY 456
Thaps3_270370(ZEP1)      LM--------------MTI----------------------------------------- 588
Phatr2_45845(ZEP1)       KM--------------MTI----------------------------------------- 538
Phatr2_56492(ZEP3)       TP--------------AKIYRD----ENGK----------FQ------FENCNYAGIVTK 465
Thaps3_261390(ZEP2)      TP--------------WSPHDD----LGKS----------WK------S-------YLTF 445
Phatr2_56488(ZEP2)       TP--------------WSPHDN----LGKS----------WK------S-------YLTF 517
Thaps3_22671             NA-----------WREWSIKA-----------------QVWAYGTLPVMRPGAASDYMTK 639
Thaps3_20663             GN-----------FR*-------------------------------------------- 577
Phatr2_43425             AR-----------FRDVFFKT-----MGAV------GIAEWVLI--SAAKPKI*------ 408


Phatr2_47925             HFVDLARQAILSSEMNNLAFLRKLSWEYPNLIR-----------NV-DVDDMTCLQKAAK 522
Thaps3_6395              ------------------------------------------------------------ 591
Phatr2_45936             ------------------------------------------------------------ 598
Thaps3_1961              AFNRQ*------------------------------------------------------ 461
Thaps3_270370(ZEP1)      ---PLLGPFFLTMTQLSMPFILRYLYTPSF*----------------------------- 615
Phatr2_45845(ZEP1)       ---PLVGPFFLFMTQVSMPFVLRFLYTPEF*----------------------------- 565
Phatr2_56492(ZEP3)       ILQP-----ILPI-FFSVQFAFLYDG----------------WKNDKQIDFKAFLGFSVL 503
Thaps3_261390(ZEP2)      FWKVRGVAPFYGNECQHISYSLLFSTQQPS------------------------------ 475
Phatr2_56488(ZEP2)       FWKPILQFAIFPM-----QFAYLYSYYPTGNMGDLPAKLEAIWKEKHKTDAEAVFEQASK 572
Thaps3_22671             VEEVVLE*---------------------------------------------------- 646
Thaps3_20663             ------------------------------------------------------------ 577
Phatr2_43425             ------------------------------------------------------------ 408


Phatr2_47925             SGHIRIA---HWLITEAGCL--VDA-RLL---NDLSIKSYMKALLKMYM*-------    562
Thaps3_6395              ---------------------------------------------------------    591
Phatr2_45936             ---------------------------------------------------------    598
Thaps3_1961              ---------------------------------------------------------    461
Thaps3_270370(ZEP1)      ---------------------------------------------------------    615
Phatr2_45845(ZEP1)       ---------------------------------------------------------    565
Phatr2_56492(ZEP3)       GGLIVSL--VLFELAEAGLGIGLGAEALLGAEGLLDFGGISAAIQDFFLGGGAAGL*    557
Thaps3_261390(ZEP2)      ---------------------------------------------------------    475
Phatr2_56488(ZEP2)       EGFVMEHEASFFKKAEVE----LSPTALAATKEELS*--------------------    604
Thaps3_22671             ---------------------------------------------------------    646
Thaps3_20663             ---------------------------------------------------------    577
Phatr2_43425             ---------------------------------------------------------    408
```

3)

### VDE/VDL/VDR Percent Identity Matrix

| | | | | | | | | | |
|---|---|---|---|---|---|---|---|---|---|
| 1: Phatr2_56450_VDR_ | 100.00 | 52.25 | 17.27 | 17.45 | 14.40 | 17.05 | 16.82 | 16.67 | 13.54 |
| 2: Thaps3_270211_VDR_ | 52.25 | 100.00 | 16.24 | 17.33 | 16.25 | 16.05 | 18.10 | 16.75 | 11.46 |
| 3: Thaps3_7677_VDE_ | 17.27 | 16.24 | 100.00 | 56.04 | 26.69 | 23.60 | 21.62 | 22.31 | 18.89 |
| 4: Phatr2_44635_VDE_ | 17.45 | 17.33 | 56.04 | 100.00 | 26.49 | 24.01 | 21.93 | 21.65 | 15.56 |
| 5: Thaps3_22076_VDL1_ | 14.40 | 16.25 | 26.69 | 26.49 | 100.00 | 65.61 | 30.81 | 30.30 | 30.21 |
| 6: Phatr2_46155_VDL1_ | 17.05 | 16.05 | 23.60 | 24.01 | 65.61 | 100.00 | 28.57 | 28.30 | 28.42 |
| 7: Thaps3_11707 | 16.82 | 18.10 | 21.62 | 21.93 | 30.81 | 28.57 | 100.00 | 53.25 | 39.29 |
| 8: Phatr2_45846_VDL2_ | 16.67 | 16.75 | 22.31 | 21.65 | 30.30 | 28.30 | 53.25 | 100.00 | 54.46 |
| 9: Phatr2_bd_1281 | 13.54 | 11.46 | 18.89 | 15.56 | 30.21 | 28.42 | 39.29 | 54.46 | 100.00 |

### VDE/VDL/VDR Alignment

```
Phatr2_56450(VDR)        MKLHRKGRYRLLVTAVLLGTVCSFVPENLRSGSVRIPRKNANAGSVPGTHTVSKQPSASA  60
Thaps3_270211(VDR)       ------------MAMVLLIRTAV-----IASYSLTL----------TSAFSSSIRPTCRT  33
Thaps3_7677(VDE)         ------------------------------------------------------------  0
Phatr2_44635(VDE)        ------------------------------------------------------------  0
Thaps3_22076(VDL1)       ------------------------------------------------------------  0
Phatr2_46155(VDL1)       ------------------------------------------------------------  0
Thaps3_11707             ----------------------------------------------MSASSSS-------  7
Phatr2_45846(VDL2)       ----------------------------------------------MK------------  2
Phatr2_bd_1281           ------------------------------------------------------------  0

Phatr2_56450(VDR)        TRHKVSQTSIDANALPIKNDLIQGPVSANKSIGSITFLLPSSGADEIKTNFGSSSPVGNP  120
Thaps3_270211(VDR)       FRQS---------------TPHHATASNVDIIGTVALLVPSSSTE--LSKYGSKSPAPRP  76
Thaps3_7677(VDE)         ----------------------------------MK--------LFL-----SLVLAAAP  13
Phatr2_44635(VDE)        ----------------------------------MKFLGVTSLLCLW-----SVVNRENV  21
Thaps3_22076(VDL1)       ----------------------------------MRP-----STSAL-----TVVLGT--  14
Phatr2_46155(VDL1)       ----------------------------------MRFAWVVAAGVVL-----TTTTQA--  19
Thaps3_11707             -------------------------TTTTTNAGKRARSWPSSSASTS-----SMPTRS--  35
Phatr2_45846(VDL2)       --------------------------RATRKRTLAATLWIAMSSVTG-----SGPGRT--  29
Phatr2_bd_1281           ------------------------------------------------------------  0

Phatr2_56450(VDR)        SLDEAVRHLANKSQYFSDGRVETRIVYVPIEEQDESVWKDLLETDVLLAMGLQYEADLAF  180
Thaps3_270211(VDR)       SYQEAAEHLARKISHFSDGRIEATVVTPSTNQDDT--DDVCLTSNALIALGITDPAEVQY  134
Thaps3_7677(VDE)         VSSFAPSNPVVSR---THSSVHSQQHN-H-------------------VLEAHND-----  45
Phatr2_44635(VDE)        SEAFAPRHQSLSR---PSSRTTSAFSRAP-------------------ILSLRK------  53
Thaps3_22076(VDL1)       -IALV------SCSQ-LN-----------------------------------NNVS-AF  30
Phatr2_46155(VDL1)       -LVPL------DCTG-MGETRTSGI--RP-------------------IRGLESNMA-RY  49
Thaps3_11707             -ILILATFLSLTSSS-SSTSVEAAFVGSP-------------------AVGLRSHTAAST  74
Phatr2_45846(VDL2)       -AAFAP----------SGNNNNNGCHGLS-------------------RVALHTTELQAH  59
Phatr2_bd_1281           ------------------------------------------------------------  0

Phatr2_56450(VDR)        A--RKLFQQRH-DRDAEHRFRQCHFAIDCAQ-SFPTMVGPYDSENPSFRAKLLPWTKHAS  236
Thaps3_270211(VDR)       L--STTFRKRRTSHQETSSYNTCQFALDCGSNNYAPLVGPWDEANPSILAEIAPWTGVAS  192
Thaps3_7677(VDE)         -----------NMDDITFSLSARNINNE-------IVE-R--------IGKV--------  70
Phatr2_44635(VDE)        ------------------YDSDSEVEND-------LLS-K--------LNPFQNW--QTA  77
Thaps3_22076(VDL1)       STRSSSLTQRHKTCTITTS-SSSLYVNP-------N-NDD--------DNSNRSQHKPNP  73
Phatr2_46155(VDL1)       ATVRHGTDQ--TNHGITSS-SERQWPFP-------R-GG----------------SSPRA  82
Thaps3_11707             SKQQSSLYAQ-KKNNIDSSDNPLSYLFD-------LSSDP--------ETKRRLQKQTAT  118
Phatr2_45846(VDL2)       SKPPHQQSPQ-T-----RYTPPSAMNIQ-------F-PEP--------DESLHVWDHVRN  97
Phatr2_bd_1281           ------------------------------------------------------------  0

Phatr2_56450(VDR)        GKRLSQQMVALLQRGNSDDF----VFAIMLF---LNQFSGSSVDWVKHSIDATWEKGPLR  289
Thaps3_270211(VDR)       GKRLTEQMNGLFEKQTSDEF----ALAVMLF---FNRFSGAAIPWVQHSIDVTWEKGLVQ  245
Thaps3_7677(VDE)         T----TSALLALTLSFSAI---------------TSPISGPNGDVLSSIPSAN----AAD  107
Phatr2_44635(VDE)        L----QSTALALTIGVASW---------------TSL------PTIVPPAFAA----TTD  108
Thaps3_22076(VDL1)       F----LSAALTAAVTTSLFLSSLPS--A-------TFASTPASTTQKYDGFAEYAKENKM  120
Phatr2_46155(VDL1)       V----ARSVATFGLGFSIALASVFGVAA-------APVGADTTPAVKYDGFAEYAQDNQM  131
Thaps3_11707             ---------LFSTLGFSALFLTNPLIPHLPFSPSLSSANAEDELYAKYGGK-----GLDT  164
```

```
Phatr2_45846(VDL2)           --------IGKVCTGFAL---AGLLSALVSF---TSPVWAENELSAKYGG------GLDT  137
Phatr2_bd_1281               ------------------------------------------------------------  0


Phatr2_56450(VDR)            NAQEVVSMVSKCGDCVVKCV-QDDNCRECLEVLTALDTR--DQVASYRTIVSYESDLLKD  346
Thaps3_270211(VDR)           NAKEIFSMITKCGPCITKCL-NDENCSQCINALDKIDTR--DQVTSYRTVVSFESELLRD  302
Thaps3_7677(VDE)             GAKIGLCLVKKCRVPLAKCI-TNPNCLANVICINSCNGKEDETGCQINCGNVFENDVVGE  166
Phatr2_44635(VDE)            SKSIVSCLFQKCPLPLAKCI-ANPKCLANVVCINTCTGRPDEIECQIECGNLFENEVVGE  167
Thaps3_22076(VDL1)           EQSDVGCFINKCGDQTKQLF-SNPRGIKGVSCLGRCKGE---QSCATRCFAEFGSEDLDN  176
Phatr2_46155(VDL1)           EQSDVGCFINKCGDQTKALF-SNPRGIKGVSCLGRCKGE---QSCATRCFAEFGSESLNA  187
Thaps3_11707                 SLVDKDCLVNQCQVQAKACLQDDPDCRKGLTCTAKCLGD---NACITGCFARYGNENLDE  221
Phatr2_45846(VDL2)           SLVDQNCLVSACSLQTKACLQDDPSCRKGLTCTAKCLGD---NACITGCMARYGNANLDN  194
Phatr2_bd_1281               -------------------------------------------------MERYSNDKLNN  11
                                                                                 : .  :

Phatr2_56450(VDR)            FSFCILQKNNIFNCDASLPTLPNVQPVATWREQPLTEDIARSLLVGHLNDEA-APESSLR  405
Thaps3_270211(VDR)           FSLCILQKNNIFECSAEIPELPVVKPMSTWRGKDVTTDVARGIMIGHLEGAGGSLEGNLQ  362
Thaps3_7677(VDE)             FNKCAVTDMTCVPQKKDDGSYP---------------VPSKDVLVQSF--------DTKL  203
Phatr2_44635(VDE)            FNKCVLTDMKCVPKQGDDGSYP---------------VPAPEIVVPKF--------DTKF  204
Thaps3_22076(VDL1)           WLSCTIEDYECVKVPKNIDN----S----------AENVGYDTTVKKF--------DPST  214
Phatr2_46155(VDL1)           WLSCTIEENECVKVPKNVDN----S----------AEDIGYSTTLRSF--------DPQS  225
Thaps3_11707                 LLKCTIEDHECIKVAILEGGGDVLG----------REPKSPAPTVQGF--------DLAS  263
Phatr2_45846(VDL2)           LLKCTIEDHECIKVAILEGGADVFG----------QEPRAPAPTVTAF--------DPKS  236
Phatr2_bd_1281               LLKGTIEDHECTKVAILEGGIEVFG----------QELRASDSTVTAF--------DPKS  53
                                  : .                           .    :  :        .

Phatr2_56450(VDR)            TDISWKVACGANEAYDKFPSQNQLFYPAARGRD---------LW----------------  440
Thaps3_270211(VDR)           LGVSWKVACGANVAYDQFPSQNQLFYPSAKGKD---------LW----------------  397
Thaps3_7677(VDE)             WNGRWFITAGQNKLFDTFPCQVHFFTETAPGKF----------VGK--------------  239
Phatr2_44635(VDE)            FDGRLYISAGQNKLFDVFPCQVHFFTETEKGKF----------FGK--------------  240
Thaps3_22076(VDL1)           LVGKWYKTDGLNPNYDLFDCQSNTFDFSDDTK------------------------KELD  250
Phatr2_46155(VDL1)           LVGTWYKTDGLNPNYDLFDCQKNTFTP--TSD------------------------KELD  259
Thaps3_11707                 MEGTWYKVAGYNPNYDCYACQRNTFSSPEGGLSDSLQLPTGGILGSLSNAVGSIGADRLQ  323
Phatr2_45846(VDL2)           LQGSWFKVVGYNPNYDCYACQRNTFSAPDSSANGNRNN---LLWSVASGNTNPAVTNQLR  293
Phatr2_bd_1281               LQSSWFGKVAHNPNYDCYACRGIQSMDV--------------LIVFHCGETK----DRDR  95
                                      . *  :* : .:

Phatr2_56450(VDR)            YDPVFRVETL--DGR--------------------------NVWCKRH------YKVRPA  466
Thaps3_270211(VDR)           YDPVFRVETI--DGR--------------------------NVWCKRH------YKVRNG  423
Thaps3_7677(VDE)             --LNWRIE--EPDGE----------------------------FFTRD----AVQEFVQDP  264
Phatr2_44635(VDE)            --LNWRVE--QPDGN----------------------------FFTRD----ALQEFVQDP  265
Thaps3_22076(VDL1)           MGIFFRVPRPEEYGG--------------------------GFWENSLTEHMIVDAVSP-  283
Phatr2_46155(VDL1)           MGIFFRVQRPPESGG--------------------------GYWENALTEHMIVDVPVQP  293
Thaps3_11707                 VDVEFSMPRYLPDGSPQPPSGVRESFISSADSMEGSGLQSVGYNQYSTHETMVFDTVKSN  383
Phatr2_45846(VDL2)           MDVEFSMPHLLPDGSPPPPSNVRESILVSGEDGSVFGSKSIALNDYRTRETMVFDQVSTG  353
Phatr2_bd_1281               KGINFFLTHFFDADTVQ*------------------------------------------  112
                                 : :      .

Phatr2_56450(VDR)            DIPG-TFRF----------------------SVLDNGITSNEFWTIVGVADD-----LSW  498
Thaps3_270211(VDR)           ETPG-TFKF----------------------SVLDNGVTSNEFWTIVGAADD-----LSW  455
Thaps3_7677(VDE)             NNPA----------HL--INH------------DNEYLHYQDDWYIVDYAADDNKEGVPP  300
Phatr2_44635(VDE)            NQPG----------HL--INH------------DNEYLHYEDDWWVIDYEYDGNKDGVPP  301
Thaps3_22076(VDL1)           -------------------ELDNPTGRTMHTAGKMYGLKFTENWYILGE--SNGDNDIPP  322
Phatr2_46155(VDL1)           PTAGTQLVASANAATGDLNDELNPTGRTMHTAGKMYGLEFTENWYILGE--SDGKGSVPP  351
Thaps3_11707                 GVGE-AVKL-----ALGKRGEEKLYSRTAHSEGEMFGLKFWENWYIIGQ--NNP--GQDE  433
Phatr2_45846(VDL2)           N----NMVF-----H-KGTTQEVSYSRTAHSEGEMFGLKFWENWYIIGE--NDP--GQPE  399
Phatr2_bd_1281               ------------------------------------------------------------  112


Phatr2_56450(VDR)            IVFHYAGAASAVGQRYLGGLLCTADGSLPDESQRPEIWRVLRSAGIQPWDLYT-VNNDLT  557
Thaps3_270211(VDR)           VVFHYAGAAGAVGQRYLGGLLCTPTGELPPEEDLGHIYNFLRSAEIEPWELFV-VDNDDQ  514
Thaps3_7677(VDE)             FAFVYYRGENDAWIGYGGAVVYTRDSKLPE-SLLPRLREAAKKVNFDFDKDFDLTDNSCK  359
Phatr2_44635(VDE)            FAFVYYRGNNDAWEGYGGVVVYTRAAQLPE-SLLPRLRVAAEKIGFDFDKDFVITDNTCP  360
Thaps3_22076(VDL1)           FKLVAYKGHTL-QGNYEEAFVYAKESVLPK-EAVGAVREAAAKAGLDFDK-FTRIDNTCP  379
Phatr2_46155(VDL1)           FKLVAYKGHTL-QGNYEGAFVYAKEATVPE-AAKPAIREAATKAGLDFDA-FTRIDNTCS  408
Thaps3_11707                 FKFVYYNGKTR-QNTYDGAFIYSRSRTLSP-ASMEKVYKIAKDAGMNPDQ-FCKIQNSCF  490
Phatr2_45846(VDL2)           FKFVYYNGKTR-QNTYEGAFVYSRSKELAP-ESMAKVYSIAKEAGMKVDQ-FCRIRNGCF  456
Phatr2_bd_1281               ------------------------------------------------------------  112


Phatr2_56450(VDR)            SPG------------AQEAGPPPLDYFRREVLAKR---------------AASSPHP*--  587
```

```
Thaps3_270211(VDR)      SPG------------ALAAGAPPLDYFRKTASVIG---------------*---------  537
Thaps3_7677(VDE)        ALEKG-EEVVLR-EKFAGKMAIQTEK-------------QLQQQAVLARTAASNTVKGEV  404
Phatr2_44635(VDE)       TDLSGKEKQILR-EKFAGKVALQTEQ-------------QLQAQVTRLRGNAVNSIKAQK  406
Thaps3_22076(VDL1)      TTTKS------LNDASAGTG-TSTTDWVDLVVGEGGVIDWV---VPGWRGEYKN*-----  423
Phatr2_46155(VDL1)      V-GDS------LNDAQAGTG-TSTTDWINLVVGEGGVIDWI---SPGWRGEYKAKR*---  453
Thaps3_11707            DGEDDKQEMMMMNPQREGLG-SPSNPFRGILASTK-VSQFLGVESVAAETTYNEPKSTIS  548
Phatr2_45846(VDL2)      SDET------VVKAPSSGLG-SQSNPFRGILASTR-ISQLLGVEPVAARDTVRRNAPT--  506
Phatr2_bd_1281          ------------------------------------------------------------  112

Phatr2_56450(VDR)       ------------------------------------------------------------  587
Thaps3_270211(VDR)      ------------------------------------------------------------  537
Thaps3_7677(VDE)        TAVEKSLQK-------IEEKALAFEKELMKDVVSVEKEIVKEVEEVEKEIVQEEQKIFGG  457
Phatr2_44635(VDE)       LFFEQGLEGAQKAYDALEETEKQFERETSQQ*----------------------------  437
Thaps3_22076(VDL1)      ------------------------------------------------------------  423
Phatr2_46155(VDL1)      ------------------------------------------------------------  453
Thaps3_11707            SNFLQGSQATNRKADAVQERPWWKEMGDY-------------LEDP-----RRHFRLMDS  590
Phatr2_45846(VDL2)      ------SPTLQPAVGTIASRPWWYEIGDY-------------LENP-----HRHFQVMDS  542
Phatr2_bd_1281          ------------------------------------------------------------  112

Phatr2_56450(VDR)       -----------------       587
Thaps3_270211(VDR)      -----------------       537
Thaps3_7677(VDE)        IR*--------------       459
Phatr2_44635(VDE)       -----------------       437
Thaps3_22076(VDL1)      -----------------       423
Phatr2_46155(VDL1)      -----------------       453
Thaps3_11707            LRTDMDWPDYIKEKNW*       606
Phatr2_45846(VDL2)      LRLPMTWAEDVKN*---       555
Phatr2_bd_1281          -----------------       112
```
