## Appendix C - Gene Models and Targeting Predictions for "Overexpression of *Thalassiosira pseudonana* violaxanthin de-epoxidase-like 2 (VDL2) increases fucoxanthin while stoichiometrically reducing diadinoxanthin cycle pigment abundance"

### 1)Thaps3_268908 (PSY1)

#### Exon 1:

>Thaps3 chr_5:1332233-1330996
TACTTTTGAAGCCTTCTCCCTACTACTTCTCAATACGTTTTGCAATCTTTGCTGCATAAA
CATTAAAGATTGATTTTCGCCGGTTGATACACAACGATACGGAAGAAGTGAAAATCAATT
GCAAAACGGAACAGTACCTCTGCCGTAGAGGAACACAACATGAGGATATCATCCATTGTA
GCAGCCACCCTGATCACGTGCGATGTTGCTTCAGCATGGTCATCGTCTGCATTCACATCA
CCGTTATCTATCAAGACACAGCATCGGTCTCAGGCATCAACCAAAAGAAGAGACGGATCA
ATCTTGATGTCTGCTGTGTCACAAAAAGAAAGTAGTAGTAGTAGCAGCAGCGGCCAAGGA
CGCACCACAAAAGAGATTTCAAACGAATTAGCATTAGGTGTTACTCTCGATGGAGGTCCG
GTGATTGATTTTGCCTCCGTCAAAGATACCACATCTCGTGCAGAAATAGCACTTGCCGAC
TCACGAAAGAAGTACGAAGCCAGTGGTGCTACAATTAGCCCTAATCCGGGAGGAAGATTA
ATGGGTATCAACGATGAGGTTGTGGCAGAGGTTGGCTATGAGATTGGGGAGTTTGCCGAA
GAATATTTGGACAAGGAGTCTGGAGAGACTGTAGATGAATTGGTGCAAAAGGTTGCTCGA
TACCTTCGTTCAAAGTCAAATACGGATACGTTCCCTGAAATGGAAGAGGAGGAAGCACCG
TTTACTAATCAGGAGAAGATTAGGTTCAATCGTCTCTTATCCCGAGCCTACGAAGAATCA
GGGATTGTCACATCAGCATTTGCCAAAACCTTTTACCTCGGTACACAAGTTCTTCCGGAA
CCATCCATGAAGGCCATTTGGGCAATTTACGTTTGGTGCCGTAGAACAGATGAAATCGTC
GATGCCCCTCGTCCTGCGGCTCACGATCCTAATGCAGAAATGTTGACAGATTTGTCGGAG
TGGGAGATTCGTCTCGAGAGACTGTTTGATCGAGGGGAGGTTGTGGATGTGCTGGACTTG
CCACTGTTGGACTGTAAGGTCAAGTATCCTACGTTGCCCATTACACCATTTTCAGATATG
ATACGTGGTATGTTGATGGACATCCCTGGGCTGGGACAAGAGAGATACGATACTTGGGAT
GAGTTGCACCTATATTGTTATCGCGTGGCCGGGACTGTGGGGTTGATGTCCATGCCAGTG
TTTGGATGTGCGGAAGGTTACACCGATGAAGTTGCAAA

#### Exon 2:

>Thaps3 chr_5:1330907-1330265
GGAGCCTGCTCTTTCGCTTGGTGTTGCATTCCAAATTACCAATATTCTCCGTGATGTAGG
AGAGGATGCTGAGAAACGCGAACGTGTATACCTTCCCCAACGAGACATGGAGAGGTTTGG
AGTTACTGAACGACAAATTTTTGACAAAGTAGTGGATGAGAACTACATTAATCTCATGAA
GTTTGAGATTGCTCGTGCCCGAATGTACTACGCTCGTGCTCTGAGAGGTGTGCCCATGCT
TCGCCCAGAGTCACGTCTGCCAGTGCAACTCTCGTTGGATGCCTACGGGAAGATATTGGA
CAAAATTGAGGAGAATGGATATGATTCGCTGACAAAACGAGCATATGTTGGTAAGTGGGA
GAAGTTGGCAGGAATCCCAGCGTCTTGGTACCGTACTCTTGATATTGCAAAGGCAATGCC
ACTCCCTGGGGATTGGGAGCGTCCGTCGTTAGAAGAATATGAAAAAAGTTTGGAACAAAT
GTTGGAGAATAAATGGGTAGAACAAAGGGGGAGCTCAGCGACAAGTGATTATTTGCAAAG
CCTCAACAAATAAGTGCAACTCGGCAAACCCTCAACGAAAGCATTGCTGCATCGCGAGGC
AGCGTTGAATAAAACAATAAATGTAGACAATGATGACGCAGTT

#### Translated:

MRISSIVAATLITCDVASAWSSSAFTSPLSIKTQHRSQASTKRRDGSILMSAVSQKESSSSSSSGQG
RTTKEISNELALGVTLDGGPVIDFASVKDTTSRAEIALADSRKKYEASGATISPNPGGRL

MGINDEVVAEVGYEIGEFAEEYLDKESGETVDELVQKVARYLRSKSNTDTFPEMEEEEAP
FTNQEKIRFNRLLSRAYEESGIVTSAFAKTFYLGTQVLPEPSMKAIWAIYVWCRRTDEIV
DAPRPAAHDPNAEMLTDLSEWEIRLERLFDRGEVVDVLDLPLLDCKVKYPTLPITPFSDM
IRGMLMDIPGLGQERYDTWDELHLYCYRVAGTVGLMSMPVFGCAEGYTDEVAKEPALSLG
VAFQITNILRDVGEDAEKRERVYLPQRDMERFGVTERQIFDKVVDENYINLMKFEIARAR
MYYARALRGVPMLRPESRLPVQLSLDAYGKILDKIEENGYDSLTKRAYVGKWEKLAGIPA
SWYRTLDIAKAMPLPGDWERPSLEEYEKSLEQMLENKWVEQRGSSATSDYLQSLNK-

##### Targeting:

SignalP 3.0 - NN = Yes, Cleavage Site ASA-WS

SignalP - HMM = Yes, Cleavage Site ASA-WS, Signal Peptide Probability = 0.993

SignalP 4.1: Yes, Cleavage Site ASA-WS, D = 0.554 (D-cutoff = 0.450)

ChloroP: Yes, Score = 0.558

#### 2)Thaps3_263269 (PSY2)

No introns.

>Thaps3 chr_7:1643508-1642761
AAAAAAAGTCGGATTGATACAACCAGAAGCGAAGTACACCGACCCAACAACAACCAACAC
CCCAACACCTCGGTCTGATCTGCTCTGCACACTCCAATGATGCTCCTACTATTATCAATA
TCGGCGTTATGCAACATCTCACTTGCTCGAGCGTTTGGGAACCTCTCACCAACGAACCGC
ATCAACCACCACCTCGACCTCTCCTCCTTTGTCCACCTATCTCGAACTCGTCTATATAGC
AGCCCACATTCAGCCAACCCAACAATATCCAATGAAATTATACAACTCATGTCAAAACAC
GATCCCATTCTATTATTTGCATCGCGACTGTTACCGTATCAAACTGCGGTCGACGCGTCG
GCTTTATACGCATGGTGTCGAAGATTGGATGAAATCACAGATGATCCATCAGCTAATGTG
CACTCTATACAACAACAACTGATTGATTGGGAGGATCGATTCAAGAAATTGTGCACTGGA
CAGCCTGTGGATGAAATGGACGATGCATTGTATCAATGCTTACAACGAAACTCAAACTCT
CTAAATGAGCGACCTTTTCAAGACATGATTGTGGGTATGAAGAGTGATGCTGTTCCAACT
ATTCGTACAATAAGCAGTATGGAGGAGTTGGAAGAATATGCCTATCAAGTCGCTGGAACT
GTGGGATTAATGCTGCTTCCGGAATCGTGTGGAGAAGGCACGTCAGCCTGCCATTGCTCT
TGGAAAGGCCATTCAACTGATTAACATA

##### Translated:

MMLLLLSISALCNISLARAFGNLSPTNR
INHHLDLSSFVHLSRTRLYSSPHSANPTISNEIIQL MSKHDPILLFASRLLPYQTAVDAS
ALYAWCRRLDEITDDPSANVHSIQQQLIDWEDRFKKLCTGQPVDEMDDALYQCLQRNSNS
LNERPFQDMIVGMKSDAVPTIRTISSMEELEEYAYQVAGTVGLMLLPESCGEGTSACHCS
WKGHSTD-

##### Targeting:

SignalP 3.0 - NN: Yes, Cleavage Site ARA-FG

SignalP 3.0 – HMM: Yes, Cleavage Site ARA-FG, Signal Peptide Probability = 1.000

SignalP 4.1: Yes, Cleavage Site ARA-FG, D = 0.826 (D-cutoff = 0.450)

ChloroP: No, Score = 0.489

If the first methionine is omitted, the ChloroP score becomes 0.499.

#### 3)Thaps3_23291 (PDS1)

No introns.

>Thaps3 chr_6:1795938-1794225
AACGCAAAGCGTTCCTCTTCGGCGTCAGCCTCCTCCTTTCTTTCACTGTCAGTCAGTTGT
GTTGAACACTTCTGCTTCTTCATTCATCTCCTCATTACTACCACTGCTATTGCAACTGGC
TGCCGTATCTGACTCTCCTTCAATTCGTCACTCCCATAGCGCCACGTTCACCATCATTTA
TCACCATGATCATTACAAATTTCATCCTCTCCACCGTCCTAGCGACATCAATGGCCTTTC
AACCACACACACCCATCCTCTCCAAACCATCCTTCTCCAACCGTGTCCATCGCTCCCCCA
AAATCGGCTCTTCCAACCTCGTTATGAAGGACTTTCCGAAACCAAATGTCGAAGATACAG
ACAACTATCGCTACGCAGAGGCCATGTCCACTAGCTTCAAGACGTCTCTCCGAGTGACGA
ATGATTCACAGAAGAAGAAGGTGGCTATCATTGGAGGAGGATTATCAGGTCTGTCTTGTG
CCAAGTACCTCTCCGATGCCGGGCATGAACCCACCGTATACGAAGCACGTGATGTACTCG
GAGGAAAGGTGTCAGCGTGGCAAGATGAAGATGGAGACTGGATCGAAACAGGTCTTCACA
TCTTCTTCGGAGCATACCCCAACGTTATGAACATGTTCGCTGAGCTTGGCATCCACGATA
GGCTTCAGTGGAAGATTCACCAAATGATTTTCGCAATGCAGGAACTTCCCGGAGAGTTCA
CTACCTTTGATTTCATCCCTGGTATTCCAGCTCCGTTCAACTTTGGATTGGCCATTCTTA
TGAATCAAAAGATGTTGACGTTGGGTGAAAAAATTCAGACCGCTCCTCCTCTTCTTCCTA
TGCTTATTGAGGGACAGTCATTCATTGATGCTCAGGATGAGTTGAGTGTGACGCAGTTCA
TGAGGAAGTACGGTATGCCTGAGAGAATCAACGAGGAGGTGTTTATTGCGATGGCCAAGG
CGTTGGACTTTATTGATCCTGATAAGTTGAGTATGACTGTGGTGCTTACGGCTATGAACA
GGTTCTTGAATGAGAGTAATGGACTTCAGATGGCATTCTTGGATGGAAATCAGCCTGATA
GGTGGTGCACTCCCACGAAGGAGTATGTGGAAGCACGCGGAGGAAAGGTCAAATTGAACT
CTCCCATTAAGGAGATTGTGACCAACGACGATGGAACTATCAATCACCTTCTCCTTCGAT
CTGGCGAGAAGATTGTGGCCGATGAATACGTCTCTGCCATGCCCGTGGACATCGTCAAAC
GTATGCTTCCCACAACGTGGCAGACTATGCCCTACTTCCGTCAGCTTGACGAACTTGAGG
GCATCCCTGTTATCAACTTGCACATGTGGTTCGATCGTAAGTTGAAAGCAGTCGACCATC
TTTGCTTCAGTCGCTCCCCACTCCTTTCCGTCTACGCCGACATGTCCGTCACATGCAAGG
AGTACGAAGATCCCAACAAGTCCATGTTGGAATTGGTCTTTGCTCCCTGCTCTCCTATTG
CCGGAGGAAATGTCAACTGGATTGGAAAGTCAGATGAGGAAATCATTGATGCTACCATGG
GTGAGCTTGCTCGCCTTTTCCCTACCGAGATTGCGAATGATGATAAGTGGCCTGCTACGA
AGATGCAGGGACCTAATGGACAGGCAAAGCTTGAGAAGTATGCTGTTGTGAAGGTGCCAA
GGAGTGTGTATGCTGCCATTCCTGGTGAGTGAAA

##### Translated:

MIITNFILSTVLATSMAFQPHTPILSKPSFSNRVHRSPKIGSSNLVMKDFPKPNVEDTD
NYRYAEAMSTSFKTSLRVTNDSQKKKVAIIGGGLSGLSCAKYLSDAGHEPTVYEARDVLG
GKVSAWQDEDGDWIETGLHIFFGAYPNVMNMFAELGIHDRLQWKIHQMIFAMQELPGEFT

TFDFIPGIPAPFNFGLAILMNQKMLTLGEKIQTAPPLLPMLIEGQSFIDAQDELSVTQFM
RKYGMPERINEEVFIAMAKALDFIDPDKLSMTVVLTAMNRFLNESNGLQMAFLDGNQPDR
WCTPTKEYVEARGGKVKLNSPIKEIVTNDDGTINHLLLRSGEKIVADEYVSAMPVDIVKR
MLPTTWQTMPYFRQLDELEGIPVINLHMWFDRKLKAVDHLCFSRSPLLSVYADMSVTCKE
YEDPNKSMLELVFAPCSPIAGGNVNWIGKSDEEIIDATMGELARLFPTEIANDDKWPATK
MQGPNGQAKLEKYAVVKVPRSVYAAIPGE-

##### Targeting:

SignalP 3.0 - NN: Yes, Cleavage Site SMA-FQ

SignalP 3.0 - HMM: Yes, Cleavage Site SMA-FQ, Signal Peptide Probability = 0.864

SignalP 4.1: Yes, Cleavage Site SMA-FQ, D = 0.603 (D-cutoff = 0.450)

ChloroP: Yes, Score = 0.550

#### 4)Thaps3_1383 (PDS2)

##### Exon 1:

>Thaps3 chr_1:1373161-1372917
CGGGGAACAGACCTCAGACGTCCTTAAGTTGGATCCTTCTCTCATCCCTACTAACCAATC
ATGAAGTTTCTTCTACCGCTGCTACCAGCAGTCGCTGGGGCCTTCTCCATAACGCACCTC
TCACAACACCCATCGTTACGCATGCATCAATCTTTGTCCACATCGCTGTACTCTTCCTCC
TCCTCCACATCGCAACGTCCAAGACGTCCAACTCCTGATCGTATTCGCAATACACAAAAC
TTTAA

##### Exon 2:

>Thaps3 chr_1:1372807-1372145
AGAAGCGAAAGAGTTGAGTCAGAAATTCATAACAGACTTTCAACAACTACAAAAGGTTGG
TAGTGGCGAACCGAAGCGAGTTGCCATATTCGGAGGTGGTCTTTCTGGTCTTTCGTGTGC
AAAGTATCTCTCAGATGCTGGTCACATCCCAACCCTGTACGAGGCTCGTGGTGTTCTCGG
GGGCAAGGTCTCGGCATGGCAAGATGAAGACGGAGATACAGTCGAGACCGGACTACACAT
TTTCTTTGGTGCTTATCCAAACATCCACAATCTCTTTGATGGGCTGAAAATACAAGACAG
ATTGCAATGGGCTCCTCACAGAATGACATTTGCCATGCAAGAGCTTCCCGGGCAATTTAC
CACCTTTGAGTTCCCTGCTGGCGTTCCTGCTCCATTGAATATGGCTGCTGCGATTCTGGG
GAATACTGAAATGCTTACGTTGGAAGAAAAGATTAAAATGGTTCCAGGGCTATTACCAAT
GCTATTGGAAGGGCAATCTTTCATTGACGAGCAAGATGAGCTTTCTGTTTTGCAATTCAT
GCGGAAGTATGGTATGCCCGAACGTATCAATGAAGAGATATTTATTGCAATGGGAAAAGC
ACTGGACTTCATCGACCCTGATCTACTGTCCATGACGGTTGTTCTCACGGCAATGAATCG
TTT

##### Exon 3:

>Thaps3 chr_1:1372066-1371065
CATCAACGAAGCAGACGGAAGCCAAACAGCTTTCCTTGATGGGAATCCCCCAGAGCGACT
ATGTCAACCTATGAAAGAGTCCATCGAAAAGAAGGGAGGAGAAGTAGTTTGCAACAGTCC

TGTAGTTGAGATTCAACTGAACGAAGAGAGTAACGTCAAGTCTCTCAAACTTGCAAATGG
AACTGAAATCACAGCAGATTATTACGTGTCGGCAGTGCCTGTGGATGTCTTCAAACGTCT
CGTGCCCACGCAGTGGTCAACAATGCCTTACTTTCGTCAACTTGATGAACTTGAAGGAAT
ACCTGTCATCAACATTCAGATTTGGTTTGACCGAAAGCTCAACTCGGTGGATGGATTGTG
CTTTAGTCGGTCTCCACTGTTGAGTGTCTATGCGGATATGTCAACGTGTTGCGAAGAATA
TGCAAGTAACGATAAATCCATGTTGGAGTTGGTGTTTGCACCGTGTTCTCCTGAGGCGGG
ATCTCCATTGAATTGGATTGCGAAGCCAGACTCTGATATCATTGACGCAACAATGAAGGA
GTTGGAGCGCCTCTTTCCCTTGGAGATCGGTCCCGATGCTCCCGAGGAGAAACGCGCCAA
TGTTGTAAAGTCTACGGTGGTCCGTGTACCTCGAAGTGTGTATGCGGCTGTTCCTGGCAG
AAACAAATATAGACCTAGTCAGGAATCACCAATCGAAAACTTCATTATGGCTGGAGATTA
TGCAACACAGAAGTACCTTGGTAGCATGGAGGGGGCTGTACTCTCAGGGAAACTTGCAGC
TGAGGTCATTTGCGACAAGTTCATGGGCAGAGCGGAGAGGAAAGGGGTCAAAGAGGTACA
CTCATCGGTGCTCACGAAGCAAATCGAAGAGAGAACCCCAGCGGGTATTGCAATGGAAAA
AGGCAGAGTGTCGCCAACATCGTATGGAGGTGGTCAACAAGGTGGCTTTGAAAATCCCTA
AAAGTTTAGACTAATGCACTGTACAATCTGCTCAGGATGTTT

#### Translated:

MKFLLPLLPAVAGAFSITHLSQHPSLRMHQSLSTSLYSSS
SSTSQRPRRPTPDRIRNTQNFKEAKELSQKFITDFQQLQKVGSGEPKRVAIFGGGLSGLS
CAKYLSDAGHIPTLYEARGVLGGKVSAWQDEDGDTVETGLHIFFGAYPNIHNLFDGLKIQ
DRLQWAPHRMTFAMQELPGQFTTFEFPAGVPAPLNMAAAILGNTEMLTLEEKIKMVPGLL
PMLLEGQSFIDEQDELSVLQFMRKYGMPERINEEIFIAMGKALDFIDPDLLSMTVVLTAM
NRFINEADGSQTAFLDGNPPERLCQPMKESIEKKGGEVVCNSPVVEIQLNEESNVKSLKL
ANGTEITADYYVSAVPVDVFKRLVPTQWSTMPYFRQLDELEGIPVINIQIWFDRKLNSVD
GLCFSRSPLLSVYADMSTCCEEYASNDKSMLELVFAPCSPEAGSPLNWIAKPDSDIIDAT
MKELERLFPLEIGPDAPEEKRANVVKSTVVRVPRSVYAAVPGRNKYRPSQESPIENFIMA
GDYATQKYLGSMEGAVLSGKLAAEVICDKFMGRAERKGVKEVHSSVLTKQIEERTPAGIA
MEKGRVSPTSYGGGQQGGFENP-

#### Targeting:

SignalP 3.0 - NN: Yes, Cleavage Site AGA-FS

SignalP 3.0 - HMM: Yes, Cleavage Site AFS-IT, Signal Peptide Probability = 0.994

SignalP 4.1: Yes, Cleavage Site AFS-IT, D = 0.465

ChloroP: Yes, Score = 0.569

### 5)Thaps3_24832 (ZDS)

No introns.

>Thaps3 chr_14:437250-439255
CTCAACCAACGAAGAAGCCTTTTTTTCACTTGTCGTCAGCTCTCTCATTCTCTGCGACCC
AACCCATAGAGACTGCCGAAAACAATTCACATCCATCGCACCTCACATAGCCACTCCGAA
ACACCTGCCATGCGTCTCTCCACAGCCTTCCTCGTCGGATGTGTCCTCCCAGCAACACAC
TCGTTCCACCTTCCCTCCGCCAGCACCTCTCTCCGTCGTCCCGTCACCGTCGCCACCACA
TCATCCCTCTCCATGTCCGCCGCCACCTCTGAAGGAGAATTCACCTCCGAATCCGCCAAA
CAACAAATCGGAAACGACTCCTTCCTCAACGAAAACCTCATGGCGCGTGCCCAAAACGGA
CCTGGCAAAGTCAACGACGAAAAGCTCAAGATTGGAGTAGTCGGCGCTGGTTTGGCGGGT
ATGGTCGCCGCGATGGACTTGGCCGATGCAGGTCATGACGTGGAGATGTTTGAGTTGAGG
CCTTTTGTGGGGGGGAAGGTTTCGTCGTGGAAGGATAAGGAGGGGAATCATATTGAGATG
GGGTTGCATGTGTTCTTTGGGTGTTATTATAACTTGTTTGGGATTATGAAGAGAACGGGA
TCGTTTGATACCGAGTTGAGGATTAAAGAGCATATTCACACTTTTGTGAATGAGGGTGGG
ATTTTGGGTGCGTTGGACTTCAAATTTCCTATTGGGGCTCCTATTTCGGGACTTCAAGCC
TTTGCTCGTACGGAGCAGTTGGGATGGGATGACAAGTTCCACAATGCACTGAGGTTGGGT
ACTTCTCCTATTGTGAGGGCGCTGTTTGACTTTGATGGTGGTATGGATATGGTGCGAGAC
TTGGATGATATCACGTTTACTGAGTGGTTTACTCAGTTGGGAGGATCACGAGGAAGTTTG
GATAGGATGTGGGATCCCATAGCGTATGCGCTTGGTTTCATTGATTGCGATCACATCTCG
GCAAGGTGCATGCTGACAATCTTTATGCTTTTCGCCATCAGAACTGAAGCGAGTGTGTTG
AGAATGCTGGAAGGAAGTCCTCAGACGTGCTTGCACGATCCTATTCTCAAGTATTTGGGT
GATCGTGGGGTCAAGATCAATACCTCCATGGGATGCAGAGAGATCGTGCACGATGTGGAT
GAGAATGGCAAGCCTATTAGAGTGACTGGAATCAAGGTTGGACCCAAGGAGGAGTTGAAG
GAGTTTGATGCCGTGGTTTGTGCATTGGATGTTCCTGGAATCAAAAAGGTCTTGCCTCAA
TCATTCAGGGATCACTACCCAATGTTTGACAACATTTACAACCTCGACACTGTTCCTATT
GCTACCGTTCAAGTCCGATTCGATGGATGGGTGACTGAAATGAATGACGACGTTCGCATG
ATGGATATTTCTGGAGATCAATCCGACGGACGTGGCGGTGGAATTGACAACCTTCTCTAC
TCTGCCGATGCTGAGTTCTCATGCTTCGCTGATCTCGCCATCACTTCTCCCGGAGAGTAC
TACAAAGAAGGGGAGGGAAGTCTCATCCAAGCAGTGTTTGACGAACGTGCCTTTGATCGT
TCCAATGACCAAATTGTCCAAGATTGCATCAGTCAATTGAACTCACTGTTCCCTTCCAGT
AAGAAGCTCAATTGCACCTGGTCAAGTGTGGTCAAGTTGGGACAATCACTGTACAGAGAG
AAGCCAGGACAGGACAAGTTCCGTCCGAAGCAGGCTACTCCCATCTCCAATTTTTTCTTG
GCTGGAAGTTACACTTACCAAGATTACTTGGATTCCATGGAGGGTGCTACGCGTAGTGGT
TTGATGGTGGCGGATGAGATTATTGCAAGAGCTGACGGACCTAATGGATTGAAGGCACAG
ACTGCCAAAGCGATGGCCAATGGGAGTGGGAAGAAGGCTATTAGTGAGGAAAAGGTTCCA
TTCTTTGCATCAGCGTCGGCATAATGGGGGTACAGCAGGCGGAGGATAATGTTCTTAATG
ATATTTAGTTGAAATACTAAGGCAGG

#### Translated:

MRLSTAFLVGCVLPATH
SFHLPSASTSLRRPVTVATTSSLSMSAATSEGEFTSESAKQQIGNDSFLNENLMARAQNG
PGKVNDEKLKIGVVGAGLAGMVAAMDLADAGHDVEMFELRPFVGGKVSSWKDKEGNHIEM
GLHVFFGCYYNLFGIMKRTGSFDTELRIKEHIHTFVNEGGILGALDFKFPIGAPISGLQA
FARTEQLGWDDKFHNALRLGTSPIVRALFDFDGGMDMVRDLDDITFTEWFTQLGGSRGSL
DRMWDPIAYALGFIDCDHISARCMLTIFMLFAIRTEASVLRMLEGSPQTCLHDPILKYLG
DRGVKINTSMGCREIVHDVDENGKPIRVTGIKVGPKEELKEFDAVVCALDVPGIKKVLPQ
SFRDHYPMFDNIYNLDTVPIATVQVRFDGWVTEMNDDVRMMDISGDQSDGRGGGIDNLLY

SADAEFSCFADLAITSPGEYYKEGEGSLIQAVFDERAFDRSNDQIVQDCISQLNSLFPSS
KKLNCTWSSVVKLGQSLYREKPGQDKFRPKQATPISNFFLAGSYTYQDYLDSMEGATRSG
LMVADEIIARADGPNGLKAQTAKAMANGSGKKAISEEKVPFFASASA-

##### Targeting:

SignalP 3.0 - NN: Yes, Cleavage Site THS-FH

SignalP 3.0 - HMM: Yes, Cleavage Site THS-FH, Signal Peptide Probability = 0.999

SignalP 4.1: Yes, Cleavage Site THS-FH, D = 0.543 (D-cutoff = 0.450)

ChloroP: Yes, Score = 0.572

#### 6) Thaps3_270357 (LCYB)

No introns.

>Thaps3 chr_2:1232356-1230385
CGGGTTGACACGTCATCAACCTTTCTATCACTCGTCCTCGCTCGGCATTCTTCCTCATGG
TGGCCGTCAATCCTCTCCACGCTGTGGCATTAGCAGTGCTTCCATCGGCGGCATATGCCT
TTGTATCTCCTACACCAGTCGTCCTCTCCCCGTCGTCGGCTAGAGTGACACCCTTGTCTG
GTGCACCTTACTGCACCTCATATCGCGTTCGTCATTCCCACTCCTTCCCGACAACGTCCA
TCCAATCTCGTACCACCTCCTCTACAATCTTTGCATCCGCGTCCAACCCAACTCTCACTC
CAACTTCTCAAGATACATGCGATGTTCTCGTCCTCGGAAGCGGACCTGCAGCACGATCCA
TCGCCACCCTCCTCTCCTCCAAAGCAAACGACAAAGCATACGACGTACTTCTAGCAGACT
CAAATTACGATCGCAGATGGGCTCCTAACTACGGTGTTTGGCAAGATGAATGGCAATCTA
TTTGCAAACTATATGAATCATTCAATCAGCCCATTGGTAATGAAGTTATTGATCGGTTGT
GGATGTCGACGGATTGTTTCTTCGGTGGCTCTTTTGACATCGCTGCGGAGCAACGGATGA
GATTGGATAGGCCATATTGCAGGATTGAGAGGGATGTGCTGAGAAGAGTGTTGAGTCCAG
CGAGTGAAGTGAAGACACAGAACGATGGAGAGGGGACAGCAAACTATCGTGTTACACGTG
CCAATCACATGTCAAAGTGCACGTCTGTGAACATTTATTCTCCCTCTGGATCAATGGTAC
ACGATGAATCTGGAACCAGTGTTTTGATTCAATCAAAGGATGGTGATATTTCTCAAGTAC
GTGCAAAGCTAGTTGTGGATTGTACGGGACATGAATCCAAAATCGTGTTGAAGGATGATC
GTATGAAATCCATTCCTCCAGGCTTTCAGATTGCCTACGGTATTCTAGCAGAAGTTGACG
AGACGAGCATACCAAACAATGATTTCTGCGGTCCGTACTTCAAAGAGGCAATGACTTTGT
TTGATTACAGAACCGATCACTTTCCGGAAGGTTCTAATGAGTTGGCTAAAGCAGAAAAGG
CACCGACGTTCATGTACGCCATGCCACTCAATGGGAATCGTATCTTTTTCGAAGAGACAT
CGTTAGTGGCTAGACCAGCATTATCATTCCAAGAATGCAAAGATCGATGCATGACACGTC
TCGAGCACTTGGGGATTACTGTTACAAAGGTTGAGGAAGAGGAGTTTTGCTACATTCCAA
TGGGAGGACCGTTGCCAGCCAAAGACCAAAGAGTTATTGGATTTGGAGGTGCGGCTGCGA
TGGTTCATCCTAGTACTGGGTACCACCTGTGTCGAGCAATGATGGGAGCAGGAGAAGTAG
CCAAAGTGATTCGTGAAGAGTTGGAAGAGAAGAACTGGAACCCAGATAGGGCAGCTGCAC
GTGCTTACAATGCAATTTGGTCACCAACGACCATTGCCCAACGAAACTTTGCAGTCTTTG

GAGGTGAATTCTTGATGAAGCAAAACGTAGTAGGGCTGCGAGGCTTCTTTGATGGCTTCT
TCAAGCTTCCTCTTGGCTTGTGGGGAGGGTTCCTCGCTGGATGGCCTGGACTGCCAAACA
ATGAGAATCACGAGACGTGGTGGGCACGGTTAGTGTTTGGGTTGACGTTTGTTTCAAAGC
TACCCGTGTCTGTGGCGGTAGATATGCTTGGATCGATAGCAACATACTCCATTTCAGAGG
GAGTACCACTGCCTCAGTCGGTGACACCGTTATTGGGATTGCCTGATGGATATGAGTACA
GGGAAAAGAAGAGTTCTATAGGTGATGTTGCAGCGAAGCATGAGGCAAGGAGAATGATTA
TGGAGTCGACTGTTGAGGAAGTAGTTCCTGTAGACTTTGAAGAAAAGACAGCGGCGTGAT
TGGATTGATAACTCTTATTTGGAATGTAGTGATAGATACACAACTACAGATT

#### Translated:

MVAVNPLHAVALAVLPSAAYAFVSPTPVVLSPSSARVTPLSG
APYCTSYRVRHSHSFPTTSIQSRTTSSTIFASASNPTLTPTSQDTCDVLVLGSGPAARSI
ATLLSSKANDKAYDVLLADSNYDRRWAPNYGVWQDEWQSICKLYESFNQPIGNEVIDRLW
MSTDCFFGGSFDIAAEQRMRLDRPYCRIERDVLRRVLSPASEVKTQNDGEGTANYRVTRA
NHMSKCTSVNIYSPSGSMVHDESGTSVLIQSKDGDISQVRAKLVVDCTGHESKIVLKDDR
MKSIPPGFQIAYGILAEVDETSIPNNDFCGPYFKEAMTLFDYRTDHFPEGSNELAKAEKA
PTFMYAMPLNGNRIFFEETSLVARPALSFQECKDRCMTRLEHLGITVTKVEEEEFCYIPM
GGPLPAKDQRVIGFGGAAAMVHPSTGYHLCRAMMGAGEVAKVIREELEEKNWNPDRAAAR
AYNAIWSPTTIAQRNFAVFGGEFLMKQNVVGLRGFFDGFFKLPLGLWGGFLAGWPGLPNN
ENHETWWARLVFGLTFVSKLPVSVAVDMLGSIATYSISEGVPLPQSVTPLLGLPDGYEYR
EKKSSIGDVAAKHEARRMIMESTVEEVVPVDFEEKTAA-

#### Targeting:

SignalP 3.0 - NN: Yes, Cleavage Site AYA-FV

SignalP 3.0 - HMM: Yes, Cleavage Site AYA-FV, Signal Peptide Probability = 0.996

SignalP 4.1: Yes, Cleavage Site AYA-FV, D = 0.670 (D-cutoff = 0.450)

ChloroP: Yes, Score = 0.570

### 7) Thaps3_263437 (BCH)

No introns.

>Thaps3 chr_8:927506-925618
CGTCATCAGTCACCTTTCTTTCTCACCTCCATCATCCACCCTGCCATCCATCATCCACCC
TCTACCACCCTATCATCTACATACATTCACCCATGCATCAGCTATCCAGACTGCTTTCCA
AACTCTTCCCAGTCATCGCTACCTACATCATTGCGAAGTATGCTCTCCCTCACATCAATA
CGATCCTTCACTGTTCAAATACCTTGAGCAACTTTGTGAGAATAACATACACCGTCCTCT
TTGCAATAGCGATGGAGTACATTTCAAGATATAGTCACTGCTACTTGTGGCATGGAAAGT
TCCTTTGGTGGATTAATGGCTCACATCATCATCAGTATCCAGCTGTTGGAAGCACACCTG
TGTATGGACACAACAATCCGTACGTCTCTCCAGCCATTGAGTTGAACGATGCATTTGCCG
TTTTCTTTGCAACGATTGCCACATTGGCAATGTGGATAGGGTCAGAGCCTCCATCAACCT
TGACAAAAGATTGTTCTATTGGTATTGGATTGGGAGTGACTCTCTATGGCCTTTCATACT

TTGTCGGTCACGACATTGTTGCTCACGAACGTTTGGGCAAGGGAGTAGCAAACGCTTTGA
GACGAGCATTTCCGTATATGGAGCAATGTGCTTCTGTTCATATCCGGTATCATCACAAGT
TGACGAAACGTAGTAATGATTCTGATCCATATGGAGCGCCTTACGGCTTTTGGTTGGGTC
CGTCGGAAGTAGAGTGTTTGAACAGAGGTCAATGGTATGCACCGATGCCTATGTCGTTGA
AAGCTATTTCTTGGATAGCAACATTGATCTTCTTCGCATCGACAATTCACAGCTCGTTGT
CTCCGGCAGCTCAAGCAATTGTGCTCCTCGGATGTGTAGGATGGTGTGGCTCCGGCACAT
TGTCAGACAATACAACAACATCTCGTCGTATCGGCAAGCTGCTCTCATTGAACTGGCAAT
CAACTCCATCGAGACTGATGCCTCACGGACTTTCTGGGCTCATTTCAGTTGGCATTGGTT
CATACCTCATCTTTGGCCACAGTCTCGTAGGCGATCTGAAACCATACACAATGCAACAAC
CACCATATCTCATCATTCTCTATGCAACAGCCACATCATGGAATGCTCTCGGAGGATATA
TGATTGTCAATACTGCTCCTCCAAATACAAGAATGCTGTTCAGGAGATGTGCAATACTTC
AGGTTTGCTTGTCGTACTTCATTGTTAGGTTCCTTCCGCACTCGTCGGTGCTGCTGATAC
GTCTCGAGAGCAACACAATTACAACATCGCTGAGATGTCTTGACTTAATCGTTACGATAT
CAGCAGTGGTGTGTACTCTATCGTTCTTTGATGCAGTCGTTGACATGTCAAAGCAGAGCG
TTGTCCTCGGACAATCAATCGCATTCGGTATCATCGGGATATTGCTCCTTTCCGTGTATC
CAATCCAACTGAGTTTGCAAGGAGAAGAGTGGTGGAGTTGTATACAAAACAGATATCCAA
TGCAAGCCAGTGGAATGATTGCTTATATTTACGTTCCAGCTACAGTGACGTTTAGTTTGT
TTCTCTTCGGAGCAACGCTGTATCAAAGAAAGATAATGTCTGCATCAGAGTATGGGATTA
TATCATTGATGGTGATACTTGTATGCCTATTGGCAACTGTGCTCAGCCAAGAGATACATA
TCCCAGACGTTTCGACGCAGAGAATCTATTTACCATGCGAAGATCCAGCAATGGATTCGT
TGGAAGAGAAAGTACTTGAAGCGTTGGACTTTTCTCGTTATGCACGTTCCATCTTGACCA
CAGTCTTGGGTATTAAATTTGAGTCACCCGCGTAGCATGATGGGAATCTCCAATTTAACT
AATTGCACATTAATCATTGTTGTATTATC

#### Translated:

MHQLSRLLSKLFPVIATYIIAKYALPHINT
ILHCSNTLSNFVRITYTVLFAIAMEYISRYSHCYLWHGKFLWWINGSHHHQYPAVGSTPV
YGHNNPYVSPAIELNDAFAVFFATIATLAMWIGSEPPSTLTKDCSIGIGLGVTLYGLSYF
VGHDIVAHERLGKGVANALRRAFPYMEQCASVHIRYHHKLTKRSNDSDPYGAPYGFWLGP
SEVECLNRGQWYAPMPMSLKAISWIATLIFFASTIHSSLSPAAQAIVLLGCVGWCGSGTL
SDNTTTSRRIGKLLSLNWQSTPSRLMPHGLSGLISVGIGSYLIFGHSLVGDLKPYTMQQP
PYLIILYATATSWNALGGYMIVNTAPPNTRMLFRRCAILQVCLSYFIVRFLPHSSVLLIR
LESNTITTSLRCLDLIVTISAVVCTLSFFDAVVDMSKQSVVLGQSIAFGIIGILLLSVYP
IQLSLQGEEWWSCIQNRYPMQASGMIAYIYVPATVTFSLFLFGATLYQRKIMSASEYGII
SLMVILVCLLATVLSQEIHIPDVSTQRIYLPCEDPAMDSLEEKVLEALDFSRYARSILTT
VLGIKFESPA-

The underlined portion aligns with BCHs from plants and green algae **(Appendix D)**.

#### Targeting:

SignalP 3.0 - NN: Yes, Cleavage Site VIA-TY

SignalP 3.0 - HMM: No, Signal Peptide Probability = 0.299

SignalP 4.1: No, D = 0.285 (D-cutoff = 0.500)

ChloroP: No, Score = 0.458

#### 8) Thaps3_9541 (LTL1)

No introns.

>Thaps3 chr_13:695539-693528
CTCCAACTTCGCATCGGTCAACACAACACTGCAAGAGCGAAGGAAAAGCCAAACCCTAAC
ATGAAGTTCACAACAGCTCTCGCCGTCCTCTGCTGGACGTCGGTGACAAATGCCTTCGTT
CCATCGTCCTTCACCTCTCCAGCGTTGAAGAATGAGCAGCAGCAAGTACGTGCATCATCT
CCACTGTACGCACTTGATACCAAAGAAAAGGAAGAAACCACCACGGCCACCTCTGCTTCC
TCTACCGACACATCTTCCACACCAGCAGCAGCAGCCACTGAAGAATCCGAAGGACTCCCA
TGGTGGTGGGAATACATCTGGAAGCTCCCCGTAATGCAACCAGCCGAACCAGGCACCGAC
ATCATCTTCGCCGACAGTGCTCGCGTCCTTCGCACGAACATTGAACAAATCTACGGAGGA
TTCCCCTCCCTCGATCAATGTCCATTGGCCGAAGGAGAAATTACCGATATTGCCGATGGA
ACAATGTTCATCGGTTTGCAGAGGTATCAACAACAGTATGGAAGTCCGTACAAACTGTGC
TTTGGTCCAAAGAGCTTCTTGGTGATTTCGGATCCAGTTCAAGCCAAACACGTTCTACGC
GATGCCAACACTCTCTACGATAAAGGAATCTTGGCTGAGATTCTTAAACCGATCATGGGG
AAGGGGTTGATTCCAGCAGACCCAGAAACGTGGTCGGTACGACGTAGGGCGATTGTTCCT
GCCTTTCACAAGGCGTGGTTGAATCATATGGTGGGATTGTTTGGTTATTGCAATGAAGGA
TTGATTGCTTCGTTGGAGGAGGCGGCGAAGAAAAATGATGCTCCTAACGGACAACAGGGT
GGAAAGATTGAGATGGAAGAAAAGTTTTGCAGTGTGGCACTGGACATCATTGGGTTGTCG
GTGTTCAACTATGAATTTGGATCAGTGAGCGAAGAATCACCAGTGATCAAGGCAGTGTAC
TCTGCATTGGTGGAGGCAGAACATCGTAGCATGACTCCCGCTCCTTACTGGGATTTGCCA
TTTGCCAACGAGGTGGTACCACGTCTACGCAAGTTCAATAGCGATCTCAAGGTCTTGGAT
GATGTGTTGACTGATTTGATTGATCGTGCCAAGAACTCACGTCAAGTGGAGGACATTGAA
GAGTTGGAGAAGCGTGATTATGCCAATGTGAAGGATCCATCGTTGTTGCGATTCTTGGTG
GATATGAGAGGTGCTGATATTGATAATAAGCAATTGAGGGATGATTTGATGACGATGCTT
ATTGCAGGGCACGAGACGACTGCGGCTGTGTTGACTTGGGCGTTGTTTGAGCTAACAAAG
CATCCGGAACAGATGGCCAAGGTTCGTGCCGAGATTGATTCTGTGTTGGGAGATCGTACA
CCGACATATGACGATATCAAAGAGATGCAGTATTTGAGGTTGGTGGTTGCAGAGACTTTG
AGGTTGTATCCTGAGCCTCCGCTGTTGATTCGTCGTTGTAGAACCGAAAACAAGTTGCCC
AAGGGAGGTGGAAGGGAGGCTACTGTGATTCGTGGTATGGATATCTTTCTATCATTGTAC
AATCTTCACCATGATGAGAGGTTCTGGCCGGAGCCTAATGAGTTCAAGCCCGAGAGATGG
GAGAGCAAGTATATCAATCCTGAGGTACCAGAATGGGCTGGATATGATCCTGCAAAGTGG
ATAAATACCAACTTGTACCCCAACGAGGTCGCGTCCGACTTTGCCTACTTGCCATTCGGA
GGAGGAGCACGAAAGTGTGTCGGAGACGAGTTCGCCACGTTGGAGGCAACTGTGACTTTG
GCAATGTTACTCCGTCGCTTTGAGTTCGAGTTTGACTCTGCCAAGCTTGCTGCTTCGAAG
ATTGACATTATGGATCATCCAGAGGATTTGGAGCATGCGGTTGGTATGAGGACCGGAGCT
ACTATTCATACTAGGAAGGGATTGCATATGGTGATTAGGAAGCGTGAGTTGTAGTTCATT
AGAGCCTAGTCTAAGCAACAACTATGGTCATT

##### Translated:

MKFTTALAVLCWTSVTNAFVPSSFTSPALKNEQQQVRASS
PLYALDTKEKEETTTATSASSTDTSSTPAAAATEESEGLPWWWEYIWKLPVMQPAEPGTD
IIFADSARVLRTNIEQIYGGFPSLDQCPLAEGEITDIADGTMFIGLQRYQQQYGSPYKLC

FGPKSFLVISDPVQAKHVLRDANTLYDKGILAEILKPIMGKGLIPADPETWSVRRRAIVP
AFHKAWLNHMVGLFGYCNEGLIASLEEAAKKNDAPNGQQGGKIEMEEKFCSVALDIIGLS
VFNYEFGSVSEESPVIKAVYSALVEAEHRSMTPAPYWDLPFANEVVPRLRKFNSDLKVLD
DVLTDLIDRAKNSRQVEDIEELEKRDYANVKDPSLLRFLVDMRGADIDNKQLRDDLMTML
IAGHETTAAVLTWALFELTKHPEQMAKVRAEIDSVLGDRTPTYDDIKEMQYLRLVVAETL
RLYPEPPLLIRRCRTENKLPKGGGREATVIRGMDIFLSLYNLHHDERFWPEPNEFKPERW
ESKYINPEVPEWAGYDPAKWINTNLYPNEVASDFAYLPFGGGARKCVGDEFATLEATVTL
AMLLRRFEFEFDSAKLAASKIDIMDHPEDLEHAVGMRTGATIHTRKGLHMVIRKREL-

##### Targeting:

SignalP 3.0 - NN: Yes, Cleavage Site TNA-FV

SignalP 3.0 - HMM: Yes, Cleavage Site TNA-FV, Signal Peptide Probability = 0.999

SignalP 4.1: Yes, Cleavage Site TNA-FV, D = 0.797 (D-cutoff = 0.450)

ChloroP: Yes, Score = 0.548

#### 9) Thaps3 270336 (LTL2)

##### Exon 1:

>Thaps3 chr_9:683521-684549
CGCCCTCCCCTCAACCACTTCCACGCTTTTTGGCTCTGGTTTAGCTGCTGATAAAAGTTG
AGAGTAGTCCAACAATACAGCAATACGACTCTCTTCGTTGCGTAGTAGTAGACTGTGTAT
CATCGGTGGAGGAAAGCTGAGTTGATTGGATAGAATTGGGCTGATACAACGGATAGAAAG
AACATCAAGATGTGCACCAAACTATCCAGCCGTCGGACGCTGTTGGCGTTGTACTTCGCC
TTTACTGGATGCACTGCCTTTCAACTACCATCGGCCACACCATCACGTGCGTCAATAACA
AAAGCATACAGTACCCACCTTGATAAAGAGATCAAAAGCAAAACGCCTCTCGTCAACCCA
AGCAAGATCTACACACAAGCAGACATAGATACTCTCGATCTCTCCTCGTACGAAAACGAA
CTCCTAGCAGCATGGGATACAGACTCATCCCTTCAACGTGGATTTGACTGGGAGATTGAA
AAGCTACGACGAAATTTTGCAGGCCTGCGCCAAAGAGAAGATGGACAGTGGGTTCGTAAA
CCAAGTCTATTCGATTTCCTCGTCACCAACACACCATCGAACGTAGTTGGTGTGAGCAAT
ACTGGTGAACGATATGAAAGTCCTCCAAAACCGGTGAATATGCTTGATGTAGGATTGCTG
ATCACCAAGAATCTATTGAACACTCTTGGATTTGGGCCGAGTCTCGGTATGGCAGCCGTG
CCGGATGCAGTGATTCAAAAGTATGAAGGAAGTTTCTTCTCCTTCATCAAGGGTGTGCTC
GGTGGTGATCTTCAAACACTCGCAGGAGGACCACTATTCCTACTCCTTGCAAAGTATTAT
CAGGACTATGGACCCATCTTTAATTTGAGCTTTGGCCCGAAGAGTTTCTTGGTCATCTCC
GATCCTGTTATGGCGAGGCATATCTTGAGGGATAGTAGTCCAGAGCAGTATTGCAAGGGA
ATGTTGGCGGAGATTTTGGAGCCAATCATGGGCGATGGTCTTATCCCTGCTGATCCAAAA
ATTTGGAAG

##### Exon 2:

>Thaps3 chr_9:684641-685198
GTACGACGACGAGCAGTCGTCCCTGGCTTCCACAAAAAGTGGCTCAACAATATGGTGACT

CTCTTTGGTGACTGTGGTGAACGTCTCGTTAACGATCTCGATGCACGGGCTACTGCTAAG
ACTCCAGTGGATATGGAAGAACGATTCTGCTCCGTAACGCTGGATATCATTGGAAAAGCA
GTCTTCAACTATGACTTTGGTTCAGTTACGAAGGAGTCTCCAATTGTCAAAGCAGTGTAC
CGCGTGCTTCGTGAGGCAGAACATCGATCATCATCTTTCATTCCGTACTGGGATTTGCCC
TATGCTGATAAGTGGATGGGAGGTCAAGTTGAATTCCGAAAAGATATGGGCATGTTGGAT
GATATCTTAACAAAACTAATCAATCGCGCTATTGAGACTAGGGACGAGGCAAGCGTAGAA
GAGTTGGAGGATAGAGATGTTGGAGATGATCCAAGTTTGTTGAGGTTCTTGGCTGATATG
AGAGGAGAGGATTTGACGAGTAAGGTGTTGAGGGATGATTTGATGACAATGCTTATTGCA
GGACACGAGACAACGGCT

#### Exon 3:

>Thaps3 chr_9:685302-685832
GCAATGCTTACTTGGACAGTGTTTGGACTTGTGAGCAATGATTCTGGTTTGATGAAGGAG
ATTCAGGCCGAAGTACGAACAGTCATGGGTGACAAATTGCGTCCAGATTACGATGACATT
GCCAAAATGAAGAAGATGAGATATGCTTTGATAGAAGCACTTCGTTTGTATCCAGAACCA
CCTGTTCTCATTCGTCGGGCAAGGTCTGAGGACAACCTTCCAGCGGGTGGGTCTGGTTTG
TCGGGTGGTGTCAAAGTATTGCGAGGAACAGACATCTTCATTTCTACATGGAATCTTCAT
CGTGCTCCAGAGTATTGGGAGAATCCGGAGAAGTATGATCCCACGCGATGGGAACGACGA
TTCAAAAACCCCGGAGTGAAGGGCTGGAATGGATACGACCCAGAGAAACAATCAGAGTCG
TCGCTGTATCCGAATGAGATCACTGCGGACTATGCATTCCTTCCGTTTGGTGCAGGGAAG
CGAAAGTGCATTGGGGATCAATTTGCAATGCTCGAAGCATCAGTCACTCTG

#### Exon 4:

>Thaps3 chr_9:685927-686236
GCCATGATCATAAACAAGTTTGACTTTACATTAGTTGGCAGTCCAAAAGATGTCGGCATG
AAAACTGGGGCAACCATTCACACCATGAATGGACTCAACTTGGTGGTGAGTCGTCGGTCG
GAAGATAATCCGATTCCGGAGACCAATGATTACTGGATACAGCAGCATTTGTCGAGAGGT
CTCAATGTCAATGGACGACCATATTCAACCAATGAAGATGCTGCCTGGACGGCATCTTCT
CGAGATAAGAATGAGGGAGTTGTCTCTCGGTTAGTTAATTAAAGTTATCTAGGATATAAG
AGGATTCTGT

#### Translate:

MCTKLSSRRTLLALYFAFTGCTAFQLPSATPSRASITKAYSTHLDKEIKSKTPLVNP
SKIYTQADIDTLDLSSYENELLAAWDTDSSLQRGFDWEIEKLRRNFAGLRQREDGQWVRK
PSLFDFLVTNTPSNVVGVSNTGERYESPPKPVNMLDVGLLITKNLLNTLGFGPSLGMAAV
PDAVIQKYEGSFFSFIKGVLGGDLQTLAGGPLFLLLAKYYQDYGPIFNLSFGPKSFLVIS
DPVMARHILRDSSPEQYCKGMLAEILEPIMGDGLIPADPKIWKVRRRAVVPGFHKKWLNN
MVTLFGDCGERLVNDLDARATAKTPVDMEERFCSVTLDIIGKAVFNYDFGSVTKESPIVK
AVYRVLREAEHRSSSFIPYWDLPYADKWMGGQVEFRKDMGMLDDILTKLINRAIETRDEA
SVEELEDRDVGDDPSLLRFLADMRGEDLTSKVLRDDLMTMLIAGHETTAAMLTWTVFGLV
SNDSGLMKEIQAEVRTVMGDKLRPDYDDIAKMKKMRYALIEALRLYPEPPVLIRRARSED
NLPAGGSGLSGGVKVLRGTDIFISTWNLHRAPEYWENPEKYDPTRWERRFKNPGVKGWNG
YDPEKQSESSLYPNEITADYAFLPFGAGKRKCIGDQFAMLEASVTLAMIINKFDFTLVGS
PKDVGMKTGATIHTMNGLNLVVSRRSEDNPIPETNDYWIQQHLSRGLNVNGRPYSTNEDA
AWTASSRDKNEGVVSRLVN-

##### Targeting:

SignalP 3.0 - NN: Yes, Cleavage Site CTA-FQ

SignalP 3.0 - HMM: Yes, Cleavage Site CTA-FQ, Signal Peptide Probability = 0.990

SignalP 4.1: Yes, Cleavage Site CTA-FQ, D = 0.724 (D-cutoff = 0.450)

ChloroP: No, Score = 0.497

#### 10) Thaps3_270370 (ZEP1)

No introns.

>Thaps3 chr_6:1420465-1418423
CTCCGTTCTCATTTTTCCTTTTCTTTCCAAACTTCTTCCGCTGCTGGAGCACCAACGGGT
AAACAGTGGCAGCAGAGGACAAATCAACTAGTACAGTACAACGAATCATGACGGTTAGAA
GAATCGCCTCGCTGGCCATTGGTATCTCGCTGTCCACCTTGACATGTGCATTTGTCACGA
TCTCCTCGTCTCGTACTACCATCAAACCTCTCAATGTCGTTGGCGAACAGGCCTCTTCCA
TCGGACCAGCAACTCTCCTCCGTAATCTCAAACAAAACCTACCACAAATAGATTGGCTAG
CGGAAGGTAAAGGCTCGCCATCCAACAAGATTGACATTCCTGATCATGTAGCTACAGTGC
TTGCTCAACCAAATGCTCCAAAGCGCGAAGCTGAGAGTGAAGAACGTACGCATAAGATCC
GCAGTAGAGCAAAGCAGGCTAGCGAAGATGCAATGGCATTGCGTGGTATGTTGATTGGAG
ATGATGACGCCAATGCATGGTGGCGTGAACAACGTTCTATTCCAGAGGGAGGACGAGTAG
TTACAACGGATGATCCATTGACTGTTCTTGTTGCTGGCGGTGGATTGGCTGGTTTGGTTG
TCGCTGCAGCATGCCATTCCAAAGGCATGAAAGTTGCATTGTTTGAGCAGGCTTCGTCTT
ATGCTCCTTACGGAGGCCCAATTCAGATTCAATCCAATGCATTGAGGGCTTTGCAGCAAA
TCAATCCTGAGATCTTTCAGGAGTTGGTTACTGCTGGAACATGCACTGCGGATCGTGTGT
CTGGATTGAAGATTGGATATAAGAAGGGAAACAAACTTGCTGGACTGTACGATGCAGGAG
ATTGGTTGGTGAGGTTTGACACTATCGGACCAGCGTTGGAAGCTGGATTGCCAGCAACTG
TGGTTGTGGATAGGCCAGTCATTCAGCAGATTCTGGTGAAATATGGCTTTCCTGAGGGCA
CCGTGCGTATCAAATCACGTATCCAATCGTATGAGGATCTGGGGAAGGGACGTGGAGTGA
GTGTCACCTTGGAAGACGGGACGAAAGCGTACGCAGATGTATTGGTAGGAGCTGATGGCA
TCTGGTCTCAAGTTAGAAAGAATCTCCACGGATTGGACGATGGAGCTGGAGGGTTCGCTG
CATCGGGCGCAGCAGGAGGTGCATTGGACGATGCCGAAGCACGCAAATTGGCACGTGATA
CAGTCGCAATTGCAGCCAAGGCCGATCGTCGCTTCTCTGGTTTCACATGTTACGCTGCAC
TGGCTCCTCATCGGGCATCCAACATCGAAAATGTGTCGTATCAAATCTTGTTGGGAGAGA
AGAAGTACTTTGTATCTACCGATGGAGGAGGAGACAGGCAACAGTGGTTTGCACTCATTC
GCGAACCTGCCGGGGGAGTGGATCCTGAGCCCACTCCCGAGGATCCTCACCCTAAGCTCA
CTCGTCTTAGGAAGGAATTTGCGTGCAATGGAAGTGGTGATGCTGATGGCAATGTGTGGG
ATCCATTTGCATTGGAGTTGATCAATGCAGCCTCGGAAGAAGACATCAAGCGTCGTGATT
TATACGACGGAGCTCCTCTCTTGACTACCCTTGACCCACAACGTTTGTTGAGTCCATGGG
CAAAGGGACCTGTGGCACTTTGCGGAGATGCAGCACATCCAATGATGCCTAACCTCGGAC
AAGGAGGATGCCAAGCCACAGAAGACGGATACCGTCTCGTCGAAGAGTTGGCAAAGGTGC
AGCATTCAAGAGATGTTCCAGGAGCACTCGGGAGATACTCTCGCGTTCGTGTGATTAGGA

CAGCCATTATCCAAGGTTTTGCTCAGCTTGGAAGTGATCTGTTGGTTGACTTTGATCTGA
TGATGACTATTCCACTCTTGGGTCCCTTCTTCCTGACAATGACACAGCTTTCCATGCCAT
TCATTTTGAGATACCTCTACACACCTTCTTTTTAAGAGAGATGCTTGTACGATTGTTGAG
AAGCAGATGTTAATGGGTAATATTCTGATAAGGTTTAAACTTTATGATGGAGTTGTCTGT
CCA

##### Translate:

MTVRRIASLAIGISLSTLTCAFVTI
SSSRTTIKPLNVVGEQASSIGPATLLRNLKQNLPQIDWLAEGKGSPSNKIDIPDHVATVL
AQPNAPKREAESEERTHKIRSRAKQASEDAMALRGMLIGDDDANAWWREQRSIPEGGRVV
TTDDPLTVLVAGGGLAGLVVAAACHSKGMKVALFEQASSYAPYGGPIQIQSNALRALQQI
NPEIFQELVTAGTCTADRVSGLKIGYKKGNKLAGLYDAGDWLVRFDTIGPALEAGLPATV
VVDRPVIQQILVKYGFPEGTVRIKSRIQSYEDLGKGRGVSVTLEDGTKAYADVLVGADGI
WSQVRKNLHGLDDGAGGFAASGAAGGALDDAEARKLARDTVAIAAKADRRFSGFTCYAAL
APHRASNIENVSYQILLGEKKYFVSTDGGGDRQQWFALIREPAGGVDPEPTPEDPHPKLT
RLRKEFACNGSGDADGNVWDPFALELINAASEEDIKRRDLYDGAPLLTTLDPQRLLSPWA
KGPVALCGDAAHPMMPNLGQGGCQATEDGYRLVEELAKVQHSRDVPGALGRYSRVRVIRT
AIIQGFAQLGSDLLVDFDLMMTIPLLGPFFLTMTQLSMPFILRYLYTPSF-

##### Targeting:

SignalP 3.0 - NN: Yes, Cleavage Site TCA-FV

SignalP 3.0 - HMM: Yes, Cleavage Site TCA-FV, Signal Peptide Probability = 0.960

SignalP 4.1: No, Cleavage Site TCA-FV predicted, D = 0.436 (D-cutoff = 0.500)

ChloroP: Yes, Score = 0.526

#### 11) Thaps3_261390 (ZEP2)

##### Exon 1:

>Thaps3 chr_2:1117778-1118447
TCCCCCTCTTCGGGTTGTGTCAGACGCAGACGTCCCTCTCGCATTTTGTAGTTCAGTCCA
AGCAACACCAATCAATTAAACAGAATGAAGCTCTCCATCGTGTGCTTCATTATATTAACG
TCGGCGACGTCAGCCTTCATCGCTCCATCAACCACACGCACATCAGTTGCCGTCACAACG
TCATCATTTGCAAATGTGCGAGGAAGTGCCCTCCAAATGGCCGATGACGAAGCCGACGCC
GACTTCAATTCTTCGGATTATGAACTCCTCGGAAGACCAGCTCGCCCTGGGCGTCCTCTC
AAAGTAGCCATTGCTGGTGGTGGTGTCGGTGGTCTCACAGCTGCGCTATGTATGTTGAAG
AAGGGATTTGACGTGACGGTGTACGAAAAGACTGCTGCCTTTGCTCGTTTCGGTGGACCC
ATTCAGTTTGCCTCAAATGCTCTTTCTGTCATCAAGGAGATTGACGAGGAGTTGTTTGAA
CGTGTAATGGATAAGTTCACCTTCACTGGTACAAGAGCTTGTGGTATCAAAGACGGTTTG
AGAGCGGATGGATCGTTCCGTATGACGAATGATTCCTTGGACTACTTGTGGAATCCCGAG
GCTCCTGCTGATTGGTTTGTCAAGTTCCCTTTGAGGCAGTGTGCTGATTTGTTTGGACTT
CCCTACACTG

#### Exon 2:

>Thaps3 chr_2:1118548-1119207
GCGTCATTGACAGACCCGATTTGCAGGAAATTCTTCTTGATGAGTGCAGAAAGATCAAGC
CCGATTTCATTCAAAATGGCAACCCAGTGAACGGATACGTTAGCAAAGGAAAAGGCAACG
GAGTGACTGTGAACCTCGCCGATGGAACAACTGCAGAGGCTGACGTCCTTGTTGGTTCGG
ATGGTATTTGGTCTGCTATTCGTGCTCAGATGTATGGGGAGGAGATTAAAAAGAGTTCAA
ACAATGCACTCAAACGTCAGGGCTGCACGTACAGTGGATATACCGTCTTTGCTGGAGAGA
CTGTGCTCAAGACGGAGGATTACTACGAGACTGGATACAAAGTGTACATTGGTCCTCAAC
GCTACTTTGTGACTTCAGATGTAGGAGACGGAAGAGTGCAGTGGTACGCTTTCTTTGCCT
TGCCGCCGGGTACGAAGAAAGCACCAAGTGGATGGGGAGGTACCGAGCGAACAGCGCAGG
ACGACCCAGAGGAGAATCTCGTAGATTACATCAAATCGTTGCATCAGGGATGGTCGGATG
AAGTCATGACTGTTCTTGATTCTACCCCTCCTGATAGTGTTGAGCAACGTGACTTGTACG
ATAGGCCACCTGAGCTATTGAGAAGTTGGGCTGATGGAAACGTCGTCCTCATTGGTGATG

#### Exon 3:

>Thaps3 chr_2:1119282-1119556
CTGTCCACCCGATGATGCCAAACCTTGGACAAGGAGGATGCCAAGCAATTGAAGATGCAT
TTGTTCTTTCTGAAACGCTGGAGGCATGCGAATCTACTCAAAAGTTGGAGGATGCTTTGC
AGGACTTTTACAAAAAGCGTATCGTTCGTGTTAGTATTGTGCAGTTCCTCAGTCGGTTAG
CGAGTGACTTGATCATCAATGCGTTTGATACACCCTGGAGTCCTCATGACGACCTCGGAA
AGTCGTGGAAGAGTTATTTGACTTTCTTCTGGAAG

#### Exon 4:
>Thaps3 chr_2:1119650-1119989
CCCATTCTTCAGTATGCCATCTTCCCTGCACAGTTTGCTTATCTTTACTCATACCACCCA
ACGGGAAACATGGGAGGTTTGCCTTCTGCTCTTGAAGCAAAGTGGAAGAAACAGCACGAA
GAGGACGCTGAGATGGCATTCAATAGGGTAGAGGAAGAGGGGCAATCAACTAGAGGACCG
AGTTTCTTCAAAATAGCAGAGTCAGAGACGGTGTTGGCTGCAAAGAAGATGTAATGACTA
AGAAGGAAGGAAGAGGGATATGATGCGTGTGTACTCAGGCTACAAGGGGTTGTTAGTGGG
TGAAGAGTGCAACTTATTTAGTACAAAAGTACAGAATGCA

#### Translated:
MKLSIVCFIILTSATSAFIAPSTTRTSVAVTT
SSFANVRGSALQMADDEADADFNSSDYELLGRPARPGRPLKVAIAGGGVGGLTAALCMLK
KGFDVTVYEKTAAFARFGGPIQFASNALSVIKEIDEELFERVMDKFTFTGTRACGIKDGL
RADGSFRMTNDSLDYLWNPEAPADWFVKFPLRQCADLFGLPYTGVIDRPDLQEILLDECR
KIKPDFIQNGNPVNGYVSKGKGNGVTVNLADGTTAEADVLVGSDGIWSAIRAQMYGEEIK

KSSNNALKRQGCTYSGYTVFAGETVLKTEDYYETGYKVYIGPQRYFVTSDVGDGRVQWYA
FFALPPGTKKAPSGWGGTERTAQDDPEENLVDYIKSLHQGWSDEVMTVLDSTPPDSVEQR
DLYDRPPELLRSWADGNVVLIGDAVHPMMPNLGQGGCQAIEDAFVLSETLEACESTQKLE
DALQDFYKKRIVRVSIVQFLSRLASDLIINAFDTPWSPHDDLGKSWKSYLTFFWKPILQY
AIFPAQFAYLYSYHPTGNMGGLPSALEAKWKKQHEEDAEMAFNRVEEEGQSTRGPSFFKI
AESETVLAAKKM-

#### Targeting:

SignalP 3.0 - NN: Yes, Cleavage Site TSA-FI

SignalP 3.0 - HMM: Yes, Cleavage Site TSA-FI, Signal Peptide Probability = 0.998

SignalP 4.1: Yes, Cleavage Site TSA-FI, D = 0.793 (D-cutoff = 0.450)

ChloroP: Yes, Score = 0.551

### 12) Thaps3_7677 (VDE)

No Introns.

>Thaps3 chr_8:842679-841117
AGGACAACAACCAACACCCTTGCCATTGAACACTCTGTTGCTGCTCTACCCAATAGACGC
CATGAAGCTGTTCCTTTCTCTCGTGCTGGCGGCTGCACCAGTGTCTTCCTTCGCACCATC
AAACCCAGTTGTATCTCGTACTCATTCATCGGTACACTCCCAACAACATAATCACGTGCT
CGAAGCTCACAACGACAACATGGATGACATTACCTTCTCCCTATCCGCCAGGAATATCAA
CAACGAGATCGTAGAGCGAATTGGCAAAGTAACCACCTCAGCTCTCCTCGCATTGACGCT
CAGTTTCTCTGCCATCACATCGCCCATCTCCGGTCCCAACGGCGACGTACTATCGTCCAT
CCCATCTGCCAACGCTGCCGATGGTGCAAAGATCGGTCTATGCCTTGTCAAGAAGTGCAG
AGTTCCTTTGGCCAAGTGTATCACCAACCCAAACTGCCTTGCTAATGTGATTTGCATCAA
TTCTTGCAACGGAAAGGAAGATGAGACTGGATGTCAGATTAATTGTGGAAACGTCTTTGA
GAATGACGTTGTTGGGGAGTTCAACAAATGTGCCGTCACCGATATGACGTGCGTTCCTCA
AAAGAAAGACGATGGAAGTTATCCCGTCCCATCCAAAGATGTATTGGTCCAATCGTTTGA
TACCAAACTATGGAACGGAAGATGGTTCATCACCGCAGGGCAGAACAAGCTCTTTGATAC
GTTCCCATGTCAAGTCCACTTCTTCACCGAGACTGCTCCAGGCAAGTTCGTCGGGAAATT
GAATTGGCGTATCGAAGAGCCTGATGGAGAATTCTTCACTCGTGATGCTGTGCAAGAGTT
TGTTCAGGATCCCAACAATCCTGCTCACTTGATCAACCACGACAATGAATATTTGCATTA
CCAAGATGATTGGTACATTGTGGATTATGCCGCGGATGATAACAAGGAGGGTGTTCCTCC
CTTTGCATTTGTGTATTACCGTGGTGAGAATGATGCATGGATTGGATACGGTGGAGCTGT
GGTGTACACTCGTGATTCAAAGTTGCCAGAGTCTCTCCTACCACGTCTTCGTGAGGCTGC
TAAGAAGGTAAACTTTGACTTCGACAAAGACTTTGATCTCACGGACAACTCGTGTAAGGC
ACTTGAGAAGGGAGAGGAGGTCGTGTTGAGGGAGAAGTTTGCTGGTAAGATGGCCATTCA
GACGGAGAAGCAGTTGCAACAGCAGGCTGTGTTGGCACGAACTGCGGCTAGTAATACTGT
AAAGGGTGAAGTGACTGCTGTTGAGAAATCGCTTCAGAAGATTGAAGAGAAGGCTTTGGC
GTTTGAGAAGGAATTGATGAAGGATGTTGTTTCAGTGGAAAAGGAGATCGTAAAGGAAGT
TGAAGAGGTAGAGAAGGAGATTGTTCAAGAGGAACAAAAGATCTTTGGTGGTATTAGATA

GACTGGTTGTAGTTAGTGGAGTAAATGCTACTCATTGACTTTATCTTGGAATGATATTTG
AGAAGACGTCTAAATGCACTGCGAGTAAGAAGATCTTTATGTGAATGGGAAAAGTATGAA
TGG

##### Translated:

MKLFLSLVLAAAPVSS
FAPSNPVVSRTHSSVHSQQHNHVLEAHNDNMDDITFSLSARNINNEIVERIGKVTTSALL
ALTLSFSAITSPISGPNGDVLSSIPSANAADGAKIGLCLVKKCRVPLAKCITNPNCLANV
ICINSCNGKEDETGCQINCGNVFENDVVGEFNKCAVTDMTCVPQKKDDGSYPVPSKDVLV
QSFDTKLWNGRWFITAGQNKLFDTFPCQVHFFTETAPGKFVGKLNWRIEEPDGEFFTRDA
VQEFVQDPNNPAHLINHDNEYLHYQDDWYIVDYAADDNKEGVPPFAFVYYRGENDAWIGY
GGAVVYTRDSKLPESLLPRLREAAKKVNFDFDKDFDLTDNSCKALEKGEEVVLREKFAGK
MAIQTEKQLQQQAVLARTAASNTVKGEVTAVEKSLQKIEEKALAFEKELMKDVVSVEKEI
VKEVEEVEKEIVQEEQKIFGGIR-

##### Targeting:

SignalP 3.0 - NN: Yes, Cleavage Site VSS-FA

SignalP 3.0 - HMM: Yes, Cleavage Site VSS-FA, Signal Peptide Probability = 1.000

SignalP 4.1: Yes, Cleavage Site VSS-FA, D = 0.682 (D-cutoff = 0.450)

ChloroP: Yes, Score = 0.529

#### 13) Thaps3_22076 (VDL1)

##### Exon 1:

>Thaps3 chr_4:141771-142931
CCCATAACAGCGCTTCCAACACCGCAACATGAGACCGTCAACCTCTGCCTTGACAGTCGT
CCTAGGCACCATTGCGCTTGTCAGCTGCAGCCAGCTCAACAACAATGTATCCGCCTTCTC
TACAAGATCATCGTCGTTGACGCAAAGACACAAGACATGCACCATCACAACATCATCATC
ATCTCTCTACGTGAACCCCAACAACGATGATGACAACTCCAACAGATCCCAACACAAACC
CAATCCCTTCCTCTCAGCGGCACTAACCGCCGCAGTCACCACATCCCTCTTCCTCTCATC
TCTCCCCAGTGCCACCTTCGCCTCCACACCGGCCTCCACAACGCAAAAGTACGACGGCTT
CGCCGAGTACGCCAAGGAAAACAAAATGGAACAATCGGACGTAGGATGCTTCATTAACAA
GTGTGGCGATCAGACGAAACAACTGTTTAGTAATCCCCGTGGTATCAAGGGGGTGTCGTG
TTTGGGACGGTGCAAGGGGGAACAATCGTGTGCTACGCGGTGTTTTGCTGAGTTTGGGAG
CGAGGATTTGGACAACTGGTTATCGTGCACTATTGAAGATTATGAATGTGTGAAAGTTCC
AAAGAATATTGACAACTCTGCCGAGAATGTGGGGTATGATACTACCGTGAAGAAGTTTGA
TCCGTCAACGTTGGTGGGAAAGTGGTACAAGACGGATGGACTGAATCCCAATTACGATCT
GTTCGATTGTCAATCTAATACGTTTGACTTTTCAGATGATACGAAAAAGGAGTTGGATAT
GGGTATCTTCTTTAGAGTGCCGCGTCCAGAAGAATACGGAGGTGGATTCTGGGAGAACAG
TCTTACAGAGCACATGATTGTTGATGCCGTATCACCCGAGTTAGACAACCCTACTGGAAG
AACGATGCATACCGCTGGTAAGATGTATGGGCTCAAATTCACTGAGAACTGGTACATACT

CGGAGAATCCAATGGTGATAATGATATCCCTCCGTTTAAGTTGGTGGCGTATAAGGGGCA
TACGTTGCAAGGGAATTATGAGGAGGCGTTTGTGTATGCGAAGGAGAGTGTTTTGCCGAA
GGAGGCTGTGGGGGCGGTGAGGGAGGCTGCGGCGAAGGCTGGACTGGACTTCGACAAGTT
CACGAGGATTGATAATACTTG

##### Exon 2:

>Thaps3 chr_4:143069-143272
TCCAACCACAACAAAATCACTGAATGATGCATCGGCTGGAACTGGAACGTCTACCACAGA
CTGGGTTGATCTTGTAGTTGGTGAAGGAGGAGTTATTGATTGGGTTGTTCCTGGATGGAG
AGGAGAGTACAAAAACTAAAACAGGTTAGAGCGTCGAAGAAGAAGATGCTTTTCTAACCT
AAAATCTGGGTATTGTCGTCCCGA

##### Translated:

MRPSTSALTVVLGTIALVSCSQLNNNVSAFSTRSSSLTQRHKTCTITTSSS
SLYVNPNNDDDNSNRSQHKPNPFLSAALTAAVTTSLFLSSLPSATFASTPASTTQKYDGF
AEYAKENKMEQSDVGCFINKCGDQTKQLFSNPRGIKGVSCLGRCKGEQSCATRCFAEFGS
EDLDNWLSCTIEDYECVKVPKNIDNSAENVGYDTTVKKFDPSTLVGKWYKTDGLNPNYDL
FDCQSNTFDFSDDTKKELDMGIFFRVPRPEEYGGGFWENSLTEHMIVDAVSPELDNPTGR
TMHTAGKMYGLKFTENWYILGESNGDNDIPPFKLVAYKGHTLQGNYEEAFVYAKESVLPK
EAVGAVREAAAKAGLDFDKFTRIDNTCPTTTKSLNDASAGTGTSTTDWVDLVVGEGGVID
WVVPGWRGEYKN-

##### Targeting:

SignalP 3.0 - NN: Yes, Cleavage Site VSC-SQ

SignalP 3.0 - HMM: Yes, Cleavage Site VSC-SQ, Signal Peptide Probability = 0.970

SignalP 4.1: Yes, Cleavage Site VSC-SQ, D = 0.694 (D-cutoff = 0.450)

ChloroP: Yes, Score = 0.568

#### 14)Thaps3_11707 (VDL2)

##### Exon 1:

>Thaps3 chr_22:194840-195897
ATTTCCTTCACTTTTACTTTGATTCAAATCTTAATAGACATAGCCTTTTTACGTGGCAAA
GGTCGCGTCATTTGCCAGGAGGTTGAAGCAAAACAGTGGACGACGTCGCTCGTGAGGACG
GACAGCGAGAGAGGCCACCCCAGCTGTTGCATCCCAACAACCATCCATCGTAGACTTCAA
TACACATCAACTGATAACAGAGTGGTGCTGATACTGTCAGTTAGAGTTCATTGGATTAGA
AAGACTGCAAGTGAGATACACCACAGTCCAGATCGCCTGAGTTGTAATAGTCAAAGTTTT
TGCTCCCTCAATACAAAACAAAATGTCGGCATCATCATCATCAACAACAACAACAACAAA
CGCTGGAAAGAGGGCACGATCGTGGCCGTCTTCATCCGCATCAACATCATCAATGCCTAC
TCGGTCGATACTGATACTGGCAACCTTTCTCTCATTGACGTCGTCGTCGTCGTCAACATC
GGTGGAAGCTGCCTTTGTCGGCAGCCCTGCCGTTGGACTTAGATCTCACACTGCTGCGTC

AACATCAAAGCAGCAGTCGTCTCTCTACGCTCAAAAGAAGAACAACATCGACTCATCCGA
CAACCCGCTATCATATCTATTCGATCTATCCTCCGATCCAGAAACCAAACGTCGCCTCCA
AAAACAAACAGCCACCCTCTTCTCCACCCTCGGCTTCTCCGCCCTCTTCCTCACCAATCC
ACTCATCCCACACCTCCCCTTCTCCCCCTCCCTCAGCAGTGCCAACGCCGAAGACGAACT
CTATGCTAAATATGGTGGCAAAGGCCTCGATACATCCTTAGTCGACAAAGACTGTCTCGT
CAATCAATGTCAAGTGCAAGCCAAAGCGTGTCTTCAGGATGATCCCGATTGTAGAAAGGG
ATTAACGTGTACTGCCAAGTGTTTGGGGGATAATGCGTGCATTACGGGGTGTTTTGCTAG
GTATGGGAATGAGAATTTGGATGAGTTGTTGAAGTGTACTATTGAGGATCATGAGTGTAT
TAAGGTGGCTATTTTGGAGGGTGGGGGCGATGTGCTTG

#### Exon 2:

>Thaps3 chr_22:195996-196062
GGCGAGAGCCAAAGTCGCCTGCTCCTACTGTTCAAGGGTTTGATCTAGCCAGTATGGAAG
GGACTTG

#### Exon 3:

>Thaps3 chr_22:196169-196611
GTATAAAGTAGCAGGCTACAACCCCAACTACGATTGCTACGCCTGCCAACGAAACACCTT
CTCCTCACCCGAAGGCGGTCTCTCCGACTCACTCCAACTACCAACAGGAGGAATACTCGG
CTCTCTATCCAACGCTGTAGGTTCCATCGGTGCAGATCGCTTACAAGTCGATGTGGAATT
CAGTATGCCGAGGTATTTGCCAGATGGTAGTCCTCAGCCACCGAGTGGAGTGCGCGAATC
ATTCATTAGTAGTGCTGATTCAATGGAAGGGAGTGGGTTGCAGAGTGTGGGGTACAATCA
GTACTCGACTCATGAGACAATGGTGTTTGATACGGTCAAGAGTAATGGTGTGGGGGAAGC
TGTGAAATTGGCTTTGGGTAAGAGGGGAGAGGAGAAGTTGTATTCGAGGACGGCTCATTC
GGAAGGAGAGATGTTTGGACTGA

#### Exon 4:

>Thaps3 chr_22:196740-197411
AATTCTGGGAGAACTGGTACATCATTGGCCAAAACAACCCCGGCCAAGACGAGTTCAAAT
TCGTCTACTACAACGGCAAGACACGTCAAAATACCTACGACGGTGCCTTCATCTACTCCC
GATCACGCACCCTCTCACCCGCGTCCATGGAGAAAGTCTACAAGATTGCCAAGGATGCGG
GTATGAATCCTGATCAGTTTTGTAAGATTCAGAACTCTTGCTTTGACGGTGAGGATGATA
AGCAAGAGATGATGATGATGAATCCTCAGAGAGAAGGACTGGGTAGTCCTTCCAATCCAT
TTAGGGGTATTTTGGCATCTACGAAAGTATCTCAATTCTTGGGAGTTGAATCTGTGGCGG
CCGAGACTACGTACAACGAACCAAAGAGTACGATATCGTCCAACTTTCTTCAAGGGAGTC
AGGCTACCAACCGCAAAGCGGATGCCGTTCAGGAGAGGCCGTGGTGGAAGGAAATGGGAG
ATTATTTGGAAGATCCTAGACGGCATTTCCGTTTGATGGATAGTCTGAGAACAGATATGG
ATTGGCCGGATTATATCAAAGAGAAGAATTGGTGAAGGAGTTCGGCATGCTGAGTAGTTA
CGAAACTACTTTTGTTTGGAGTATGTGTACTATTTCTATGATCCAAAGATGTAAGGGGTA
ACTTAAAGAAAC

#### Translated:

MSASSSSTTTTTN
AGKRARSWPSSSASTSSMPTRSILILATFLSLTSSSSSTSVEAAFVGSPAVGLRSHTAAS

TSKQQSSLYAQKKNNIDSSDNPLSYLFDLSSDPETKRRLQKQTATLFSTLGFSALFLTNP
LIPHLPFSPSLSSANAEDELYAKYGGKGLDTSLVDKDCLVNQCQVQAKACLQDDPDCRKG
LTCTAKCLGDNACITGCFARYGNENLDELLKCTIEDHECIKVAILEGGGDVLGREPKSPA
PTVQGFDLASMEGTWYKVAGYNPNYDCYACQRNTFSSPEGGLSDSLQLPTGGILGSLSNA
VGSIGADRLQVDVEFSMPRYLPDGSPQPPSGVRESFISSADSMEGSGLQSVGYNQYSTHE
TMVFDTVKSNGVGEAVKLALGKRGEEKLYSRTAHSEGEMFGLKFWENWYIIGQNNPGQDE
FKFVYYNGKTRQNTYDGAFIYSRSRTLSPASMEKVYKIAKDAGMNPDQFCKIQNSCFDGE
DDKQEMMMMNPQREGLGSPSNPFRGILASTKVSQFLGVESVAAETTYNEPKSTISSNFLQ
GSQATNRKADAVQERPWWKEMGDYLEDPRRHFRLMDSLRTDMDWPDYIKEKNW-

Based on the targeting analysis of the above peptide (a delayed signal sequence and clear chloroplast targeting), it appears that the protein starts at the second in-frame methionine. The following analysis pertains to the latter.

##### Targeting:

SignalP 3.0 - NN: Yes, Cleavage Site SSS-TS

SignalP 3.0 - HMM: Yes, Cleavage Site VEA-AF, Signal Peptide Probability = 1.000

SignalP 4.1: Yes, Cleavage Site SSS-TS, D = 0.799 (D-cutoff = 0.450)

ChloroP: Yes, Score = 0.554

#### 15) Thaps3_270211 (VDR)

##### Exon 1:

>Thaps3 chr_2:1186955-1187856
GGGGGTGATGAGGAGCCGAGGGGCGCCGAGCAACGACGCGAGCAACTACACAACGAGCAA
CCCCACAACAAGAACAATGGCAATGGTGCTGCTGATACGGACAGCAGTGATAGCTTCATA
TAGCCTAACATTGACATCGGCATTTTCCTCATCAATAAGGCCGACTTGCCGGACATTTCG
TCAGAGTACACCACATCATGCAACCGCTTCCAACGTCGACATCATCGGTACAGTGGCACT
TCTCGTGCCTTCTTCGTCCACTGAGCTGTCAAAGTATGGATCCAAATCTCCAGCACCTCG
ACCATCCTATCAAGAAGCAGCTGAACACTTGGCTCGCAAAATAAGTCACTTCTCTGACGG
TCGAATAGAAGCAACAGTAGTAACACCATCAACTAATCAAGATGACACGGACGACGTCTG
CCTTACATCTAATGCACTGATTGCTTTGGGAATCACTGACCCAGCGGAAGTCCAATACCT
TTCCACGACGTTTCGTAAACGTCGCACATCTCATCAAGAAACGTCCTCTTATAACACGTG
TCAATTTGCATTAGACTGCGGCAGCAACAACTATGCACCTCTCGTTGGACCATGGGATGA
AGCCAATCCATCCATTCTCGCCGAAATTGCTCCGTGGACAGGTGTGGCATCTGGCAAACG
TCTGACAGAACAAATGAATGGCTTGTTTGAAAAGCAAACATCTGATGAGTTTGCCTTGGC
AGTTATGCTCTTTTTCAATCGGTTTTCTGGCGCTGCTATTCCTTGGGTGCAACACTCAAT
TGATGTAACTTGGGAGAAGGGATTGGTCCAGAATGCAAAGGAGATCTTCTCTATGATTAC

AAAGTGTGGACCTTGCATTACCAAGTGTTTGAACGATGAGAATTGTTCTCAGTGTATCAA
CG

#### Exon 2:

>Thaps3 chr_2:1187934-1188181
CACTTGACAAGATCGACACACGAGACCAAGTCACAAGTTATAGAACAGTCGTGTCGTTTG
AAAGTGAACTGCTTAGGGATTTCAGCTTGTGTATTTTGCAAAAGAATAACATCTTCGAGT
GCTCAGCTGAGATCCCAGAGTTGCCAGTTGTCAAACCGATGAGCACATGGAGAGGGAAGG
ATGTTACGACGGACGTCGCGAGAGGCATTATGATTGGGCACTTAGAAGGAGCGGGGGGAT
CGCTAGAG

#### Exon 3:

>Thaps3 chr_2:1188256-1188336
GGGAATTTGCAACTTGGTGTCTCTTGGAAGGTGGCTTGCGGAGCAAACGTTGCCTATGAT
CAGTTTCCCTCTCAAAATCAG

#### Exon 4:

>Thaps3 chr_2:1188422-1189161
TTGTTTTACCCATCTGCAAAGGGGAAGGATCTTTGGTACGACCCTGTATTCCGAGTAGAA
ACAATTGACGGTAGAAATGTCTGGTGCAAACGTCACTACAAAGTCAGGAATGGAGAAACT
CCTGGTACGTTCAAATTCTCGGTATTGGACAATGGCGTAACGAGCAACGAGTTCTGGACG
ATTGTCGGGGCTGCTGACGACTTGTCGTGGGTTGTATTTCATTACGCTGGAGCTGCTGGC
GCTGTAGGCCAGAGGTATTTGGGAGGACTGCTGTGCACACCAACGGGAGAGCTGCCACCA
GAAGAAGACCTTGGACACATCTACAATTTTCTTCGATCGGCGGAAATTGAACCGTGGGAG
TTATTTGTTGTGGACAATGATGATCAGTCGCCTGGTGCGTTAGCAGCAGGCGCTCCACCG
CTGGATTACTTCAGAAAGACTGCATCGGTCATTGGCTGAGCATTCGCCTGTACTATGTTT
AATCCTTAAGCCAAGATACTCTCAAGCAACGGCGAATGCTCCGTCTTGCGAAAGTTATGC
ATTACACCCCAGTGCTCATCGTCGAACCGAAATATCACCGCAACGTCAAGAGGCAAGTCG
CTAATAAAAGATGCAGAAGGCAACTGACAAAGCAGCTTTGATCCTGGGAACGTCGCCTTT
TTTGTAACACTTGCCAAGCTTGTAATTCTGCCGACACGTCCGTTGAGGAATGGTAACAAA
GGAACCTCATTTGACTCGTA

#### Translate:

MRSRGAPSNDASNYTTSNPTTRTMAMVLLIRTAVIASYSLTLTSAFSSSIRPTCRTFR
QSTPHHATASNVDIIGTVALLVPSSSTELSKYGSKSPAPRPSYQEAAEHLARKISHFSDG
RIEATVVTPSTNQDDTDDVCLTSNALIALGITDPAEVQYLSTTFRKRRTSHQETSSYNTC
QFALDCGSNNYAPLVGPWDEANPSILAEIAPWTGVASGKRLTEQMNGLFEKQTSDEFALA
VMLFFNRFSGAAIPWVQHSIDVTWEKGLVQNAKEIFSMITKCGPCITKCLNDENCSQCIN
ALDKIDTRDQVTSYRTVVSFESELLRDFSLCILQKNNIFECSAEIPELPVVKPMSTWRGK
DVTTDVARGIMIGHLEGAGGSLEGNLQLGVSWKVACGANVAYDQFPSQNQLFYPSAKGKD
LWYDPVFRVETIDGRNVWCKRHYKVRNGETPGTFKFSVLDNGVTSNEFWTIVGAADDLSW
VVFHYAGAAGAVGQRYLGGLLCTPTGELPPEEDLGHIYNFLRSAEIEPWELFVVDNDDQS
PGALAAGAPPLDYFRKTASVIG-

#### Targeting:

SignalP 3.0 - NN: No

SignalP 3.0 - HMM: No, Signal Peptide Probability = 0.163

SignalP 4.1: No, D = 0.173 (D-cutoff = 0.500)

ChloroP: Yes, Score = 0.579

Based on the clear predicted chloroplast targeting, the analysis was repeated with the peptide starting with the second in-frame methionine, with the following results:

SignalP 3.0 - NN: Yes, Cleavage Site TSA-FS

SignalP 3.0 - HMM: Yes, Cleavage Site TSA-FS, Signal Peptide Probability = 0.975

SignalP 4.1: Yes, D = 0.654 (D-cutoff = 0.450)

ChloroP: Yes, Score = 0.561

Based on the clear ER and chloroplast predicted targeting, it is likely that the peptide begins at the second in-frame methionine.

#### 16)Thaps3_bd_1474

No available RNA-seq data for the unmapped “bottom drawer” sequences.

Gene model predicted by JGI (below) cuts off abruptly on the N-terminal side due to the way the unmapped sequences were assembled. Some N-terminal sequence is likely missing.

##### JGI-Predicted Exon 1:

>Thaps3 bd_37x91:13831-12903
CTTATGAATCAAAAGATGTTGACGTTGGGTGAAAAAATTCAGACCGCTCCTCCTCTTCTTCCTATGCT
TATTGAGGGACAGTCATTCATTGATGCTCAGGATGAGTTGAGTGTGACGCAGTTCATGAGGAAGTACGGT
ATGCCTGAGAGAATCAACGAGGAGGTGTTTATTGCGATGGCCAAGGCGTTGGACTTTATTGATCCTGATA
AGTTGAGTATGACTGTGGTGCTTACGGCTATGAACAGGTTCTTGAATGAGAGTAATGGACTTCAGATGGC
ATTCTTGGATGGAAATCAGCCTGATAGGTGGTGCACTCCCACCAAGGAGTATGTGGAAGCACGCGGAGGA
AAGGTCAAATTGAACTCTCCCATTAAGGAGATTGTGACCAACGACGATGGAACTATCAATCACCTTCTCC
TTCGATCTGGCGAGAAGATTGTGGCCGATGAATACGTCTCTGCCATGCCCGTGGACATCGTCAAACGTAT
GCTTCCCACAACGTGGCAGACTATGCCCTACTTCCGTCAGCTTGACGAACTTGAGGGCATCCCTGTTATC
AACTTGCACATGTGGTTCGATCGTAAGTTGAAAGCAGTCGACCATCTTTGCTTCAGTCGCTCCCCACTCC
TTTCCGTCTACGCTGACATGTCCGTCACATGCAAGGAGTACGAAGATCCCAACAAGTCCATGTTGGAATT
GGTCTTTGCTCCCTGCTCTCCTATTGCCGGAGGAAATGTCAACTGGATTGGAAAGTCAGATGAGGAAATC
ATTGATGCTACCATGGGTGAGCTTGCTCGCCTTTTCCCTACCGAGATTGCGAATGATGATAAGTGGCCTG
CTACGAAGATGCAGGGACCTAATGGACAGGCAAAGCTTGAGAAGTATGCTGTTGTGAAGGTGCCAAGGAG
TGTGTATGCTGCCATTCCTG

##### JGI-Predicted Exon 2:

>Thaps3 bd_37x91:12821-12475

GACGTAACAAATACCGCCCCAGTCAGACCTCCCCCATC
CCACACTTCACCATGGCTGGATGCTATACCTCACAAAAGTTCCTCGGATCCATGGAGGGTGCCACCCTCG
CCGGGAAGCTTGCTGCCGAGGTCATTGCCAACCGTGCCCTCGGAAATGCGGATAAGCCAGTCAAGGAGAT
TCAGCAACACATTATCGACTCGGCTAGTAAGCATGTTGTGAAGGAGCCAGTGGGTGTGAAGGGAGAGGGA
GCGATTGCATTTGGAGGGGGGTATACTGTTGGAAAGAAGGAGGAGGATTTGTTGAGGGAGTCGGATCCTG
CTCAGTATGAGTTGGCAGTAGCCAAGTAA

#### Translated:

MNQKMLTLGEKIQTAPPLLPMLIEGQSFIDAQDELSVTQFMRKYGMPERINEEVFIAMA
KALDFIDPDKLSMTVVLTAMNRFLNESNGLQMAFLDGNQPDRWCTPTKEYVEARGGKVKL
NSPIKEIVTNDDGTINHLLLRSGEKIVADEYVSAMPVDIVKRMLPTTWQTMPYFRQLDEL
EGIPVINLHMWFDRKLKAVDHLCFSRSPLLSVYADMSVTCKEYEDPNKSMLELVFAPCSP
IAGGNVNWIGKSDEEIIDATMGELARLFPTEIANDDKWPATKMQGPNGQAKLEKYAVVKV
PRSVYAAIPGRNKYRPSQTSPIPHFTMAGCYTSQKFLGSMEGATLAGKLAAEVIANRALG
NADKPVKEIQQHIIDSASKHVVKEPVGVKGEGAIAFGGGYTVGKKEEDLLRESDPAQYEL
AVAK-

Genomic sequence disregarding the predicted intron:

>Thaps3 bd_37x91:13831-12475
CTTATGAATCAAAAGATGTTGACGTTGGGTGAAAAAATTCAGACCGCTCCTCCTCTTCTTCCTATGCT
TATTGAGGGACAGTCATTCATTGATGCTCAGGATGAGTTGAGTGTGACGCAGTTCATGAGGAAGTACGGT
ATGCCTGAGAGAATCAACGAGGAGGTGTTTATTGCGATGGCCAAGGCGTTGGACTTTATTGATCCTGATA
AGTTGAGTATGACTGTGGTGCTTACGGCTATGAACAGGTTCTTGAATGAGAGTAATGGACTTCAGATGGC
ATTCTTGGATGGAAATCAGCCTGATAGGTGGTGCACTCCCACCAAGGAGTATGTGGAAGCACGCGGAGGA
AAGGTCAAATTGAACTCTCCCATTAAGGAGATTGTGACCAACGACGATGGAACTATCAATCACCTTCTCC
TTCGATCTGGCGAGAAGATTGTGGCCGATGAATACGTCTCTGCCATGCCCGTGGACATCGTCAAACGTAT
GCTTCCCACAACGTGGCAGACTATGCCCTACTTCCGTCAGCTTGACGAACTTGAGGGCATCCCTGTTATC
AACTTGCACATGTGGTTCGATCGTAAGTTGAAAGCAGTCGACCATCTTTGCTTCAGTCGCTCCCCACTCC
TTTCCGTCTACGCTGACATGTCCGTCACATGCAAGGAGTACGAAGATCCCAACAAGTCCATGTTGGAATT
GGTCTTTGCTCCCTGCTCTCCTATTGCCGGAGGAAATGTCAACTGGATTGGAAAGTCAGATGAGGAAATC
ATTGATGCTACCATGGGTGAGCTTGCTCGCCTTTTCCCTACCGAGATTGCGAATGATGATAAGTGGCCTG
CTACGAAGATGCAGGGACCTAATGGACAGGCAAAGCTTGAGAAGTATGCTGTTGTGAAGGTGCCAAGGAG
TGTGTATGCTGCCATTCCTGGTGAGTGAAAAAGAGTGTGCGGTGAATCTTCGTCGTCTACCTTGGCCAAC
TACTGACTCTTGTTCTCTTTAAATCATATCAGGACGTAACAAATACCGCCCCAGTCAGACCTCCCCCATC
CCACACTTCACCATGGCTGGATGCTATACCTCACAAAAGTTCCTCGGATCCATGGAGGGTGCCACCCTCG
CCGGGAAGCTTGCTGCCGAGGTCATTGCCAACCGTGCCCTCGGAAATGCGGATAAGCCAGTCAAGGAGAT
TCAGCAACACATTATCGACTCGGCTAGTAAGCATGTTGTGAAGGAGCCAGTGGGTGTGAAGGGAGAGGGA
CGATTGCATTTGGAGGGGGGTATACTGTTGGAAAGAAGGAGGAGGATTTGTTGAGGGAGTCGGATCCTG
CTCAGTATGAGTTGGCAGTAGCCAAGTAA

Peptide sequence disregarding the predicted intron:

MNQKMLTLGEKIQTAPPLLPMLIEGQSFIDAQDELSVTQFMRKYGMPERINEEVFIAMA

KALDFIDPDKLSMTVVLTAMNRFLNESNGLQMAFLDGNQPDRWCTPTKEYVEARGGKVKL
NSPIKEIVTNDDGTINHLLLRSGEKIVADEYVSAMPVDIVKRMLPTTWQTMPYFRQLDEL
EGIPVINLHMWFDRKLKAVDHLCFSRSPLLSVYADMSVTCKEYEDPNKSMLELVFAPCSP
IAGGNVNWIGKSDEEIIDATMGELARLFPTEIANDDKWPATKMQGPNGQAKLEKYAVVKV
PRSVYAAIPGE-

No predicted targeting to the ER or chloroplast for either peptide, however due to the high likelihood of a missing N-terminus, such analysis is inconclusive.

#### 17)Thaps3_21900

No introns.

>Thaps3 chr_3:1688852-1690753
CAACGACCGACGGCACACAACCCCAACACCAAAATGATCCGTTCAATATCAGCGTTGGCA
CTCCTGGCAGCCTGCTGCCCATCCGTTTTTTCCTTCGCTCCCCTATCAGTATTTAGAGCA
AATGCACCATCTTCACTGGCATCGACCACCTATCAAGATGAAGTAGACTGCATCGTTATC
GGCAGTGGCATCGGTGGTCTCTCGTGTGCAGCCCTCTTAGCAGCCACAGGTCGTACCGTA
CGCGTCCTAGAGCAACATTATGAAATAGGAGGATGCGCACATGCATTTTACATGGATATG
AATGGCAAGACGGTACCTTCGTCTGCGCTGAAGGATGACCCTACAAAGAAAGGAGAGTTG
TTTCATTTCGAAGCAGGGCCTAGTTTGTACAGTGGACTATCGGAGGAGAGAACACCGAAC
CCGCTTAAGCACATCTATCAAATGATCGAGGAAGAGCCAGAGTGGTTAACGTACGATCAG
TGGGGTGCCTTCTTGCCTGAAGCCCCAGAGGGATATCAGATGAGTATTGGAGCAGAAAAC
TTTTGCAAGATCCTCGAGACTTATGGAGGCGAGGGCGCAGTTGAAGATTGGGAAAAGCTT
GCCGAGCAATTGAGACCAATGGCGGGGGGTATCAAAGGCATTCCACATGCAGCAATACGC
GGTGATTGGGGTATCTTCCTGACTCTCATCTTGAAGTATCCTCTCTCCTTCATGAATGTG
CTCAAGTATGCTCCTGCATTCACTGCTCCTTTTGATTTGGACAAGTTGGGCGTGACAAAC
AAGTTCTTGCGGAATTATCTTGAAATGCTGGCTTTCCTTTTGCAAGGCTTGCCAGCTGAT
CAAACATTGACCGTAGTCATGGCGTATATGGTAGAGGACTTCTTTCGTGAAAATGCAGTG
ATGGATTTCCCAAAAGGAGGATCTGGTGAGCTTATGGGTGCTTTGGCGAGAGGAGTGACA
AAACGTGAGGGGTGTTCCGTCGAAGTATCTACATCTGTGGACGAAGTTATTGTTGAGAAT
GGACGTGCTGTTGGTGTGAAATTGGCAAAGAGCGGACGTATCATCAAAGCGAAGGAAGCT
GTTATCAGCAATGCTGATCTTTACAACACATACAAGTTTGTTCCAGAGGGAAAGCACGAG
GGATTCGACAAAGAGAGAATTGAGTACCTTGGTCTTACTGCAAAGCCGAAAGATGGCTCA
GTTCCATTCTGCAAATCATTCATGCACTTGCATCTCGCTGTAAAGGCAGAACTCATACCA
GAAGATGCGCCTCCACAATGGACTGTTGTTCAGGATTGGGATAAAGGTATCGATGCAACT
GGAAACGTAGTGGTTGTATCGGTCGGAAGCAAACTTGATCAGTCATTAGCACCGCCTGGA
TATCACGTTATCCATGCTTACACAGCGGGAAATGAATCTTACGAAGACTGGGAACAATTT
GAACATCTGATGGATGATGCTGCCGTCAGAGACAAAGATGCAGCATACCAAACATTCAAA
GACGAGCGAGCTCAGCCGATTTGGGATGCCATCCAAAAGCGTGCCCCTGCCGTCGTCAAG
GGTGCTTGTGTTATAGAAAAGGTAGCCACCCCATTGACTCACGCTCGATTCCTCAATAGA
CATCGTGGAAACTACGGATTAGCCATTGCGCCGGATAATGCAGAAGGCTGGAAGTTCCCA
GATGTAAAGACGCCTCTTGAAGGATACTACAGGTGCGGTGATTCCACAACGTCTGGCATT
GGAGTTCCGGCGACGGCAAGTAGTGGAGCCGTTTGTGCTAATGCGATCATGTCCGTTTGG

GATCAGCTTTCATTGAATCAAAAGATCAAAATGCCGTGAACGAGTAAGTATAATCAATCG
TTCTTTCTCTGTAATATATTTAAACTTCAAGCAGCTCAATAC

##### Translated:

MIRSISALALLAACCPSVFSFAPLSVFRANAPSSLASTTYQDEVDCIVI
GSGIGGLSCAALLAATGRTVRVLEQHYEIGGCAHAFYMDMNGKTVPSSALKDDPTKKGEL
FHFEAGPSLYSGLSEERTPNPLKHIYQMIEEEPEWLTYDQWGAFLPEAPEGYQMSIGAEN
FCKILETYGGEGAVEDWEKLAEQLRPMAGGIKGIPHAAIRGDWGIFLTLILKYPLSFMNV
LKYAPAFTAPFDLDKLGVTNKFLRNYLEMLAFLLQGLPADQTLTVVMAYMVEDFFRENAV
MDFPKGGSGELMGALARGVTKREGCSVEVSTSVDEVIVENGRAVGVKLAKSGRIIKAKEA
VISNADLYNTYKFVPEGKHEGFDKERIEYLGLTAKPKDGSVPFCKSFMHLHLAVKAELIP
EDAPPQWTVVQDWDKGIDATGNVVVVSVGSKLDQSLAPPGYHVIHAYTAGNESYEDWEQF
EHLMDDAAVRDKDAAYQTFKDERAQPIWDAIQKRAPAVVKGACVIEKVATPLTHARFLNR
HRGNYGLAIAPDNAEGWKFPDVKTPLEGYYRCGDSTTSGIGVPATASSGAVCANAIMSVW
DQLSLNQKIKMP-

##### Targeting:

SignalP 3.0 - NN: Yes, Cleavage Site VFS-FA

SignalP 3.0 - HMM: Yes, Cleavage Site VFS-FA, Signal Peptide Probability = 0.998

SignalP 4.1: Yes, Cleavage Site VFS-FA, D = 0.663 (D-cutoff = 0.500)

ChloroP: Yes, Score = 0.517

#### 18)Thaps3_25361

##### Exon 1:

>Thaps3 chr_18:367805-365028
CTCGCGAAATGAACCTCGTGATCCTCTTCGTCAAACCACCAACGCCGAACACAAGCACAC
CGTATTGACATATCCAATACGCTACGCGCTACCAATAGAGCTATACTACTAGACGTATCT
TGAACCGTCATCATCAAGCAGCTACGCTAACACGAGACACATCGCCAACACACAGGACAC
CATCCATCATGCGTATGGGACGCCCCAACAAAAAGCTCCGCTCCACCTCCAAGCAAACCA
CCAATCCCAATCCACCCAAATACTCCTCGCCCACATTAGTAGTAGGACAGGTGTCCTCCA
ACGTCATCCATTCCATCTATGGGCCGGCGTTGACGAAGCTGGCCGTGGAGAGCGTGGAGG
AGTATGCAGATGCTGTTTTGAGGTGGGAGGCGAGTTTGCCGGAGGTACTTGTCAAGCCAT
CACAGCTTGATGATGCAGAGGATATTGATGTTGATGCCGATGGCACCTTCGAAAAGGGAG
AGGAGGTGGAAGTCGACCTCGACGGAAGCATCCTGCCCAGCCACGACAACGACGACGACA
AGACATCTTCACCCACAATGCCCACCTCCAACCGCCTCTTCACCACCCAATCATCCATCG
ACAACCTCACCGCCCTCCTCACGGACACATCCCAACACTTCTCTACCACCAACGCATGGA
AGATACACGCCAACGCCGCCAAATTCGAACGTCTATTAGATGAAAAATACGGACGTTTCC
GTCCCTTTATTGAATCTCATCCAGAGTTAGAGGTGTTTATCAAAAAGGTTCAAAGGAAGT
ATGCCATGGGACAATTCAGCCCCCTTAGGAAAGGAGAGGGACCGATGAGTACTACCAGTA
GTATTATGCTGTTGTTCATGATGCATAGGAATGGGGTTCGGAAGGAGTTGGTGGCATTAG
TGGCGTTGTTCACTCTGGTGGGATTGGAGCCATGGGCGTTGGTCGGATTGGTGTGTGTTG
GAAAGTATTCCGTGGATCAAAGGAGGAGGAAACGGATCGGTGGTATGCCCAAGAAGGTCA
AGGTTGTGGAATCGTATTATGCACATGGTGTGGTTGGGGAGGAGGAAGAGGAGAGCGAAG

AAGTGGAACGGAGCAAAAAGTATGCTATTTTGGAAAAGCCCGTGGGTACCATCTTCAATC
CGGCGGATTTAAGTTTGAGAGATGAAGAGTACGATGTCATCTTGTTGGGATGTGGTCCGG
AAGTGCTGTACACTGCATCGTTGCTATCACGAGCGGGCAAAAAGACGTTGGTATTGTCAC
CACGAGAGGATGCCTCTGGGTGCTTGACATTGCAGAACGGAAAGACGAATGTTCCTTTTG
ATATTGACGGAAGTAACATTGCCCATTTGGCAAGACAACAGTCGCTGTTGGCTCCGGCAC
TTTGTACAACGACGGATACTCAGGGTGGCATTCGCTTTGCACGCATCGGAAGCGAGGTGG
ATGGCTATGCTCATTCTATTCTCTCGGTGCCAGGCCTGGGCACGGATTCTATCTCGAATG
AGTGTATTCCGATCGTATTGACGGCGGAAGGTGAAGTTGCCTTGGCGGAGTACTGCTCGA
CGTATCTTGGAGATGCATTTCCTGGTACTGATTTGGATGGAAACGATAATGGCAACTCTA
CTTCATTGAGCTATCTCAAGGCATGTGGACAAATCAACGCTGGGTCTGGTGACTTTTACT
TGGCCAAGCTTTTTCCCAAGGCAGCTGAATCGTTCAAATCTTCCGACTCCAATGTCTACC
AACAAGCTTCTATTCGTCCAGCATCGACATTCCTTAACAAGTGCCTACCGCTCAATACTC
ACGTGCGTGCTCTCATGGCGGCCATTGGAATGGCGAATGAGAACTTGAGTCCAGATAAGA
CGAGCATGGCGGCTCATGTTACCAATGTTTGTGCCATGACTAGTACGGAGGGCTATGCGT
ATCCTGTTGGAGGTCCGAGGGCGTTGTGTCATGCATTGACGAGCGTGATAGAACAGAATG
GTGGAAGAGTTGTGAGTGGTGTTTTGTTGCAGGAGTTGCTGTTTGAGAAGTTGGAAAAGA
AGGAACCAAAGGAAGAGACTAAAGATGGCGAGTCAAAGGAGCCAAAGCCTCGTTGCAAGG
GGATAAGATTGGAGAATGGCTTAGAGTTGTCGGTTTCAGACAAGGGAGCTGTCGTTTCGT
TCATGGGAATGATACCCACCTTTTTGCAACTCGTATCTCCTGATGTACGAACTGCCGAGG
GAGTTCCTGCCGGCCTGCCAGCACTAGAGGAACGCCGTCCCTTGATGAGGGTCATGATTA
GTCTCAAAGGAAATAAGGACGACTTGAACTTGACGGGAGCCGATTGGTATCGCTTGCCCA
ATGCCACCTTGCCGAGGGATGAGTTGGATCCAATGACCGGTCAGGTAAAATTCGGAACGA
TTGGTGTAGACGACGATAATACTGGCGCAAGCGAGGAGTTGATACTTGGCGAAGCTACAG
ATGAGACAGAAGCAACGACTAGTCACACACGAGGCAAGCGAAACAAAGCAGCCACGTCGA
AGGCGCCACGATCCAAGTTTACATCTGGAGTATCATGGATGAAAGTATCATTTCCAAGTG
CCAAGGATCCAAGTTGGCAAGATCGACATGGAGACGTTTCCACTTGCGTTGTAACAGTCG
AGGCAGACGACGACTTTGTCCAAATGTTTGATACAAAGCCAAAGATTTACTCGGTGTTGA
AAGCGGGTAATGGCGAGAGAGAACGATTGCGGGACCGAGTGTTGAAAGATTTATTGGAGA
CATTTCCTCAGCTTCAAG

#### Exon 2:

>Thaps3 chr_18:364943-364515
GCCAGTTGGAGACTGTCCAGATATGTGGACCCGTGCGGTCTGGGCTTACTCACAATGGTC
CCAGATTTGCCATCAAAGGAAATCGTCCAGAAACTCCGTACCCCGGTCTGTACATTGGTG
GAGCGGATCTTACTGTGGGTGATTCCTTCTCTGGTGCAATCGTTGGTGGATGGTTGGCTG
CTAATGCGATCATGGGTTACAGTTTCATGGATCATATGTATCTCGGGAAGAACATCACTT
CGGACCTGCAGCAGTTCATAGAGGAACCGATTTTGGCAACTGAAAGGAATGGTGTCATAG
TGGATGACGTTGCTGTTCCTTTCAAGGAGGTTGTTGTTGATATGCAGAAAGGAATCACGG
ATGCAGATAGAAGCACCGCAGCTGAATCTAGTAAAGAGGAGTAATCGTAATCATAGGATG
CTATTGAAT

#### Translated:

MRMGRPNKKLRSTSKQTTNPNPPKYSSPTLVVGQVSSNVIHSIYGPALTKLAVESVEE
YADAVLRWEASLPEVLVKPSQLDDAEDIDVDADGTFEKGEEVEVDLDGSILPSHDNDDDK
TSSPTMPTSNRLFTTQSSIDNLTALLTDTSQHFSTTNAWKIHANAAKFERLLDEKYGRFR

PFIESHPELEVFIKKVQRKYAMGQFSPLRKGEGPMSTTSSIMLLFMMHRNGVRKELVALV
ALFTLVGLEPWALVGLVCVGKYSVDQRRRKRIGGMPKKVKVVESYYAHGVVGEEEEESEE
VERSKKYAILEKPVGTIFNPADLSLRDEEYDVILLGCGPEVLYTASLLSRAGKKTLVLSP
REDASGCLTLQNGKTNVPFDIDGSNIAHLARQQSLLAPALCTTTDTQGGIRFARIGSEVD
GYAHSILSVPGLGTDSISNECIPIVLTAEGEVALAEYCSTYLGDAFPGTDLDGNDNGNST
SLSYLKACGQINAGSGDFYLAKLFPKAAESFKSSDSNVYQQASIRPASTFLNKCLPLNTH
VRALMAAIGMANENLSPDKTSMAAHVTNVCAMTSTEGYAYPVGGPRALCHALTSVIEQNG
GRVVSGVLLQELLFEKLEKKEPKEETKDGESKEPKPRCKGIRLENGLELSVSDKGAVVSF
MGMIPTFLQLVSPDVRTAEGVPAGLPALEERRPLMRVMISLKGNKDDLNLTGADWYRLPN
ATLPRDELDPMTGQVKFGTIGVDDDNTGASEELILGEATDETEATTSHTRGKRNKAATSK
APRSKFTSGVSWMKVSFPSAKDPSWQDRHGDVSTCVVTVEADDDFVQMFDTKPKIYSVLK
AGNGERERLRDRVLKDLLETFPQLQGQLETVQICGPVRSGLTHNGPRFAIKGNRPETPYP
GLYIGGADLTVGDSFSGAIVGGWLAANAIMGYSFMDHMYLGKNITSDLQQFIEEPILATE
RNGVIVDDVAVPFKEVVVDMQKGITDADRSTAAESSKEE-

##### Targeting:

SignalP 3.0 - NN: No

SignalP 3.0 - HMM: No, Signal Peptide Probability = 0.004

SignalP 4.1: No, D = 0.101 (D-cutoff = 0.450)

ChloroP: Yes, Score = 0.543

Analysis using peptide sequences starting with any of the next four in-frame methionines downstream the first one did not result in clear predicted ER or chloroplast targeting.

#### 19)Thaps3_14875

Most of the region was not covered by RNAseq reads. However, there was an open reading frame, that encompassed an approximately 450 base pair region with reads mapped to it.

##### Open reading frame:

>Thaps3 chr_3:2375339-2377698
ATGGACAAAAATAGAGATGCCTTGGCCGACGAGAACGCCGCACCGCGAAAGCATTTGGCA
CTCCATCTCACTCATAACCGTGTTCATCATCACTGGCACACCAACGATGGACAATAGACC
TCTCTCCTGTCTGACGCCAGAGTGCACAATAGATCGCCATCCCAACGCTACTACTTCTTC
GTTTCAATCGTTTGGAGGTGCAGCAACCTGCAGTGGATCCTCCACTGCATCCACGACGAC
TCATCTACTCCTGTCATCCATTCTCATCGGCCTCCTCTCTCCCATCATCTCATGTCTCTT
CGTCATTCTGCTCCTCATGTTCTTCAAACGACGTCAAACCCGTCATGAATTGGAAGGAGG
CAAGTGCAATCTACCCACTGTCGTTTGGAGACCAAGATTTATGAACTACACGTCCAAGGA
TGAAGGATCCGATGAGGACATTGAGATCGATGATTATGAAGCATGGGCAAGAGAGTATGC
ACGTTCACTGCAGTCAGACGACGGCAACAACAGCGGCACATCAACTCATATGAAGAAGTT
GGGATCCTCGGCAATAACCAACATATTACCAAGAATGGAACGCCTCAATGGTCCGTATGG
AATGTACGCCACCGTTTACGGAGTGTCCACGAAGGTATTGCACGTGGCTCATCCAGTTCC
CGCCAGAGCTATTCTGACGGGGAGTGGAGTTGTAGATGTTGGAGGAATGAACAATGGAAT

CGGAGAATGCTTTGAACGACAGAATAGTTCGTTTTTAGGAGAAATATCACAATCAGTGAC
TCGGCCGTTCAAACGATTGTCATCGGGAATGGAGGGTGCTGCTATCAGTCCATCCGAAGA
ACGAAAACAGCGTCGTAGATCTTCTGCACTGCGATTGCTCACCGGCTCAACAAAGTATCC
AGCCTATGACCACTTTAAGAACTTTTCAGGGGATGGAGTCTTCACCGCTGATGGTTCTGA
CTGGAAAGCGAAACGTGCTAGCGTCTTGCACTGTTTATTGAGAAGTGGGGGGGCCGATTG
CATGTTGGAAAAGGAGATTAATAGGGCTGCTGACTCTTTTGAGAGGGAGGTTACGTGGGC
GAAACAAACAATGAATAAGGAGGGCGATGATAAGGATGGTCCAGTGATGAATGTGGTGAC
AATGTTACAAAGGTCGACGATTGGTCTCATTTATCGCATCATTACACACCACAATGTGGA
GTTCAGTCCAGACATTGATACAAACGAGCAATTCATTTGTTCTCCAAAGAGCTCAGCAGC
ATCTCTCACATCCTTGGACAAGAACCAACACAACGGTGCTAAGGCATCAGAAGATGATAA
CCACACCAAACCCGATGTAAAGAAGGACTCACAGATGAAGTTACTTCTACCAATCTACCT
CGATGCAGTCACCAAAATACGAATGATTGTCCTCGCTCAGTCCAGATCTATTTGGTATCT
TCTGCCACGATGGGCCTATCGCACATTCTCTCCCATGTATCGTGACGAAGAAAGAACAAT
GGTTCCGATTAGACAGTTTGCCAGATTGGCGTGTGAGAATGCAGTGGAGGGAAGCCCGTT
GGAATTGCTGAGTCAAAGGAGTAGTCACGCTTCAAAAGAGGGCGAAGCGACCAGTGCAGT
CTCGAAGGATTTGTTGGATGAGGCCATTACTCTCCTATTTGCTGGACAGGATACTTCTGC
TGCCACCTTGTCGTGGACACTGCATCTACTCTCACTTCATCCACAGGAGCAGCAAAAGGT
AGTGGAGGAGGTTCGTTCAGTACTGTCATCTTTGGATGAGGGCGAAATGGTATCCAAGAA
CACCATCTCTCAGCTGCCATATTTGGATGCAGTCATCAAGGAATCGATGAGACTTTATCC
TGTTGCACCATTCATCGTTCGAAAGCTTACCACGGACATGACTATTCCCATCGAAAGTCA
GTCTGTAGAAGATGATGCCACAACAACTACCATCCCCGAATCAACCTTTGCATGCATATG
GATATACGCACTCCAACGAAACCCCAAGCTATGGACAAAACCAGACGAATTCATCCCCGA
ACGATGGATCGATCCTGATCTACGAAGCAACGACCTCGGCCAACAAGAGGTTGGCTCATA
CATGCCATTTGCGCTCGGTCCTCGTAATTGCTTGGGGCAACCAATAGCTCAAGTCATCTT
AAGAGTACTATTGGCGAGGATACTGAACAAGTATGAAGTGAGGGATCCCAAGTTTGATGC
CTTGCAGAGGTTGGGGGAGGAAACGGGGGAGGCATTTGATACCAAGTATCTTCTCAAGGA
TATGCAAGCAGGATTTACTGTTCTTCCTTCAAACGGATTGAGAATCAAGTTAGTGGAGAG
GTGCTAATTGTAGTGGGGTT

#### Translated:

MPWPTRTPHRESIWHSISLITVFIITGTPTMDNRPLSCLTPECTIDRHPNATTSS
FQSFGGAATCSGSSTASTTTHLLLSSILIGLLSPIISCLFVILLLMFFKRRQTRHELEGG
KCNLPTVVWRPRFMNYTSKDEGSDEDIEIDDYEAWAREYARSLQSDDGNNSGTSTHMKKL
GSSAITNILPRMERLNGPYGMYATVYGVSTKVLHVAHPVPARAILTGSGVVDVGGMNNGI
GECFERQNSSFLGEISQSVTRPFKRLSSGMEGAAISPSEERKQRRRSSALRLLTGSTKYP
AYDHFKNFSGDGVFTADGSDWKAKRASVLHCLLRSGGADCMLEKEINRAADSFEREVTWA
KQTMNKEGDDKDGPVMNVVTMLQRSTIGLIYRIITHHNVEFSPDIDTNEQFICSPKSSAA
SLTSLDKNQHNGAKASEDDNHTKPDVKKDSQMKLLLPIYLDAVTKIRMIVLAQSRSIWYL
LPRWAYRTFSPMYRDEERTMVPIRQFARLACENAVEGSPLELLSQRSSHASKEGEATSAV
SKDLLDEAITLLFAGQDTSAATLSWTLHLLSLHPQEQQKVVEEVRSVLSSLDEGEMVSKN
TISQLPYLDAVIKESMRLYPVAPFIVRKLTTDMTIPIESQSVEDDATTTTIPESTFACIW
IYALQRNPKLWTKPDEFIPERWIDPDLRSNDLGQQEVGSYMPFALGPRNCLGQPIAQVIL
RVLLARILNKYEVRDPKFDALQRLGEETGEAFDTKYLLKDMQAGFTVLPSNGLRIKLVER
C-

#### Targeting:

SignalP 3.0 - NN: Yes, Cleavage Site TPT-MD

SignalP 3.0 - HMM: Yes, Cleavage Site TPT-MD, Signal Peptide Probability = 0.147, Signal Anchor Probability = 0.762

SignalP 4.1: Yes, Cleavage Site ITG-TP, D = 0.475 (D-cutoff = 0.450)

ChloroP: No, Score = 0.473

Analysis using peptide sequences starting with any of the next four in-frame methionines downstream the first one did not result in clear predicted ER and chloroplast targeting.

### 20)Thaps3_bd_518

No available RNA-seq data for the unmapped "bottom drawer" sequences.

Gene model predicted by JGI (below) cuts off abruptly on the C-terminal side due to the way the unmapped sequences were assembled. Some C-terminal sequence is likely missing, as evidenced by the lack of an in-frame stop codon. The translated product of the JGI model does not start with a methionine, but can be extended to one upstream. There is only one such methionine that is in frame after the closest stop codon.

#### JGI Model:

>Thaps 3 bd_35x67:17227-17611

AAATGCTATTGGCATTGACTGGGGACTCTCTTTGCAATACAACTT
ACCAATGCTTGCAAAAGTGTGATTACCTTGATGCAGTGGCTCGTGAAACGCTACGCCTTTATCCTCCGGC
TGCAAGCACTCGTTGGGCGACAGATGCAAAGGGTGCGAATGCAGGTGGCTTCAACTTGGAAAAGAGTGTT
GTTCATGTCAACTTCTATGCAATTCAGCGAGATCCTGACGTTTGGGAGAATCCCGTCTCGTTTGTTCCTG
AACGTTTCCTTGGCGAAGAAGGAAGGAAGAGGATACTGTCGTATTCGTTCTTGCCATTCAGTAAAGGATC
ACGCGACTGCATTGGCAAGT

#### Translated:

MLLALTGDSLCNTTYQCLQKCDYLDAVARETLRLYPPAASTRWATDAKGANAGGFNLEKS
VVHVNFYAIQRDPDVWENPVSFVPERFLGEEGRKRILSYSFLPFSKGSRDCIGK
SignalP 3.0 - NN: =No

SignalP 3.0 - HMM:No, Signal Peptide Probability = 0.004

SignalP 4.1: No, D = 0.203 (D-cutoff = 0.450)

ChloroP: No, Score = 0.440

If translated in a different frame, starting downstream the start codon used above, the peptide fragment is not targeted to the ER, and therefore, the chloroplast.

MQWLVKRYAFILRLQALVGRQMQRVRMQVASTWKR

VLFMSTSMQFSEILTFGRIPSRLFLNVSLAKKEGRGYCRIRSCHSVKDHATALAS

#### Targeting:

SignalP 3.0 - NN: Yes, Cleavage Site LVG-RQ

SignalP 3.0 - HMM: No, Signal Peptide Probability = 0.070

SignalP 4.1: No, D = 0.381 (D-cutoff = 0.450)

ChloroP: Yes, Score = 0.531

### 21)Thaps3_6395

No introns.

>Thaps3 chr_6:1467629-1469626
GCTTGATGAGGCTACCATTGGCATTGCTTTCAGCTTGTACTTCGGCTTCATTTCGGAATC
GTCATATCGTGCATAGCTTTACTTTCGGCAGGTCACATAGAGATTGCACAGGAATAATCC
CATCGTTTCGCTTTCGGGTGTCTCCTACAATGTCCTCCTCTGATAGACACAGCTCACAAC
TGAATCAACCCAAACGACCTCGTCACGAGGAACCATCACACAGTATCATGGCATTTGACT
TGAAGAATAGATCTCCGTACGAGAAGAAAATGGAGAGAGTCATTACAGCGTGTCCAAAGT
GTAACGGAGAAGGAAAAGTGCGAGCTCCGTTATCAAAGAAGGCTCGTGCCCAACGCAAAC
GAATGCAACAGAGCCAAACAGGAGATACAACTAATGCACCAAATCTGGCTATTCTGAAGA
AACCGTGTAAGGAGTGTGATGGATCTGGTTTGATTGCCATCAATCCTTTGGATACAACCG
AAAGAAAGCAGACACCACCACAGATTCAACCCAACTTTTCGGTAGCCATTGTAGGCGGTG
GTATTGGTGGCATTGCATTGGCTGCCGCACTACAGCATCGCAACATTCCATGTATTGTTT
ATGAACGAGATTTGTCGTTTGAGGAAAGAAAACAGGGATACGGACTAACGATGCAACAAG
GAGCACGAGCTCTAAGATCCTTGGGCTTCTTTTCATTCTCTGACGATGGAGAGGACGACA
ACAACAATTGTAGTGGCAAAAAAGCAGTGGATGAGAATACTTCAAATACAAAGCAAAAGT
TTGGAATCCACTCAACTCGTCACGTAGTTCACAAGCCAGATGGAACTGTAGTAGGTGAAT
GGGGTATGAAAGTCTGGGGTGGTCGATTCGAGAAGAACGGCAGGAAGCACGCCAAGCGAC
AAAATGCACACATCTCTAGGCAAAATCTTCGCCAGCTGTTGATGGAGATGCTGCATCCTG
GTACAATACAATGGGGGCAAAAGTTTGTGGGTTATTCGGGACAGTCTAGTGATGACGATT
CCTCACAGGATCAACCATCATTGCAAGTCAGATTTCGACGCAGAAGTAACGATTGTGATG
AGGAGGTCGCTACAACTGCGTCTGTACTTGTCGGATGCGATGGTATCCGATCGTCTGTAC
GATCTGCAAAGTTGGGTGAGGACGGAACACCACTCCGTTACCTGGATTGCATTGTCATTC
TTGGCATTGCTCCGTCGCCAACCTCGGCGTTAACTGATGGTGAAACTGTATTTCAGACGG
CAGATGGCATAACTCGTCTGTATGTCATGCCGTTTGCGGAAGCTGGAGATGACTCGTCTG
GTTTATCAACTGACAACACTAAAGGATTGAGCATGTGGCAGCTCTCGTTTCCGATGGACG
AGACTGATGCAACAAGGCTGAGTCAACTTGGATCGTCTGCGTTGAAAGAAGAGGCCCTCA
AACGATGTGGTGCATGGCATGATCCAATATTAAAGCTGTTACGTTCCACACCAGAGGATT
TCATTACTGGATATCCGTGTTATGATCGTGCCCTTGTCGAGAGAAAAGAGCTTCGAGATG
GATGTGATAAATCTCAATCTGCAAACGCCTTTGTGACTCTACTCGGCGATGCTTGTCATC
CTATGTCCCCCTTCAAAGGTCAAGGGGCGAATCAAGCTCTTTTGGATGCCGTGCTATTAA
GCCAAAAGCTCTTCGATATATCTCGTATTCATAACGGGAAAACGAACGTCAACGAACAGC
AACCTACCATATCACTCAATGAAAGCACACCACAGGCATTGGCAGAGTTCGAAAACGACA

TGCTACAAAGGTGTGAAGTCAAAGTTAAAAAGTCGGCAGATGCAGCAAAGTTCTTGCATA
GTGACGTCGCTATTCAAGAGGGAAACATTACACGAGGAGCGGCAGCGTTGGATGCGAAGG
GATGAGAGCGAGATTCTTCTGGGATCTAGTTGGATACAATCATAAGATAACCTAAGTTAC
CCCTGTAATGAGCGAACA

##### Translated:

MRLPLALLSACTSASFRNRHIVHSFTFGRSHRDCTGIIPSFRFRVSPTMSSSDRHSSQL
NQPKRPRHEEPSHSIMAFDLKNRSPYEKKMERVITACPKCNGEGKVRAPLSKKARAQRKR
MQQSQTGDTTNAPNLAILKKPCKECDGSGLIAINPLDTTERKQTPPQIQPNFSVAIVGGG
IGGIALAAALQHRNIPCIVYERDLSFEERKQGYGLTMQQGARALRSLGFFSFSDDGEDDN
NNCSGKKAVDENTSNTKQKFGIHSTRHVVHKPDGTVVGEWGMKVWGGRFEKNGRKHAKRQ
NAHISRQNLRQLLMEMLHPGTIQWGQKFVGYSGQSSDDDSSQDQPSLQVRFRRRSNDCDE
EVATTASVLVGCDGIRSSVRSAKLGEDGTPLRYLDCIVILGIAPSPTSALTDGETVFQTA
DGITRLYVMPFAEAGDDSSGLSTDNTKGLSMWQLSFPMDETDATRLSQLGSSALKEEALK
RCGAWHDPILKLLRSTPEDFITGYPCYDRALVERKELRDGCDKSQSANAFVTLLGDACHP
MSPFKGQGANQALLDAVLLSQKLFDISRIHNGKTNVNEQQPTISLNESTPQALAEFENDM
LQRCEVKVKKSADAAKFLHSDVAIQEGNITRGAAALDAKG-

##### Targeting:

SignalP 3.0 - NN: No

SignalP 3.0 - HMM: Yes, Cleavage Site VHS-FT, Signal Peptide Probability = 0.974

SignalP 4.1: No, D = 0.207 (D-cutoff = 0.450)

ChloroP: Yes, Score = 0.554

Analysis using peptide sequences starting with either of the next two in-frame methionines downstream the first one did not result in a more clear targeting prediction.

#### 22)Thaps3_10233

##### Exon 1:

>Thaps3 chr_15:679062-677604
CGTTGACTCTCTTTCAACGACGACACCCACCAAGTCATCCATCTTAAACTTGTTGGTAGT
ACGCCACTCTTGTTGAAAAGTAGATCCAACGACCATGATCGGACGCAAGTACAGTCTCGC
AGCCTCAGCTCTGGCAATTATCGCCTCCCTCACCTCCACAACCGCATTTGCACCACCCTC
CTCCTCTCTCCTACGCTCCACTCGCCTCCACTCCACCGTCGAAGAAACCACCAACGGAGA
AGCCGCAACAAACACCAATGTCGAACAAATCAAAGACACCTCACGTGACAAAGTAATGAC
ATTTTCCTACGACATGTCCATTGAACCCAAGTACGAAAAGCCAACGTATCCAGGAACAGG
CAACGGCATGTCAGGCGACTCTGGTGAATACGACATTATCGTTATTGGATCAGGAATGGG
CGGACTCGCCTGTTCGGCACTCTCTGCCAAGTACGGCTCACGAGTCTTATGTTTGGAAAG
TCACATTAAAGTGGGAGGAAGTGCTCATACCTTTAGCCGAATGCACAATGGGGGCAAGTA
CAGTTTCGAAGTCGGACCGAGTATCTTTGAGGGATTGGATCGTCCATCGTTGAATCCACT
GAGAATGATCTTTGACATTTTGGAGGAAACCATGCCGGTCAAAACGTACAAGGGATTGGG
ATATTGGACTCCTTCTGGCTACTGGCGTTTCCCCATTGGATCACGTGAGGGATTTGAACA

GTTGTTGATGGAACAGTGTGGTGAGGATGGAGAAAAGGCTATTGGAGAGTGGAAGGCGTT
GAGAGAGCGATTGAGGACTTTGGGTGGAAGTACGCAGGCCGTGGCATTGTTGAATTTGAG
GCAAGATGCGGGATTTTTGGCAACTACTGCCGGTTCATTGCCGTTTGTGGTGACTCATCC
TGATGTGTTTGGGGATTTGTCATTGACGTTTGATGATTTGAGCAAGACGGTCGATGAGTT
TGTGACTGTGCCTTTCTTGAGGAATTTTATTGATACGATGTGCATCTTTTGTGGATTCCC
TGCAAAGGGAGCCATGACGGCACACTTGTTGTACATTCTCGAGAGGTTCTTTGAGGAGAC
TGCAGCTTTCTCTGTTCCTATTGGTGGAACATGCGAGCTTGGAAACACTCTTCAGCGTGG
ATTGGAGAAGTATGGAGGTAAATTGCAACTCAATGCTCATGTCGATGAAATTCTCGTAGA
GAACGGACGTGCCGTCGGAGTCCGTCTCATGAATGGAAATGTCGTCAAGGCACGCAAGGC
AGTGGTCAGTAATGCTACTCCCTTTGATACTGTCAAGTTGATGCCCAAGGCGGAGGGTGA
GCCCAAGGGATTGACTAAGTGGAGAGAGGAGTTGGGAAAGCTTCCTAGGCATGGAGCTAT
CAGTCATTTGTTCTTGGCGATTGATGCGGAGGGCTTGGACTTGAGTCATATTCAGGATCC
TGCTCATTTGGTAGTTCAG

#### Exon 2:

>Thaps3 chr_15:677527-676741
GATTGGGATCGTTCTCTTCAAGACTCCCAAAATCTTTGCTCCTTCTTCATCCCATCCATC
CTCGACAAAACGCTATGCCCAGAAGGCAAGCACGTCATTCATGTCTACTCTTCTGGTGGA
GAACCTTACGAACCATGGGAAAAGCTCACACCTGGCTCAGAAGAATACGAAGCATACAAG
AACGAACGTGCCGAGGTTCTCTGGCGTGCTGTGGAGCGTTGCATTCCCGATGTTCGTGAT
CGTGTGGAGTTTTCTATCGTTGGTTCACCTTTGGCTCACGAGGCTTTCCTCCGCAGAGAT
AGGGGAACATACGGTATGGCTTGGGCTGCTGGTTCGTCGGCTCCTCAATCTGGTATCCTT
GGAAGTGTTCTTCCTTTCCCATTCCCCAACTTGAAGACTCCAGTGGACGGTTTGTTGCGT
TGTGGTGACTCATGCTTCCCTGGTATTGGAACTCCGTCGGCTGCTGCTAGTGGTGCCATT
GCTGCCAATACAATGACTCACGTTGATAACCATTTGAAGATGCTGTCGGAGGCGAGTAAG
TTGGATCCTATGTACAAATTCTTGGATGCTGGTATCATGGGGCAGGTTTACAAACCCTTG
GTGCAGGGATTCACTCCTAGTCCAGAGTTGAGGACCGACCAATACGTTTCGGGTGCTGGG
GTTGCACCTGTCGATTACACTGCCACTGATCCTAGTGTCAGCGAGAGGATTGATTTGTAA
GAGTGGTAAAATAGTGGGGGTAAATATAGCGTAGTCAGAGTTATTTGTCACGTGAAGAGA
GCAATTG

#### Translated:

MIGRKYSLAASALAIIASLTSTTAFAPPS
SSLLRSTRLHSTVEETTNGEAATNTNVEQIKDTSRDKVMTFSYDMSIEPKYEKPTYPGTG
NGMSGDSGEYDIIVIGSGMGGLACSALSAKYGSRVLCLESHIKVGGSAHTFSRMHNGGKY
SFEVGPSIFEGLDRPSLNPLRMIFDILEETMPVKTYKGLGYWTPSGYWRFPIGSREGFEQ
LLMEQCGEDGEKAIGEWKALRERLRTLGGSTQAVALLNLRQDAGFLATTAGSLPFVVTHP
DVFGDLSLTFDDLSKTVDEFVTVPFLRNFIDTMCIFCGFPAKGAMTAHLLYILERFFEET
AAFSVPIGGTCELGNTLQRGLEKYGGKLQLNAHVDEILVENGRAVGVRLMNGNVVKARKA
VVSNATPFDTVKLMPKAEGEPKGLTKWREELGKLPRHGAISHLFLAIDAEGLDLSHIQDP
AHLVVQDWDRSLQDSQNLCSFFIPSILDKTLCPEGKHVIHVYSSGGEPYEPWEKLTPGSE
EYEAYKNERAEVLWRAVERCIPDVRDRVEFSIVGSPLAHEAFLRRDRGTYGMAWAAGSSA
PQSGILGSVLPFPFPNLKTPVDGLLRCGDSCFPGIGTPSAAASGAIAANTMTHVDNHLKM
LSEASKLDPMYKFLDAGIMGQVYKPLVQGFTPSPELRTDQYVSGAGVAPVDYTATDPSVS
ERIDL-

#### Targeting:

SignalP 3.0 - NN: Yes, Cleavage Site TTA-FA

SignalP 3.0 - HMM: Yes, Cleavage Site TTA-FA, Signal Peptide Probability = 0.998

SignalP 4.1: No, D = 0.619 (D-cutoff = 0.450)

ChloroP: Yes, Score = 0.560

### 23)Thaps3_33926

#### Exon 1:

>Thaps3 chr_4:1242467-1242109
TTCGCATTTTGCTCCTCGTCCTCCTCCTCACAGCTCTCATCCATCCAACACCACACCACA
CTACCGCAATGACAATCCCCGGCACCACCACAAGCGGTATGCCCTTCGGCTTCGGCTCCT
CCACAGGACCATCCGACAACCTCCTAATCGGCCTAACATGCCTCGCCATCTTCTCCCTCT
TTTACGCCTTCTTCGTCGTCCGCCCTAGGACGAAGGGAGGACCACATGCACCACCGGTGG
TTACGTCGAGCCCTGTCTCAAGTCTTCCTGTCGTTGGGACTATTGTGGAGTTTGGCAAGA
GTCCTGTGAAGATGGTTCAGAGGTGTTATGAGGATTACGGTCCTGTGTTTACTGTGCCG

#### Exon 2:

>Thaps3 chr_4:1241947-1240596
TTCTTCCACAAACGTCTCACCTTCCTCATCGGCCCCGAAGCCCAAGAACCATTCTTCAAA
GCACCCGACGAAGTCCTCTCCCAAAACGAAGTCTACGGCTTTATGAAACCCGTCTTTGGA
CCTGGAATCGTCTACGATGCCTCCAAAAAGAACCGTCAAGTTCAATTCCAATCAATGGCT
AACGGCCTTCGCACTGCTCGTCTTAAGGGATACACTGCCAAGATTGAACGTGAGACGCGT
CAGTACCTCGAATCTTGGGGAGAGTCCGGAGAGCTTGATCTATTCCATGCTCTTTCGGAG
TTGACTATTCTTACTGCCTCTCGTTGTCTTCACGGAGATGATGTTCGTGAAAATCTCTTC
AAGGAGGTTTCGGAATTGTACCACGATCTTGACCAGGGCTTGACTCCACTCACCGTGTTC
TTCCCCAATGCTCCTACAAAGTCTCACATGAAACGCAATGCGGCACGTGCCAAAATGGTG
GAGTTGTTCTCCAAAGTGATTAAGAATCGTAGGGATAATCCTGATGTGCAACACTCGGAT
GGTACGGATATCCTCTCCATCTTCATGGATGTCAAGTACAAAGATGGATCAAACATTACC
GACGAGCAAGTGACTGGGCTTTTGATCGCATTGTTGTTCGCTGGGCAGCATACGAGTTGC
ATTACTTCTACATGGACGAGTCTCTTTATACTCAACAACCCTGCCATTCTCAAGCGTATT
ATTGCTGAGCAGAATGACGTCTTTGGTTCTCAACCGGATGCCGATGTGGATTACAAGATG
GTGAACGAGGATATGCCCTTGTTGCACAACTCGATGAAGGAGGCTTTGCGTTTGTGCCCT
CCGTTGATTCTTCTCATCCGTTATGCTCTCAAAGACGTGAAGGTGAAAGCTGCCGGAAAG
GACTACACCATTCCTAAGGGCGATATGGTGCTCATTAGTCCATCTGTTGGTATGAGGATT
CCCGAGGTGTTTAAGGAACCTAATACCTTTGATCCTGATCGTTTCGGTCCTGATAGGGAG
GAGGACAAGTCAAGTCCATTCGCTTACATGGGCTTTGGAGGAGGTATGCACAGTTGCATG
GGACAGAACTTTGCGTTTGTTCAGGTCAAGACGATTCTTAGCGTGTTGTTCCGTGAGTTT
GAGTTGGAGATGGTTTCGGAGACGATGCCCGACATTGATTATGAAGCCATGGTTGTTGGA
CCCAAGGGAGATTGCCGTGTTAGGTACAAGAGGCGTCAGTAGATGATGGGTACCATGTCG

AGTTGCAACTGTCTTGATCAAACATAGATGTTTTGAGAAACTGGGGTTGATTTCATCGTA
TTGATTTTAATTGAATAAAGAAGCTAGAGATT

##### Translated:

MTIPGTTTSGMPFGFGSSTGPSDNLLIGLTCLAIFSLF
YAFFVVRPRTKGGPHAPPVVTSSPVSSLPVVGTIVEFGKSPVKMVQRCYEDYGPVFTVPF
FHKRLTFLIGPEAQEPFFKAPDEVLSQNEVYGFMKPVFGPGIVYDASKKNRQVQFQSMAN
GLRTARLKGYTAKIERETRQYLESWGESGELDLFHALSELTILTASRCLHGDDVRENLFK
EVSELYHDLDQGLTPLTVFFPNAPTKSHMKRNAARAKMVELFSKVIKNRRDNPDVQHSDG
TDILSIFMDVKYKDGSNITDEQVTGLLIALLFAGQHTSCITSTWTSLFILNNPAILKRII
AEQNDVFGSQPDADVDYKMVNEDMPLLHNSMKEALRLCPPLILLIRYALKDVKVKAAGKD
YTIPKGDMVLISPSVGMRIPEVFKEPNTFDPDRFGPDREEDKSSPFAYMGFGGGMHSCMG
QNFAFVQVKTILSVLFREFELEMVSETMPDIDYEAMVVGPKGDCRVRYKRRQ-

##### Targeting:

SignalP 3.0 - NN: No

SignalP 3.0 - HMM: Signal Anchor Predicted, probability = 0.936

SignalP 4.1: No, D = 0.136 (D-cutoff = 0.500)

ChloroP: Yes, Score = 0.543

Analysis using peptide sequences starting with either of the next two in-frame methionines downstream the first one did not result in a more clear targeting prediction.

#### 24)Thaps3_1549

No introns.

>Thaps3 chr_1:1808544-1810261
CTCGTCGACATCGAAAGACGGCAAACAGAAACAGCAACACAACGGAACAGCAGATTGATA
CATGGTGCATGTTTGATACCATGACAGCGCCATCGTCACCCTCCTCCTCTCGCTCTGCCC
TCGTCACGGCGGCAGCAGCATCCCTCGTCTCTCTCTCCTTGCTCTCCATCTACAAGCGTC
GCCGTACTACGTCCAACAATGAACTTCCCTACCCACCCACCCCTCCCGACAGGAACTACT
TCCTCGGCCATGCAATGTCTCTCCGACGAGTTCCCGGAGAACCAAAGAAATCGCACGATC
TTCTCTTCTTGAACTGGATGAACAAACTCAACAGCAAAGTAGTAATGTTTGAGCTCCCAT
TTCTTGGACGGCTGTTCGGTCTCGGGCGTATGATTTGCGTAGGTGATGCCGAGATTGCTC
GGCACATACTCGTTACGGCGAACTACAACAAGTCTCCCACCTACAGTGTGTTACAGCCAC
TCATCGGCATGAGTTCCATGGTTGCCACGGAGGGAAAGATGTGGAAGGATCAAAGAAAGT
TGTACAATCCTGGATTTTCTCCAGAGTTTCTTCGCAATTGTGTATCGACAATTATTGAGA
AGTGTAATAGATTCATCGCCAGATGTGATGGTGATGTTGAGAATGGTGTAGCGACGGATA
TGTTGGCGAGATCCATTGACCTCACTTCTGATGTGATTGTGCAGGTAGCATTTGGAGAGG
ACTGGGGAGTTGATAGCAAGGACAAACATGGTATCGAGACACTGCAAACAATACGAGATC
TTACGGTAGCCGTTGGGGAAAATATGACCAACCCATTACGCAAATACTTTGGATTACGAA
GCATTTGGAGAACGAGGAGACTCTCGGCAGCTCTGGATCAGGATATGCAAAATCTTGTGA
AGAGAAGACTTGCTCAGGTGTTGGCTGGAGATGCTGATTTAGAGAAGGATATCTTATCAT

TGACGTTGTCTGGTGTTTTGGAAGCAAAACAGGAATCTAAGTCAGGCGCCATCTCGTTGA
GCAAGGACGAAATGGAAAGGATGACATCGCAACTTAAGACTTTTTACTTCGCTGGGCACG
ATACCTCTTCATCCGCTATCGCATGGGCGTATTGGCTGTTGACAAAACATCCAGAATCAC
TCCAACGAGCTAGAGAGGAAGTTGTATCACACCTCGGAAGAGATTGGTCGGACGAGGCAT
TGACTGGGGACTCTCTTTGCAATACAACTTACCAATGCTTGCAAAAGTGTGAGTACCTTG
ATGCAGTGGCTCGTGAAACGCTACGCCTTTATCCTCCGGCTGCAAGCACTCGTTGGGCGA
CAGATGCAAAGGGTGCGAATGCAGGTGGCTTCAACTTGGAAAAGAGTGTTGTTCATGTCA
ACTTCTATGCAATTCAGCGAGATCCTGACGTTTGGGAGAATCCCGACTCGTTTGTTCCTG
AACGTTTCCTTGGCGAAGAAGGAAGGAAGAGGATACTGTCGTATTCGTTCTTGCCATTCA
GTAAAGGATCACGCGACTGCATTGGCAAGTACTTTGCTCTTCTTGAAATAAAGATTGCAT
TGGCTGCTTTGATTTCTCGGTACGATGCATCAGTTGTGAATGAAAATGAGCAGTATGTTA
TCCGTTTAACATCTGTTCCTCACGACGGATGCAAAGTGAATCTCTCTCGTCGCAGGAAAT
AAACGTGCTGTAAAGAAATCCTAAACTAACTTAGTATT

##### Translated:

MFDTMTAPSSPSSSRSALVTAAAASLVSLSLLSIYKRR
RTTSNNELPYPPTPPDRNYFLGHA MSLRRVPGEPKKSHDLLFLNWMNKLNSKVVMFELPF
LGRLFGLGRMICVGDAEIARHILVTANYNKSPTYSVLQPLIGMSSMVATEGKMWKDQRKL
YNPGFSPEFLRNCVSTIIEKCNRFIARCDGDVENGVATDMLARSIDLTSDVIVQVAFGED
WGVDSKDKHGIETLQTIRDLTVAVGENMTNPLRKYFGLRSIWRTRRLSAALDQDMQNLVK
RRLAQVLAGDADLEKDILSLTLSGVLEAKQESKSGAISLSKDEMERMTSQLKTFYFAGHD
TSSSAIAWAYWLLTKHPESLQRAREEVVSHLGRDWSDEALTGDSLCNTTYQCLQKCEYLD
AVARETLRLYPPAASTRWATDAKGANAGGFNLEKSVVHVNFYAIQRDPDVWENPDSFVPE
RFLGEEGRKRILSYSFLPFSKGSRDCIGKYFALLEIKIALAALISRYDASVVNENEQYVI
RLTSVPHDGCKVNLSRRRK-

##### Targeting:

SignalP 3.0 - NN: No

SignalP 3.0 - HMM: Signal Anchor Predicted, probability = 0.936

SignalP 4.1: No, D = 0.136 (D-cutoff = 0.500)

ChloroP: Yes, Score = 0.543

Analysis using peptide sequences starting with either of the next two in-frame methionines downstream the first one did not result in a more clear targeting prediction.

#### 25)Thaps3_264647

##### Exon 1:

>Thaps3 chr_19c_29:130307-130912
CAAACATCACTATTTTTGGCAAGGGAGGATGTCTTCATTCTGAGTAGTCTCCAATATATT
TCGCTACAAATCCGGTCATACTCTCCTTATAATTTTTGATATGCCTTCTTCAAGGTTCGA
TGCTGCTCGATTTGTATCTTTGGTTGTTCACACAGTCAGCAGTGAGAAGTGAGGAGCTCT
CTCTGCATTTGGATATATAAAACTGATAGAAGAGATCAGCCAACACACATAACAATTTGG

TGGAGGAGCGAGAGATTTCATTGCACCATCGAGAGAGCGGCGAGGCAGGAGGCGACGGTG
CATCCATACTTGATCCATCGCATCGGACGGTCAGATTTGCAAATCGGCAATCGTACCGAC
ACATCAACTGCTCCCACTTCCCAGTGTTCCTTTCATCCCTGATCAGTCCTTCTTCCGCAG
CAGCGATAGTATGATCAGTAATATGACGATGGCCCTTTCAACACTGGCATTGTATCTTAC
ACCAACCACAATTATCACCCTTCTGCTATGTCTCTTAGTGACTCGGTTCATCCAATGGAA
GAATCACCTTGCGAATATGAAGAGCCAGACTCCCTTTCCTGGAATTCCAGTAGTTCCCGA
TGCTCA

#### Exon 2:

>Thaps3 chr_19c_29:131012-133019
CTGGCTACTCGGCCACTATCCCTTGTTCGTCAACCCCGACAAACACCACCAAACCTTCAC
CGCCCACGCCACCCCCTCCGGCATATCCGCCCTCTGGGGTCCGTCCACTGATAAATTCTT
CTCCTCCGTCCGAGCCGATCACTGCCGTTCCATCCTTCGTCAAAGCTCCAGTAGAAACTT
TGTCTCGTTCATTGTACGGCATGGACGGAGGACATTGGGAGAGGAGAGTATCATTTTGAT
TAACGGAGGAAAGAGGTGGAAGAGGCAACGAAAGGTGATTCAGAAGGCGTTTCATTTGGA
GGTTGTGAAAGGTAGGAGGGAGGCTGTGGGGGAGGTGGCGGATGTTGTTGTGGATTGGAT
ACTGAGGGCTTGTAGTGGTAGGAGTGATAGCGGTGTGCATGATGGAGATTGTGGAGGGAG
ATTGGTTGGTGCGGATGGGAATGAGAAGGTTTGTGTGGAGGCTGAAGACTTCTTTAAGTT
GTTTGCGTTGGAGGTGTTCGGAAAGGTAGCAATGGGATATGATTTTCGGTGCTTTCCTTC
TCTTGCTACTTCGGACGACAACGGCAACAGCAACAGTAATGTTGCCTTACACAACAAACA
AGATGTGGTGAGCAATGGCACCCATACATACAACGACAACGCGTGCAACTGTCTCCAAAT
GCCACCCGATGCACAGTCCTTTGACTTTCTGAATGTCGACATCGGGAACCGCTCAACACC
AACGAGCCTGATGAATCCGTGCATGCAATTCTACTCTATACCCACTCCACACAACAAAAA
GTATCATCATCATATGGACAGAATTAAGGGACTGGTTGGTAAGATTATCGGACTGCAACT
GAACAGGTTGTGCAGTGACGGCGGTGTCATGGAAGGCGATACGAACATGATTACCCACTT
GTTACAATCTACAATCGAAGAGAACTTCAGTCCTACCGATACAGATGGGAACACTGGCTG
TCCATTCTCATCATCCTTACCATCAAAGTCTATTCCAGATACAGTCATCTCCAACCTCAC
TCCATCAGATAAAGATCAAATCATTGAAAGCGTCTCCAAGATGCTCATCACATTCCTCAT
GGCAGGCTACGAAACCACTGCAATCTCAATGTCGTTTGTAGTGTACTTCCTTTCCAAATA
CAAACGATGCCAAGAAAGATGTGCAGAGGAAGCGAGAAGAGTTCTGGGGCGCTGTGGAGT
GCATGGAACGGACATTGACGATGACGAGCTGGTATACTGCCGTGCTGTCTTCATGGAAAC
GATACGGTTGCATCTGCCAGTCATGTTCACAACTCGTGTGACAGAGAAGGAAATGTCCTT
TGATACAGGGCTGGAGGAAGGCCATAATGTGACAATACCAAAGGGAACGAGGTGTGTCGT
TTGTCCTACAGTAGTTCATATGGATGAGCGTAATTTTGAACGAGCCGAGGAGTTCTTACC
TGAGAGATGGGTGCGGTGGGAGAGGGGCAGGTGGGTTGAACGAGATTACGAAACTGAAGG
ATTGAAGTCAACAGCATTGCCATCAATTACTGAAGATGAACAAGATTCTCCTCCCATATC
TGCAAAGTACGACGAAGAGAATAATTCTGCCAGTTCAATCTCTGCGGCTGATCCACACAA
TTTCTTCTCGTTCTCAGATGGAGCAAGGAATTGTGTTGGGAAACGTCTTGCAATTATGGA
GTCTACAATCTTGATTGCGGTATTGCTTCGTGACGTGTGCGTCGACTTCGCAGAGGAGGG
ATTTGAGATGAAAAAGGTACGGCGGTTCGTCACGTGTGGCCCTGAGAGTTTACCCGTCGT
GTTTTGGAGGAGGGAGTGAAGAAAGCTTGTAGTAGGTCTTAGTTTTTTTGCAGAAGAACC
GTAGCGAAATGCGGCTGTAACCAAGCTTCTTTAAGTAGACACACTATTGTGTGTAAAGTC
GTGATCATGTATTCCATGAAGCCGATCTAATGACTGGAGATGCTGTTCAGTTAAGATGCA
CTTACATGTGAAACTAACAGTTGAGGAC

#### Translated:

MISNMTMALSTLALYLTPTTIITLLLCLLVTRFIQWK
NHLAN MKSQTPFPGIPVVPDAHWLLGHYPLFVNPDKHHQTFTAHATPSGISALWGPSTDK
FFSSVRADHCRSILRQSSSRNFVSFIVRHGRRTLGEESIILINGGKRWKRQRKVIQKAFH
LEVVKGRREAVGEVADVVVDWILRACSGRSDSGVHDGDCGGRLVGADGNEKVCVEAEDFF
KLFALEVFGKVAMGYDFRCFPSLATSDDNGNSNSNVALHNKQDVVSNGTHTYNDNACNCL
QMPPDAQSFDFLNVDIGNRSTPTSLMNPCMQFYSIPTPHNKKYHHHMDRIKGLVGKIIGL
QLNRLCSDGGVMEGDTNMITHLLQSTIEENFSPTDTDGNTGCPFSSSLPSKSIPDTVISN
LTPSDKDQIIESVSKMLITFLMAGYETTAISMSFVVYFLSKYKRCQERCAEEARRVLGRC
GVHGTDIDDDELVYCRAVFMETIRLHLPVMFTTRVTEKEMSFDTGLEEGHNVTIPKGTRC
VVCPTVVHMDERNFERAEEFLPERWVRWERGRWVERDYETEGLKSTALPSITEDEQDSPP
ISAKYDEENNSASSISAADPHNFFSFSDGARNCVGKRLAIMESTILIAVLLRDVCVDFAE
EGFEMKKVRRFVTCGPESLPVVFWRRE-

#### Targeting:

SignalP 3.0 - NN: Unclear

SignalP 3.0 - HMM: Signal Anchor Predicted, probability = 0.894

SignalP 4.1: No, D = 0.340 (D-cutoff = 0.500)

ChloroP: No, Score = 0.455

Analysis using peptide sequences starting with any of the next four in-frame methionines downstream the first one did not result in a chloroplast targeting prediction.

### 26)Thaps3_4026

No introns.

>Thaps3 chr_3:2380128-2379629
TCAACAAAATCATGTCCAAGAACGCAACTACGCCCTCTTCACCTCCCGCAGCCGCAGGCA
TGGATGGAATCCTAGCCGAACGAATGGCCGAGGGAGGTGGAGCACTCTGCAACGATGAAA
ACGGATTGTGTCTAGGTTCACGTGGGGATATTGATACTAGTCAGAGTGGTTGTTATACGG
CAATTGCAAAACTTGCGAGTCAGTTGGATAACACGGTGGATGGACAGGGCACTAAGAATA
ACAACAACAACGGGGAAATGCCATTGGTGACCATTCAGACTGAAAAGGCGGCACTGTTGG
TGAAGGAATATGGGGGAAGAACGGTGGTGTTTCGTGTGCCGACAGAGGTGAATGTGAACG
GACAACAATGTGGGAGTGAACTGGTGATGAACGGGGAGCTCGACAATCTGAACGAAGGAG
TTTGATGGGGATGCATCCCGAAACAACAATCCCACTCGGTGTATATTAGGAACGGTATAT
AATACAAGAGGTTACAACAG

#### Translated:

MSKNATTPSSPPAAAGMDGILAERMAEGGGALCNDENGLCLGSRGDIDTSQSGCYTA
IAKLASQLDNTVDGQGTKNNNNNGEMPLVTIQTEKAALLVKEYGGRTVVFRVPTEVNVNG
QQCGSELVMNGELDNLNEGV-

##### Targeting:

SignalP 3.0 - NN: No

SignalP 3.0 - HMM: No, probability = 0.011

SignalP 4.1: No, D = 0.139 (D-cutoff = 0.450)

ChloroP: No, Score = 0.447

Analysis using peptide sequences starting with either of the next two in-frame methionines downstream the first one did not result in a more clear targeting prediction.

#### 27)Thaps3_5221

##### Exon 1:

>Thaps3 chr_5:652777-650770
TTTCATCCTCCTTCTACACTTCATCAACGTCCGCTAGACTGCCGCCGCTGGCAACACATT
TGTCCCTCAACTCCTGACACAGCCGAAAGAGAGACTCTCGCCATGAAGCTGTTGTTTGTA
GCGTCGACGTTGATCGGCGTACTCTCGTTCACGCCCCCTCAAGTTCTGACCGACACACGT
CATCGGCCACCAGCTCTGTGTAACTCAGGCGATGCAACAAACGATGGTGGTGATGAAGTG
CACGAAGTCGATATTGCTGTCGTTGGAGCGGGGATAGGTGGTCTCTGTGCCGGCGCCATA
CTCAACACACTTTACGACAAGAAGGTTGGGGTGTATGAATCTCACTATTTAGCCGGAGGA
TGTGCACACAGTTTCAGCCGTAGTGTAAAAATTGGAGACGATGAACAGCCAACAACGTTC
ACATTTGACTCTGGGCCTACCATAGTATTGGGATGCAGCAAAGAACCGTACAATCCTCTG
CAACAAGTACTACGTGCAGTGGGGGTAGATGATCAAATAGAGTGGCTTCCTTACGACGGG
TGGGGAATGATCGAGCATCCAATGCAACCGAAGGAAAAGAGATGGAAGTTCAAAGTTGGA
CCGAATCACTTTGAGGACGGTCCTCTTCAAGTGTTTGCATCAAATCTTAATGCTCTTGAG
GAGTTCAATCAATTGAGAGAAATTACAAAGCCTCTTGTCACAGGAGCCGCTACCATTCCA
GCCATGGCCATGAGACCAGGACAATCAGCTCTAGTTCCGTTGTTGAGATATCTTCCATCA
TTGATCTCAATCATTAGCAATGGAGTTGAAGCATCGACTGGACCTTTTGCTCCCTACATG
AACGGCCCAATATTTACTGTAAAAGATCCGTGGCTACGAAGCTGGTTGAATGCGTTGGCG
TTCAGTTTGAGTGGTCTCCCCGCAGATCGTACCAGTGCTGGTGCAATGGCGTATGTGCTA
TTTGATATGCACAGAGAAGGGGCAGCGCTAGATTATCCCCGGGGAGGACTTGGAGAAGTA
GTCAAAGCATTGGTCAACGGCGTGGAGCAAAAGAGTATTGGATCAAAAGTACATCTTAGC
AGACACGTAGAAAGTATTGATACCAACGAAGAAGGAGATAGAGTCATTGGATTGACTGTT
CGTAAGAATGGAGGGAAGAAGGTCATCGTCAAGGCCAAAGAAGGTGTCGTGTGTAACGTG
CCGATGTGGTCACTCCGAAAGTTGATCAAGAATAGGAATGCACTGAGTGTTCTGGGTGGA
GACAAGGCAACTTCTTCATCAAGTGGCTTGAAAGCAAAACAATCTTGGATGACGTCTTTT
GATACAGACCCAAGTACTGGAAGAGGAAGCGTACTTCGTCCAAAACCAGCTGAGGACACG
ACAATAGAAAAAAGTCTCTTAGAGAAGTGTGACTCTGCAGAAATGACTGGCTCATTTCTT
CACCTGCATCTCGCTCTCAATGCTACTGGACTTGATCTTCAGTCTCTTGAGCCTCACTAC
ACTGTCATGGATCGTGGTTTGGAAGGCGATGGGAAAGTTATTGATGGGGTTAAGGATGAT
TCAAGCGGCGAGCTGAATATGATTGCTGTATCTAATCCTTGTGTGTTGGACAATACTTTG
GCACCAGAGGGATTTATCATCATGCATGCCTATGGTGCAGGTAACGAGCCTTTCGAGATA
TGGAAACCACCAACTGCAAGTAAAGGCAATGCTTCACCAAATACTGCAGGAGAAGGAGAA
ATTATTGGAGGGGAACGATGCTCACCATCGACGTACCAGGCATTGAAAGATAGCCGATCG

AAGGTACTATGGAGAGCTGTGGAGTCTGTTATACCTGACGCACGTGAACGTACTGTGCTT
GCTCTCATCGGATCTCCTCGAACACACGAACGATTTCTACGTCGTCCATGCGGCTCGTAC
GGTGCAGCGTTTGAGGATTGTTTGAAGGACGGAAGCACTCCAATATCTAACTTGGTCTTA
TCTGGTGACGGTGTCTTTCCTGGTATTG

#### Exon 2:

>Thaps3 chr_5:650680-650446
GCATTCCTGCTGTAGCACTCAACGGGGCTAGCGCAGCGAATGGATTCGTTGGCATATTTG
ATCAGTGGAGATGTATGGATTATCTTAAGGCCAAAGGAATCATTGCCTAGATAGCCGAGA
AGTTGGACTTCGTCGGCAGCAGTATTTTAGGGTTCCGTGTGCCTTCTTCAACTGAGTATC
GCCCATTGTACTCAGTCGCTTTCAAGCTACTGTCTCTTTCAACGAGTTCACAAAT

#### Translated:

MKLLFVASTLIGVLSFTPPQVLTDTR
HRPPALCNSGDATNDGGDEVHEVDIAVVGAGIGGLCAGAILNTLYDKKVGVYESHYLAGG
CAHSFSRSVKIGDDEQPTTFTFDSGPTIVLGCSKEPYNPLQQVLRAVGVDDQIEWLPYDG
WGMIEHPMQPKEKRWKFKVGPNHFEDGPLQVFASNLNALEEFNQLREITKPLVTGAATIP
AMAMRPGQSALVPLLRYLPSLISIISNGVEASTGPFAPYMNGPIFTVKDPWLRSWLNALA
FSLSGLPADRTSAGAMAYVLFDMHREGAALDYPRGGLGEVVKALVNGVEQKSIGSKVHLS
RHVESIDTNEEGDRVIGLTVRKNGGKKVIVKAKEGVVCNVPMWSLRKLIKNRNALSVLGG
DKATSSSSGLKAKQSWMTSFDTDPSTGRGSVLRPKPAEDTTIEKSLLEKCDSAEMTGSFL
HLHLALNATGLDLQSLEPHYTVMDRGLEGDGKVIDGVKDDSSGELNMIAVSNPCVLDNTL
APEGFIIMHAYGAGNEPFEIWKPPTASKGNASPNTAGEGEIIGGERCSPSTYQALKDSRS
KVLWRAVESVIPDARERTVLALIGSPRTHERFLRRPCGSYGAAFEDCLKDGSTPISNLVL
SGDGVFPGIGIPAVALNGASAANGFVGIFDQWRCMDYLKAKGIIA-

#### Targeting:

SignalP 3.0 - NN: Yes, Cleavage Site VLS-FT

SignalP 3.0 - HMM: Yes, Cleavage Site VLT-DT, Signal Peptide Probability = 0.999

SignalP 4.1: Yes, Cleavage Site VLS-FT, D = 0.635 (D-cutoff = 0.500)

ChloroP: No, Score = 0.460

Analysis using peptide sequences starting with either of the next two in-frame methionines downstream the first one did not result in a more clear different targeting prediction.
