## Appendix D - Additional Sequence-Based Analyses for "Overexpression of *Thalassiosira pseudonana* violaxanthin de-epoxidase-like 2 (VDL2) increases fucoxanthin while stoichiometrically reducing diadinoxanthin cycle pigment abundance"

### 1)PDS

#### Peptide Alignment (With the JGI-Predicted Intron for Thaps3_bd_1474)

```
PDS1          MIITNFILSTVLATSMAFQPHTPILSKPSFSNRVHRSPKIGSSNLVMKDFPKPNVEDTDN    60
bd_1474       ------------------------------------------------------------    0


PDS1          YRYAEAMSTSFKTSLRVTNDSQKKKVAIIGGGLSGLSCAKYLSDAGHEPTVYEARDVLGG    120
bd_1474       ------------------------------------------------------------    0


PDS1          KVSAWQDEDGDWIETGLHIFFGAYPNVMNMFAELGIHDRLQWKIHQMIFAMQELPGEFTT    180
bd_1474       ------------------------------------------------------------    0


PDS1          FDFIPGIPAPFNFGLAILMNQKMLTLGEKIQTAPPLLPMLIEGQSFIDAQDELSVTQFMR    240
bd_1474       ------------------MNQKMLTLGEKIQTAPPLLPMLIEGQSFIDAQDELSVTQFMR    42
                                ******************************************

PDS1          KYGMPERINEEVFIAMAKALDFIDPDKLSMTVVLTAMNRFLNESNGLQMAFLDGNQPDRW    300
bd_1474       KYGMPERINEEVFIAMAKALDFIDPDKLSMTVVLTAMNRFLNESNGLQMAFLDGNQPDRW    102
              ************************************************************

PDS1          CTPTKEYVEARGGKVKLNSPIKEIVTNDDGTINHLLLRSGEKIVADEYVSAMPVDIVKRM    360
bd_1474       CTPTKEYVEARGGKVKLNSPIKEIVTNDDGTINHLLLRSGEKIVADEYVSAMPVDIVKRM    162
              ************************************************************

PDS1          LPTTWQTMPYFRQLDELEGIPVINLHMWFDRKLKAVDHLCFSRSPLLSVYADMSVTCKEY    420
bd_1474       LPTTWQTMPYFRQLDELEGIPVINLHMWFDRKLKAVDHLCFSRSPLLSVYADMSVTCKEY    222
              ************************************************************

PDS1          EDPNKSMLELVFAPCSPIAGGNVNWIGKSDEEIIDATMGELARLFPTEIANDDKWPATKM    480
bd_1474       EDPNKSMLELVFAPCSPIAGGNVNWIGKSDEEIIDATMGELARLFPTEIANDDKWPATKM    282
              ************************************************************

PDS1          QGPNGQAKLEKYAVVKVPRSVYAAIPGE--------------------------------    508
bd_1474       QGPNGQAKLEKYAVVKVPRSVYAAIPGRNKYRPSQTSPIPHFTMAGCYTSQKFLGSMEGA    342
              ***************************.

PDS1          ------------------------------------------------------------    508
bd_1474       TLAGKLAAEVIANRALGNADKPVKEIQQHIIDSASKHVVKEPVGVKGEGAIAFGGGYTVG    402


PDS1          ---------------------         508
bd_1474       KKEEDLLRESDPAQYELAVAK         423
```

#### Peptide Alignment (Disregarding the JGI-Predicted Intron for Thaps3_bd_1474)

```
bd_1474_No_Intron      ------------------------------------------------------------    0
PDS1                   MIITNFILSTVLATSMAFQPHTPILSKPSFSNRVHRSPKIGSSNLVMKDFPKPNVEDTDN    60


bd_1474_No_Intron      ------------------------------------------------------------    0
PDS1                   YRYAEAMSTSFKTSLRVTNDSQKKKVAIIGGGLSGLSCAKYLSDAGHEPTVYEARDVLGG    120
```

```
bd_1474_No_Intron      ------------------------------------------------------------   0
PDS1                   KVSAWQDEDGDWIETGLHIFFGAYPNVMNMFAELGIHDRLQWKIHQMIFAMQELPGEFTT   180

bd_1474_No_Intron      ------------------MNQKMLTLGEKIQTAPPLLPMLIEGQSFIDAQDELSVTQFMR   42
PDS1                   FDFIPGIPAPFNFGLAILMNQKMLTLGEKIQTAPPLLPMLIEGQSFIDAQDELSVTQFMR   240
                                         ******************************************

bd_1474_No_Intron      KYGMPERINEEVFIAMAKALDFIDPDKLSMTVVLTAMNRFLNESNGLQMAFLDGNQPDRW   102
PDS1                   KYGMPERINEEVFIAMAKALDFIDPDKLSMTVVLTAMNRFLNESNGLQMAFLDGNQPDRW   300
                       ************************************************************

bd_1474_No_Intron      CTPTKEYVEARGGKVKLNSPIKEIVTNDDGTINHLLLRSGEKIVADEYVSAMPVDIVKRM   162
PDS1                   CTPTKEYVEARGGKVKLNSPIKEIVTNDDGTINHLLLRSGEKIVADEYVSAMPVDIVKRM   360
                       ************************************************************

bd_1474_No_Intron      LPTTWQTMPYFRQLDELEGIPVINLHMWFDRKLKAVDHLCFSRSPLLSVYADMSVTCKEY   222
PDS1                   LPTTWQTMPYFRQLDELEGIPVINLHMWFDRKLKAVDHLCFSRSPLLSVYADMSVTCKEY   420
                       ************************************************************

bd_1474_No_Intron      EDPNKSMLELVFAPCSPIAGGNVNWIGKSDEEIIDATMGELARLFPTEIANDDKWPATKM   282
PDS1                   EDPNKSMLELVFAPCSPIAGGNVNWIGKSDEEIIDATMGELARLFPTEIANDDKWPATKM   480
                       ************************************************************

bd_1474_No_Intron      QGPNGQAKLEKYAVVKVPRSVYAAIPGE      310
PDS1                   QGPNGQAKLEKYAVVKVPRSVYAAIPGE      508
                       ****************************
```

Genomic Alignment (Extended Sequences Contain 10kb downstream the JGI-Predicted Gene Models)

The abrupt start of sequence differences is highlighted in red.

```
PDS1_Extended         GCAAAGCGTTCCTCTTCGGCGTCAGCCTCCTCCTTTCTTTCACTGTCAGTCAGTTGTGTT    60
PDS1                  GCAAAGCGTTCCTCTTCGGCGTCAGCCTCCTCCTTTCTTTCACTGTCAGTCAGTTGTGTT    60
bd_1474_Extended      ------------------------------------------------------------    0
bd_1474               ------------------------------------------------------------    0

PDS1_Extended         GAACACTTCTGCTTCTTCATTCATCTCCTCATTACTACCACTGCTATTGCAACTGGCTGC    120
PDS1                  GAACACTTCTGCTTCTTCATTCATCTCCTCATTACTACCACTGCTATTGCAACTGGCTGC    120
bd_1474_Extended      ------------------------------------------------------------    0
bd_1474               ------------------------------------------------------------    0

PDS1_Extended         CGTATCTGACTCTCCTTCAATTCGTCACTCCCATAGCGCCACGTTCACCATCATTTATCA    180
PDS1                  CGTATCTGACTCTCCTTCAATTCGTCACTCCCATAGCGCCACGTTCACCATCATTTATCA    180
bd_1474_Extended      ------------------------------------------------------------    0
bd_1474               ------------------------------------------------------------    0

PDS1_Extended         CCATGATCATTACAAATTTCATCCTCTCCACCGTCCTAGCGACATCAATGGCCTTTCAAC    240
PDS1                  CCATGATCATTACAAATTTCATCCTCTCCACCGTCCTAGCGACATCAATGGCCTTTCAAC    240
bd_1474_Extended      ------------------------------------------------------------    0
bd_1474               ------------------------------------------------------------    0

PDS1_Extended         CACACACACCCATCCTCTCCAAACCATCCTTCTCCAACCGTGTCCATCGCTCCCCCAAAA    300
PDS1                  CACACACACCCATCCTCTCCAAACCATCCTTCTCCAACCGTGTCCATCGCTCCCCCAAAA    300
bd_1474_Extended      ------------------------------------------------------------    0
bd_1474               ------------------------------------------------------------    0

PDS1_Extended         TCGGCTCTTCCAACCTCGTTATGAAGGACTTTCCGAAACCAAATGTCGAAGATACAGACA    360
PDS1                  TCGGCTCTTCCAACCTCGTTATGAAGGACTTTCCGAAACCAAATGTCGAAGATACAGACA    360
bd_1474_Extended      ------------------------------------------------------------    0
bd_1474               ------------------------------------------------------------    0
```

```
PDS1_Extended        ACTATCGCTACGCAGAGGCCATGTCCACTAGCTTCAAGACGTCTCTCCGAGTGACGAATG    420
PDS1                 ACTATCGCTACGCAGAGGCCATGTCCACTAGCTTCAAGACGTCTCTCCGAGTGACGAATG    420
bd_1474_Extended     ------------------------------------------------------------    0
bd_1474              ------------------------------------------------------------    0

PDS1_Extended        ATTCACAGAAGAAGAAGGTGGCTATCATTGGAGGAGGATTATCAGGTCTGTCTTGTGCCA    480
PDS1                 ATTCACAGAAGAAGAAGGTGGCTATCATTGGAGGAGGATTATCAGGTCTGTCTTGTGCCA    480
bd_1474_Extended     ------------------------------------------------------------    0
bd_1474              ------------------------------------------------------------    0

PDS1_Extended        AGTACCTCTCCGATGCCGGGCATGAACCCACCGTATACGAAGCACGTGATGTACTCGGAG    540
PDS1                 AGTACCTCTCCGATGCCGGGCATGAACCCACCGTATACGAAGCACGTGATGTACTCGGAG    540
bd_1474_Extended     ------------------------------------------------------------    0
bd_1474              ------------------------------------------------------------    0

PDS1_Extended        GAAAGGTGTCAGCGTGGCAAGATGAAGATGGAGACTGGATCGAAACAGGTCTTCACATCT    600
PDS1                 GAAAGGTGTCAGCGTGGCAAGATGAAGATGGAGACTGGATCGAAACAGGTCTTCACATCT    600
bd_1474_Extended     ------------------------------------------------------------    0
bd_1474              ------------------------------------------------------------    0

PDS1_Extended        TCTTCGGAGCATACCCCAACGTTATGAACATGTTCGCTGAGCTTGGCATCCACGATAGGC    660
PDS1                 TCTTCGGAGCATACCCCAACGTTATGAACATGTTCGCTGAGCTTGGCATCCACGATAGGC    660
bd_1474_Extended     ------------------------------------------------------------    0
bd_1474              ------------------------------------------------------------    0

PDS1_Extended        TTCAGTGGAAGATTCACCAAATGATTTTCGCAATGCAGGAACTTCCCGGAGAGTTCACTA    720
PDS1                 TTCAGTGGAAGATTCACCAAATGATTTTCGCAATGCAGGAACTTCCCGGAGAGTTCACTA    720
bd_1474_Extended     ------------------------------------------------------------    0
bd_1474              ------------------------------------------------------------    0

PDS1_Extended        CCTTTGATTTCATCCCTGGTATTCCAGCTCCGTTCAACTTTGGATTGGCCATTCTTATGA    780
PDS1                 CCTTTGATTTCATCCCTGGTATTCCAGCTCCGTTCAACTTTGGATTGGCCATTCTTATGA    780
bd_1474_Extended     -----------------------------------------------------CTTATGA    7
bd_1474              -----------------------------------------------------CTTATGA    7
                                                                          *******

PDS1_Extended        ATCAAAAGATGTTGACGTTGGGTGAAAAAATTCAGACCGCTCCTCCTCTTCTTCCTATGC    840
PDS1                 ATCAAAAGATGTTGACGTTGGGTGAAAAAATTCAGACCGCTCCTCCTCTTCTTCCTATGC    840
bd_1474_Extended     ATCAAAAGATGTTGACGTTGGGTGAAAAAATTCAGACCGCTCCTCCTCTTCTTCCTATGC    67
bd_1474              ATCAAAAGATGTTGACGTTGGGTGAAAAAATTCAGACCGCTCCTCCTCTTCTTCCTATGC    67
                     ************************************************************

PDS1_Extended        TTATTGAGGGACAGTCATTCATTGATGCTCAGGATGAGTTGAGTGTGACGCAGTTCATGA    900
PDS1                 TTATTGAGGGACAGTCATTCATTGATGCTCAGGATGAGTTGAGTGTGACGCAGTTCATGA    900
bd_1474_Extended     TTATTGAGGGACAGTCATTCATTGATGCTCAGGATGAGTTGAGTGTGACGCAGTTCATGA    127
bd_1474              TTATTGAGGGACAGTCATTCATTGATGCTCAGGATGAGTTGAGTGTGACGCAGTTCATGA    127
                     ************************************************************

PDS1_Extended        GGAAGTACGGTATGCCTGAGAGAATCAACGAGGAGGTGTTTATTGCGATGGCCAAGGCGT    960
PDS1                 GGAAGTACGGTATGCCTGAGAGAATCAACGAGGAGGTGTTTATTGCGATGGCCAAGGCGT    960
bd_1474_Extended     GGAAGTACGGTATGCCTGAGAGAATCAACGAGGAGGTGTTTATTGCGATGGCCAAGGCGT    187
bd_1474              GGAAGTACGGTATGCCTGAGAGAATCAACGAGGAGGTGTTTATTGCGATGGCCAAGGCGT    187
                     ************************************************************

PDS1_Extended        TGGACTTTATTGATCCTGATAAGTTGAGTATGACTGTGGTGCTTACGGCTATGAACAGGT    1020
PDS1                 TGGACTTTATTGATCCTGATAAGTTGAGTATGACTGTGGTGCTTACGGCTATGAACAGGT    1020
bd_1474_Extended     TGGACTTTATTGATCCTGATAAGTTGAGTATGACTGTGGTGCTTACGGCTATGAACAGGT    247
bd_1474              TGGACTTTATTGATCCTGATAAGTTGAGTATGACTGTGGTGCTTACGGCTATGAACAGGT    247
                     ************************************************************

PDS1_Extended        TCTTGAATGAGAGTAATGGACTTCAGATGGCATTCTTGGATGGAAATCAGCCTGATAGGT    1080
PDS1                 TCTTGAATGAGAGTAATGGACTTCAGATGGCATTCTTGGATGGAAATCAGCCTGATAGGT    1080
bd_1474_Extended     TCTTGAATGAGAGTAATGGACTTCAGATGGCATTCTTGGATGGAAATCAGCCTGATAGGT    307
bd_1474              TCTTGAATGAGAGTAATGGACTTCAGATGGCATTCTTGGATGGAAATCAGCCTGATAGGT    307
```

```
                        ************************************************************

PDS1_Extended           GGTGCACTCCCACGAAGGAGTATGTGGAAGCACGCGGAGGAAAGGTCAAATTGAACTCTC    1140
PDS1                    GGTGCACTCCCACGAAGGAGTATGTGGAAGCACGCGGAGGAAAGGTCAAATTGAACTCTC    1140
bd_1474_Extended        GGTGCACTCCCACCAAGGAGTATGTGGAAGCACGCGGAGGAAAGGTCAAATTGAACTCTC    367
bd_1474                 GGTGCACTCCCACCAAGGAGTATGTGGAAGCACGCGGAGGAAAGGTCAAATTGAACTCTC    367
                        ************* **********************************************

PDS1_Extended           CCATTAAGGAGATTGTGACCAACGACGATGGAACTATCAATCACCTTCTCCTTCGATCTG    1200
PDS1                    CCATTAAGGAGATTGTGACCAACGACGATGGAACTATCAATCACCTTCTCCTTCGATCTG    1200
bd_1474_Extended        CCATTAAGGAGATTGTGACCAACGACGATGGAACTATCAATCACCTTCTCCTTCGATCTG    427
bd_1474                 CCATTAAGGAGATTGTGACCAACGACGATGGAACTATCAATCACCTTCTCCTTCGATCTG    427
                        ************************************************************

PDS1_Extended           GCGAGAAGATTGTGGCCGATGAATACGTCTCTGCCATGCCCGTGGACATCGTCAAACGTA    1260
PDS1                    GCGAGAAGATTGTGGCCGATGAATACGTCTCTGCCATGCCCGTGGACATCGTCAAACGTA    1260
bd_1474_Extended        GCGAGAAGATTGTGGCCGATGAATACGTCTCTGCCATGCCCGTGGACATCGTCAAACGTA    487
bd_1474                 GCGAGAAGATTGTGGCCGATGAATACGTCTCTGCCATGCCCGTGGACATCGTCAAACGTA    487
                        ************************************************************

PDS1_Extended           TGCTTCCCACAACGTGGCAGACTATGCCCTACTTCCGTCAGCTTGACGAACTTGAGGGCA    1320
PDS1                    TGCTTCCCACAACGTGGCAGACTATGCCCTACTTCCGTCAGCTTGACGAACTTGAGGGCA    1320
bd_1474_Extended        TGCTTCCCACAACGTGGCAGACTATGCCCTACTTCCGTCAGCTTGACGAACTTGAGGGCA    547
bd_1474                 TGCTTCCCACAACGTGGCAGACTATGCCCTACTTCCGTCAGCTTGACGAACTTGAGGGCA    547
                        ************************************************************

PDS1_Extended           TCCCTGTTATCAACTTGCACATGTGGTTCGATCGTAAGTTGAAAGCAGTCGACCATCTTT    1380
PDS1                    TCCCTGTTATCAACTTGCACATGTGGTTCGATCGTAAGTTGAAAGCAGTCGACCATCTTT    1380
bd_1474_Extended        TCCCTGTTATCAACTTGCACATGTGGTTCGATCGTAAGTTGAAAGCAGTCGACCATCTTT    607
bd_1474                 TCCCTGTTATCAACTTGCACATGTGGTTCGATCGTAAGTTGAAAGCAGTCGACCATCTTT    607
                        ************************************************************

PDS1_Extended           GCTTCAGTCGCTCCCCACTCCTTTCCGTCTACGCCGACATGTCCGTCACATGCAAGGAGT    1440
PDS1                    GCTTCAGTCGCTCCCCACTCCTTTCCGTCTACGCCGACATGTCCGTCACATGCAAGGAGT    1440
bd_1474_Extended        GCTTCAGTCGCTCCCCACTCCTTTCCGTCTACGCTGACATGTCCGTCACATGCAAGGAGT    667
bd_1474                 GCTTCAGTCGCTCCCCACTCCTTTCCGTCTACGCTGACATGTCCGTCACATGCAAGGAGT    667
                        ********************************** *************************

PDS1_Extended           ACGAAGATCCCAACAAGTCCATGTTGGAATTGGTCTTTGCTCCCTGCTCTCCTATTGCCG    1500
PDS1                    ACGAAGATCCCAACAAGTCCATGTTGGAATTGGTCTTTGCTCCCTGCTCTCCTATTGCCG    1500
bd_1474_Extended        ACGAAGATCCCAACAAGTCCATGTTGGAATTGGTCTTTGCTCCCTGCTCTCCTATTGCCG    727
bd_1474                 ACGAAGATCCCAACAAGTCCATGTTGGAATTGGTCTTTGCTCCCTGCTCTCCTATTGCCG    727
                        ************************************************************

PDS1_Extended           GAGGAAATGTCAACTGGATTGGAAAGTCAGATGAGGAAATCATTGATGCTACCATGGGTG    1560
PDS1                    GAGGAAATGTCAACTGGATTGGAAAGTCAGATGAGGAAATCATTGATGCTACCATGGGTG    1560
bd_1474_Extended        GAGGAAATGTCAACTGGATTGGAAAGTCAGATGAGGAAATCATTGATGCTACCATGGGTG    787
bd_1474                 GAGGAAATGTCAACTGGATTGGAAAGTCAGATGAGGAAATCATTGATGCTACCATGGGTG    787
                        ************************************************************

PDS1_Extended           AGCTTGCTCGCCTTTTCCCTACCGAGATTGCGAATGATGATAAGTGGCCTGCTACGAAGA    1620
PDS1                    AGCTTGCTCGCCTTTTCCCTACCGAGATTGCGAATGATGATAAGTGGCCTGCTACGAAGA    1620
bd_1474_Extended        AGCTTGCTCGCCTTTTCCCTACCGAGATTGCGAATGATGATAAGTGGCCTGCTACGAAGA    847
bd_1474                 AGCTTGCTCGCCTTTTCCCTACCGAGATTGCGAATGATGATAAGTGGCCTGCTACGAAGA    847
                        ************************************************************

PDS1_Extended           TGCAGGGACCTAATGGACAGGCAAAGCTTGAGAAGTATGCTGTTGTGAAGGTGCCAAGGA    1680
PDS1                    TGCAGGGACCTAATGGACAGGCAAAGCTTGAGAAGTATGCTGTTGTGAAGGTGCCAAGGA    1680
bd_1474_Extended        TGCAGGGACCTAATGGACAGGCAAAGCTTGAGAAGTATGCTGTTGTGAAGGTGCCAAGGA    907
bd_1474                 TGCAGGGACCTAATGGACAGGCAAAGCTTGAGAAGTATGCTGTTGTGAAGGTGCCAAGGA    907
                        ************************************************************

PDS1_Extended           GTGTGTATGCTGCCATTCCTGGTGAGTGAAAAATAGTGTGCGGTGAATCTTCGTCGTCTA    1740
PDS1                    GTGTGTATGCTGCCATTCCTGGTGAGTGAAAAATAGTGTGCGGTGAATCTTCGTCGTCTA    1740
bd_1474_Extended        GTGTGTATGCTGCCATTCCTGGTGAGTGAAAAAGAGTGTGCGGTGAATCTTCGTCGTCTA    967
bd_1474                 GTGTGTATGCTGCCATTCCTGGTGAGTGAAAAAGAGTGTGCGGTGAATCTTCGTCGTCTA    967
                        ********************************* **************************

PDS1_Extended           CCTTGGCCAACTACTGACTCTTGTTCTCTTTAAATCATATCAGGACGTAACAAATACCGC    1800
PDS1                    CCTTGGCCAACTACTGACTCTTGTTCTCTTTAAATCATATCAGGACGTAACAAATACCGC    1800
bd_1474_Extended        CCTTGGCCAACTACTGACTCTTGTTCTCTTTAAATCATATCAGGACGTAACAAATACCGC    1027
```

```
bd_1474             CCTTGGCCAACTACTGACTCTTGTTCTCTTTAAATCATATCAGGACGTAACAAATACCGC    1027
                    ************************************************************

PDS1_Extended       CCCAGTCAGACCTCCCCCATCCCACACTTCACCATGGCTGGATGCTATACCTCACAAAAG    1860
PDS1                CCCAGTCAGACCTCCCCCATCCCACACTTCACCATGGCTGGATGCTATACCTCACAAAAG    1860
bd_1474_Extended    CCCAGTCAGACCTCCCCCATCCCACACTTCACCATGGCTGGATGCTATACCTCACAAAAG    1087
bd_1474             CCCAGTCAGACCTCCCCCATCCCACACTTCACCATGGCTGGATGCTATACCTCACAAAAG    1087
                    ************************************************************

PDS1_Extended       TTCCTCGGATCCATGGAGGGTGCCACCCTCGCCGGGAAGCTTGCTGCCGAGGTCATTGCC    1920
PDS1                TTCCTCGGATCCATGGAGGGTGCCACCCTCGCCGGGAAGCTTGCTGCCGAGGTCATTGCC    1920
bd_1474_Extended    TTCCTCGGATCCATGGAGGGTGCCACCCTCGCCGGGAAGCTTGCTGCCGAGGTCATTGCC    1147
bd_1474             TTCCTCGGATCCATGGAGGGTGCCACCCTCGCCGGGAAGCTTGCTGCCGAGGTCATTGCC    1147
                    ************************************************************

PDS1_Extended       AACCGTGCCCTCGGAAATGCGGATAAGCCAGTCAAGGAGATTCAGCAACACATTATCGAC    1980
PDS1                AACCGTGCCCTCGGAAATGCGGATAAGCCAGTCAAGGAGATTCAGCAACACATTATCGAC    1980
bd_1474_Extended    AACCGTGCCCTCGGAAATGCGGATAAGCCAGTCAAGGAGATTCAGCAACACATTATCGAC    1207
bd_1474             AACCGTGCCCTCGGAAATGCGGATAAGCCAGTCAAGGAGATTCAGCAACACATTATCGAC    1207
                    ************************************************************

PDS1_Extended       TCGGCTAGTAAGCATGTTGTGAAGGAGCCAGTGGGTGTGAAGGGAGAGGGAGCGATTGCA    2040
PDS1                TCGGCTAGTAAGCATGTTGTGAAGGAGCCAGTGGGTGTGAAGGGAGAGGGAGCGATTGCA    2040
bd_1474_Extended    TCGGCTAGTAAGCATGTTGTGAAGGAGCCAGTGGGTGTGAAGGGAGAGGGAGCGATTGCA    1267
bd_1474             TCGGCTAGTAAGCATGTTGTGAAGGAGCCAGTGGGTGTGAAGGGAGAGGGAGCGATTGCA    1267
                    ************************************************************

PDS1_Extended       TTTGGAGGGGGGTATACTGTTGGAAAGAAGGAGGAGGATTTGTTGAGGGAGTCGGATCCT    2100
PDS1                TTTGGAGGGGGGTATACTGTTGGAAAGAAGGAGGAGGATTTGTTGAGGGAGTCGGATCCT    2100
bd_1474_Extended    TTTGGAGGGGGGTATACTGTTGGAAAGAAGGAGGAGGATTTGTTGAGGGAGTCGGATCCT    1327
bd_1474             TTTGGAGGGGGGTATACTGTTGGAAAGAAGGAGGAGGATTTGTTGAGGGAGTCGGATCCT    1327
                    ************************************************************

PDS1_Extended       GCTCAGTATGAGTTGGCAGTAGCCAAGTAAGGAAGAGTATTATTAAAGTAAACAGCTAGA    2160
PDS1                GCTCAGTATGAGTTGGCAGTAGCCAAGTAAGGAAGAGTATTATTAAAGTAAACAGCTAGA    2160
bd_1474_Extended    GCTCAGTATGAGTTGGCAGTAGCCAAGTAAGGAAGAGTATTATTAAAGTAAACAGCTAGA    1387
bd_1474             GCTCAGTATGAGTTGGCAGTAGCCAAGTAA------------------------------    1357
                    ******************************

PDS1_Extended       TTGATTTTGAAGGAGTTTTGTATGTAGCTTAGTACCTTTAAGGTACTTGATGTACTTACT    2220
PDS1                TTGATTTTGAA-------------------------------------------------    2171
bd_1474_Extended    TTGATTTTGAAGGAGTTTTGTATGTAACTTAGTACCTTTAAGGTACTTGATGTTCTTACT    1447
bd_1474             ------------------------------------------------------------    1357


PDS1_Extended       TTCCGCTTGGAGAGCCATTTCTTTGATTATATGACAGAATGGAGTTTTCCCTTCATCCTT    2280
PDS1                ------------------------------------------------------------    2171
bd_1474_Extended    TTCCGCTTGGAGAGCCATTTCTTTGATTTTACGACAGAATTGAGTTTTCCCTTCATCCTT    1507
bd_1474             ------------------------------------------------------------    1357


PDS1_Extended       CTCGTTGATTGATTTTGGATCAGATGATATTTATGCATCTCTCACCTGGTGCCACTAACG    2340
PDS1                ------------------------------------------------------------    2171
bd_1474_Extended    CTCGTTGATTGATTTTGGATCAGATGATATTTATGCATCTCTCACCTGGTGCCACTAACG    1567
bd_1474             ------------------------------------------------------------    1357


PDS1_Extended       ATACATACGAAAGAACCTCCCCCCACCTTGCCACACCAACTCATTCATCTCATCTCCTTA    2400
PDS1                ------------------------------------------------------------    2171
bd_1474_Extended    ATACATACGAAAGAACCTCCC-CCACCTTGCCACACCAACTCATTCATCTCATCTCCTTA    1626
bd_1474             ------------------------------------------------------------    1357


PDS1_Extended       TTTGATTTTGTGCTCCGTAAACAGAACTCTCCTCCTGACAATCGGATCAAATTTTACGAA    2460
PDS1                ------------------------------------------------------------    2171
bd_1474_Extended    TTTGATTTTGTGCTCCGTAAACAGAACTCTCCTCCTGACAATTGGATCAAATTTTACGAA    1686
bd_1474             ------------------------------------------------------------    1357


PDS1_Extended       CTTGAACTTCTCTGGTGTCTTCGATACGTTTCGTCGTGTGGTGTAGAAGAAGCCAGTCCC    2520
PDS1                ------------------------------------------------------------    2171
```

```
bd_1474_Extended      CTTGAACTTCTCTGGTGTCTTCGATACGTTTCGTCGTGTGGTGTAGAAGAAGCCAGTCCC    1746
bd_1474               ------------------------------------------------------------    1357

PDS1_Extended         CGCCGAGCTTAAGAGCTGCGTTGTTAACGACGTGAATGGAAAGAGGAGTCAGTCAGTGTT    2580
PDS1                  ------------------------------------------------------------    2171
bd_1474_Extended      CGCCGAGCTTAAGAGCTGCGTTGTTAACGACGTGAATGGAAAGAGGAGTCAGTCAGTGTT    1806
bd_1474               ------------------------------------------------------------    1357

PDS1_Extended         GGATGAGATGATATGCAGCATAGCCACACCTCTTCAATCATACAAAGGCTGATTTGCACA    2640
PDS1                  ------------------------------------------------------------    2171
bd_1474_Extended      GGATGAGATGATATGCAGCATAGCCACAACTCTTCAATCATACAAAGGCTGATTTGCACA    1866
bd_1474               ------------------------------------------------------------    1357

PDS1_Extended         CAACGTGTACCGTCACCTCATCTGACGGAACAGCACCTCCCACTCACTTTGATGGGGATC    2700
PDS1                  ------------------------------------------------------------    2171
bd_1474_Extended      CAACGTGTACCGTCACCTCATCTGACGGAACAGCACCTCCCACTCACTTTGATGGGGATC    1926
bd_1474               ------------------------------------------------------------    1357

PDS1_Extended         GTCTTGCCCTTCCCTCTGGCCATGTGAAGAATAGCGGTCGGCTGCATGGTATTTTACTCT    2760
PDS1                  ------------------------------------------------------------    2171
bd_1474_Extended      GTCTTGCCCTTCCCTCTGGCCATGTGAAGAATAGCGGTCGGCTGCATGGTATTTTACTCT    1986
bd_1474               ------------------------------------------------------------    1357

PDS1_Extended         TCTCGATAATGCAAACGTTTTCTAAGATGGGTTGTTGCCGGTTGTGGTGAAGGGGTGACG    2820
PDS1                  ------------------------------------------------------------    2171
bd_1474_Extended      TCTCGATACTGCAGACGTTTTCTAAGATGGGTTGTTGCCGGTTGTGGTGAAGGGGTGGCG    2046
bd_1474               ------------------------------------------------------------    1357

PDS1_Extended         GTGGTGGTGGTTGTGGTTGTTGTTTTGTGTGCGTGACGAGTTGTGATGTTGTGGTTTTGT    2880
PDS1                  ------------------------------------------------------------    2171
bd_1474_Extended      GTGGTGGTCGTTGTGGTTGTTGTTTTGTGTGCGTGACGAGTTGTGATGTTGTGGTTTTGT    2106
bd_1474               ------------------------------------------------------------    1357

PDS1_Extended         GAACAGTTCAGTAACTTGTTGTCGTGACGGACTAACGTGATTCCGTGGATACCAAGAAGC    2940
PDS1                  ------------------------------------------------------------    2171
bd_1474_Extended      GAACAGTTCTGTAACGTGTTGTCGTGACGGACTAACGTGATTCCGTGGATACCAAGAAGC    2166
bd_1474               ------------------------------------------------------------    1357

PDS1_Extended         ACTAACATGGTAACCCCC--CCCCCAGACCAACAAATCATTTCATCCCCACCAACTCCAT    2998
PDS1                  ------------------------------------------------------------    2171
bd_1474_Extended      ACTAACATAGTAACCCCACCCCCCCAGACCAACAAATCATTTCATCCCCACCAACTCCAT    2226
bd_1474               ------------------------------------------------------------    1357

PDS1_Extended         CACACTCCCTCATCGATACATCCAGCACAAACCATGATAGTTGATAGTCTCCATTGATTC    3058
PDS1                  ------------------------------------------------------------    2171
bd_1474_Extended      CACACTCCCTCATCGATACATCCAGCACAAACCATGATAGTTGATAGTCTCCATTGATTC    2286
bd_1474               ------------------------------------------------------------    1357

PDS1_Extended         CTCGTCGCAGTCTGCTACAGCGATGCCTCCACAAACAGCCGTCACGATGATAAACATTGT    3118
PDS1                  ------------------------------------------------------------    2171
bd_1474_Extended      CTCGTCGCAGTCTGCTACAGCGATGCCTCCACAAACAGCCGTCACATTGATAAACATTGT    2346
bd_1474               ------------------------------------------------------------    1357

PDS1_Extended         ACTCCCACTCCAGACTGGGTGGAGTCGACAGTGAAGTCAAGACGATGATGAATGTTGGCT    3178
PDS1                  ------------------------------------------------------------    2171
bd_1474_Extended      ACTCCCACTCCAGACTGGGTGGAGTCGACAGTGAAGTCAAGACGATGATGAATGTTGGCT    2406
bd_1474               ------------------------------------------------------------    1357

PDS1_Extended         GCAAACGCACAACCCCGAGCCCAAGTGAACGATGGCACTACAATCATCCCGCCGCCATGC    3238
```

```
PDS1                  ------------------------------------------------------------    2171
bd_1474_Extended      GCAAACGCACAACCCCGAGCCCAAGTGAACGATGGCACTACAATCATCCCGCCGCCATGC    2466
bd_1474               ------------------------------------------------------------    1357

PDS1_Extended         CTTTGAAACCAATTCACTAGACCACTGCGAAACACCACTTTGTGCCTACGAAAATGTTCA    3298
PDS1                  ------------------------------------------------------------    2171
bd_1474_Extended      CTTTGAAACCAATTCACTAGACCACTGCGAAACACCACTTTGTGCCTACGAAAATGTTCA    2526
bd_1474               ------------------------------------------------------------    1357

PDS1_Extended         ACCAGTCCTCGAGATGATGGCGAAGCATCTTCACGTCCAACCATCAATGTTACGCATCTG    3358
PDS1                  ------------------------------------------------------------    2171
bd_1474_Extended      AACAGTCCTCGAGATGATGGCGAAGCATCTTCACGTCCAACCATCAAAGTTACGCATCTG    2586
bd_1474               ------------------------------------------------------------    1357

PDS1_Extended         GGATCCGTATTATTGCGATGGGACTGTCAAGCAACATCTAGCCTCTTTGGGATACGATCG    3418
PDS1                  ------------------------------------------------------------    2171
bd_1474_Extended      GGATCCGTATTATTGCGATGGGACTGTCAAGCAACATCTAGCCTCTTTGGGATACGATCG    2646
bd_1474               ------------------------------------------------------------    1357

PDS1_Extended         TGTGATTAACGAAAACTTTGACTTTTACAAGCGAGTAGAGGACAACACTATCCCCGAGCA    3478
PDS1                  ------------------------------------------------------------    2171
bd_1474_Extended      TGTGATTAACGAAAACGTTGACTTTTACAAGCGAGTAGAGGACAACACTATCCCCGAGCA    2706
bd_1474               ------------------------------------------------------------    1357

PDS1_Extended         CGATGTCTTGTTGACGAATCCTCCGTACAGCGGCGATCACATCGAACGATTGCTTAAGTT    3538
PDS1                  ------------------------------------------------------------    2171
bd_1474_Extended      CGATGTCTTGTTGACGAATCCTCCGTACAGCGGCGATCACATCGAACGATTGCTTAAGTT    2766
bd_1474               ------------------------------------------------------------    1357

PDS1_Extended         TGTTACGACGGTGAATGATAAGCCATTTTGTCTACTAATGCCAAATTGGGTTGCGAGAAA    3598
PDS1                  ------------------------------------------------------------    2171
bd_1474_Extended      TGTTACGACGGTGAATGATAAGCCATTTTGTCTACTAATGCCAAATTGGGTTGCGAGAAA    2826
bd_1474               ------------------------------------------------------------    1357

PDS1_Extended         GAAGGAGTACAAATCCATCATTGGTAAAACGAATCTGTTTTACGTATCCCCCATCGAGGT    3658
PDS1                  ------------------------------------------------------------    2171
bd_1474_Extended      GAAGGAGTACAAATCCATCATTGGTAAAACGAATCTGTTTTACGTATCCCCCATAGAGGT    2886
bd_1474               ------------------------------------------------------------    1357

PDS1_Extended         ATACACGTATGCTATGCCAACTTGGAATTCGAAACCGGAACACGTCGACGAGGAGACGGG    3718
PDS1                  ------------------------------------------------------------    2171
bd_1474_Extended      ATACACGTATGCTATGCCAACTTGGAATTCGAAACCGGAACACGTCGACGAGGAGACGGG    2946
bd_1474               ------------------------------------------------------------    1357

PDS1_Extended         AAAGACGACTCCCTATTTGAGTTCTTGGTATGTATCGTTAAGAAGCAATAGTGAGGCAAC    3778
PDS1                  ------------------------------------------------------------    2171
bd_1474_Extended      AAAGACGACACCCTATTTGAGTTCTTGGTATGTATCGTTAAGAAGCAATAGTGAGGCAAC    3006
bd_1474               ------------------------------------------------------------    1357

PDS1_Extended         GAGTAGGATAGAGAATAAGCTTGATTCTATTGCAAAACGGCAGCAACCACCTGTTTGGGT    3838
PDS1                  ------------------------------------------------------------    2171
bd_1474_Extended      GGGTAGGATAGAGAATAAGCTTGATTCTATTGCAAAACGGCAGCAACCACCTGTTTGGGT    3066
bd_1474               ------------------------------------------------------------    1357

PDS1_Extended         TGTAGCCAAGACGGTCAAAGGGCTAAAGTGGAAGATTCAAAAGGTGAAAGAAAAGAAGAG    3898
PDS1                  ------------------------------------------------------------    2171
bd_1474_Extended      TGTAGCCAAGACGGTCAAATCGACAGGAGCACCCAATTCCTTGACAACATTGTTGTGTGT    3126
bd_1474               ------------------------------------------------------------    1357
```

```
PDS1_Extended        GTGATTGGAGATTTT-----CTCCTCGGAGATGCTAAATCGCTTAGCAATGGAAGTGAGA    3953
PDS1                 ------------------------------------------------------------    2171
bd_1474_Extended     CTCAATGGGAGCGATGAAATCGCATGGGACAATCTTCGTGCCTTGATGCAATAATTGATA    3186
bd_1474              ------------------------------------------------------------    1357

PDS1_Extended        A-------------CTGAATTGTCAATGATACACGTTATATATGTCATAAACTGAAGTAC    4000
PDS1                 ------------------------------------------------------------    2171
bd_1474_Extended     GCTAGGACCAAGTGTTGACGACTGTGAGGTTCAGATTATGGCT-----AGTTTTACGAAC    3241
bd_1474              ------------------------------------------------------------    1357

PDS1_Extended        CCATCCCACTCAATCGACA----ATAAG----------AGTTCATACCCAACGA----AC    4042
PDS1                 ------------------------------------------------------------    2171
bd_1474_Extended     TACGAGAAACGAATGGCCAGCATCTGAGAACTCAGTCAAGTCTATGACCTACGAAGGACA    3301
bd_1474              ------------------------------------------------------------    1357

PDS1_Extended        GGTTTCCCCCGTTTTATCACTCCGGAGTTGCTCACTCGAGGACGAGTTCGATTTGTTTGC    4102
PDS1                 ------------------------------------------------------------    2171
bd_1474_Extended     GACACCCCCCATCATAGCACTTGACAACAAAT-----GTCGTCCAGAAC-----------    3345
bd_1474              ------------------------------------------------------------    1357

PDS1_Extended        TTGATGGTAGTCATTACTCCCGTTGCCTCTCTCAAAGTTTGTTTGTTGTACTTGATACTC    4162
PDS1                 ------------------------------------------------------------    2171
bd_1474_Extended     -----------------------TACATAGATGAGAGATGGTGTCATTTGCATGACAACT    3382
bd_1474              ------------------------------------------------------------    1357

PDS1_Extended        TATCCCCCCCAACCCGTCGCACCCTACAAAGCTTCGTTTCATTTCATTTCATTGCCCAGC    4222
PDS1                 ------------------------------------------------------------    2171
bd_1474_Extended     GTTGCGAAGCAACAGCAAGTCCAAAA--GAGAAGATTTGGAAGGC---AGAACGCGAAGA    3437
bd_1474              ------------------------------------------------------------    1357

PDS1_Extended        CCCCTCCAATGTGCCGCTTTTTATTGAGTCAATCCTGTTCTTTGTCTTGATCCCCATCAA    4282
PDS1                 ------------------------------------------------------------    2171
bd_1474_Extended     CATGAAGAAGCTGACGCGGTACGTGTACCTGACTCC------GGGAGTGGTGGCCAGTCA    3491
bd_1474              ------------------------------------------------------------    1357

PDS1_Extended        AAGTAAACATAATTAGAAGTGAGATGGTTCAGATTTCACCTTCACAGTGTACATGAACAA    4342
PDS1                 ------------------------------------------------------------    2171
bd_1474_Extended     GTG-----AGACTTAGA-------GCGGTGGTGTGACCCG---------------GACCA    3524
bd_1474              ------------------------------------------------------------    1357

PDS1_Extended        TCGTCTCACCTTCTCCTCTCTCCACCCAATCCTCCACCACTACCAAACCTCGCTAACTTT    4402
PDS1                 ------------------------------------------------------------    2171
bd_1474_Extended     TGTGATGATGTCTTTGTATCTGGCGTGAAGGATACACGAGGAGGAGAAGTGATAGTACTG    3584
bd_1474              ------------------------------------------------------------    1357

PDS1_Extended        CCAT-------AGGAGCAGCACCTCCACCACTAAAATCAGCAGCCTTCACATA------C    4449
PDS1                 ------------------------------------------------------------    2171
bd_1474_Extended     GAGTCAGTGCTAACGGTAGTGTTGTCATGGGAATAATGAGCAGCCAACGCATTCTGACGA    3644
bd_1474              ------------------------------------------------------------    1357

PDS1_Extended        AACACATTATTACACCGAATCAACACCTCCCCCAACGTACCCGCAAACTGTCCATCAATG    4509
PDS1                 ------------------------------------------------------------    2171
bd_1474_Extended     AATGAAGAATAACGGGACATGAAAGAGTC-------------GCGAAGATGGCGAAAATG    3691
bd_1474              ------------------------------------------------------------    1357

PDS1_Extended        AACTCCTCCGTCTCCTCCAATTGTAAATTCATGTAGGCATCCGAGGAGGCGAGCTTGCCT    4569
PDS1                 ------------------------------------------------------------    2171
bd_1474_Extended     AGCACTGTCTTTCATCTCTTTTG--------TGAAAAGATCCGCGGGATTGATCTTGCCA    3743
bd_1474              ------------------------------------------------------------    1357
```

```
PDS1_Extended        TGGTATTCTTGGCCCCAC------TTGA----------GACGGACGCGGACGTGTTTGCC    4613
PDS1                 ------------------------------------------------------------    2171
bd_1474_Extended     GAAACATGTTTCACTTCAATGAGTTTGTGCTGCACCCACTCGCGCACAGCGTTTTCTTTC    3803
bd_1474              ------------------------------------------------------------    1357

PDS1_Extended        TGTTAAGTCGGC---------TAGGA------------AGGGTTTGGGGTTGACGATTAG    4652
PDS1                 ------------------------------------------------------------    2171
bd_1474_Extended     AGTTCCATGTGTCGAATAGCTTTGGAAGTTGTGTTGTGGGTCCATGAGACGCAAGATTCA    3863
bd_1474              ------------------------------------------------------------    1357

PDS1_Extended        TTGTGACTGTTGATGAGAGTTTGGTGGAGGAGGAGGAGATTTATAGGTGAGAAGGTGGGC    4712
PDS1                 ------------------------------------------------------------    2171
bd_1474_Extended     TTGTAATTCCAGATAGTAGTG---------------------------------------    3884
bd_1474              ------------------------------------------------------------    1357

PDS1_Extended        AGAGGCATCAGACGAAGGCATCGGCCTCACCATTGAACGAGCAGATAATAGATTAATCAA    4772
PDS1                 ------------------------------------------------------------    2171
bd_1474_Extended     -GCACCGTCAGCGTCAGAAATGGGAACGCCAATAGAACAGAAACCATTGGCGATGTTGCG    3943
bd_1474              ------------------------------------------------------------    1357

PDS1_Extended        TCCCTTCTAGATGGGCTTACCATTGTATTCGGGGCCGTTAGCGTTCGAAAGAGTGAAGCG    4832
PDS1                 ------------------------------------------------------------    2171
bd_1474_Extended     AATAGCCAAAGTGAGTTTCCCAGCGACATTGGTG--------GCTCGAATCTCTGCTTCA    3995
bd_1474              ------------------------------------------------------------    1357

PDS1_Extended        GG----TAAAGGCACCGACCGTCTTGTCGTCGTCGTCGTGGTGAACCAAATGGTCCCGCC    4888
PDS1                 ------------------------------------------------------------    2171
bd_1474_Extended     CAAGAACTTAGTGAAGTTCTGTCTTGGCGTTCTGATGCCCAT----GAAATCGGTCCGCC    4051
bd_1474              ------------------------------------------------------------    1357

PDS1_Extended        GTTTCTGAAAACAACAAT---AGTGACAATATGCTGTTAAACGAGAGGTTGGGAGTAACC    4945
PDS1                 ------------------------------------------------------------    2171
bd_1474_Extended     AGATCGAAACACAATAGCACCACTCATACTACGAAGCTTGAAGAGGGGAAGGAGCGTT-C    4110
bd_1474              ------------------------------------------------------------    1357

PDS1_Extended        AAGCCAAGCCCGCAACGAAGGCAACAGGAGACCAACAA----------------------    4983
PDS1                 ------------------------------------------------------------    2171
bd_1474_Extended     CTTCATGTACCGCGTTGCCGATTTGACTACCCCAACATGCATCACTGTAAGTAGTCAGTT    4170
bd_1474              ------------------------------------------------------------    1357

PDS1_Extended        --ACAACCAGAACTGATCAAAACAGCCCAAACACCCAACAATGAACACGCTAATTGCACG    5041
PDS1                 ------------------------------------------------------------    2171
bd_1474_Extended     TGCTTTCCATACCTGATCTTGGTGGTGTTGCATCACAATAAGCTTCA-----GTGTCCGA    4225
bd_1474              ------------------------------------------------------------    1357

PDS1_Extended        TGGCGCCGAGAAGTCAAGGTGTTT--GTTGCGCACTAGCATCAAAT--------------    5085
PDS1                 ------------------------------------------------------------    2171
bd_1474_Extended     GGAATCAGGAAAGTGAAGATATGTATGGAGAGGATTACGGTTTTCTGACGTGAAGGTAAT    4285
bd_1474              ------------------------------------------------------------    1357

PDS1_Extended        --------CAAGTGCACTCTCGACGTCGGCGGG------------TGCATCCT---CTCC    5122
PDS1                 ------------------------------------------------------------    2171
bd_1474_Extended     GCCATAGTCATGGGTAGAATGGATATAGTGCAGTGCATGAAGGGCTGCTTTCATGTGACC    4345
bd_1474              ------------------------------------------------------------    1357

PDS1_Extended        AAGAAGTGATGTG---TCAGCACGTAATCAGCCGCACATGATGCATGCTACAACATCATC    5179
PDS1                 ------------------------------------------------------------    2171
bd_1474_Extended     AGGAGAAGGCTTGTTGTTGTACGATGATAAGAAGGAGTGGACTGGAGCAATGTCAGGACG    4405
bd_1474              ------------------------------------------------------------    1357
```

```
PDS1_Extended        GGCGTCTGCTGGCAACTGCTACACTAACATCCAACCATTCTCGGC--------CAATACC    5231
PDS1                 ------------------------------------------------------------    2171
bd_1474_Extended     AGTGCATTGTGCCAACCA--TCCAATAGATCCAACAAGACTCATATATGCCTCCGAGCAT    4463
bd_1474              ------------------------------------------------------------    1357

PDS1_Extended        TCCAGTTGGACCCGCAACT---TCTCATC---GGAAGCACCCTCACAACAACAACAACAA    5285
PDS1                 ------------------------------------------------------------    2171
bd_1474_Extended     CGTTGTTGGGCAGTCGAATGCGTTTCATTGTTGGAGTCAGCGATTGCATCAATAGGAATG    4523
bd_1474              ------------------------------------------------------------    1357

PDS1_Extended        CAACAACGTCCCAGCGCCATCGAAAAAGCCAACCCCCCCCTCCTAACACCCGGACACGCC    5345
PDS1                 ------------------------------------------------------------    2171
bd_1474_Extended     CCAGATCGATACGGAGTGATGGTAGGAGAAG---GAGTCTTAGAATCA-ACTTGGAATCG    4579
bd_1474              ------------------------------------------------------------    1357

PDS1_Extended        ATCCGCATCCTTCAAATCCACGGAGCCAACCAAGCACAACTCATCCGTCGTTCAGACTTT    5405
PDS1                 ------------------------------------------------------------    2171
bd_1474_Extended     TTCCGCAAGATTT--------GCTGCAAAACCAGCCTGATTCATGTGCACTTTAACTGTT    4631
bd_1474              ------------------------------------------------------------    1357

PDS1_Extended        GTCAAACTATG------TGAATCATCCCGTCCTGGAAAGAAACGGGATGCCAAGGTGATT    5459
PDS1                 ------------------------------------------------------------    2171
bd_1474_Extended     GATGGAGTAAGACTCCAAGAGAAGTGTGTGCCCAAAAACCATTCAACAACGCCCATGAAG    4691
bd_1474              ------------------------------------------------------------    1357

PDS1_Extended        GCTACGGCTTTGAGGGAGTTCAAACGCAATAACAAGTTTGTGCTTCATAGGGAGGGAGCG    5519
PDS1                 ------------------------------------------------------------    2171
bd_1474_Extended     TCAACGGCGATGAGGTTTGACAGAATTTCCTGGAACTTTGTC---TCTACAGAGTCGTCG    4748
bd_1474              ------------------------------------------------------------    1357

PDS1_Extended        AGTGCCGCTGTGGTGGGGATGATGAGGTCCAGTATTCCGAATTATAAGGTGGTTTATGGA    5579
PDS1                 ------------------------------------------------------------    2171
bd_1474_Extended     GGTG--------------AGTAATACACGAAGTCATCAACATACAAACCAAGTGTTAGTG    4794
bd_1474              ------------------------------------------------------------    1357

PDS1_Extended        AAGCCTAGGGTGGAGGCTGCGGTGTTTGTGGCGGAGCAGGTTTTGGATGAGAGTACGGGG    5639
PDS1                 ------------------------------------------------------------    2171
bd_1474_Extended     GTGCCGTGGACGG---------------TGGTACAGATGGATTG---TTAGGGTC----G    4832
bd_1474              ------------------------------------------------------------    1357

PDS1_Extended        TTGTACTTTGCAGTGAAGATTGAGGATGTGGATGGTGTGTTGAGTGAGTTGCAGCAGGGG    5699
PDS1                 ------------------------------------------------------------    2171
bd_1474_Extended     ATGATGTGTCGAGTAAAGAGGCATGGATCCGA-------TGGATTGGGTTGCAAACCCAT    4885
bd_1474              ------------------------------------------------------------    1357

PDS1_Extended        TTGAGTGAGTTGGAGGAACGGGGTATGAACGTTAGACTTGCAGCTCCAAATGAAGAAGAG    5759
PDS1                 ------------------------------------------------------------    2171
bd_1474_Extended     CGACA---TGAGGATGGAGCGGATCTTATCGTACCAGTGACGGGGGCT-ACGACGAAGAC    4941
bd_1474              ------------------------------------------------------------    1357

PDS1_Extended        GTCGAAGAGAGTGGTGATGAGTGCAAGATGCTACAAG-----------------------    5796
PDS1                 ------------------------------------------------------------    2171
bd_1474_Extended     CGTACAATGTCTTCTTGAGAAGCCAGTATTCGCCAGGCTTAGCGTCGGGATCACCGGATG    5001
bd_1474              ------------------------------------------------------------    1357

PDS1_Extended        -----------ATG-------CTCAGCGTGTCACGGAAGGATTGATGAAGCTATTGGTA-    5837
PDS1                 ------------------------------------------------------------    2171
bd_1474_Extended     GAGGGCGGACGATGGTGACTTCGTCAGGTGGCAAAATGCCATTGCAGAAGGCATTTTTGC    5061
```

```
bd_1474                 ------------------------------------------------------------   1357

PDS1_Extended           --------------AAGCGAAGATCTCGTCCAGAGAACAAGATGAAGAAGAGAGCAAAGC   5883
PDS1                    ------------------------------------------------------------   2171
bd_1474_Extended        AATCACCTTGCTTGAGGACACGACATTGTTCGATAGCCAAACTG-ACAAGGTACCGGAGG   5120
bd_1474                 ------------------------------------------------------------   1357

PDS1_Extended           GAGGGTATCTCAAACTCCTTCAACTGAAC-----GATGGCCCGGAAGACAGCACCTTGAA   5938
PDS1                    ------------------------------------------------------------   2171
bd_1474_Extended        GAGTCTTGA-CGAAGAACTGGTGCGTAACGTTGAGATTTCGACCAAGAGCGATCTTCGAG   5179
bd_1474                 ------------------------------------------------------------   1357

PDS1_Extended           ACTGGCAACACAAATCGCTCTTGCTATTGGGGGATCTGCTTCTGCGAGGAAGAACATTAT   5998
PDS1                    ------------------------------------------------------------   2171
bd_1474_Extended        GTTACCAAGGACAACGATGCGTGATTTTGC----------ACGAAGCGGCATGAGATTTT   5229
bd_1474                 ------------------------------------------------------------   1357

PDS1_Extended           TGCTCCGTATGCTGATGC-----CTTTTGGAGTGGAGAGGTTGACGAGTCTATTCTTCAG   6053
PDS1                    ------------------------------------------------------------   2171
bd_1474_Extended        CATCCTTTTTGATGGTAAGAACACACATGGTAGGTATGGCTTTGGGAGCACCTTTCTCAC   5289
bd_1474                 ------------------------------------------------------------   1357

PDS1_Extended           TTGGTTGCGGATGCTGAG----GCGAAGGAGTTGGCGGAGAAGGAGAGTGTCGAGGCTGC   6109
PDS1                    ------------------------------------------------------------   2171
bd_1474_Extended        GCAATGCCCGATATTAACCGACAGTGAGGCGTTGGAAGGTACCCATACTTTCAATAGAAC   5349
bd_1474                 ------------------------------------------------------------   1357

PDS1_Extended           CGCCGCCGCAGCTGCTGCTGCAGAAGAGGAGGAGGCATCTGAAGCGGGGGATGGTGAAGT   6169
PDS1                    ------------------------------------------------------------   2171
bd_1474_Extended        CCT-------TTTCTTCATAGTAACTTTGAAGCCATATTT--------------------   5382
bd_1474                 ------------------------------------------------------------   1357

PDS1_Extended           ACACGATGGCAGTGAAAATGATTCAGAACAATCAGAAGAGTCAACGGAGGAAACTAAGTC   6229
PDS1                    ------------------------------------------------------------   2171
bd_1474_Extended        -CTCGATCGGGATGGTTGTCGGCGAGAGCACGAAGGAGAGTGGCAGGA---C----AATC   5434
bd_1474                 ------------------------------------------------------------   1357

PDS1_Extended           TTAAGGTAATATACTGTGTAAGGTAGTACTACACCTGTTGTTATCCGATGGGGTCCGGTT   6289
PDS1                    ------------------------------------------------------------   2171
bd_1474_Extended        TTGATG-------------GAGGTTGATGGCACTGACGTGATGTGCAACTGGATCAAACG   5481
bd_1474                 ------------------------------------------------------------   1357

PDS1_Extended           GCTTGGCTGTCGAGGTAGGGGGGAGAGGTGTCAGCCTCGTCTACTGGGTGGTTGTAACAC   6349
PDS1                    ------------------------------------------------------------   2171
bd_1474_Extended        TGCTACCGGTAGAGGGAATGCAAAGG--AATGA-------TGAATGATGACCCGGAAGAA   5532
bd_1474                 ------------------------------------------------------------   1357

PDS1_Extended           GCCTTCCTCACGCTCCGCGCTCCCCCCCCACCAACCACAATCCAAAAGCAAACGCCGCCA   6409
PDS1                    ------------------------------------------------------------   2171
bd_1474_Extended        GAATGCCTTGAACACATAGATCAGTCCAATTGTATGGCAGGTCAGGAAGAGCAACCCCCC   5592
bd_1474                 ------------------------------------------------------------   1357

PDS1_Extended           CTGCATCGTCCTACCCTGTTGTGC----TTCCCTGACAGCGCCACTCTTAAATTGACCCT   6465
PDS1                    ------------------------------------------------------------   2171
bd_1474_Extended        ATTCCTCTTTGCGTGCATTTGGATGGCGTTTATACACAAACCGGTATCCATTGTCGGACT   5652
bd_1474                 ------------------------------------------------------------   1357

PDS1_Extended           TACACTGCTGATATATGCCATCGGTGCCGTATTGATTGAACGTCGTCCTACGATTTGCAA   6525
PDS1                    ------------------------------------------------------------   2171
```

```
bd_1474_Extended      TACCCAAGTAACCTTTGTAGTATTGTCCCTCATGCTCATAAGTGATCTTACTGTTCAGTT    5712
bd_1474               ------------------------------------------------------------    1357


PDS1_Extended         ACATTAAT---CAGAGAACATCAGCTCACACATCATCGGCATCGGTGCACGTCAACAACC    6582
PDS1                  ------------------------------------------------------------    2171
bd_1474_Extended      GAAGAAACGGTGGAAGGAGAGTGGAGGAGTCATCATCTGACGAGGGAGCTG---------    5763
bd_1474               ------------------------------------------------------------    1357


PDS1_Extended         ACCGCAATCATGCTTTCCGAAACAGGCAAAACAGTCCTAGAACAAAGTCCCTACACCCTC    6642
PDS1                  ------------------------------------------------------------    2171
bd_1474_Extended      --------GAGGCGTTGGGATTAAGTTGGGCATGTCCGAGAGGGAGATATAGGCAGTCGT    5815
bd_1474               ------------------------------------------------------------    1357


PDS1_Extended         AACATGGTGGAGTACGATTTCGAGAACTATGCTCTTACCCAAGGTAAGAAATCCGCCTAT    6702
PDS1                  ------------------------------------------------------------    2171
bd_1474_Extended      AGAGTCAT-----CAAATTGGATGAGATAGTCACTGTGAGAAGGAGAGGAC---------    5861
bd_1474               ------------------------------------------------------------    1357


PDS1_Extended         GACTTGCCCAAGTGATCTACGATGCAGTAGTTCAGCAGAGTAATTGTAA-ATGATCCTAA    6761
PDS1                  ------------------------------------------------------------    2171
bd_1474_Extended      GGGAGGGGGATGTCCATGACAGTGCCGGATTTGAGGA---TGTTTGTGTGGGGATCCACG    5918
bd_1474               ------------------------------------------------------------    1357


PDS1_Extended         CCTGTTAATCACTGCCATGAACGAATGACGTCTTGCGTCTTCACCCAAGGTATTCCCGAA    6821
PDS1                  ------------------------------------------------------------    2171
bd_1474_Extended      CGTTCTACGCGAGTGCCTGGAGGATAGGGTTCG-TCGAAGGAAGGAGTGGTGTCACGATA    5977
bd_1474               ------------------------------------------------------------    1357


PDS1_Extended         ACACTCGACCCTCTCCCCTACCGCAAGGCTGGCTCCG----TCTTCGCCACGGGCGAAGA    6877
PDS1                  ------------------------------------------------------------    2171
bd_1474_Extended      TAAGGAGCAGAACAAGCCCCCATCATAGGTGATTGTAGGATAAACCGAAACTGGCAAACG    6037
bd_1474               ------------------------------------------------------------    1357


PDS1_Extended         GGCCGAGCCGGTATTAACCTCCGCCAAGCACGATCCTAGGTTACGTACCGTGGTGTGTCG    6937
PDS1                  ------------------------------------------------------------    2171
bd_1474_Extended      AAACGGATCAA--TACGATAGCTGTCAGGCTCATAGTACTGTTTGTTGCGTGGATTGTAG    6095
bd_1474               ------------------------------------------------------------    1357


PDS1_Extended         TCACTGGTTGCGTGATTTGTGTATGAAGG---GAGCGGCGTGTGAGTTTTTGCATCAATA    6994
PDS1                  ------------------------------------------------------------    2171
bd_1474_Extended      ACCCGAATAGCATTGGATGTAGTGGAACGACCGATGACGATACCATCCATCGTATGAGCC    6155
bd_1474               ------------------------------------------------------------    1357


PDS1_Extended         TGATTTATCCAA---------GATGCCACTGTGTCGCCATG-------------------    7026
PDS1                  ------------------------------------------------------------    2171
bd_1474_Extended      TGATTCTTAGAACGAACACAAGAGCCGTCTTTGTTGTGGTGGAAGTAACAGACGGAGAAG    6215
bd_1474               ------------------------------------------------------------    1357


PDS1_Extended         ---GCGATCGGTGTAAAGTGAAGGATTGT----CCATTTAGACACATCAACGAAGCGGAT    7079
PDS1                  ------------------------------------------------------------    2171
bd_1474_Extended      ATAGGGATCCAAGTACGCTGATCAGCTTTCTCACCGTGTACGAGCATGAAGGGTGATGCT    6275
bd_1474               ------------------------------------------------------------    1357


PDS1_Extended         CGGTTGGAGTGTGTCTTTTACAGTCAGGGATTCTGCATTCACGGTCCATTCTGCAGGTAT    7139
PDS1                  ------------------------------------------------------------    2171
bd_1474_Extended      AATTTACCGTGGAACTTT-------------CCTGGAATCATG-TTCATCATGCG---AG    6318
bd_1474               ------------------------------------------------------------    1357


PDS1_Extended         AGGCACGTGAGGAGGGAGAGGGCCGACTTGCCAGTGGTGGCTGATTTTACATTGGGGTTG    7199
```

```
PDS1                 ------------------------------------------------------------   2171
bd_1474_Extended     CGGCATGAGAGAT-GGAGTGAAACCAAAAGCGACGCGGCATTTGTTTTTCAGTGAGGTAA   6377
bd_1474              ------------------------------------------------------------   1357

PDS1_Extended        AGTCAAATGCAAGCTGGGAAGGATGGAGTGTCGGCTGTACGACGACCTGCTCCGAAGCCA   7259
PDS1                 ------------------------------------------------------------   2171
bd_1474_Extended     GCA----CGAGACATGTGAACCATCACC--------TTCCAATGGCTTTCAACCAAACCG   6425
bd_1474              ------------------------------------------------------------   1357

PDS1_Extended        AATGAGTTCTT-------CA---AAGTTAGTTTGTGTAAGCACTTTTTGAGTG--GGGAG   7307
PDS1                 ------------------------------------------------------------   2171
bd_1474_Extended     TTGGAGGACTGGCGCCCTGCAGGAGCGGCAATAATGTTGGAATCTTTGTCTTGGAGGTAA   6485
bd_1474              ------------------------------------------------------------   1357

PDS1_Extended        TGTCCCTTTGG-AGATGGATGTCACTTTGCTCATG-------------------------   7341
PDS1                 ------------------------------------------------------------   2171
bd_1474_Extended     GCTCGGATGGCAGTACCGAAGAGCTTTTCGTCACAGTCACAGCGAAAACATTTGGCGAAA   6545
bd_1474              ------------------------------------------------------------   1357

PDS1_Extended        -GGGAGGCAGAGTTGAGGAGGCCAGGGCAAGAACCGACGAAGACTGTGGGGGAGGATGGG   7400
PDS1                 ------------------------------------------------------------   2171
bd_1474_Extended     CGGCCGGCTTCACTCCTGAATTGGTTGAAAGCAGCGATGATGTCTTGTGGAGAGAGCGAG   6605
bd_1474              ------------------------------------------------------------   1357

PDS1_Extended        GAAATTGCGGAGAAC---------ATGTTTGGGACGCAGGAGACGACTACGGTGGATTAT   7451
PDS1                 ------------------------------------------------------------   2171
bd_1474_Extended     CGAAGACCAAAGACCCAAGTGTATCTGGTTGCTCCATCTGCGAAGACAAGAGCATA----   6661
bd_1474              ------------------------------------------------------------   1357

PDS1_Extended        TATCAGGGTGGAGCTGCCGGTGGGGGTAAGCCAACG------CCCATTTTGGAACCCGAG   7505
PDS1                 ------------------------------------------------------------   2171
bd_1474_Extended     ---------CATCGTGCCACCAAGGGCAACCCCATCACCAAAACCAATGTCAACATGAAC   6712
bd_1474              ------------------------------------------------------------   1357

PDS1_Extended        AATGCATCATTCTTTATTCTGCGGGCGGCGACCTATCGTG-------ATTGTGCCATCTC   7558
PDS1                 ------------------------------------------------------------   2171
bd_1474_Extended     GATGTCCAAATACCTG---TAACGAGTACGATCGATTGGGGTACCACGTTTTGCCTTTGG   6769
bd_1474              ------------------------------------------------------------   1357

PDS1_Extended        TACTGTTCGTAATGAATGGATGGTACAACGAAAGCATGCAGAGGCAATCAAC---AAAGC   7615
PDS1                 ------------------------------------------------------------   2171
bd_1474_Extended     TATAGTAGCGTACGATCCAAGAGCTAATGGAAACTCACCAGAATCCGTCCACACTCCATC   6829
bd_1474              ------------------------------------------------------------   1357

PDS1_Extended        TCATGAGGGTGGGAGACAAGTCATGTTCTTCTTTACAGTTGGGGATTCCAACCACATTCA   7675
PDS1                 ------------------------------------------------------------   2171
bd_1474_Extended     GCGAGAAGTTTGGAGAAGGTGCTTGTAGTTCTTGAAACGACGGC-----AACCAGTGATA   6884
bd_1474              ------------------------------------------------------------   1357

PDS1_Extended        GGGGGCAGCACTGCTGACTTCCTCCGCATCGTAT------GTTGAATCGGCAGATCACAA   7729
PDS1                 ------------------------------------------------------------   2171
bd_1474_Extended     CGGTGGAGTTCCTCTGCTGACCAATGGGTTTTGGTGTCACTCCCATTGGGCGTGTCACAT   6944
bd_1474              ------------------------------------------------------------   1357

PDS1_Extended        ACAACCAAAGAACGGAGCTCCTACAAACGCCAACGCAAACAAAGATGGAGCATTCTGCTA   7789
PDS1                 ------------------------------------------------------------   2171
bd_1474_Extended     GGACGGATGGGA---------------GGAGGAGACGTACCGGGATGGTGGATGAGGCGG   6989
bd_1474              ------------------------------------------------------------   1357
```

```
PDS1_Extended        CCGATTCAA---GTGTGAATGGT---ATCGTACTTGTGAGCTGCCCATCTCCACTGCATT    7843
PDS1                 ------------------------------------------------------------    2171
bd_1474_Extended     AGGATGTCGGTGGTGGAAATGGTGGAAAGGAGCCGAGGTGCTGCGGATACGGGCTTGTCA    7049
bd_1474              ------------------------------------------------------------    1357

PDS1_Extended        GGAAGCTGCTCCCGATCTCTTGCTTCCCACATCAACTACACAATTCTGTCAAGATATGAA    7903
PDS1                 ------------------------------------------------------------    2171
bd_1474_Extended     ACAAGTGGGATGC-----GTTGTTTGATCTCATCAGGAGCCAAGTCAGACATTGAA----    7100
bd_1474              ------------------------------------------------------------    1357

PDS1_Extended        GTCTAAAACTGGGGAAGCAGTGATGAAGGCAATTTGGAACTCGCCTCTTTGTACGTTGTA    7963
PDS1                 ------------------------------------------------------------    2171
bd_1474_Extended     --------CGAGGCA------------CT-GAGTTGGAAGGAGGCTGGGTGTCAGGTGGA    7139
bd_1474              ------------------------------------------------------------    1357

PDS1_Extended        CGAGTCATGGAATGGAGGGGACTACGATGAAATGGGCGGGAAGGACTCACGTCCTCCTCC    8023
PDS1                 ------------------------------------------------------------    2171
bd_1474_Extended     A-TGACATGTAGTACCGGCGGAGATGACGAAGTGGCTGCATTGTCCACA--------ATG    7190
bd_1474              ------------------------------------------------------------    1357

PDS1_Extended        GCCAGTGGGAGATGCAATATTGACAGACTTTCGTTGTCCTTTGCCGGAGGAGGT---GTC    8080
PDS1                 ------------------------------------------------------------    2171
bd_1474_Extended     GCAACCGGGCGAGCAGAGA-CCCGAGGCTGAGCGTAATCTAACTCCGAAGCGATGACGTT    7249
bd_1474              ------------------------------------------------------------    1357

PDS1_Extended        GTGGCCTACGCTTCCA------GGTCCGGGGTTCATATTTGG-GTGCAACTCGCAGACTA    8133
PDS1                 ------------------------------------------------------------    2171
bd_1474_Extended     GGTGCCTAACGGTTCATATAAAAGATGACAGTCCAAGGTTGTGTTGACAGTGAGAATGAA    7309
bd_1474              ------------------------------------------------------------    1357

PDS1_Extended        TGGATGAATGTTTAGGAAGAGGATTGTTTGGTCTACCAGCTCATATGAAGGTGAGACTTT    8193
PDS1                 ------------------------------------------------------------    2171
bd_1474_Extended     TGAAGGAAAGTAGACATACATACCCCCCATGTCAGCGTCACCAATGAAACCGCAGCCTTT    7369
bd_1474              ------------------------------------------------------------    1357

PDS1_Extended        GTTGGGA----GGGCACAATGTAGTTTGAGACTATTTCCTTAAT-GCTCACTTGGACTCG    8248
PDS1                 ------------------------------------------------------------    2171
bd_1474_Extended     CTGGAGTAAGTGAGACCGGAGACTGTAGAGAGGATTGCGAAGAGCCGGAACATGCAGACA    7429
bd_1474              ------------------------------------------------------------    1357

PDS1_Extended        TTTGCAAC-----------TATTCCTCTAGATTGCTGCTGCAGGTATCCGACCAG-----    8292
PDS1                 ------------------------------------------------------------    2171
bd_1474_Extended     GTTCCGAACCATAATCTTCTTTCCATTGAGAGAGAAGACGGCAGTACCCACCCCACAGAC    7489
bd_1474              ------------------------------------------------------------    1357

PDS1_Extended        --GCTCTTCCATCTT-TCTCTACAACGTCAGTGAACGTCTCATCTTTGGTATCTTTGAAG    8349
PDS1                 ------------------------------------------------------------    2171
bd_1474_Extended     TGGAGCCAACGACTTGTTTCCCATGCGCACCTGCAG-GTGAGTGATGGGTTTGTAGGAGA    7548
bd_1474              ------------------------------------------------------------    1357

PDS1_Extended        CTCTTACACCGGCTATCATGAACATGGAACCAAAGGCATTCTCAAAGAATCCAAAAGCTC    8409
PDS1                 ------------------------------------------------------------    2171
bd_1474_Extended     CAAATACCGACTTGTCAGGGAACATGTGA------------TCAGTCGCACCAGAGTCAG    7596
bd_1474              ------------------------------------------------------------    1357

PDS1_Extended        AAACCTCACCTTTCCCTGTACAAATCCGTGTACGAATCAGTTTGGAGTGTCCTCCCGTCC    8469
PDS1                 ------------------------------------------------------------    2171
bd_1474_Extended     CAACCAAAAGTGCTGAGGATG----GAGTGGAGGTGGTGATGTGGGCTGGTGTAGTGTCC    7652
bd_1474              ------------------------------------------------------------    1357
```

```
PDS1_Extended        ATGAC---G--------------ACGATCCAGCGTTGA-----------ATGATGTGTTG    8501
PDS1                 ------------------------------------------------------------    2171
bd_1474_Extended     GAAATGGAGCCCTCCAATAAGAGACGGACCGACGTAGGGAGCCGAAACACAGTAGTGGTG    7712
bd_1474              ------------------------------------------------------------    1357

PDS1_Extended        AGAGCTCGTGTTGGCGGAAGAATTGGACCACTGACGTTTGCACAGTCGGAGGCATTGGCA    8561
PDS1                 ------------------------------------------------------------    2171
bd_1474_Extended     GAGGACGATGCCGATGGAGGAGATGGACGAATGGGAGAAGGACAAGAAGGGGCAGTGGAG    7772
bd_1474              ------------------------------------------------------------    1357

PDS1_Extended        AGTTTGATTGCCAACCAATGTGGGGCATTGAGTTACATGACAGAGTATCGCCGTAGTATT    8621
PDS1                 ------------------------------------------------------------    2171
bd_1474_Extended     GAGACAATGTACTTGGAACTGGAGTCGTTATGTTTAGCGACAACA--GAAGCATAAGAGG    7830
bd_1474              ------------------------------------------------------------    1357

PDS1_Extended        GAAACGAGACAGTTGAACGTTGCA---GCTCCTCCAATTGCATTGCCCCCTCGTAAAATT    8678
PDS1                 ------------------------------------------------------------    2171
bd_1474_Extended     CAGACGAAGTCGTATGATCTATAACAGCCTCATCGCCATCCCAAGCAAAGTCATTGAACT    7890
bd_1474              ------------------------------------------------------------    1357

PDS1_Extended        GGAACGGCTTAAGGGGGACAATCTTGACTGTTTAAAAGTACACATGAATTGATATTTTAA    8738
PDS1                 ------------------------------------------------------------    2171
bd_1474_Extended     CCTCAGGCGTCGAGTACCCCCTATCGTCC-------GCACCAGATGACG-----CGGTAA    7938
bd_1474              ------------------------------------------------------------    1357

PDS1_Extended        CGATACTTCTTGTTCTCCTCAGTATTCACAATATATGCCATGCTTCGAAAGAGCTTATAC    8798
PDS1                 ------------------------------------------------------------    2171
bd_1474_Extended     CCTGACTGACAG-----CTGGCTGTGTGCTAGACCCCCCCTCATTCAGCTGAGACACAGG    7993
bd_1474              ------------------------------------------------------------    1357

PDS1_Extended        A-TAAGGGAATGTCCACACTCGTCAGTGCCCATCCATGAGATTGAAGCCG----------    8847
PDS1                 ------------------------------------------------------------    2171
bd_1474_Extended     AGAAGAGGGAGGAGGTGAAGGGTCAGTGACAGTAGGTTGGACAGCAGCAGGTTGAGCGGA    8053
bd_1474              ------------------------------------------------------------    1357

PDS1_Extended        GCAACACTTTTAAATTCATCTCGGCAAAAACCTGCATCGTGCTGTTGCTGTCACTGAAAT    8907
PDS1                 ------------------------------------------------------------    2171
bd_1474_Extended     GGGAAGAGACGACTTGCCTTCGCGAGTGACGGTGA-----GATGTGCCTTTTTCAGCAAC    8108
bd_1474              ------------------------------------------------------------    1357

PDS1_Extended        GGGTCTGGAGTACTTTCTGTCGTGCTACCTCAGCTTTGTTGGTGTTTTCGTTTAGCTCCA    8967
PDS1                 ------------------------------------------------------------    2171
bd_1474_Extended     TGGCACTTTGTGATGTTGTGACAGCGACGTCGCATGCA--GATGCAGTTGGACTTCTTGA    8166
bd_1474              ------------------------------------------------------------    1357

PDS1_Extended        GCAGCGAGGTAAGCTCTT------------CGGGTACAATAACTGGGGAATGATATTGGA    9015
PDS1                 ------------------------------------------------------------    2171
bd_1474_Extended     GAAAGAAGGCAAGAACGTCCTCCTGCTTGAGGGCACCAATCCATTCCCACGGAGTCTTGA    8226
bd_1474              ------------------------------------------------------------    1357

PDS1_Extended        AAAGAG----AACGGAGCGTGTGGTTGGAGAAGAGGGTGCTGTTGATGCTTGACGGTCG    9071
PDS1                 ------------------------------------------------------------    2171
bd_1474_Extended     AGGTTGAAGGGGCCTGGCTACTGGAAGGAGTGGTGGAAGCTGCTGCAGCTGCAGGAGCAG    8286
bd_1474              ------------------------------------------------------------    1357

PDS1_Extended        CAGAGCACTCGGCAGAACTCCCTCACACCCTCGGGAGTTATGTTCTCGCTTGTGACGTCG    9131
PDS1                 ------------------------------------------------------------    2171
bd_1474_Extended     GAGCAGGAGTAGCCGCGGCAGATCGAGTCCCAGGAGCATCGG--------TCGAACAAGG    8338
bd_1474              ------------------------------------------------------------    1357
```

```
PDS1_Extended        AGTATTTCTAATGTTGTGTTACCAACCAAGCTAATTGCCAATCCCCTCAATACTTCATTG    9191
PDS1                 ------------------------------------------------------------    2171
bd_1474_Extended     AGCCCGATCAAACTTCTGGACAAGATCAGACACCTT------AGACATTGTCATGGAGTC    8392
bd_1474              ------------------------------------------------------------    1357

PDS1_Extended        GTAACCGCGTTGCAACCGAGATCGACAACTTCTAGCAATGTGTGGTTACTTTGGAA----    9247
PDS1                 ------------------------------------------------------------    2171
bd_1474_Extended     G---AAGGAGAGCTCACCAGAGCGAAA---ACGGTCAATGAGTGGCTCGTATTCAGCACG    8446
bd_1474              ------------------------------------------------------------    1357

PDS1_Extended        --GACCGTCGACAACAACTGCAATCCAGCTGCAGTAAGACTACGATTGTCGGATAA----    9301
PDS1                 ------------------------------------------------------------    2171
bd_1474_Extended     AAGACCCATTACAAAGAACATGACCTGAAGTTTGGAAGACACCAATTGTCCCGAGGAGGA    8506
bd_1474              ------------------------------------------------------------    1357

PDS1_Extended        ---ATCGAGA-----CGCCTCAACTTGGAGTTGGTTATTAATCATTTGTCAATGCAAGCA    9353
PDS1                 ------------------------------------------------------------    2171
bd_1474_Extended     AGTACGGAGGAACATCCCCTCCAACCGAGATTGGTAGGAAGTGACTGGTTCCGACACCCC    8566
bd_1474              ------------------------------------------------------------    1357

PDS1_Extended        TAGCTTCGTCTTCAAACTGATTGCTTGATACCACAAGAGTCTCCATTGCTGTGTTTGGAT    9413
PDS1                 ------------------------------------------------------------    2171
bd_1474_Extended     ----------TTGGCGGTTGTTGGGTAAATCACGGAGAATCTCAAATGTAACCATATCAT    8616
bd_1474              ------------------------------------------------------------    1357

PDS1_Extended        TTTGCAGTATGGAGGACCGAAAGAAGCGTGACCATCCTCGTCCTGTCGCTCTCCCGTTGT    9473
PDS1                 ------------------------------------------------------------    2171
bd_1474_Extended     CAC--------ACGGAGAGAAATGGTTGT-----TGATTGCATCAACCATCTCAAATCCT    8663
bd_1474              ------------------------------------------------------------    1357

PDS1_Extended        TACCGAGAGTTAAAGTCTTCAGGTTTGTGTTTGCAGCGAGCCCATTGGCCAAAGACAGAA    9533
PDS1                 ------------------------------------------------------------    2171
bd_1474_Extended     TTGCCATCATACAAATC---AC-----------------CCTTGTTCGCAAAGAGCAACT    8703
bd_1474              ------------------------------------------------------------    1357

PDS1_Extended        GTGATTCGTCGTCAAGACCACAGCATGAGGCGTTCAATTCTTTCAACGCTGAATTGGGCA    9593
PDS1                 ------------------------------------------------------------    2171
bd_1474_Extended     GTGCCTCGGAGCCATCGACA----------------------------------------    8723
bd_1474              ------------------------------------------------------------    1357

PDS1_Extended        TAACTTCTGAAAGCGCCTGCCAACCAATACTGGTTATATCAGGATTCAAGCTCATAGTCA    9653
PDS1                 ------------------------------------------------------------    2171
bd_1474_Extended     ------------------------------------AGGCAGGTGTGTACCTTCAACTCC    8747
bd_1474              ------------------------------------------------------------    1357

PDS1_Extended        ATGTTTTCAATGTGAAATTGCTGCTCAACCCATGTGATAATGCG----------GCAATA    9703
PDS1                 ------------------------------------------------------------    2171
bd_1474_Extended     AGCTGACGACTCTGAGCAGCGTTGTCAACAGTGGTGACCAGAGGGTCACTTGAGGAGACA    8807
bd_1474              ------------------------------------------------------------    1357

PDS1_Extended        TCTTCATCCGTCAGCCCAGACGAGATGATGATCAGGTGCTCCACTGTGGAAGTACTCAAA    9763
PDS1                 ------------------------------------------------------------    2171
bd_1474_Extended     TTGTAGTAGCTGCGGAGGAAGAAACGAGTGCTGAAGATCCAAGAGGAGTAAGAAG-----    8862
bd_1474              ------------------------------------------------------------    1357

PDS1_Extended        GCAGATGCAAGTGACTGCCATGTTC------TCACCGTTGAGAACAAGCAATTCGCG---    9814
PDS1                 ------------------------------------------------------------    2171
bd_1474_Extended     AGAGATCCGAAAGTTTCGGAAGCTCATACTTACGAGCTTGAATGGCATCGTAGAGGGAGA    8922
```

```
bd_1474               ------------------------------------------------------------    1357

PDS1_Extended         --AGGTTCAATGTTTGCAGTGTCGAGTTGCCACTCAAACCACCAGAGAGCATTGATATTC    9872
PDS1                  ------------------------------------------------------------    2171
bd_1474_Extended      GCTGGTGTGATGGAGACTGAATCAAATTGGTGGTAGAG-ACCTCGAGAGCAAGACCCTCC    8981
bd_1474               ------------------------------------------------------------    1357

PDS1_Extended         CATCGTCATTCATCTC--------A---------------TTATTGCTGAGGTCAAGCCG    9909
PDS1                  ------------------------------------------------------------    2171
bd_1474_Extended      AATTGGTCCTCCTCAGGGACACACAATGACAACGAAGCGGGGATGGTCGAAGCCACAGTG    9041
bd_1474               ------------------------------------------------------------    1357

PDS1_Extended         TTTCAATTGGGAATTTGAGCTTTTCAGCATGTTAGATACTCCCGTCATGAGTACACAGTA    9969
PDS1                  ------------------------------------------------------------    2171
bd_1474_Extended      CTAGAAGCCGAAGGTGGAGCCGGCGAGGGTGCAGGATGACGCTTAGAAGGG---------    9092
bd_1474               ------------------------------------------------------------    1357

PDS1_Extended         TGAACGATTTGTGAGCGAGCTTGCATCAGAGAGTACTAATGTTTCAAGAGCCGAAGCTGT    10029
PDS1                  ------------------------------------------------------------    2171
bd_1474_Extended      GGGAGGATAGAGGCAAGAACATCAATG-------TCGTCCACCTCATCATCCGAATCGGA    9145
bd_1474               ------------------------------------------------------------    1357

PDS1_Extended         CAATCCATTAGACAGGGCACCAATC-------------CCT--------C--TACCATCT    10066
PDS1                  ------------------------------------------------------------    2171
bd_1474_Extended      CCCCCCATGATCGGCGGCAAGACTCTGCGATGCCAAATGCTGAGCCGCGCGACGCCGTGT    9205
bd_1474               ------------------------------------------------------------    1357

PDS1_Extended         GAAAGGCTATCTTTGATTGTAAGCGTTCTCATATTGTTGTTACTGGAAAGAAGAGTCTCA    10126
PDS1                  ------------------------------------------------------------    2171
bd_1474_Extended      ACGTTGTGATTTCTTCTTCCGAGATTGATAAGATTGGTTGTTGTATAACGCCGATGGAGG    9265
bd_1474               ------------------------------------------------------------    1357

PDS1_Extended         ATTGCTTCGCAGCTTTTCTTTCCGACGTAGCTACCCATCACAGAGAGACTGGCTAGATTG    10186
PDS1                  ------------------------------------------------------------    2171
bd_1474_Extended      CTCCA----GCTGAACCCGGGCCGCAGAGGACAAAGATGACACTGAAG-TGCGTCCACTG    9320
bd_1474               ------------------------------------------------------------    1357

PDS1_Extended         GGGTTGGCAGCCAATGCCTCTATTATAAGCCTGTTTGCTTCGTCGGTACGAACAATACGT    10246
PDS1                  ------------------------------------------------------------    2171
bd_1474_Extended      GAGGGGGTAG----------------ACCGAAGGGAGC-GGGCAGGCGAGAACAACAAGT    9363
bd_1474               ------------------------------------------------------------    1357

PDS1_Extended         ACCT-----CCCATTGATGTTGGTGAATATTCACACGTTTCAGGGATGTG--TTTCCTAG    10299
PDS1                  ------------------------------------------------------------    2171
bd_1474_Extended      GCTGAGGCACGAAGTCGTCATCAGGAGGATCCACGCCAAGCCGAGAAGGGGGCTGAGTGG    9423
bd_1474               ------------------------------------------------------------    1357

PDS1_Extended         GAGCATCGATGCAATACATGATGGATGAATGACAAGACATTCCTCCAAGTCGAGTTCGGT    10359
PDS1                  ------------------------------------------------------------    2171
bd_1474_Extended      AAGGAAG--AGGACGAGTTGATG-------CAGAAGCTCGCGAGAGAAGCCAAGTAGGAG    9474
bd_1474               ------------------------------------------------------------    1357

PDS1_Extended         AAGATGCCCGTCTGCGAACAGGGGCTCAAGTTCTGACCAATCAGCTCCGATAAAGCACCT    10419
PDS1                  ------------------------------------------------------------    2171
bd_1474_Extended      GAGACGCGATGGGGTTGCGTGAAGGTGTATTTCC-----CACAGCCACGGAAAAGGATG-    9528
bd_1474               ------------------------------------------------------------    1357

PDS1_Extended         GCATAACTGCAGCCTTATGATTGATCTGTTGCGA-----GATACCCCCT---CACAAAAG    10471
PDS1                  ------------------------------------------------------------    2171
```

```
bd_1474_Extended      ACTGAGAGGGATGGTCATACTCCTCCTGCTGCAATTGGCGATGCAGGAGAGAGAGACGAC    9588
bd_1474               ------------------------------------------------------------    1357


PDS1_Extended         CTTCTAAACAGAAGTCTGAAGTGACCTGGTTCGTCTTCATCTCTATATCCACGGATGACC    10531
PDS1                  ------------------------------------------------------------    2171
bd_1474_Extended      CCTCATGGAAGATGACAGGGCTACTCTGGAGGAGATCGGCGAGAAGGTCGACGGGTGCAC    9648
bd_1474               ------------------------------------------------------------    1357


PDS1_Extended         AACTGTTTCAG--------TTGCGTGTTCTCCCCGATATGTCTCCCAAGTTCCCCCAGTG    10583
PDS1                  ------------------------------------------------------------    2171
bd_1474_Extended      CATTGTTTGCCGCAGAGAACTGGGCAATGGCCGCGTCCTGTGTCAGGGAAGTGGCGATCG    9708
bd_1474               ------------------------------------------------------------    1357


PDS1_Extended         CATCGTCTATTTCTTCTGTAAAACGTACCCTCAGTGAAGTAATGGTAG----------GA    10633
PDS1                  ------------------------------------------------------------    2171
bd_1474_Extended      CGGTGTG-AGTGATGACACATCCCGTGTCCTCAAGGTAGGCAATGAGAGATAGGGCATCA    9767
bd_1474               ------------------------------------------------------------    1357


PDS1_Extended         TCGTTGGATCTAATCTTCGACATGATGTCTTCGCTATAATCTACGCACGCATAGTCAAGT    10693
PDS1                  ------------------------------------------------------------    2171
bd_1474_Extended      GGGTCGGAGGGATCAGGGCAGAAGACGTGCCCGATAAGACCAGCACCGTCGGAGGGGGTG    9827
bd_1474               ------------------------------------------------------------    1357


PDS1_Extended         GACGAAACGAGATCGGTGAGCAAATCTTCATCATAAGATGCGTTGAGCCTCGCTATTTTG    10753
PDS1                  ------------------------------------------------------------    2171
bd_1474_Extended      GA---------GGTGATGATAACAGCACCTTCCTCGGACGTCATGGTCGGCACGACGAGG    9878
bd_1474               ------------------------------------------------------------    1357


PDS1_Extended         CTCAA-----GAGATCTTCAACCGCGTTGAAGCTATCCATTGTACTAGTTCTAGACGTCA    10808
PDS1                  ------------------------------------------------------------    2171
bd_1474_Extended      CCAGATCCAATGGGTAGGTAGGTAGAACGAGACTATAGAGTCG--AGGGTCTATAGGGAA    9936
bd_1474               ------------------------------------------------------------    1357


PDS1_Extended         AAG---------------------TGAGCCGTTGTGGTTGAGTATC----------GGTA    10837
PDS1                  ------------------------------------------------------------    2171
bd_1474_Extended      AGGAAGGATAGCGAGATAACGGTGCTCGCCATGGAGGGGGAGCACCGTCTCGTCGGTATC    9996
bd_1474               ------------------------------------------------------------    1357


PDS1_Extended         ATCTTG-GGCCGCGAATCCGTGCAGGGGAAGGAATGGAAGGGGAACGTGTGTGAAGAAAA    10896
PDS1                  ------------------------------------------------------------    2171
bd_1474_Extended      AAGTTCACGCGCTCAATTCGTGAGGTTCAGT----TTATGGCTA-GTTTTACGAACTACG    10051
bd_1474               ------------------------------------------------------------    1357


PDS1_Extended         GATACGTTGTTGTCTCCTCGTAATATCGTTTACCTGTAGAGGGGTCTATATGTAGAGACA    10956
PDS1                  ------------------------------------------------------------    2171
bd_1474_Extended      AGAAACGAATGGCCAGCATCTGAGAACTCAGTCAAGTCTAGGACCTACGAAGGACAGACA    10111
bd_1474               ------------------------------------------------------------    1357


PDS1_Extended         ATTTATGGTCCTCGATGGGAAGGTAGATAAACAGGGGACAACTGATTTTCATTCTGTACA    11016
PDS1                  ------------------------------------------------------------    2171
bd_1474_Extended      GGGTGGCGGCGGAGAGGGGA--------GAGGAGAGGAGAGATGTTGTCCCCTTA--ATG    10161
bd_1474               ------------------------------------------------------------    1357


PDS1_Extended         AACTGTTCCCGTTAGTAACACTTTTTATTTTAA-------------AGGGGTTAATTTAG    11063
PDS1                  ------------------------------------------------------------    2171
bd_1474_Extended      AAACACACACTTTAGAAGGAAACTTTATTGTACCCATACAATAGAAATAAATTATATTAG    10221
bd_1474               ------------------------------------------------------------    1357


PDS1_Extended         TAGACAATTTATGGACACGATC-TCCCCACGGTGCTCGTGTCAGCAAACCG-CAACGAAC    11121
```

```
PDS1                  ------------------------------------------------------------   2171
bd_1474_Extended      GGGATTAAAAATGGACAATAAGATGGATGCCGAGTTCATAACCAACAAAAAAAATGGTAC   10281
bd_1474               ------------------------------------------------------------   1357

PDS1_Extended         CCTCT-CTT--CCGTTCCTACCCTCCCTCCTCTCCACGCCCACCACCACTCAACATGTCG   11178
PDS1                  ------------------------------------------------------------   2171
bd_1474_Extended      CCTTCACCTTCACCCTGCTGCTCATCCTCCCTTATGCAAT-----CGACTCGAATCTTCA   10336
bd_1474               ------------------------------------------------------------   1357

PDS1_Extended         ACCCCCTCCCAGCTCCACA-----ACCCCAAAGCAGTATGGTTTCTCGGTGCACCAGTCT   11233
PDS1                  ------------------------------------------------------------   2171
bd_1474_Extended      ATGGCCTACTTTCATCAGTAAGCATCGGCGACATCACATCTTCTCTACGTCGAG----GT   10392
bd_1474               ------------------------------------------------------------   1357

PDS1_Extended         CCAAAGCAATAGCTGCATCAACCGCTGCGATTTATGTGTTGGCGGAGATGAACAAATGGC   11293
PDS1                  ------------------------------------------------------------   2171
bd_1474_Extended      CGAGCCCAGTCTTTG-ATGTCCGAGGAGGAAGTCAATGT------GGCTCGGAAGGTGGC   10445
bd_1474               ------------------------------------------------------------   1357

PDS1_Extended         ACGAGGCTCTTGTGTTCGGTGAGTGGTGCATGTGATCTGTGTTTAG--GGATGTGGCGTT   11351
PDS1                  ------------------------------------------------------------   2171
bd_1474_Extended      AACAGTCAGTGGATTGGG---------GTAATTGATCTTCGTCCAGAAGCTGAACCCGAT   10496
bd_1474               ------------------------------------------------------------   1357

PDS1_Extended         GCTATTGAGCT---------------TGTAGTTCTTGAAGCCGTTGTTTTGTTGTCTTTC   11396
PDS1                  ------------------------------------------------------------   2171
bd_1474_Extended      AGTATTACGTTGGTCCAAGACAATTCACAAGTTACTGAAAGAGCTGTATTCGTCTCTGAT   10556
bd_1474               ------------------------------------------------------------   1357

PDS1_Extended         TAATT-AA--TGTTCACAACTCTA-----------------------------GATACCT   11424
PDS1                  ------------------------------------------------------------   2171
bd_1474_Extended      ACAAGTACCGTCATCACAGCTGTTGGTAGTGAAGCTGAAACGATTCAACAAACAGCAGCT   10616
bd_1474               ------------------------------------------------------------   1357

PDS1_Extended         CCAAGATATTTGACCAGGCTCAGTTCTACAGAATCTTTGTTTGCAATTTGACGTTTGCGT   11484
PDS1                  ------------------------------------------------------------   2171
bd_1474_Extended      GCCTTCTCCATTGGCAAGCTCAGTTCAACCATATATCTTGTGATATCCTATGACTTT-GA   10675
bd_1474               ------------------------------------------------------------   1357

PDS1_Extended         CGATTGGAGAACTCGTCTTCGGGTTGTTGGCTCTGTGTCC----TTTGATGCGTCGATTT   11540
PDS1                  ------------------------------------------------------------   2171
bd_1474_Extended      GCATGGGA--AGACAGTGTTGCATCGTACACTGGGTGGCAGCAAGTTGATGAATTTTGTG   10733
bd_1474               ------------------------------------------------------------   1357

PDS1_Extended         GAACGTGAGGTATGTTACGTTG---TAACTTATTGATATCCGCTGAATTGCATTGATTCT   11597
PDS1                  ------------------------------------------------------------   2171
bd_1474_Extended      GAT-GGAATGCGAGAACGGTGGAACGAACTTCATGCACAAATCAATTTGGTACTTGTTCT   10792
bd_1474               ------------------------------------------------------------   1357

PDS1_Extended         CTTGCTTGTTTCTTCAACTCATACGAAACGTATCCAC---TTCTTCACTCACAAGATGGG   11654
PDS1                  ------------------------------------------------------------   2171
bd_1474_Extended      CGAGCCTTCTTCGGGAGCTGCAACATCGTCTTTTCTTTCCTGCCTCCCTTCAAGTACTGA   10852
bd_1474               ------------------------------------------------------------   1357

PDS1_Extended         TAGCAGAAAGTTTGGTGCATTCATAATCTATTCATCAGTCCTCTCAACGATATTTGAGTT   11714
PDS1                  ------------------------------------------------------------   2171
bd_1474_Extended      TGATAAGGAGAGCGAGGAAACAAACAACGAAACCTGGGGCATTGCAAACACGGTGGCGTT   10912
bd_1474               ------------------------------------------------------------   1357
```

```
PDS1_Extended        GGTATT-CTTCAATATATTCTTTGACACAGAGCGATACTCCGGTCCATATCCACAACTGG    11773
PDS1                 ------------------------------------------------------------    2171
bd_1474_Extended     GGGTCTCACAGCACACACTGAATG--------GAAT-------------------ACCAC    10945
bd_1474              ------------------------------------------------------------    1357

PDS1_Extended        GGGCAGTCCTTGCAATGTATCACAAATTCGCACCTCGGCTGCATCCTAAGTTCTTTGGAG    11833
PDS1                 ------------------------------------------------------------    2171
bd_1474_Extended     GGGCAGAGATCACCTTGTTG----------AAAG---GC---TGGTGACATACTTTCAGC    10989
bd_1474              ------------------------------------------------------------    1357

PDS1_Extended        TGCTAGGATACGACTTCTCTGAGAAGTCACTCACGTATGGTTTATGTGCTCA-AGTGATT    11892
PDS1                 ------------------------------------------------------------    2171
bd_1474_Extended     TGGGAGGAGAACAATACGC-------------CAATATCAATCCATTTCACAAGGTGGAC    11036
bd_1474              ------------------------------------------------------------    1357

PDS1_Extended        CTTTCGGGTGGTTTGAGCACGGCTATCCCTACAATCTTTGGCTTCATCTCGGGTATGCTT    11952
PDS1                 ------------------------------------------------------------    2171
bd_1474_Extended     GTTATTGGTGCTTGG-GATAGACAG-------------AAGGAGCATAGTGAATCAGATG    11082
bd_1474              ------------------------------------------------------------    1357

PDS1_Extended        AGTGTGAGCCTCAGTCAGCACGAGCTACCAGAAATCGTGTACACTGCTGCTGGAACTCTT    12012
PDS1                 ------------------------------------------------------------    2171
bd_1474_Extended     AAAA-------CAAGCAAAGCAAGCTAGACGAC----TCCA---CGATGAAGGTAATCTT    11128
bd_1474              ------------------------------------------------------------    1357

PDS1_Extended        GG-------TAAAGCATTTGTGGACGATGCACCCGCAATCATGATGGCACGGACAGTACA    12065
PDS1                 ------------------------------------------------------------    2171
bd_1474_Extended     ACGCCATGCCGAGTTGTTCAGTGACGTTGGTGCTGCAATCACCT--------CCAACAAA    11180
bd_1474              ------------------------------------------------------------    1357

PDS1_Extended        GCGT-GGAGGAAGGAATCAGCAAAGACGAGCTTCCACCGGTGCAAGGGGAGGAAGGGATG    12124
PDS1                 ------------------------------------------------------------    2171
bd_1474_Extended     GAGTCATTCCAAGCTGTCATCCAAGAGGTGTTTGAATCTGTTGGTGGAAAAGGATGCTTG    11240
bd_1474              ------------------------------------------------------------    1357

PDS1_Extended        C-----CACGGCGGGA------------------------G--CTC--------------    12139
PDS1                 ------------------------------------------------------------    2171
bd_1474_Extended     ACTTTCCTCTGAGATACTATGGTCGTTGACTACTAATTAACCTCTCTGATATATCCCTTG    11300
bd_1474              ------------------------------------------------------------    1357

PDS1_Extended        --------CGGCAGCGGCACTTCCTGCTGTGCAGCCTC-----------------CT       12171
PDS1                 ---------------------------------------------------------       2171
bd_1474_Extended     TCTATTCTTTGCAAACACACTTCAATCATCCCACGCTGACATGATCTTCGCAAAGCT       11357
bd_1474              ---------------------------------------------------------       1357
```

### Peptide Alignment (62.9% Sequence Identity)

```
PDS1       MIITNFILSTVLATSMAFQPHTPILSKPSFSNRVHRSPKIGSSNLV---MKDFP-KPNVE       56
PDS2       ---MKFLLPLLPAVAGAFSI-THLSQHPSLRMHQSLSTSLYSSSSSTSQRPRRPTPDRIR       56
               :*:*  : *.: **.  * : .:**:  :   * .: **.         *   .:.

PDS1       DTDNYRYAEAMSTSFKTSL---RVTNDSQKKKVAIIGGGLSGLSCAKYLSDAGHEPTVYE       113
PDS2       NTQNFKEAKELSQKFITDFQQLQKVGSGEPKRVAIFGGGLSGLSCAKYLSDAGHIPTLYE       116
           :*:*:: *: :* .* *.:   : ....: *:***:****************** **:**

PDS1       ARDVLGGKVSAWQDEDGDWIETGLHIFFGAYPNVMNMFAELGIHDRLQWKIHQMIFAMQE       173
PDS2       ARGVLGGKVSAWQDEDGDTVETGLHIFFGAYPNIHNLFDGLKIQDRLQWAPHRMTFAMQE       176
           **.*************** :*************: *:*  * *:*****  *:* *****
```

```
PDS1      LPGEFTTFDFIPGIPAPFNFGLAILMNQKMLTLGEKIQTAPPLLPMLIEGQSFIDAQDEL      233
PDS2      LPGQFTTFEFPAGVPAPLNMAAAILGNTEMLTLEEKIKMVPGLLPMLLEGQSFIDEQDEL      236
          ***:****:*  *:***:*:. *** * :**** ***: .* *****:******* ****

PDS1      SVTQFMRKYGMPERINEEVFIAMAKALDFIDPDKLSMTVVLTAMNRFLNESNGLQMAFLD      293
PDS2      SVLQFMRKYGMPERINEEIFIAMGKALDFIDPDLLSMTVVLTAMNRFINEADGSQTAFLD      296
          ** ***************:****.********* ************:**::* * ****

PDS1      GNQPDRWCTPTKEYVEARGGKVKLNSPIKEIVTNDDGTINHLLLRSGEKIVADEYVSAMP      353
PDS2      GNPPERLCQPMKESIEKKGGEVVCNSPVVEIQLNEESNVKSLKLANGTEITADYYVSAVP      356
          ** *:* * * ** :* :**:*  ***: **  *::..:: * * .* :*.** ****:*

PDS1      VDIVKRMLPTTWQTMPYFRQLDELEGIPVINLHMWFDRKLKAVDHLCFSRSPLLSVYADM      413
PDS2      VDVFKRLVPTQWSTMPYFRQLDELEGIPVINIQIWFDRKLNSVDGLCFSRSPLLSVYADM      416
          **:.**::** *.*****************:::******::** **************

PDS1      SVTCKEYEDPNKSMLELVFAPCSPIAGGNVNWIGKSDEEIIDATMGELARLFPTEIANDD      473
PDS2      STCCEEYASNDKSMLELVFAPCSPEAGSPLNWIAKPDSDIIDATMKELERLFPLEIGPDA      476
          *. *:** . :************ **. :***.* *.:****** ** **** **. *

PDS1      KWPATKMQGPNGQAKLEKYAVVKVPRSVYAAIPGE-------------------------      508
PDS2      PE--------EKRANVVKSTVVRVPRSVYAAVPGRNKYRPSQESPIENFIMAGDYATQKY      528
                    : :*:: * :**:********:**.

PDS1      ------------------------------------------------------------      508
PDS2      LGSMEGAVLSGKLAAEVICDKFMGRAERKGVKEVHSSVLTKQIEERTPAGIAMEKGRVSP      588


PDS1      --------------  508
PDS2      TSYGGGQQGGFENP  602
```

### 2)BCH

*T. pseudonana* “BCH” **(Appendix C)**

MHQLSRLLSKLFPVIATYIIAKYALPHINT
ILHCSNTLSNFVRITYTVLFAIAMEYISRYSHCYLWHGKFLWWINGSHHHQYPAVGSTPV
YGHNNPYVSPAIELNDAFAVFFATIATLAMWIGSEPPSTLTKDCSIGIGLGVTLYGLSYF
VGHDIVAHERLGKGVANALRRAFPYMEQCASVHIRYHHKLTKRSNDSDPYGAPYGFWLGP
SEVECLNRGQWYAPMPMSLKAISWIATL IFFASTIHSSLSPAAQAIVLLGCVGWCGSGTL
SDNTTTSRRIGKLLSLNWQSTPSRLMPHGLSGLISVGIGSYLIFGHSLVGDLKPYTMQQP
PYLIILYATATSWNALGGYMIVNTAPPNTRMLFRRCAILQVCLSYFIVRFLPHSSVLLIR
LESNTITTSLRCLDLIVTISAVVCTLSFFDAVVDMSKQSVVLGQSIAFGIIGILLLSVYP
IQLSLQGEEWWSCIQNRYPMQASGMIAYIYVPATVTFSLFLFGATLYQRKIMSASEYGII
SLMVILVCLLATVLSQEIHIPDVSTQRIYLPCEDPAMDSLEEKVLEALDFSRYARSILTT
VLGIKFESPA-

The underlined portion is what corresponds to BCHs from other organisms in the alignment below.

Plant and Green Algal BCH Sequences (non-heme di-iron):

*Vitis vinifera*, GenBank accession number AAM77007.

```
  1 matgisasln smscrlgrns ftatgpssvi slssfltpvt hlkgnifplq rrrslkvclv
 61 lekeiedgie ieddspessn raserlarkk aerytylvaa mmsslgitsm aivavyyrls
121 wqmeggeipv lemlgtfals vgaavgmefw arwahkalwh aslwhmhesh hrpregpfel
181 ndvfaiinav paisllsygl fnkglvpglc fgaglgitvf gmaymfvhdg lvhrrfpvgp
241 ianvpylrkv asahqlhhsd kfngvpyglf lgpmeleevg gmeelekeis rrikssdss
```

*Haematococcus pluvialis* (*Haematococcus lacustris*), GenBank accession number AAO53295.

```
  1 ittmlsklqs isvkarrvel arditrpkvc lhaqrcslvr lrvaapqtee avgtqqaaga
 61 gdehsadval qqldraiaer rarrkreqls yqaaaiaasi gvsgiaifat ylrfamhmtv
121 ggavpwgeva gplllvvggq lgmemyarya hkaiwhespl gwllhkshht prtgpfeand
181 lfaiikglpa mllctfgfwl pnvlgtacfg aglgitlygm aymfvhdglv srrfptgpia
241 glpymkrltv ahqlhhsgky ggapwgmflg pqelqhipga aeeverlvle ldwskr
```

*Capsicum annuum*, CAA70888.

```
  1 ttgryhyqlv wcqisfssts rtsyyrhspf lgpkptpttp svypitpfsp nlgsilrcrr
 61 rpsftvcfvl eddkfktqfe ageediemki eeqisatrla eklarkkser ftylvaavms
121 sfgitsmavm avyyrfywqm eggevpfsem fgtfalsvga avgmefwarw ahkalwhasl
181 whmheshhkp regpfelndv faiinavpai alldygffhk glipglcfga glgitvfgma
241 ymfvhdglvh krfpvgpvan vpylrkvaaa hslhhsekfn gvpyglflgp keleevggle
301 elekevnrrt ryikgs
```

```
T_pseudonana        ------------------------------------------------------------      0
V_vinifera          ---------matgisaslnsmscrlgrnsftatgpssvislssfltpvthlkgnifplqr      51
C_annuum            ttgryhyqlvwcqisfsstsrtsyyrhspflgpkptpttpsvypitpfspnlgsilrcrr      60
H_pluvialis         -------------------------------------ittmlsklqsisvkarrvelard      23


T_pseudonana        ------------------------------------------------------MHQLSR      6
V_vinifera          rrslkvclvle--------------kei--------ed-gieieddspessnraserlar      88
C_annuum            rpsftvcfvle--------------ddkfktqfeagee-diemkieeqisatrlaeklar      105
H_pluvialis         itrpkvclhaqrcslvrlrvaapqteeavgtqqaagagdehsadvalqqldraiaerrar      83
                                                                   .: :*

T_pseudonana        LLSKLFPVIATYIIAKYALPHINTI---LH---------CSNTLSNFVRITYTVLFAIAM      54
V_vinifera          kkaerytylvaammsslgitsmaivavyyrlswqme-ggeipvlemlgtfalsvgaavgm      147
C_annuum            kkserftylvaavmssfgitsmavmavyyrfywqme-ggevpfsemfgtfalsvgaavgm      164
H_pluvialis         rkreqlsyqaaaiaasigvsgiaifatylrfamhmtvggavpwgevagplllvvggqlgm      143
                       :     .: : :. .:  :  .    :              .    :   *   :.*

T_pseudonana        EYISRYSHCYLWHGKFLWWINGSHHHQYPAVGSTPVYGHNNPYVSPAIELNDAFAVFFAT      114
V_vinifera          efwarwahkalwhas--lwhmheshhr-pr--------------egpfelndvfaiinav      190
C_annuum            efwarwahkalwhas--lwhmheshhk-pr--------------egpfelndvfaiinav      207
H_pluvialis         emyaryahkaiwhesplgwllhkshht-pr--------------tgpfeandlfaiikgl      188
                    *  :*::*  :** .   *   . **  *                  :* ** **:: .

T_pseudonana        IATLAMWIGSEPPSTLTKDCSIGIGLGVTLYGLSYFVGHDIVAHERLGKGVANALRRAFP      174
V_vinifera          paisllsyglfnkglv-pglcfgaglgitvfgmaymfvhdglvhrrfpvgpia----nvp      245
C_annuum            paialldygffhkgli-pglcfgaglgitvfgmaymfvhdglvhkrfpvgpva----nvp      262
H_pluvialis         pamllctfgfwlpnvl-gtacfgaglgitlygmaymfvhdglvsrrfptgpia----glp      243
                     *      *    . :    .:* ***:*::*::*:. ** :. .*:  *        .*

T_pseudonana        YMEQCASVHIRYHHKLTKRSNDSDPYGAPYGFWLGPSEVECLNRGQWYAPMPMSLKAISW      234
V_vinifera          ylrkvasahqlhh--------sdkfngvpyglflgpmeleevggmeele--keisrriks      295
C_annuum            ylrkvaaahslhh--------sekfngvpyglflgpkeleevggleele--kevnrrtry      312
H_pluvialis         ymkrltvahqlhh--------sgkyggapwgmflgpqelqhipgaaeev--erlvleldw      293
                    *:.: : .*  :*        . .  *.*:*::*** *:: :
```

```
T_pseudonana      IATLIFFASTIHSSLSPAAQAIVLLGCVGWCGSGTLSDNTTTSRRIGKLLSLNWQSTPSR      294
V_vinifera        sdss--------------------------------------------------------      299
C_annuum          ikgs--------------------------------------------------------      316
H_pluvialis       skr---------------------------------------------------------      296

T_pseudonana      LMPHGLSGLISVGIGSYLIFGHSLVGDLKPYTMQQPPYLIILYATATSWNALGGYMIVNT      354
V_vinifera        ------------------------------------------------------------      299
C_annuum          ------------------------------------------------------------      316
H_pluvialis       ------------------------------------------------------------      296

T_pseudonana      APPNTRMLFRRCAILQVCLSYFIVRFLPHSSVLLIRLESNTITTSLRCLDLIVTISAVVC      414
V_vinifera        ------------------------------------------------------------      299
C_annuum          ------------------------------------------------------------      316
H_pluvialis       ------------------------------------------------------------      296

T_pseudonana      TLSFFDAVVDMSKQSVVLGQSIAFGIIGILLLSVYPIQLSLQGEEWWSCIQNRYPMQASG      474
V_vinifera        ------------------------------------------------------------      299
C_annuum          ------------------------------------------------------------      316
H_pluvialis       ------------------------------------------------------------      296

T_pseudonana      MIAYIYVPATVTFSLFLFGATLYQRKIMSASEYGIISLMVILVCLLATVLSQEIHIPDVS      534
V_vinifera        ------------------------------------------------------------      299
C_annuum          ------------------------------------------------------------      316
H_pluvialis       ------------------------------------------------------------      296

T_pseudonana      TQRIYLPCEDPAMDSLEEKVLEALDFSRYARSILTTVLGIKFESPA   580
V_vinifera        ----------------------------------------------   299
C_annuum          ----------------------------------------------   316
H_pluvialis       ----------------------------------------------   296
```

Percent Identity Matrix

```
1: T_pseudonana  100.00   25.73   26.21   27.05
2: V_vinifera     25.73  100.00   68.56   42.96
3: C_annuum       26.21   68.56  100.00   42.45
4: H_pluvialis    27.05   42.96   42.45  100.00
```

### 3)LTL

Predicted protein sequence alignment of the two *T. pseudonana* LTLs.

```
LTL1      --MKFTT--ALA--VLCWTSVTNAFVPSSFTSPA------LKNEQQQVRASSPLYALDTK      48
LTL2      MCTKLSSRRTLLALYFAFTGCTAFQLPSATPSRASITKAYSTHLDKEIKSKTPLVNPSKI      60
             *:::  :*    :.:*. *   :**:  * *       .: ::::::.:**   ..

LTL1      EKEETTTATSASSTDTSSTPAA--AATEESEGLPWWWEYI--------------WK----      88
LTL2      YTQADIDTLDLSSYEN-ELLAAWDTDSSLQRGFDWEIEKLRRNFAGLRQREDGQWVRKPS      119
           .:    : . ** :. .  **  : :. ..*: *  * :              *

LTL1      -LPVMQPAE----------------PGTDIIFADSARVLRT-------------------      112
LTL2      LFDFLVTNTPSNVVGVSNTGERYESPPKPVNMLDVGLLITKNLLNTLGFGPSLGMAAVPD      179
           : .:                    * . : : * . :: .

LTL1      -NIEQIYGGFPSLDQCPLAEGEITDIADGTMFIGLQRYQQQYGSPYKLCFGPKSFLVISD      171
LTL2      AVIQKYEGSFFSF-IKGVLGGDLQTLAGGPLFLLLAKYYQDYGPIFNLSFGPKSFLVISD      238
```

```
                *::  *.* *:    :  *::  :*.* :*: * :* *:**  ::*.***********

LTL1      PVQAKHVLRDAN-TLYDKGILAEILKPIMGKGLIPADPETWSVRRRAIVPAFHKAWLNHM      230
LTL2      PVMARHILRDSSPEQYCKGMLAEILEPIMGDGLIPADPKIWKVRRRAVVPGFHKKWLNNM      298
          ** *:*:***:.   * **:*****:****.*******: *.*****:**.*** ***:*

LTL1      VGLFGYCNEGLIASLEEAAKKNDAPNGQQGGKIEMEEKFCSVALDIIGLSVFNYEFGSVS      290
LTL2      VTLFGDCGERLVNDLDARAT--------AKTPVDMEERFCSVTLDIIGKAVFNYDFGSVT      350
          * *** *.* *: .*:  *.            ::***:****:***** :****:****:

LTL1      EESPVIKAVYSALVEAEHRSMTPAPYWDLPFANEVVPRLRKFNSDLKVLDDVLTDLIDRA      350
LTL2      KESPIVKAVYRVLREAEHRSSSFIPYWDLPYADKWMGGQVEFRKDMGMLDDILTKLINRA      410
          :***::**** .* ****** :  ******:*:: :    :*..*: :***:**.**:**

LTL1      KNSRQVEDIEELEKRDYANVKDPSLLRFLVDMRGADIDNKQLRDDLMTMLIAGHETTAAV      410
LTL2      IETRDEASVEELEDRDVG--DDPSLLRFLADMRGEDLTSKVLRDDLMTMLIAGHETTAAM      468
           ::*:  .:****.** .  .********.**** *: .* *****************:

LTL1      LTWALFELTKHPE-QMAKVRAEIDSVLGDR-TPTYDDIKEMQYLRLVVAETLRLYPEPPL      468
LTL2      LTWTVFGLVSNDSGLMKEIQAEVRTVMGDKLRPDYDDIAKMKKMRYALIEALRLYPEPPV      528
          ***::* *..: .  * :::**: :*:**:  * **** :*: :* .: *:********:

LTL1      LIRRCRTENKLPKGGGR---EATVIRGMDIFLSLYNLHHDERFWPEPNEFKPERWESKYI      525
LTL2      LIRRARSEDNLPAGGSGLSGGVKVLRGTDIFISTWNLHRAPEYWENPEKYDPTRWERRFK      588
          ****.*:*::** **.      ..*:** ***:* :***:  .:* :*:::.* *** ::

LTL1      NPEVPEWAGYDPAKWINTNLYPNEVASDFAYLPFGGGARKCVGDEFATLEATVTLAMLLR      585
LTL2      NPGVKGWNGYDPEKQSESSLYPNEITADYAFLPFGAGKRKCIGDQFAMLEASVTLAMIIN      648
          ** *  * **** *  ::.*****:::*:*:****.* ***:**:** ***:*****::.

LTL1      RFEFEFDSAKLAASKIDIMDHPEDLEHAVGMRTGATIHTRKGLHMVIRKREL--------      637
LTL2      KFDFTLVGS----------------PKDVGMKTGATIHTMNGLNLVVSRRSEDNPIPETN      692
          :*:* : .:                : ***:******* :**::*: :*.

LTL1      --------------------------------------------      637
LTL2      DYWIQQHLSRGLNVNGRPYSTNEDAAWTASSRDKNEGVVSRLVN      736
```

Percent Identity Matrix

```
1: LTL1        100.00   47.78
2: LTL2         47.78  100.00
```
