## Appendix E - Plasmid Maps and Primers for "Overexpression of *Thalassiosira pseudonana* violaxanthin de-epoxidase-like 2 (VDL2) increases fucoxanthin while stoichiometrically reducing diadinoxanthin cycle pigment abundance"

1) VDL2 overexpression, nitrate reductase promoter and terminator.

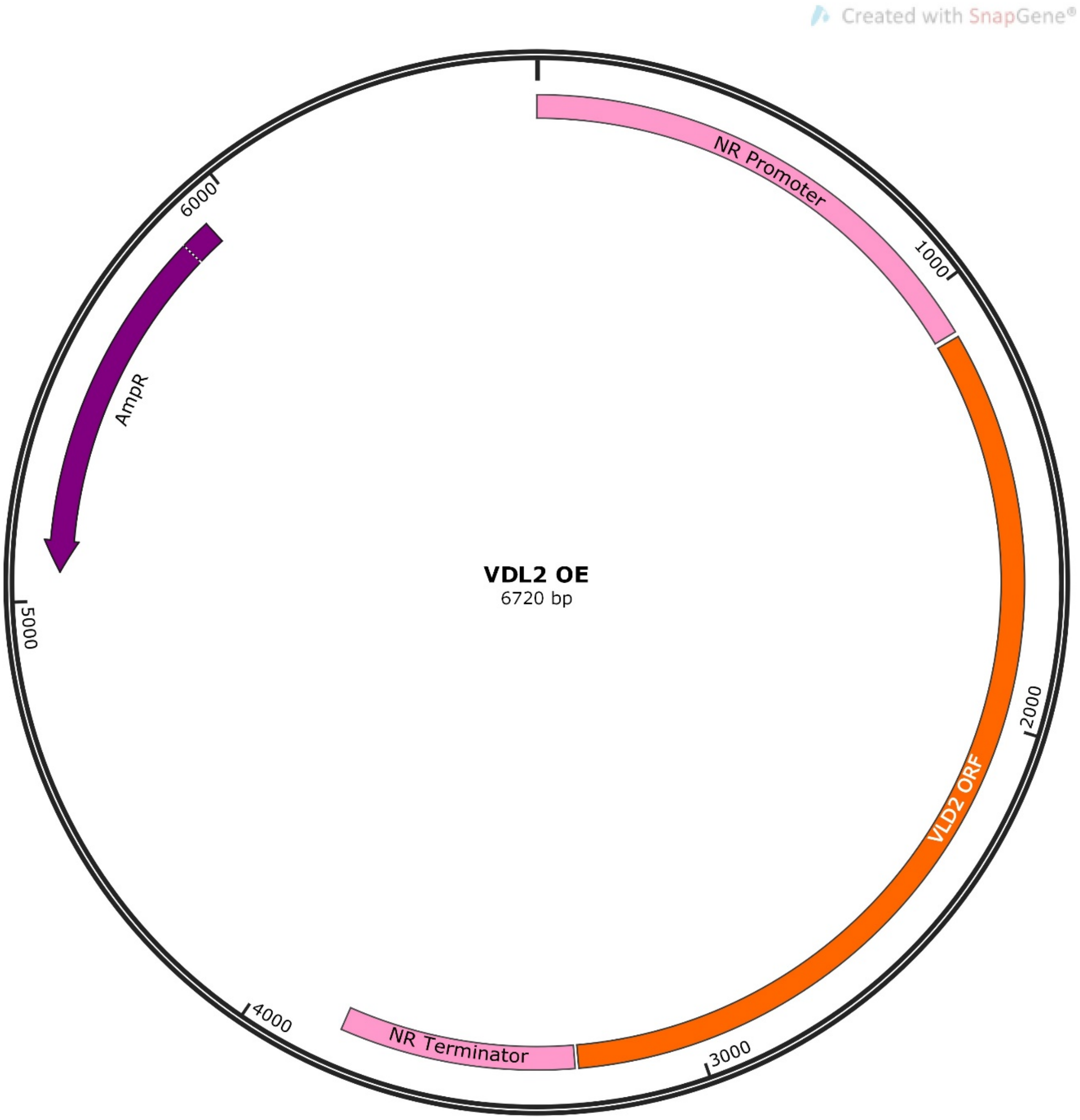


Primers for amplifying the VDL2 open reading frame:

Fwd: 5'ggggacaagtttgtacaaaaaagcaggctATGTCGGCATCATCATCATCAAC3'

Rev: 5'ggggaccactttgtacaagaaagctgggtaTCACCAATTCTTCTCTTTGATATAAT3'

Lower case sequences add Att sites for Gateway cloning.

2) NAT, acetyl CoA carboxylase promoter and terminator.

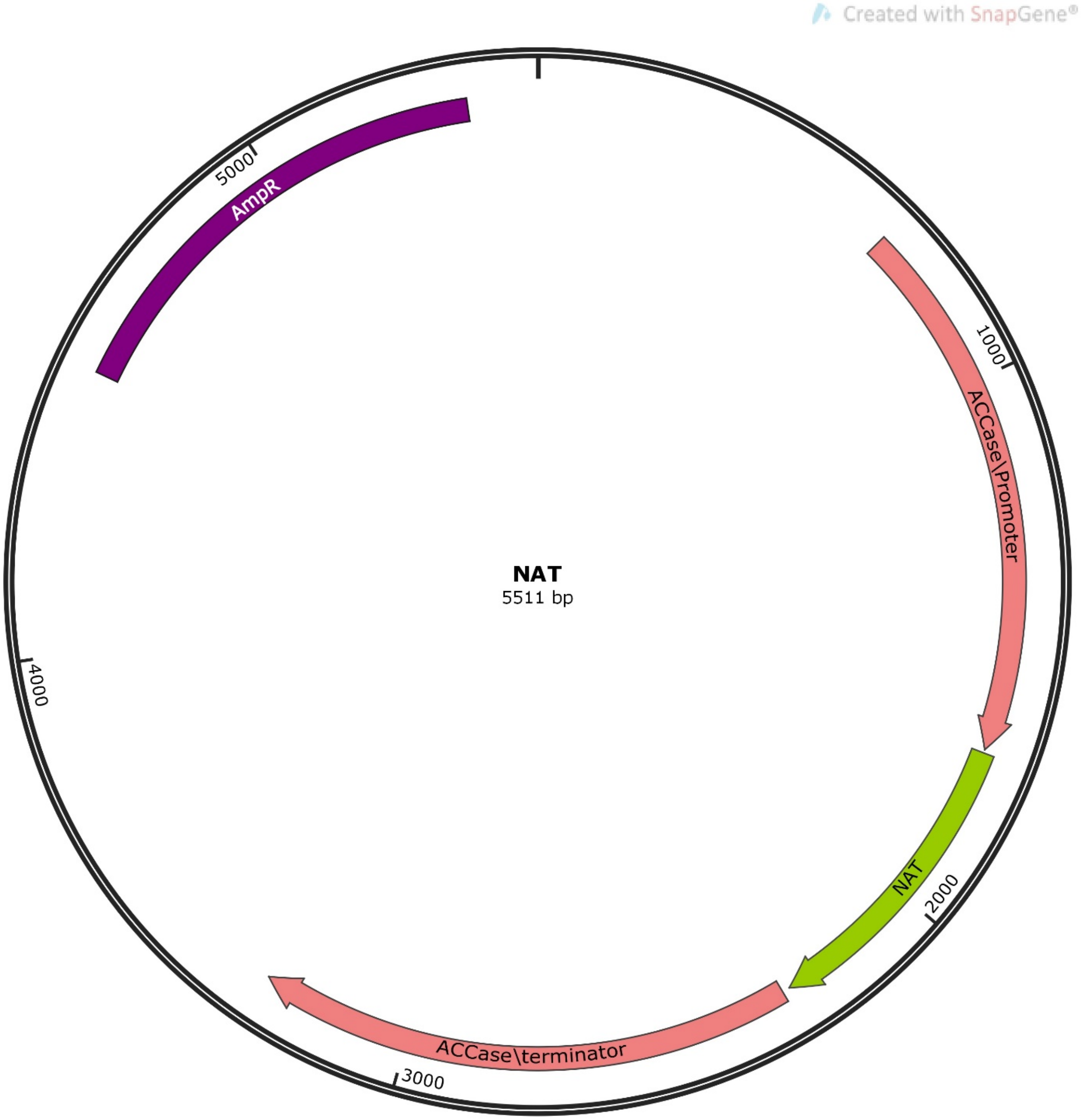

3) LTL KD, ribosomal protein 41 promoter and terminator, NAT on the same transcript.

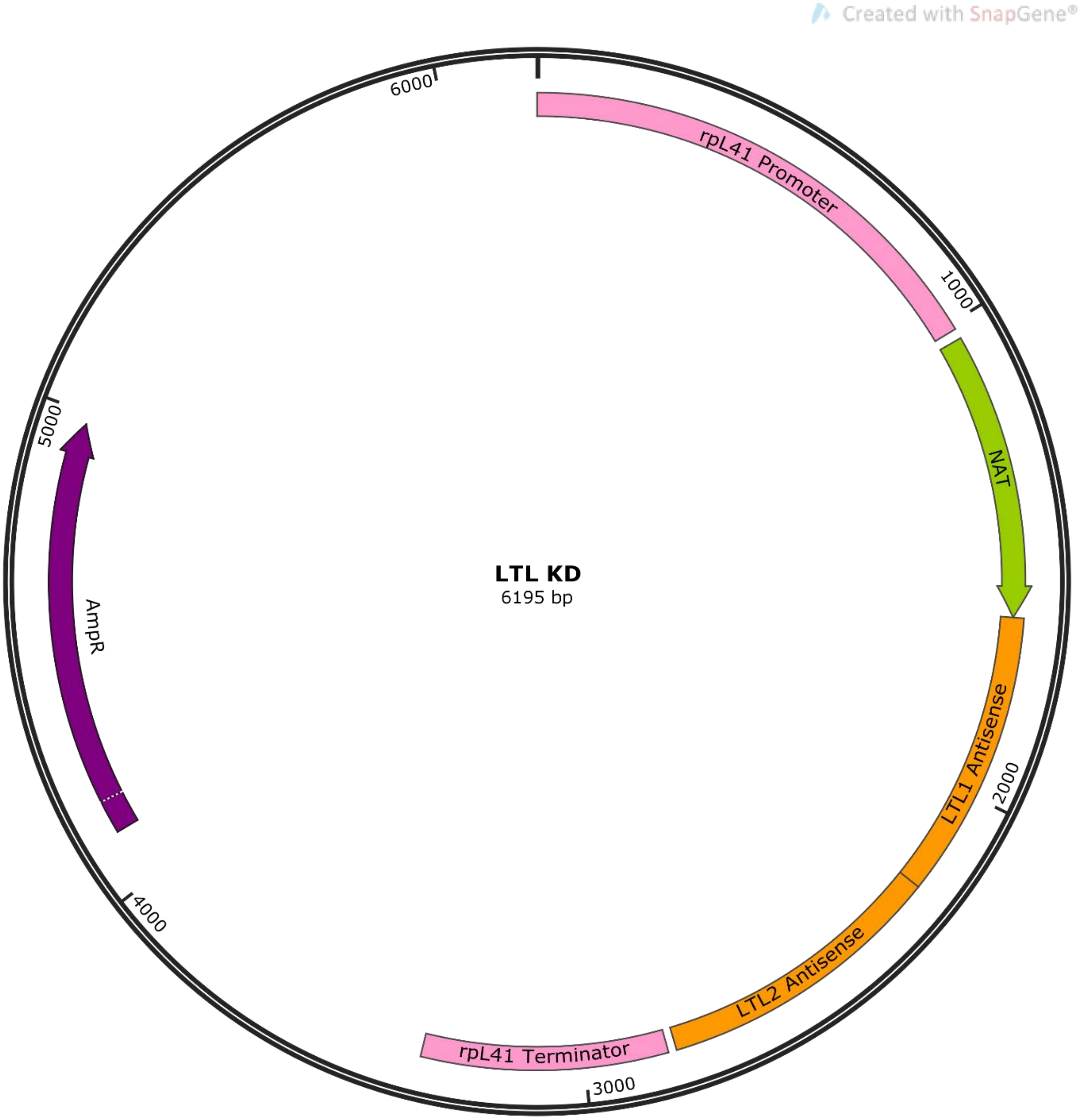


Primers to amplify antisense regions:

Fwd, LTL1: 5'ggggacaagtttgtacaaaaaagcaggctTCCAATGACCATAGTTGTTG3'

Rev, LTL1: 5'ggggacaacttttgtatacaaagttgTGCAGAGACTTTGAGGTTG3'

Fwd, LTL2: 5'ggggacaactttgtatacaaaagttgCTCTTATATCCTAGATAACTTT3'

Rev, LTL2: 5'ggggaccactttgtacaagaaagctgggtGGGTGGTGTCAAAGTATTG3'

Lower case sequences add Att sites for Gateway cloning.

4)VDL1 or VDL2 KD, acetyl CoA carboxylase promoter and terminator, NAT on the same transcript.

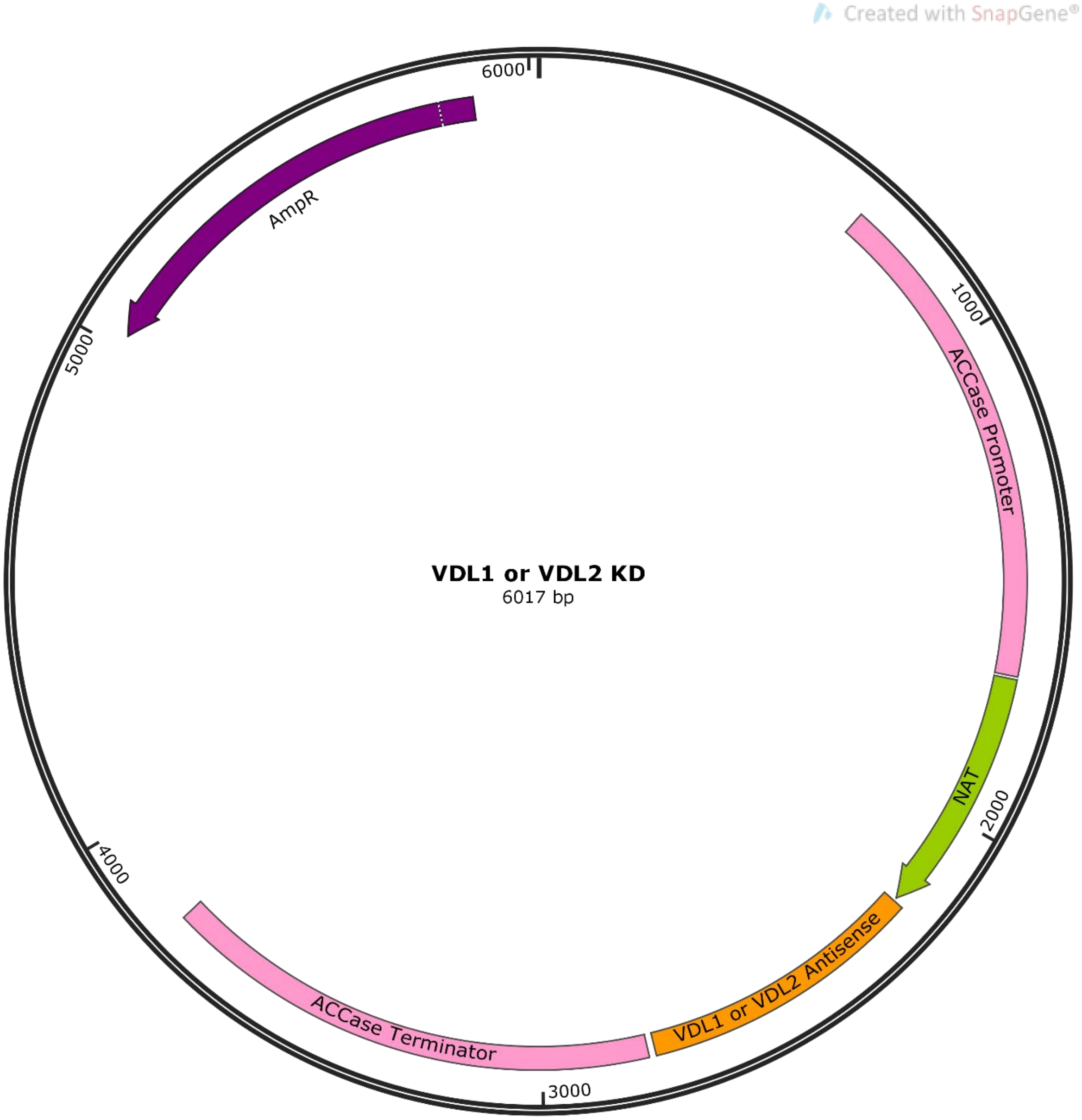


Primers to amplify antisense regions:

VDL1, Fwd: 5'ggggacaagtttgtacaaaaaagcaggctCAGTTGTCCAAATCCTCGCTCC3'

VDL1, Rev: 5'ggggaccactttgtacaagaaagctgggtaACAGTCGTCCTAGGCACCATTG3'

VDL2, Fwd: 5'ggggacaagtttgtacaaaaaagcaggctCCCAAACACTTGGCAGTACACG3'

VDL2, Rev: 5'ggggaccactttgtacaagaaagctgggtaTCAATGCCTACTCGGTCGATAC3'

Lower case sequences add Att sites for Gateway cloning.
